# Exploring Potential Signals of Selection for Disordered Residues in Naturally Occurring Prokaryotic and Eukaryotic Proteins

**DOI:** 10.1101/2020.03.10.979443

**Authors:** Arup Panda, Tamir Tuller

**Affiliations:** Department of Biomedical Engineering, Tel Aviv University, Tel Aviv 69978, Israel

**Keywords:** Intrinsically disordered protein, Comparative genomics, Gene function, Proteome evolution

## Abstract

Intrinsically disordered proteins (IDPs) were recognized as an important class of proteins in all domains of life for their functional importance. However, how nature has shaped the disorder potential of prokaryotic and eukaryotic proteins is still not clearly known. Randomly generated sequences are free of any selective constraints thus these sequences are commonly used as null models. Considering different types of random protein models here we seek to understand how disorder potential of natural eukaryotic and prokaryotic proteins differs from random sequences. Comparing proteome-wide disorder content between real and random sequences of 12 model organisms we noticed that while in eukaryotes natural sequences tend to be more disordered than random sequences prokaryotes follow an opposite trend. By analyzing position-wise disorder profile, here we showed that there is a general trend of higher disorder near the N and C-terminal regions of eukaryotic proteins as compared to the random models; however, either no or a weak such trend was found in prokaryotic proteins. Moreover here we showed that this preference is not due to the biases either in the amino acid or nucleotide composition or other factors at the respective sites. Instead, these regions were found to be endowed with a higher fraction of protein-protein binding sites suggesting their functional importance. Here, we proposed various explanations for this pattern such as improving the efficiency of protein-protein interaction, ribosome movement, and post-translational modification, *etc.* However, further studies are needed to clearly understand the biophysical mechanisms causing the trend.

## Introduction

Until the early 1990s, molecular biology studies were mainly focused on globular proteins with the view that protein function is inherently encoded in their folded 3D structures. However, recent studies suggested that a large number of naturally occurring proteins do not fold into specific three-dimensional structures in their native state [1–6]. These proteins are commonly known as intrinsically disordered proteins (IDPs) or intrinsically unstructured proteins (IUPs).

Disordered proteins follow unique sets of biophysical characteristics that are very distinct from that of well structured globular proteins. At the primary structure level, IDPs are enriched by the presence of numerous uncompensated charged groups resulting in low mean hydrophobicity and a high net charge at neutral pH [1–6]. Disordered regions are found to be encoded mainly by polar and charged (specifically, alanine, glycine, arginine, glutamine, serine, glutamic acid, lysine, and proline) amino acids and are devoid of hydrophobic and aromatic amino acids [1–5]. Due to the relatively higher rates of amino acid substitutions and fixation of insertion and deletions, disordered regions are known to evolve at significantly higher rates than ordered regions [7–10].

Despite their unordered structure, IDPs take central roles in several biological processes [4–6]. IDPs can compensate the functions of globular proteins and could carry out several other functions those can‟t be achieved by using globular proteins [4–6]. Specifically, IDPs were found to play significant roles in the processes like signaling, transcription and various regulatory processes, such as control of cell division, apoptosis and post-translational modifications, *etc.* [4–6].

Due to their inherent structural flexibility, IDPs could bind with a large number of partner proteins [1], [4−6]. Thus, IDPs could serve as the structural basis for binding promiscuity of hubs proteins (proteins those bind with multiple partners in protein-protein interaction networks) [5,6,9]. IDPs often act as flexible linkers between globular domains facilitating their binding diversity. Another important feature of IDPs is that many of these proteins can undergo coupled folding and binding process *i.e*. they can adopt stable secondary structures upon binding with partner molecules [4–6]. Binding of disordered proteins with their partner molecule may also be mediated by molecular recognition features or MoRFs [2,5,6,11,12]. Computational predictions suggested that disordered proteins, in general, are highly enriched with MoRFs indicating their high interaction promiscuity [5,6,11,12]. Considering their importance, previously a number of initiatives have been undertaken to estimate their abundance in different domains of life. These studies suggested that disordered residues, in general, are more prevalent in complex organisms such as multi-cellular eukaryotes than unicellular bacterial and archaeal genomes [13–15]. Disordered residues were suggested to be necessary for complex organisms to sustain their functional and regulatory complexities [13]. IDPs were also considered to play significant roles in the evolutionary adaptation of various prokaryotic and eukaryotic organisms [16, 17].

The sequences those are found in nature are considered to be only a small subset of all possible sequences extremely refined and edited by the million years of evolutionary constraints [18]. Randomly generated artificial sequences were considered as an important tool to understand the direction of this refinement. Natural sequences evolve under constraints imposed by their structural and functional requirements [18]. Random DNA sequences being free from such pressure were widely used as null models to explore the extent of selection in different traits of naturally occurring protein and DNA sequence [19, 20]. Random DNA sequences were often used for exploring the evolutionary signatures those discriminate real sequences from random copolymers [18, 21]. Further, random sequences were suggested to provide important insight regarding the structural and functional basis of extant protein and DNA sequences [18–20], [22].

Earlier a number of studies have been conducted in order to understand how the structural potential of naturally occurring proteins differs from that of randomly generated sequences [23, 24]. Understanding how the disorder potential of natural protein sequences differs from that of random sequences could provide insights into their evolutionary history. Moreover, since previous studies analyzed the disorder level of complete proteins there is no clear understanding of the regions in the proteins those might be under evolutionary pressure for strong or weak folding and their functional implications. Therefore, in this analysis, we took a major initiative to explore these crucial aspects. Here our major objectives are two folds (*i*) first to test whether there is any preference or abomination for disordered residues in naturally occurring eukaryotic and prokaryotic proteins as compared to random expectations, and (*ii*) to check whether there is any site-specific variation in this preference or abomination for disordered residues along the protein length.

In order to test these aspects here we generated three kinds of random protein models: (*i*) that preserve the fundamental properties of real proteins such as their overall amino acids frequencies and length, (*ii*) that preserve the characteristics of terminal regions, and *(iii)* that preserve position-wise amino acid frequencies at each position of the naturally occurring proteins (**Figure 1**). Order or disorder status of both real and random protein models was predicted based on the mutual agreement among four disorder prediction algorithms specifically chosen for this analysis. Further, we checked all the major results with another set of algorithms considering majority-vote consensus approach. To get a general evolutionary trend, here we first compared the overall disorder propensities of real and random proteins of each species then compared their disorder scores position-wise. Considering both these approaches, here we noticed that naturally occurring eukaryotic proteins show an overall higher percentage of disordered residues (here referred as protein disorder content) as compared to the corresponding random sequences and this preference is more pronounced at the terminal regions of eukaryotic proteins than the other regions. Considering several factors which may cause this trend here we argued that this is a guanine trend irrespective of selection for any other traits. In the next few pages of this article we emphasis on the functional significance of the observed trends. We believe that our study will help better understanding the forces shaping disorder propensity of natural sequences.

**Figure 1.**
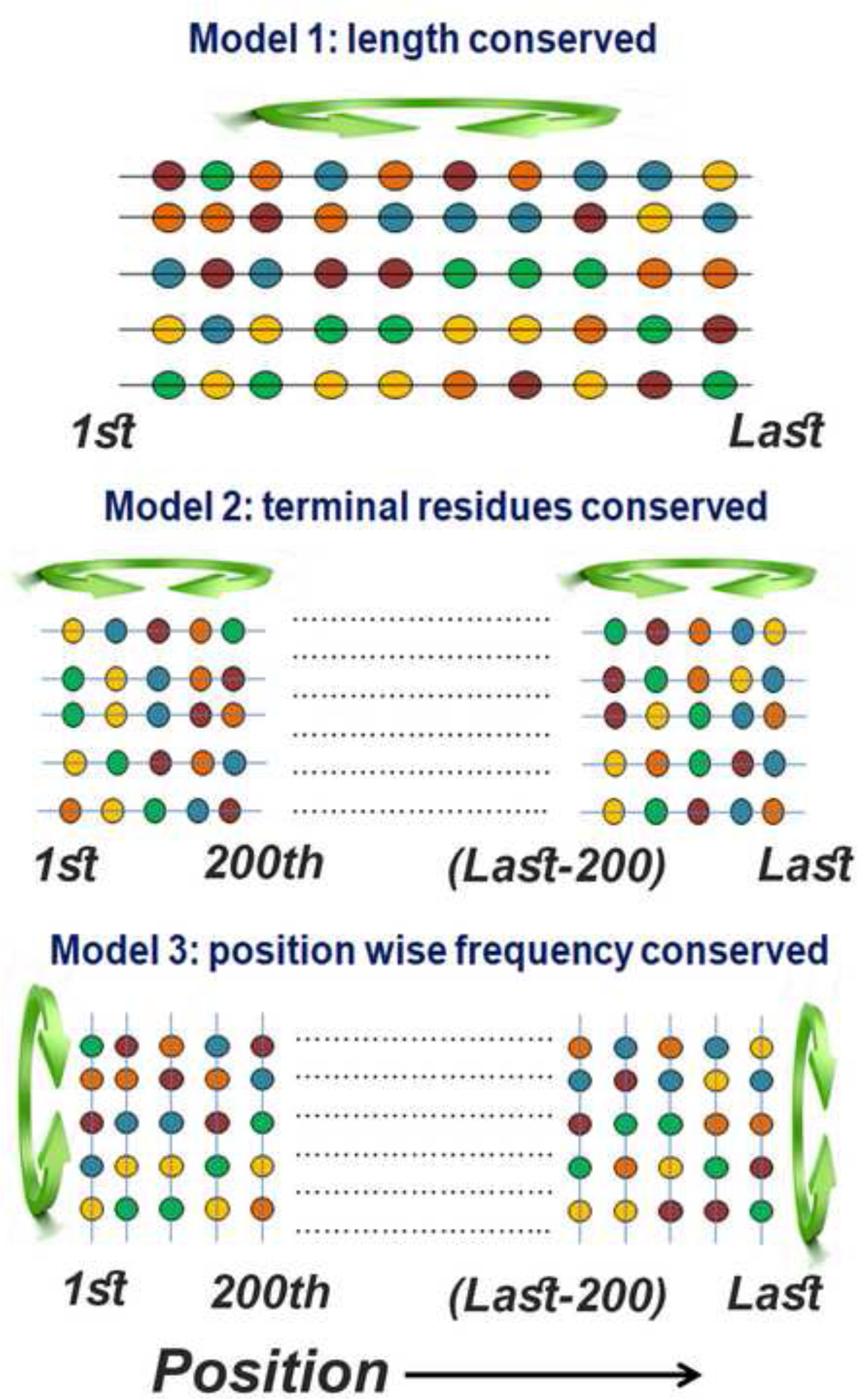
Generation of random protein models. For each naturally occurring protein sequence, we generated three randomized variants. **Model 1**. Random shuffling of amino acids of each real protein. This random model, designated as length conserved random model, maintains overall amino acid composition and length of each real protein. **Model 2**. Shuffling of amino acids (up to the first and last 200 positions) at the N and C-terminals of each real protein. This random model was designated as terminal residues conserved random model which preserves the amino composition of the respective terminal regions. **Model 3**. Shuffling of amino acids in each position of real proteins. For this random model (column-wise random model), naturally occurring proteins of each organism were aligned from both ends and then shuffled position-wise. This model preserves the overall amino acid frequencies at each position of the alignment (see main text for details).

## Results

### Different trends of protein disorder in eukaryotes and prokaryotes

Comparing disorder content of 10,000 real and randomly generated protein models previously it was shown that real proteins are more disordered than the random sequences [23]. However this study considered sequences from different organisms together, therefore till date, we have no clear idea whether all organisms show a similar trend or there is any species-wise variation. In this systematic study, we compared protein disorder content between real and random sequences species-wise. For each organism, we separately generated random artificial protein models preserving the overall amino acid frequencies and length of its real sequences (length conserved random sequences). Here, we considered two measures of protein disorder content such as percentages of all predicted disordered residues and the percentage of disordered residues within long disordered segments and compared between real and random sequences of each test species. These two measures of protein disorder content showed similar distribution when tested with different datasets, except in a few cases.

Proteome-wide average disorder content of real and random sequences of each species is shown in **Figure 2**. In contrast to earlier speculation of high protein disorder among real sequences [23], here we showed that natural sequences could have more or less disorder content compared to the random sequences depending on the characteristics of the species. Specifically, in eukaryotes (*Homo sapiens*, *Drosophila melanogaster*, *Caenorhabditis elegans*, *Saccharomyces cerevisiae*, *Aspergillus oryzae* and *Neurospora crassa*) real sequences were found to be more disordered while in prokaryotes (*Bacillus subtilis, Escherichia coli, Deinococcus radiodurans, Methanosarcina mazei, Haloferax volcanii,* and *Thermococcus gammatolerans*) real sequences were found to be less disordered compared to their corresponding random sequences (Figure 2). These results suggested a general trend that while in eukaryotes natural sequences are more disordered (in terms of percentages of all predicted disordered residues and percentage of disordered residues within long disordered segments) than their corresponding random sequences, prokaryotes follow opposite behavior (Figure 2). We tested all these results with another set of prediction algorithms (see materials and methods, consensus approach 2) and found overall similar results (Figure S1). Thus the earlier speculation of high protein disorder among naturally occurring sequences [23] seemed to be valid in the eukaryotic domain but not in the prokaryotic domain.

**Figure 2.**
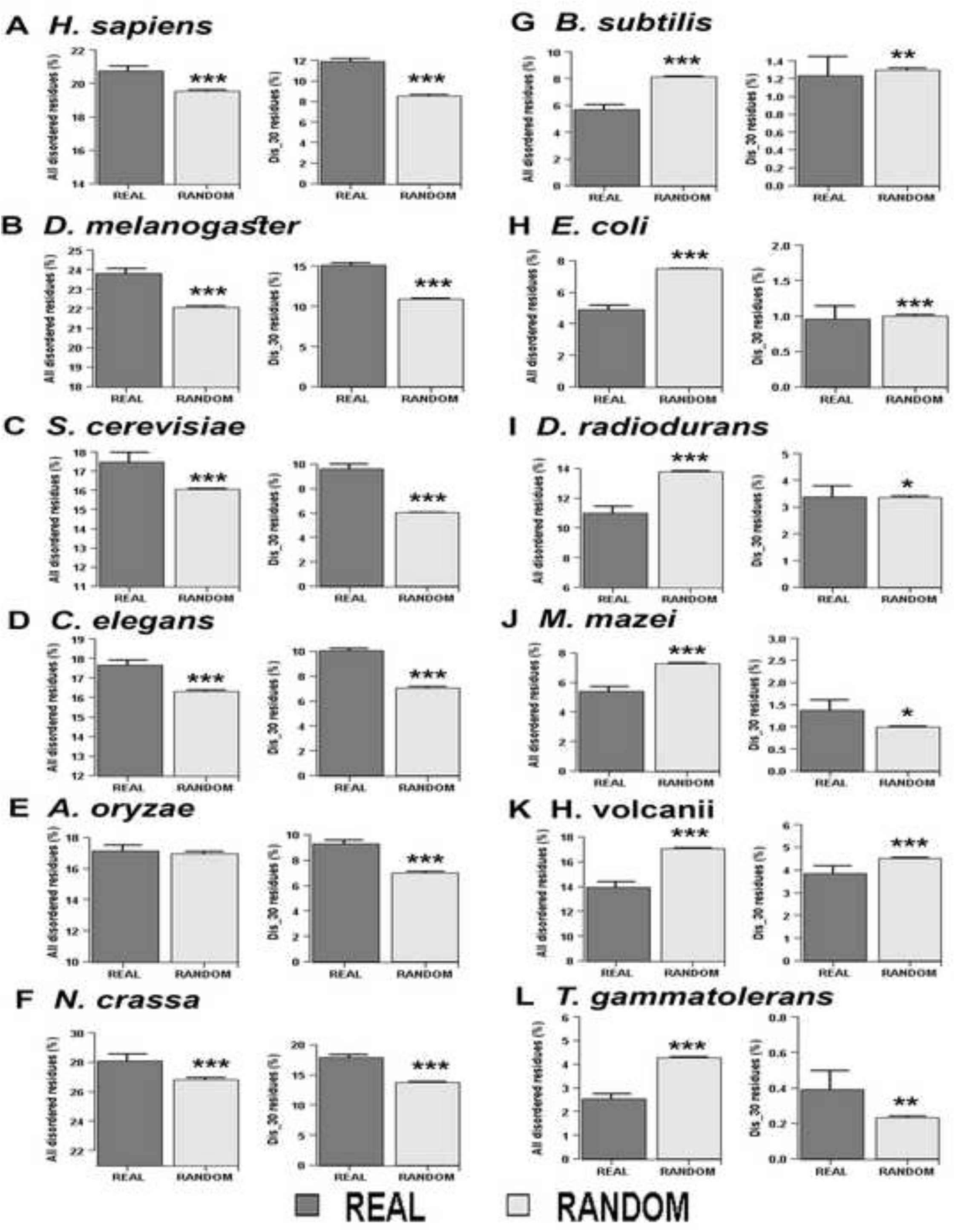
Proteome-wise comparison of disorder content between real and random sequences predicted by consensus approach #1. The graph shows the average disorder content of real and random proteins in 6 eukaryotic: **A**. *H. sapiens*; **B**. *D. melanogaster*; **C**. *S. cerevisiae*; **D**. *C. elegans*; **E**. *A. oryzae*; **F**. *N. crassa*; and 6 prokaryotic organisms: **G**. *B. subtilis*; **H**. *E. coli*; **I**. *D. radiodurans*; **J**. *M. mazei*; **K**. *H. volcanii*; **L**. *T. gammatolerans*. Disordered content is calculated as the percentage of disordered residues in each protein (predicted by the consensus approach #1, see main text for details) then averaged over all the proteins in each group. Disordered content is calculated considering all predicted disordered residues (denoted as percentages of all disordered residues) and considering disordered residues only in long disordered regions (30 or more consecutive disordered residues) (denoted as % of DIS_30 residues). So, there are two plots in each panel, showing the proportion of disordered residues between real and random sequences calculated by these two approaches. *P*-values were calculated by Mann-Whitney U test comparing disordered content between real and random sequences of each species. Significant difference between real and random sequences was shown with *, where * denotes *P* < 0.05, ** denotes *P* < 1×10^-4^ and *** denotes *P* < 1×10^-6^. Here error bars show standard error at 95% confidence interval. Number of proteins in the real datasets are as follows *H. sapiens*: 16,384; *D. melanogaster*: 24,799; *C. elegans*: 21,187; *S. cerevisiae*: 4772; *A.oryzae*: 9830; *N. crassa*: 8899; *B. subtilis*: 2588; *E. coli*: 2838; *D. radiodurans*: 2080; *M. mazei*: 2063; *H. volcanii*: 2410; *T. gammatolerans*: 1346. In each organism, number of proteins in the random datasets is 10 times than the real dataset.

Disordered regions, in general, are encoded by polar and charged residues [1−3], therefore proteomes enriched with these types of amino acids are expected to be more disordered. When we compared the proportion of polar (S, T, N, D, E, Q, R, H, K, Y) and charged (D, E, R, H, K) residues in the proteomes of our test organisms we did not find any general trend in the distribution of charged residues (Figure S2). Instead, our result suggested that prokaryotes may have a higher or lower percentage of charged residues as compared to eukaryotic genomes. However, here we found a distinct pattern in the distribution of polar residues that proportion of these residues are lower in prokaryotic genomes than the eukaryotic genomes considered in this study (Figure S3). Therefore, it may be assumed that the higher proportion of disordered residues in the eukaryotic proteomes is due to their excess polar residues. However, it is worth noting that in each proteome we compared disorder content of real sequences with the random sequences specifically derived from those real sequences preserving the overall amino acid composition and length of the proteins. Thus these results are not expected to be biased by factors such as amino acid composition or length of the protein.

Considering proteins of various genic GC content, Ángyán et al. speculated that structural preferences of real and random proteins depend highly on the GC content of their protein coding sequences [24]. To check whether the disparity of protein disorder content between extant and random sequences, as was observed in this study, depends on the GC content of the coding sequencing here we grouped the naturally occurring proteins of each species according to their genic GC content and compared their disorder content with their corresponding length conserved random sequences. Except for few cases, we noticed an overall similar trend as we found considering all proteins without categorization according to GC content (Figures S4 and S5) which suggested that the observed results are independent of the genomic GC content. Overall this study suggested that previous speculation of high protein disorder among the naturally occurring protein sequences [23] is not a general trend among all organisms; instead, it is evident that prokaryotic and eukaryotic organisms show distinct trends.

### Position-wise enrichment of disorder residues along the protein sequence

To identify the positions those may show preference or abomination for disordered residues, we compared the extent of these residues between natural and random sequences in a position-wise manner (up to the first 150 and last 150 positions). At each position, disorder propensity of naturally occurring proteins was compared with that of two kinds of random protein models (illustrated in Figure 1): one generated by random permutation of amino acids in the terminal regions of each real protein (terminal residue conserved random model) and another generated by permuting the amino acids in each position of the alignment of extant proteins in each species (column-wise random model).

Here it is noteworthy that we found bias in disorder prediction for both the terminal regions (discussed in the next section). Therefore, for each position we computed a Z-score which signifies up to what extent disorder scores of real proteins at any position deviates from that of random proteins at that position (in standard deviation unit). A higher Z-score indicates statistically more significant difference, which is further accessed through *P*-values. Since Z-score compares disorder scores of real and random proteins predicted using same disorder prediction algorithms it is expected to compensate the end bias in disorder prediction in the view that it will have a similar bias for real and random sequences.

Position-wise Z-score profiles of the 12 species considered in this study were shown in **Figure 3** (by consensus approach #1) and Figure S6 (by consensus approach #2), while detailed graphs for each species can be found in Figures S7−S18. These figures show the position-specific disorder profile for the first and last 150 positions of extant proteins of each species and their random variants predicted by consensus approach #1 (panels A, B, C, D) and consensus approach #2 (panels E, F, G, H) respectively.

**Figure 3.**
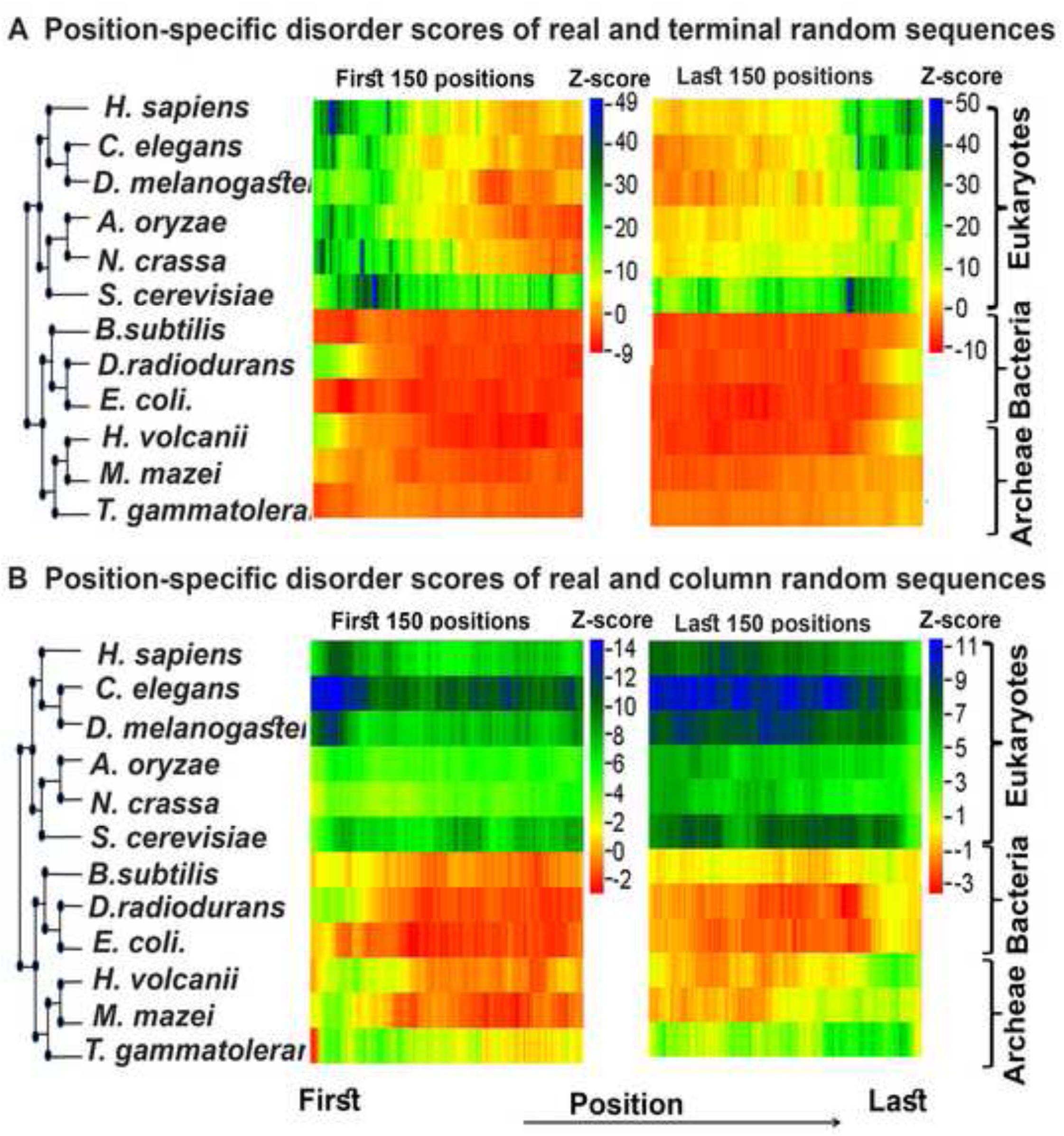
Z—score profile for the position-specific disorder score of each species predicted by consensus approach #1. This figure shows the extent of protein disorder between real and random protein models for the first 150 and last 150 positions of 12 organisms predicted by consensus approach #1 (see main text for details). **A**. Z-scores were calculated comparing position-specific disorder scores of real and terminal residue conserved random sequences of each species and **B**. Z-scores were calculated comparing position-specific disorder scores of real and column random sequences of each species. Z-scores were coded in color scheme (color legend). Here positive Z-score indicates enrichment of protein disorder in naturally occurring sequences while negative Z-score indicates the reverse. Organisms are arranged according to their mid-point rooted species tree retrieved from NCBI taxonomic database with the help of their species taxonomic identifier.

It is evident from Figures 3A, S6A, S6B, and S7−S12 (panels A, B, E, F) that in eukaryotes there is a clear increase of protein disorder (associated with high positive Z-scores) up to the first 50 to 100 residues near the N- and C-terminus of naturally occurring proteins in comparison to the terminal residue conserved random proteins. Further, the trend of higher disorder seems to be stronger near the N-terminal regions rather than the C-terminal regions (Figures 3A). However, in most of the prokaryotes organisms studied here we noticed significantly lower disorder score near the terminal regions of real sequences when compared with similar type of random model (associated with negative or weakly positive Z-score) (Figures 3A, S6A, S6B, S13, S14, S16, and S18, panels A, B, E, F) except in *D. radiodurans* (Figures S15, panels A, B, E, F) and *H. volcanii* (Figures S17, panels A, B, E, F) where we found a weak positive signals up to the first and last 10−15 residues. These results indicated that there is a clear signal for the enrichment of disordered residues near the terminal regions of eukaryotic proteins; however, there is either a week or no such signal in prokaryotic proteins.

When compared with the random model which preserve position-wise amino acid composition (column-wise random sequences) we observed that in eukaryotes natural sequences have a higher proportion of disordered residues than random sequences throughout the considered regions leading to positive but more or less similar Z-scores (Figures 3B, S6C, S6D, S7−S12, panels C, D, G, H). However considering column random sequences as reference in most of the prokaryotic organisms studied here, we did not notice any significant difference in the position-specific disorder scores between real and random sequences at any position (Figures 3B, S6C, S6D and S13−S18, panels C, D, G, H). To ensure that these trends are not the artifacts of disorder prediction algorithms used in this study we tested these results with our new sets of disorder prediction algorithms (consensus approach #2). With this new approach, we found overall similar results (Figures S6–S18, panels E, F, G, H) except at the very extreme ends of proteins (up to the first and last 5−6 residues) where we found relatively weaker Z-score compared to the results obtained by our consensus approach #1. Further, since most of the algorithms used in this study are trained on extant protein datasets their accuracy on randomly generated sequences (such as ours) is questionable. Therefore, here we concentrated specifically on one method IUPred (updated version IUPred2A [25]), which has never been trained on any specific dataset, and calculated Z-score solely based on the prediction of this method. Similar results (Figure S19−S29) as based on consensus approach (Figures S7–S18) imply that our results do not depend on whether the algorithms were trained on real protein datasets or not.

Considering both these approaches (consensus approach 1 and consensus approach 2), we can conclude the following trends (*i*) along the position eukaryotic proteins are more disordered compared to the random sequences, (*ii*) this trend is stronger near the terminal regions (specifically near the N-terminal) of eukaryotic proteins rather than at the center of proteins, and (*iii*) in prokaryotes either there is no such signal or weekly positive signal of selection for disordered residues near the terminal regions. In the next few sections, we explored the possible cause(s) and consequence(s) of these observed trends.

### Bias in disorder prediction at the terminal regions cannot explain the trends

We noticed that almost all the prediction algorithms predict very high disorder near the protein ends. Therefore, one probability may be that the ends of the proteins (irrespective of whether terminal regions of or not), in general, show the trends due to biases in disorder prediction. To check this possibility we removed the terminal regions (up to the first and last 50 residues where we found highly positive Z-score) from the extant protein sequences of six eukaryotic organisms showing the trend. We generated random protein models (both terminal residue conserved random sequences and column random sequences) from these truncated sequences, predicted disordered residues freshly in these (real and random) sequences and compared their disorder score following the Z-score approach as described before. Considering two types of random models (column random and terminal residue random models), overall we found similar results corresponding to the analogous (50−150^th^) position in the corresponding full-length sequences (Figures S30–S35). For instance, in our main analysis when we compared the disorder score of full-length real sequences in reference to terminal residue conserved random model we found weekly positive Z-scores (as compared to the end positions) after 50th position from both the terminal regions. Considering terminal residue conserved random model, in this analysis (with truncated sequences), we did not find any significant trend (high positive Z-score) near the C-terminal region of most of our test eukaryotic organism (Figures S30–S35, panel B). No positive trend (Z-score) is also noticed near the N-terminal region of *H. sapiens* (Figure S33A) and *C. elegans* (Figure S31A) proteins. However, a significant trend was noticed near the N-terminal region of other eukaryotes such as *S. cerevisiae* (Figure S35A), *N. crassa* (Figure S34A), and *D. melanogaster* (Figure S32A). This is generally weaker than the N-terminal regions of their full-length sequences and similar to the trend that was found near the 50^th^ position of their full-length protein sequences. Considering column random sequences, in our full-length protein datasets, we found more or less similar (and positive) Z-score along the length of the protein. Here we also found similar, however generally weaker trend near the terminal regions (of truncated sequences) when we considered column random sequences as reference (Figures S30–S35, panels C, D). Overall, these results suggest that results obtained are overall similar to the results obtained at the analogous position of full-length protein sequences. Thus, the trends that we found near the terminal regions of eukaryotic proteins cannot be iterated in any other position generating artificial protein ends.

### Selection for high GC content at the nucleotide level cannot explain the trends

Results presented so far revealed a general (proteome-wise disorder content) and regional (near the terminal regions) enrichment of disordered residues among eukaryotic proteins as compared to their corresponding random models. Protein disorder content was suggested to depend on a number of factors among which genomic GC content [16,17,26] was considered as most significant. Further, significant correlations (**Table 1**) were found between protein disorder content and genomic GC content in each species suggesting that the observed trends may be due to the selection for high or low GC content at the nucleotide level. In the previous section, we showed that the trend that we found considering disorder content of full-length protein is valid over the entire GC ranges of their coding sequences (Figures S4 and S5). Now, here we checked the second possibility whether the enrichment of protein disorder near the N- and C-terminal regions of eukaryotic proteins are the results of selection for high GC content at the respective sites. To explore this possibility, we compared GC content of real protein-coding sequences of each test species with that of randomly generated nucleotide sequences. For this, we generated random nucleotide models analogous to terminal residue conserved and column random protein models and calculated Z-score which signifies the deviation of GC content between real and random sequences.

**Table 1.** Correlation between GC content and protein disorder content.

| Lineage | Species name | Sample size (N) | Correlation between GC content and % of Dis_all residues | Correlation between GC content and % of Dis_30 residues |
| --- | --- | --- | --- | --- |
| Mammal | *Homo sapiens* | 16,384 | $\rho = 0.235; P < 1 \times 10^{-6}$ | $\rho = 0.189; P < 1 \times 10^{-6}$ |
| Insect | *Drosophila melanogaster* | 24,799 | $\rho = 0.300; P < 1 \times 10^{-6}$ | $\rho = 0.279; P < 1 \times 10^{-6}$ |
| Worm | *Caenorhabditis elegans* | 21,187 | $\rho = 0.387; P < 1 \times 10^{-6}$ | $\rho = 0.312; P < 1 \times 10^{-6}$ |
| Fungi | *Saccharomyces cerevisiae* | 4772 | $\rho = 0.114; P < 1 \times 10^{-6}$ | $\rho = 0.049; P = 2.05 \times 10^{-4}$ |
| Fungi | *Aspergillus oryzae* | 9830 | $\rho = 0.228; P < 1 \times 10^{-6}$ | $\rho = 0.194; P < 1 \times 10^{-6}$ |
| Fungi | *Neurospora crassa* | 8899 | $\rho = 0.082; P < 1 \times 10^{-6}$ | $\rho = 0.068; P < 1 \times 10^{-6}$ |
| Bacteria | *Bacillus subtilis* | 2588 | $\rho = 0.075; P = 2 \times 10^{-6}$ | $\rho = -0.028; P = 7.2 \times 10^{-2}$ |
| Bacteria | *Escherichia coli* | 2838 | $\rho = 0.101; P < 1 \times 10^{-6}$ | $\rho = 0.061; P = 1.11 \times 10^{-4}$ |
| Bacteria | *Deinococcus radiodurans* | 2080 | $\rho = 0.150; P < 1 \times 10^{-6}$ | $\rho = 0.066; P = 2.13 \times 10^{-4}$ |
| Archaea | *Methanosarcina mazei* | 2063 | $\rho = 0.135; P < 1 \times 10^{-6}$ | $\rho = 0.071; P = 4.50 \times 10^{-5}$ |
| Archaea | *Haloferax volcanii* | 2410 | $\rho = 0.072; P < 8 \times 10^{-6}$ | $\rho = 0.086; P < 1 \times 10^{-6}$ |

*Note:* This table shows the correlation between disorder content of naturally occurring proteins in each species with the GC content of their coding sequences. Disorder content of a protein was calculated considering (i) all the predicted disordered residues (denoted as percentages of all disordered residues) and (ii) considering disordered residues only in long disordered regions
(denoted as % of DIS\_30 residues). In each protein, disordered residues were predicted by consensus-based approach #1 (see main text). Following non-parametric distribution of protein disorder content here we calculated Spearman's Rank correlation coefficient $\rho$ , where $P$ -values stand for significance level. Here $N$ stands for sample size *i.e.* the number of proteins and their coding sequences considered in each species for this test.

The GC profile near the protein terminus with respect to terminal residue conserved and column random nucleotide models were shown in Figures S36−S48 respectively. These figures suggest that in each of the test species position-wise GC content of real sequences varies in a narrow range along the reading frame while random sequences show almost no variation. When we computed the difference in terms of Z-score we found either no or week evidence of preference (*P* > 0.05) for high or low GC near the protein terminal regions in most of the species. If GC content has any impact we could expect a concomitant trend with what we observed for disordered residues in all the tested species. Further, to probe the possibility of any hidden link between selection at nucleotide and protein level we correlated Z-scores for disordered residues with corresponding Z-scores for GC content. Here we did not find any significant correlation between these two measures in any of the tested species. Overall, these results suggested that the observed trends of high disorder near the terminal regions of eukaryotic proteins are independent of the selection for higher GC content at the nucleotide level.

### Splice junctions cannot explain the trends

Previously it was shown that disordered residues are more prevalent near the splice junctions of their coding sequences [27]. Therefore, there is a high possibility that trend that we observed near the terminal regions of eukaryotic proteins may have caused by proteins having excessive splice junction in their coding sequences near those regions. To check this possibility, here we considered only proteins without any splice junction in their coding sequences up to the first and last 100 positions (*i.e.* either encoded by a single CDS or the first and last CDS more than 300 base pair in length) of 6 eukaryotic organisms where we mainly found a higher fraction of protein disordered residues in real sequences as compared to random models. When compared position-specific disorder scores of these proteins with their corresponding terminal residue conserved random sequences and column random sequences (specifically generated from these sequences) in most of the eukaryotes we found similar trends as we found considering all proteins (Figure S49−54) which suggest that splice junctions merely have any effect on our observed trends.

### High solvent accessibility near the terminal regions of proteins cannot explain the trends

Being located on the surface of the proteins, terminal regions were considered to be solvent exposed [28]. This solvent exposed nature of terminal regions was suggested to arise from excessive use of hydrophilic and polar residues [28] which were further known to increase the propensity of a protein to be disordered [1–3]. Therefore high protein disorder near the terminal regions of eukaryotic proteins may be a side effect of charged residues selected mainly for solvent-exposed nature of these regions. To test this possibility here we calculated Z-score profile for predicted solvent-accessibility following similar approach as we did for predicted disordered residues. Considering both the random models (our terminal residue random model and column random model) we did not find any strong evidence in any of our test organism that real proteins show preference for higher solvent accessibility near their terminal regions as compared to random expectations (Figures S55−S67). Moreover, we did not find any significant correlation between the Z-score for predicted disordered regions and the Z-scores for predicted solvent accessibility which further suggested that trend that we observed near the terminal regions of eukaryotic proteins is independent of their high solvent exposed nature.

### Most of human GO slim functional categories show the trends

Disordered proteins were shown to execute a high level of functional specificity as compared to ordered globular proteins [3−6]. Therefore one pertinent question may be whether the signal for high protein disorder near the terminal regions of eukaryotic proteins is function specific? To dig deeper into this aspect we grouped human proteins according to their GO-slim functional ontologies. Considering three broad GO ontologies biological process (BP), molecular function (MF), and cellular component (CC) here we found 124 GO slim functional keywords with at least 100 proteins. Next, we compared position-specific disorder scores of proteins under each such GO-slim ontology with their corresponding terminal residue conserved random sequences (model 2) and column random sequences specifically generated for each GO category (model 3) (see material and methods). Proteins associated with most of these ontologies showed a preference for high disorder near the N and C-terminal regions. For instance, ∼78% (51 out of 65) of GO BP, ∼71% (25 out of 35) of MF, and ∼69% (16 out of 23) of CC ontologies showed strong to moderate signal for the preference of high protein disorder near the N-terminal regions when compared with terminal residue conserved random sequences. The exceptional cases where we found either negative or relatively weaker signals were associated with different types of metabolic and developmental processes for BP, different types of enzymatic functions such as helicase, isomerase, oxidoreductase, ligase *etc.* for MF and protein extra-cellular matrix, ribosome, Golgi apparatus *etc*. ontologies for CC (**Figures 4** and S68−S78). Probable explanations for this pattern are discussed in discussion. When we compared with column random sequences we did not find any specific trend near the terminal regions, however, we noticed positive signal through-out the considered regions in most of the functional classes. These results suggest that the trends reported in the previous section are in general not specific to proteins belonging to any particular functional category; however, the strength of the signal (positive z-score) may not be identical in all such groups.

**Figure 4.**
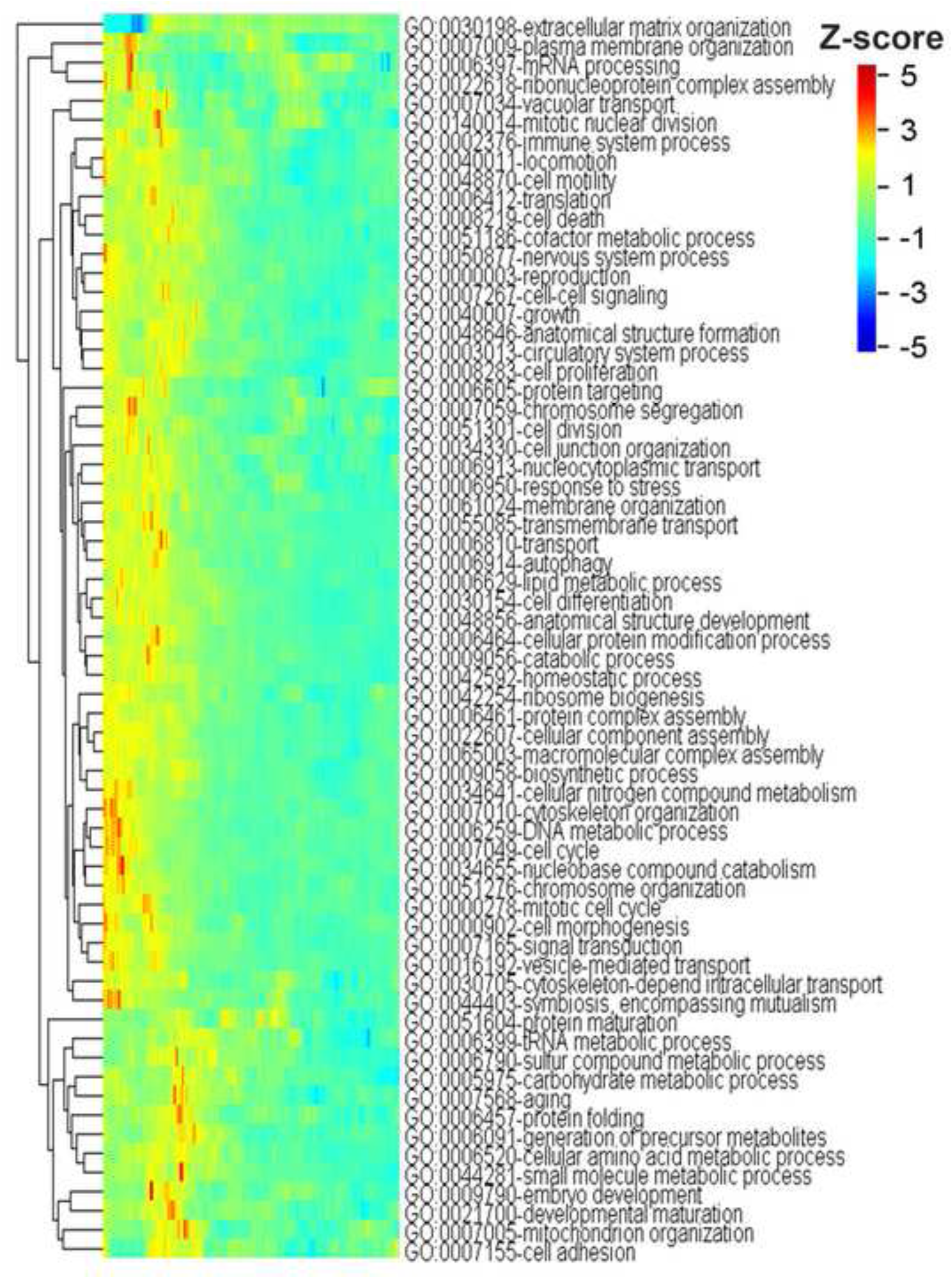
Mean Z—score profile for the first 150 positions of human proteins under GO slim biological process ontologies in reference to terminal residue conserved random model. This heat map represents position-wise Z-score of predicted disordered residues (predicted by consensus approach #1) for the first 150 positions at the N-terminal region of human proteins under each GO slim biological process ontology. In each functional category, Z-scores for predicted disordered residues were calculated considering proteins (more than 200 residues in length) under that category in reference to their corresponding terminal residue conserved random sequences. Here rows represent the positions along the protein sequence. Z-scores were coded in color scheme (color legend). Only the ontologies with more than 100 proteins were shown here.

### Highly and lowly expressed genes show species-specific trends

IDPs were shown to express at a lower level than well structured globular proteins [29, 30]. Therefore, we found it interesting to analyze whether the trends that we observed near the terminal regions of eukaryotic and prokaryotic vary according to their gene expression level. Thus here we compared Z-scores (that measures the extent of preference for disordered residues in real sequences as compared to their corresponding random models) between the proteins encoded by highly and lowly expressed genes of three species *H. sapiens*, *S. cerevisiae*, and *E. coli* with high throughput gene expression data. In each of those species, Z-score was computed for highly and lowly expressed proteins separately in reference to their terminal residue conserved and column random sequences (see material and methods). When we compared Z-scores obtained in reference to terminal residue conserved random model, lowly expressed proteins of *H. sapiens* and *S. cerevisiae* showed higher Z-score compared to highly expressed proteins near both the terminal regions (Figure S79). However, in *E. coli* we did not find a prominent difference in Z-score between these two groups of proteins. When we compared Z-scores obtained in reference to column random sequences we noticed similar trends (Figure S80).

### Regions showing enrichment of disordered residues in eukaryotic proteins are enriched with disordered binding sites

To test whether the higher fraction of disordered residues near the terminal regions of eukaryotic proteins compared to random protein models has any role in protein-protein interactions here we predicted potential interaction sites within those regions using ANCHOR [31]. ANCHOR predicts probable interaction sites within disordered regions thus suggested to provide an unbiased estimate of interaction potential that conferred from disordered residues [31]. Our results presented in **Figure 5** showed that eukaryotic (*H. sapiens*, *D. melanogaster*, *C. elegans*, *S. cerevisiae*, *A. oryzae* and *N. crassa*) proteins, in general, have more protein-protein interaction sites near the terminal regions compared to the corresponding terminal residue conserved random sequences while no such specific trend is observed in prokaryotic (*B. subtilis, E. coli, D. radiodurans, M. mazei, H. volcanii,* and *T. gammatolerans*) organisms. When we compared with column random sequences we found consistently higher Z-score for predicted ANCHOR residues throughout the considered regions in eukaryotes. Overall, these results are consistent with the position-specific disorder profile of eukaryotic and prokaryotic proteins near the terminal regions. These results may be taken as an indication that disordered residues are preferred specifically near the terminal regions of eukaryotic proteins to promote protein-protein interaction sites.

**Figure 5.**
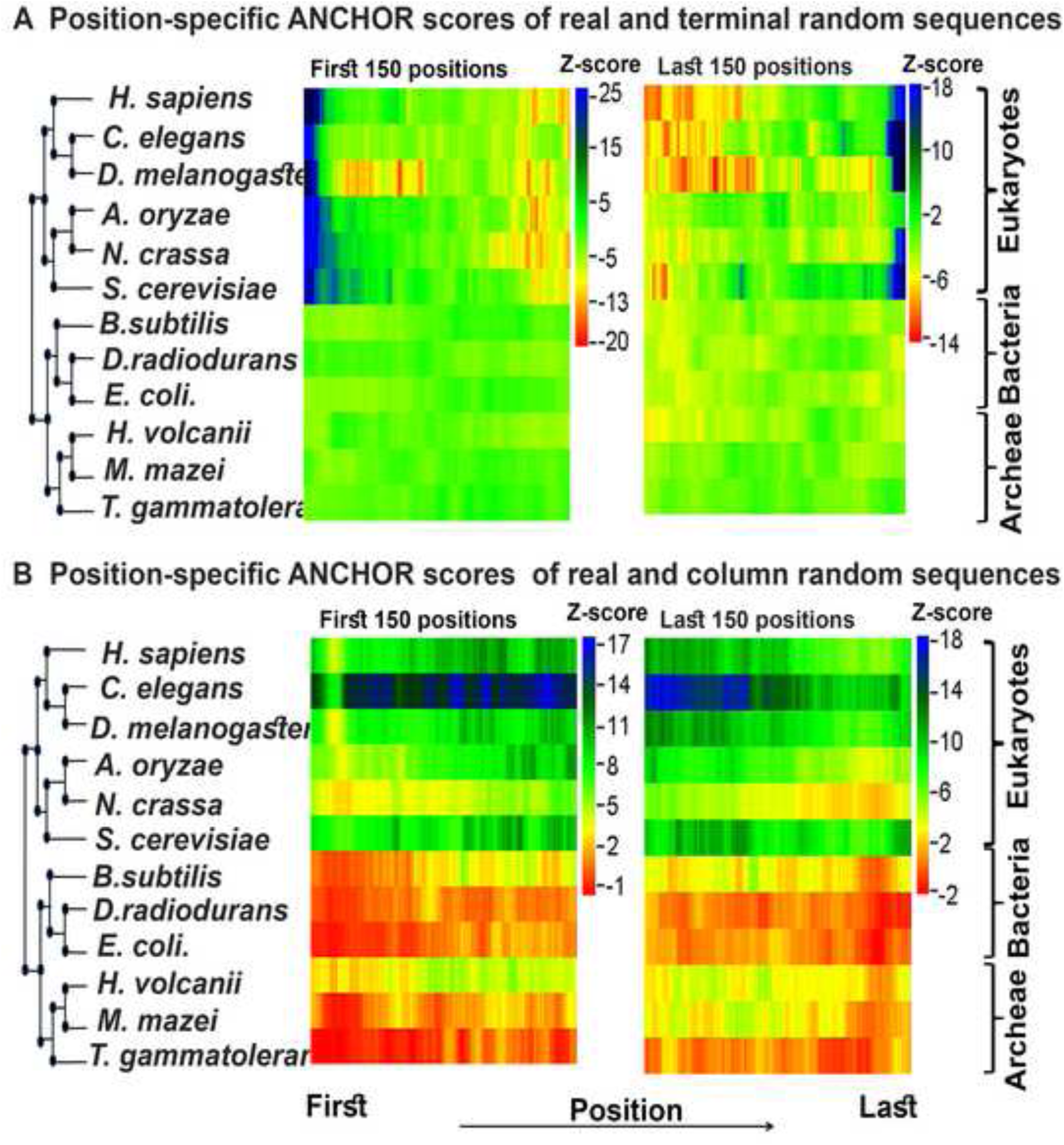
Z—score profile for the position-specific comparison of ANCHOR predicted protein binding residues. This figure shows the extent of protein binding sites within disordered regions between real and random protein models for the first 150 and last 150 positions of 12 organisms considered in this study. **A**. Z-scores were calculated comparing predicted ANCHOR residues of real and terminal residue conserved random sequences of each species and **B**. Z-scores were calculated comparing that of real and column random sequences of each species. Z-scores were coded in color scheme (color legend). Here positive Z-score indicates enrichment of ANCHOR predicted protein binding sites in naturally occurring sequences while negative Z-score indicates the reverse. Organisms are arranged according to their mid-point rooted species tree retrieved from NCBI taxonomic database with the help of their species taxonomic identifier.

## Discussion

One of the major goals of protein structural biology is to understand the structural and functional characteristics of intrinsically disordered proteins. Extensive research over the past few decades, reviewed in references [1−6], has facilitated our understanding of these proteins. However, to date, there are many fundamental issues which are not clearly understood. How nature has shaped the disorder potential of naturally occurring eukaryotic and prokaryotic proteins is one of those unresolved questions which still remain elusive. Considering naturally occurring proteins from several model organisms and artificially generated random protein models here we seek to understand whether there is any preference for disorder residues among the extant proteins as compared to random expectations.

For our study, we generated three kinds of random models. First, to compare proteome-wise disorder content between real and random sequences of each species we considered a general random model that preserves the overall amino acid composition and length of the proteins but not the order of the amino acids (length conserved random sequences). However, at any position disorder potential of natural proteins may be influenced by any other trait(s) (discussed later) that depends on amino acid composition at that position. Therefore, here we considered two other random models one by random shuffling only the terminal residues and another that maintains overall frequencies of amino acids at any particular position of real proteins and hence expected to preserve the features induced by the biased distribution of amino acids.

At first, we compared proteome-wide protein disorder content between real and random sequences in each of the 12 selected genomes. Comparing disorder content of few selected proteins mainly short peptides collected from UniRef databases with random sequences of same overall amino acid frequencies, Yu et al. suggested that high protein disorder among natural sequences is a general evolutionary trend [23]. By contrast, in this systematic analysis, we compared the disorder content of real sequences of each species with their corresponding length conserved random sequences. Thus our study allowed us to investigate the trend species-wise. Our results suggested that, depending on the characteristics of the species, natural sequences may have more or less disorder content compared to the random sequences and the trend is highly species-specific. A general pattern emerged from these results that in eukaryotes, at least for the species considered in this study, naturally occurring proteins are more disordered compared to random sequences but not in prokaryotes which may suggest that disordered residues are more preferred in eukaryotic proteins rather than in prokaryotic proteins. However, here we remark a note of caution that a number of caveats may explain the observed trends which have been discussed shortly. Previously, numerous studies have indicated that eukaryotic proteins are in general more disordered than prokaryotic ones [13−15]. However, to the best of our knowledge, no study has ever analyzed whether there is any disparity in selection for disordered residues between eukaryotic and prokaryotic proteins as we did in this study. Here, our results suggested for the existence of differential selection for disordered residues which may explain the variation in disorder content between eukaryotic and prokaryotic proteins. Disordered residues were supposed to play crucial roles for the rise of complex eukaryotic organisms [4,5,13,16]. Most of the novel functions such as transcription factors, transmembrane receptors, signaling proteins, intra-cellular communication, cytoskeletal proteins, and chromatin organization, *etc.* which appear early in eukaryotes were noted for their elevated level of protein disorder. Considering their importance in higher organisms it was suggested that proteomic disorder content of a species is linked with its genomic complexity [4, 13]. Thus the general preference for disordered residues (as compared to random expectation) among eukaryotic proteins as we observed in this study may be an evolutionary relic of the role IDPs played in these organisms.

Next, we looked for regions which may have selected for high or low disorder within the proteins. It is known from earlier studies that N-terminus regions of DNA binding homeodomain proteins are generally disordered by nature due to their high net charge [32]. Disordered tails at the N-terminal regions was suggested to be advantageous for the DNA binding proteins to serve as an anchor for high specificity and low-affinity binding (fly-casting mechanism) with cognate DNA molecules [32]. However, there is no general understanding of whether disordered residues are uniformly distributed along the protein or there is any site-specific variation. As with proteome-wise disorder content here we noticed a prominent difference in the observed trends between eukaryotic and prokaryotic proteins. Our study suggested that during the course of evolution, eukaryotic proteins have specifically accumulated a higher fraction of disordered residues near the terminal regions (particularly near N-terminus) more than what could be expected from the random distribution. However, we did not find such a clear trend in the prokaryotic organisms except in *D. radiodurans* and *H. volcanii* where we found a weak signal for high disorder in the first and last few positions. On the basis of our results here we propose that it is not only the DNA binding homeodomain proteins but high disorder near the terminal regions specifically near the N-terminus is a more general trend among eukaryotic proteins.

Below are some probable explanations for the above-mentioned trends.

First, it is possible that the trends that we observed in this study may have resulted from the constraints imposed by any other factors. Specifically, since the codons for most of the disorder-promoting amino acids are GC rich, a strong association has been proposed between genomic GC composition and protein disorder content [16,17,26]. Earlier, Ángyán et al. suggested that disorder potential of probable de-novo proteins is a function of GC content of their coding sequences [24]. In order to disentangle the impact of GC on the observed disparity in proteome-wide disorder content between the real and random sequences of prokaryotic and eukaryotic species, in each species, we compared disorder content of real sequences pooled from different genomic GC background with that of their corresponding length conserved random sequences (*i.e*. in GC bins). Except for few cases, in each species, we noticed an overall similar trend as we found considering all proteins (without any GC bin) which suggested that genomic GC has merely any impact on the observed trends. Next, we analyzed whether the trends that we observed near the terminal regions of eukaryotic proteins have resulted from selection for high GC at the nucleotide level. Constructing similar random models as for proteins we did not find a common trend between predicted disorder and GC content in most of the tested species suggesting that our observed trend is independent of selection at the nucleotide level. Actually, in many of our test organisms the GC content at the 5’end of the mRNA is relatively lower (probably related to weak mRNA folding in these regions [33, 34]). Thus, this feature cannot describe our reported intra-protein pattern.

Second, it is possible that the amino acid bias at the end of proteins may explain this signal. Amino acids are not uniformly distributed along the sequences, specifically, terminal regions were shown to prefer charged residues for their solvent exposed and flexible nature [28]. On the other hand, polar and charged residues are also known to increase the propensity of a protein to be disordered [1–3]. Thus, the trend that we observed near the terminal regions of eukaryotic proteins may have arisen because of the higher prominence of charged residues selected mainly for the solvent-exposed nature of these regions. However, here we did not find any parallel trend of preference for solvent accessibility (as compared to random expectation) as we found for disordered residues, which suggest that our observed trend is independent of the solvent-exposed nature of these regions. Moreover, the random models that we used to compare the position-specific disorder score between real and random sequences were generated in view to preserve the characteristics of terminal regions. For our second random model (terminal residue conserved) we shuffled the amino acids only in the terminal regions and for our third random model (column random) we shuffled the amino acids in each position of real proteins. Thus the random sequences generated by these way supposed to maintain the regional characteristics of the terminal regions. High protein disorder with respect to random sequences with similar amino acid background suggests that our observed results cannot be explained simply by the charged surface exposed nature of the terminal regions.

Third, it is possible that this enrichment is partially related to the lack of selection at the ends of the protein. Studies that have linked evolutionary rate with protein structure have consistently found that exposed sites are more tolerant of amino acids substitutions and evolve at a higher rate than buried sites [35]. This, in turn, suggests that terminal regions being solvent exposed by nature evolve under weaker evolutionary constraint than the central regions. Thus terminal regions were considered as “evolutionary playgrounds” for the innovation of new functions [36]. This reduced efficacy of selection at the protein terminus, especially in eukaryotes because of their lower effective population size compared to prokaryotes [37] may have provided the permissive environment for the fixation of disordered promoting amino acids which are generally associated with high rates of insertions, deletions and substitutions [7,8,10]. It is possible that disordered residues in these regions tend to be less deleterious in eukaryotes than in prokaryotes facilitating their fixation. Nevertheless, the fact that the amino acid distribution at the end cannot explain these patterns, as we showed based on our null models, supports the conjecture that lack of selection for certain amino acids is not the only explanation; if there is no selection we expect to see a similar pattern in the null models. Moreover, the results are not always in the direction what could be expected from the population genomic consideration. According to population genomic model [37], disordered residues, if deleterious, are expected to be purged from the genomes of higher effective population size due to their higher efficacy of selection. However, as an example, here we noticed comparable trends near the terminal regions of both *S. cerevisiae* and *H. sapiens* in spite of wide variation in their effective population size. This suggests that reduced efficacy selection as expected from the perspective of effective population size is not the major cause for the trends that we observed in eukaryotic genomes.

Fourth, disordered residues may have selected specifically near the terminal regions of eukaryotic proteins because of functional reasons especially for the higher proximity of protein-protein interactions (explained in **Figure 6**). Due to their inherent structural flexibility, disordered residues could form binding sites for a large number of partner proteins [1–6]. It is probable that this specific enrichment of disordered residues near protein terminus generates two ’arms’ which mediate non-specific weak protein-protein interaction that improve the search for the specific interacting protein partner (see Figure 6). To check whether high disorder at the N-terminal regions of eukaryotic proteins has any role in protein-protein interactions here we tested the distribution disordered binding motifs in real and random sequences of each test species. Disordered binding motifs are short stretches of disordered residues that undergo order to disorder transition upon binding and were considered to be crucial in molecular recognition for their binding capacity [5,11,12]. Moreover, disordered binding sites were suggested to act as a flexible linker for protein-protein interactions [6, 31]. Thus, the higher proportion of disordered binding residues near the terminal regions of eukaryotic proteins may help these proteins to attain structural flexibility for binding promiscuity. In support of this view, disordered N-terminal tails of homeodomain proteins were shown to facilitate DNA search and accelerate specific binding with partner DNA molecules which may be associated with our results [32]. This is possibly more important in the larger, more complex eukaryotic cells rather than in prokaryotic cells.

**Figure 6.**
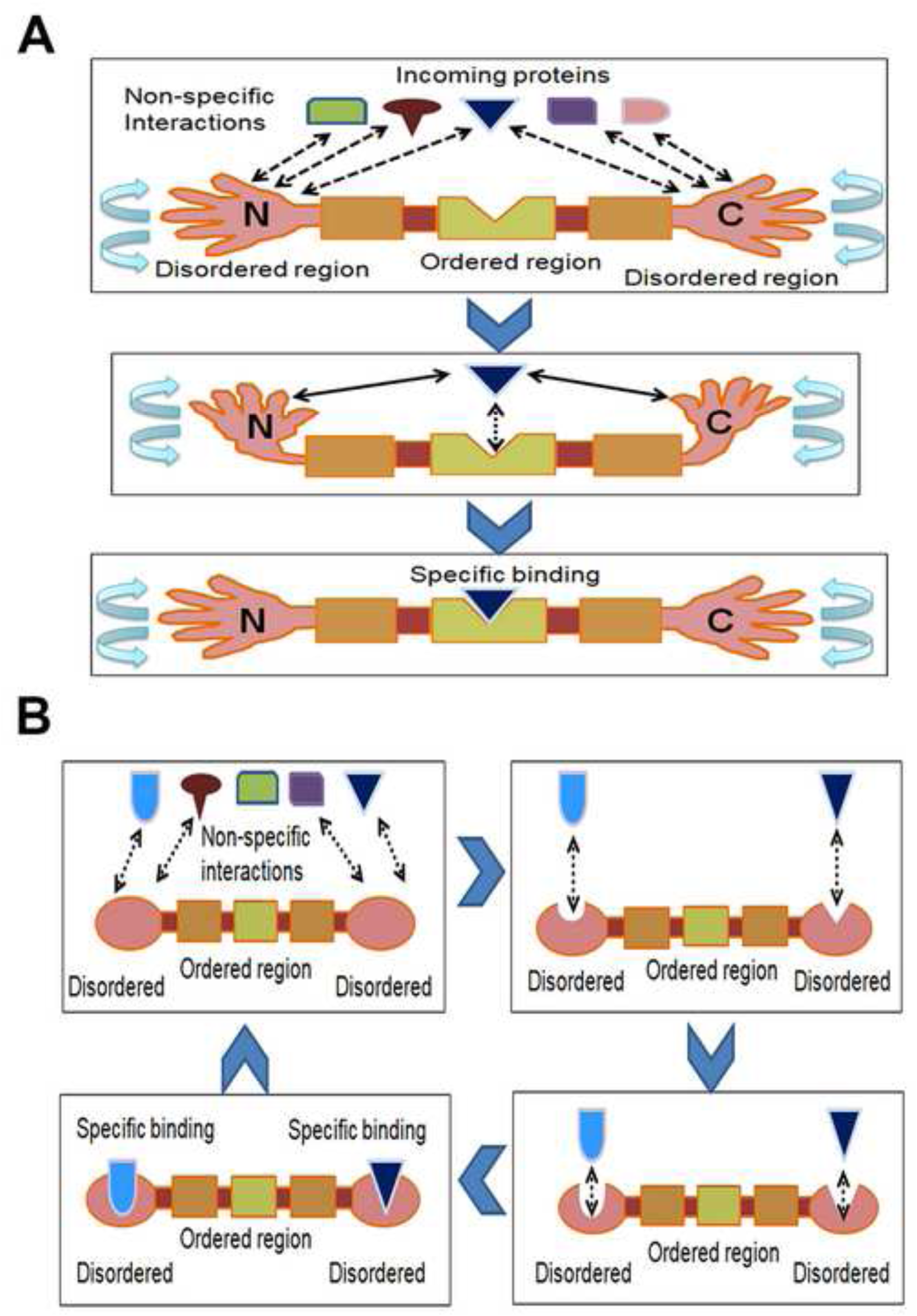
Potential roles of disordered terminus in protein-protein interaction. A. Disordered tails of eukaryotic proteins may act like two arms to improve the search for the specific interacting protein partners. Disordered regions bind with the partners with weak and non-specific interaction which may help to mediate more specific interactions. B. Being flexible in nature, disordered tails may form binding sites for large number of protein partners.

Fifth, terminal regions of the eukaryotic protein may have evolved a higher fraction of disordered residues because of other functional advantages. It is possible that on average the proper functionality of residues or domains near the end of the proteins requires a higher level of disordered residues. Further, the conformational plasticity of disordered residues may be exceptionally advantageous for the terminal regions than the core regions. This is evident from the fact that depending upon the length, 30 to 97% of human proteins were predicted to have disordered stretches near their N and/or C-terminus [38]. Multifarious roles of disordered residues in protein terminal regions were reviewed elsewhere [39]. In brief, disordered residues at the terminal regions were considered to be advantageous for G-protein-coupled receptors, voltage-gated potassium channels and ligand binding in the transmembrane region, *etc*. among several others [39]. Disordered regions are often targeted by several types of post-translational modifications (PTMs) and alternative splicing (AS) [1], [4–6]. AS and PTMs are important means of generating structural and functional diversity of eukaryotic proteins without much affecting genome size. A higher fraction of disordered residues near the protein termini may have evolved for the functional exaptation in form of sites for AS and/or PTMs which shown to be more prominent in the terminal regions and especially in eukaryotic proteins [40].

Sixth, one probable explanation may be that the signal related to protein disorder in the eukaryotes is different than prokaryotes due to the differences in protein folding pathways including the way nascent chain acquires structure in eukaryotic and prokaryotic proteins and differences in the ribosomes of the two domains. Particularly, a possible explanation for not finding any significant trend of high disorder in prokaryotic organisms may lie in their growth kinetics. Maximization of growth rate is a fundamental aspect of prokaryotic biology [41]. Most of the bacteria grow extremely fast (generation time usually ranges from few minutes to several hours) [41], while a typical eukaryotic (human) cell takes about 24 hours to divide [42]. To achieve their higher growth rates, prokaryotes are under strong selection to adopt several strategies such as increasing their replication and translational speed and efficiency, *etc.* [41]. Recent experiments suggested that the speed with which nascent peptides emerge from the ribosome is an important parameter that determines subsequent folding and can alter the overall conformation of the protein. When peptides are synthesized, nascent peptides pass through negatively charged ribosomal exit tunnel. Therefore, positively charged residues were suggested to retard the protein translational rate [43]. Later, considering more diverse datasets it was shown that it is not only the positively charged residues, however, charged residues (positive or negative), in general, may show stalling effect [44] which may have positive effect on fitness when it is at the 5’end of the mRNA [45, 46]. Disordered regions, in general, are enriched with polar and charged residues [1–3]. Hence, high amount of disordered residues especially near the rate-limiting initiation site (N-terminal regions) may be detrimental for the higher growth rates of prokaryotic organisms; specifically, it is possible that the relation is direct -- the fact that disordered residues may increase interaction with the ribosomal exit tunnel. Therefore, it is conceivable that prokaryotes will tend to use a lower amount of disordered residues compared to random expectation and especially near the protein termini.

Here we discussed some probable explanations for the enrichment of disordered residues near the protein ends in eukaryotes, however, further studies and experiments will be needed to understand their relevance fully.

A position-dependent relationship was proposed between protein disorder and function suggesting that the relative position of the unstructured region within a protein provides clues regarding its functionality [47]. Specifically, proteins forming different kinds of ion channels and those involved in transcription factor activation or repression were supposed to encode higher fraction of disordered residues near their N and C terminals respectively, as compared to the proteins related to transcription regulator, RNA pol II transcription factor, and DNA binding, *etc.* processes which contain disordered regions near the interior [47]. However, it is not clear from that study whether this trend is due to selection for high or low disorder among those functional classes. Here we asked whether the signal for high disorder near the N-and C-terminus of eukaryotic proteins is specific to any functional class. Considering 124 general functional annotations from GO slim database for human proteins we showed that except for few specific functional categories mainly related to different types of enzymatic functions and metabolic processes high disorder at the terminal regions is a common feature of human proteins belonging to the most of the other functional classes. Overall, these results are in line with previous observations suggesting that disordered residues specifically enriched among the functional classes such as signaling, transcription, cell division, apoptosis, post-translational modification, and various regulatory processes while depleted among proteins involved in enzymatic and catalytic functions [3−6]. Disordered regions were ascribed to be advantageous for the above-mentioned processes because of their functional prerequisites such as high specificity and low-affinity binding, ease of regulatory control, *etc*. which cannot be achieved from ordered proteins [3−6]. However, those studies mainly considered disorder content of entire protein without looking for any site-specific signature. Here we showed each of those functional classes bears a specific signature for high or low disorder near protein terminus.

High expression of disordered proteins was supposed to be detrimental for cell survival because their harmful effects when over expressed [11,29,30]. Since a high concentration of disordered proteins may lead to several disease conditions, cells were proposed to develop several mechanisms to control their expression level below a certain level [11,29,30]. This notion is supported by the observation that in higher organisms such as *H. sapiens* and *S. cerevisiae* disordered proteins are generally expressed at a lower level than the globular proteins [29, 30]. This may suggest that in eukaryotes highly expressed genes would show weaker selection for disordered residues than lowly expressed genes. Indeed, when we compared Z-score for predicted disordered residues between proteins encoded by highly and lowly expressed genes, the lowly expressed group showed stronger signal (higher Z-score) in *H. sapiens* and *S. cerevisiae* but not in *E. coli*. Previously, a weak positive correlation was noted between gene expression level and protein disorder in *E. coli* suggesting that since prokaryotes encode relatively lower fraction of disordered residues expression level may not be a strong burden for disordered proteins in these organisms [48]. In accordance, in *E. coli* we did not find a prominent difference in Z-score between highly and lowly expressed groups which may suggest that the trends that we observed at the protein level are independent of gene expression at the transcript level in prokaryotes.

In conclusion, in this paper, we analyzed the signals related to the selection of protein disorder in several eukaryotic and prokaryotic organisms. Our study suggested that disordered residues are preferred in eukaryotic proteins over random expectations and this preference is stronger near the protein terminals. On the other hand, prokaryotic proteins show either no or weak such signal. Based on our observations we proposed several explanations, however here we would like to re-emphasize that most of these are predictive in nature. Therefore, further experiments are needed to understand the cause(s) and consequence(s) of the trends shown here. Moreover, in this analysis, we mainly compared protein disorder between the two major groups, eukaryotes and prokaryotes where the observed trends are distinct and very clear. However, there is wide variation in the strength of selection among the genomes of each domain (eukaryotes or prokaryotes). Each organism was suggested to specifically tailor the level of disordered residues in its proteome according to its functional and environmental prerequisite [13–15]. Therefore, one important direction for future research could be to explore the factors responsible for this intra-domain variation in protein disorder.

## Materials and methods

### Data collection

For our study we considered 12 model species, six prokaryotes and six eukaryotes: *H. sapiens*, one insect: *D. melanogaster*, one worm: *C. elegans*, three fungi: *S. cerevisiae*, *A. oryzae* and *N. crassa*, three bacteria: *B. subtilis*, *E. coli*, *D. radiodurans* and three archaea: *M. mazei*, *H. volcanii*, *T. gammatolerans*. All these organisms are listed in Table S1 with details of their characteristics such as their accession number, strain description, genome size, number of predicted proteins, *etc*. We choose these organisms following Faure et al. [49] considering our objective to get a general overview in the eukaryotic and the prokaryotic domain. Except for *H. sapiens* (downloaded from Ensembl v-90 [50]) and *S. cerevisiae* (downloaded from the *Saccharomyces* Genome Database [51]) protein coding sequences and complete proteomes of all the remaining species were retrieved from the NCBI Reference Sequence database (RefSeq) [52] (release 88) (last accessed 6^th^ March‟ 2018). Here we choose our genomes mostly from the NCBI RefSeq database because the protein sequences in RefSeq are non-redundant, well-annotated, and explicitly linked with their nucleotide sequences [52]. To ensure the quality of genomes, RefSeq imposes strict evaluation criteria and corrects annotation manually when necessary [52]. Further, RefSeq considers spliced variants only when there is experimental evidence against their full-length nature thus only a minimal set of splice variants are included in this database. Therefore, chances that our results may be biased by the over-representation of some specific splice variants are very low. On the other hand, Ensembl is also a database of high quality genomes [50], however it includes a huge number of transcript variants. Therefore, to remove redundancy in our dataset (human) we took the longest isoform when we found more than one isoform for a gene.

For a species, if genomes of multiple strains are available we choose the strains annotated as the reference or representative strain. In each species, proteins containing ambiguous amino acids (B, J, O, U, X, Z) and internal stop codons or partial codons in their corresponding CDS sequences were removed and only proteins more than 50 amino acids in length were considered for proteome-wide disorder prediction.

### Generation of random models

To serve specific purposes of our study we generated three kinds of random protein models for each species. First to compare the protein-wise disorder score between real and random sequences we generated random sequences preserving the overall amino acid composition and length of each real protein (designated as length conserved random model). For this purpose, we randomly shuffled the amino acids of each real protein 10 times (total number of length conserved random sequences of each species = 10 × number of real sequences). Next, to compare position-wise disorder score between real and random sequences we looked for random models that would preserve position-specific amino acid characteristics. For this, we at first generated random sequences (designated as terminal residues conserved random model) by shuffling the amino acids at the N and C-terminals of each real protein. This random model is analogous to our length conserved random model however here we considered only the first and last 200 amino acids (of real proteins more than 200 amino acids in length) for random shuffling. As with our length conserved random, we generated 10 such random sequences for each terminal corresponding to each real protein (total number of terminal residue random sequences for each species = 10 × 2 × number of real sequences). Next, we aligned the naturally occurring proteins of each species from both ends (*i.e.* from the first amino acid position to the end of the protein and from the last amino acid position to the beginning of the protein) and shuffled the amino acids in each position of the alignment (up to 200 positions) column-wise. Thus it preserves the overall amino acid frequencies at each position of real proteins. To account for the variation in protein length here we considered up to 200 positions from both ends considering only proteins more than 200 amino acids in length. For each species, we generated 200,000 (100,000 for each terminal) such random sequences (designated as column-wise random model). All these three types of random models are illustrated in Figure 1. Here it is noteworthy that for the generation of random sequences (model 2, 3) we considered up to 200 positions from both ends, however, position-specific disorder scores were calculated up to 150 positions (discussed in the next section).

### Prediction of disordered residues

#### Consensus approach #01

Ordered or disordered status of each residue in each protein was estimated by consensus-based approach. At first, the per-residue disorder scores were predicted using four different disorder prediction algorithms namely IUPred [53, 54], VLS2B [55], MoreRONN [56], and DisEMBL [57]. If a residue is predicted as disordered (or predicted disorder score > 0.5) by three of these four algorithms then the residue is considered as a disordered residue. We choose these algorithms because these algorithms do not use any homology profile thus were expected to give an unbiased estimate of disorder scores when applied to random protein models [24]. Among these, IUPred predicts disorder scores based on inter-residue interactions probabilities (estimated through pair-wise interaction energies) [54]. There are two different variants of IUPred, IUPred-S: optimized for short disordered regions and IUPred-L: optimized for long disordered regions [54]. Here we used IUPred-L, which was shown to perform better than IUPred-S in general benchmark datasets [58]. VLS2B, a neural network based algorithm gives a weighted average of disordered scores from two different disorder prediction algorithms trained on short and long proteins [55]. VSL2B was shown to be among the top performing algorithms when compared with 16 other disorder predictors with a benchmark dataset of 514 experimentally verified disordered proteins [59]. MoreRONN is the upgraded version of well known neural network based disordered predictor RONN [56], which was considered as the best of all other 9 disorder prediction algorithms in CASP6 assessment [60]. The standalone version of MoreRONN (https://app.strubi.ox.ac.uk/MoreRONN/) was kindly provided by the RONN [56] developing team prior to their publication. DisEMBL is an artificial neural networks based algorithm which is trained on three different types of disorder dataset: coils, hot loops and missing coordinates in X-Ray structure [57]. Here we considered the version of DisEMBL which defines disordered based on missing coordinates in X-Ray structure (DisEMBL-465). Considering segment overlap measure (SOV), DisEMBL-465 was suggested as the best of 10 other disorder prediction algorithms in a recent assessment [61]. All these algorithms were run locally using default parameter settings. Most of these disorder prediction algorithms use sequence context (*i.e.* neighborhood) therefore for prediction of disordered residues of random protein models generated from truncated sequences (model 2, 3) we utilized extra 50 residues from both ends *i.e.* for disorder prediction we used sequences of 200 residues however considered up to first 150 positions in all subsequent disorder calculation.

#### Consensus approach #02

Except for MoreRONN [56] (upgraded in 2017), all other algorithms like IUPred [53, 54], DisEMBL [57] and VSL2B [55] were published more than a decade ago. Therefore, during the revision process, we were suggested to check our results with comparatively newer disorder prediction algorithms. To further validate our results here we considered three algorithms which are single sequence based, not used in our initial analysis and published or upgraded in a recent time frame. In the way of our search for new algorithms, we noticed that a newer version of IUPred (IUPred 2A) has been published last year [25]. Considering the fact that we have a very heterogeneous dataset (real and random proteins) which may have long or short disordered regions we decided to use both the verities of IUPred2A *i.e*. short and long. We found another well known single sequence based disorder predication algorithm Espritz, which is published few years back [62]. To predict disordered regions, ESpritz employs bi-directional recursive neural network-based machine learning method and was tested to perform better than most of the other disorder prediction algorithms [62]. Based on the training dataset, Espritz has three variants (*i*) Espritz-N (trained on NMR mobility dataset), (*ii*) Espritz-X (trained on proteins with known X-ray crystal structure), and (*iii*) Espritz-D (trained on proteins from DisProt database) [62]. Here we considered all these three variants with the option to maximize accuracy threshold (best Sw score) and ignoring PSI blast option. Recently a new single-sequence based disorder prediction algorithm was developed which was suggested to be more accurate than sequence profile based prediction algorithms for proteins with limited number of homologous [63]. This algorithm knows as SPOT-Disorder-Single is based on an ensemble of current state-of-art-neural networks [63]. Here we downloaded its standalone version and run with default settings. To determine consensus between these algorithms first we considered each variant as independent predictors and calculated their majority vote *i.e.* a residue was considered to be disordered if at least 4 of 6 algorithms (2 variants of IUPred2A, 3 variants of Espritz and SPOT-Disorder-Single) suggested so. Next, we calculated consensus between three algorithms IUPred2A, Espritz and SPOT-Disorder-Single considering their equal weightage. For this, we first inferred ordered or disordered status of each residue by integrating prediction from the different variants of IUPred2A and Espritz into a final prediction (here denoted as IUPred2A-meta and Espritz-meta). For IUPred2A-meta, we considered a residue as disordered if average disorder score predicted by its two variants (long and short) >0.5. For Esprtiz-meta, we considered mutual agreement between 2 of its 3 variants. The prediction from IUPred2A-meta and Espritz-meta is then combined with that of SPOT-Disorder-Single and the final consensus was calculated considering their majority vote. Overall, we found very similar results by both these ways of calculating consensus; therefore, here we presented results obtained by one of these methods (consensus calculated considering equal weightage of each algorithm).

We tested the performance of the algorithms used for this study on experimentally known disorder protein dataset (check Table S2 for details). Here we measured correlations between the proportions of disordered residues predicted by each of these algorithms in terminal regions (up to the first and last 150 residues) of these proteins with that calculated based on experimental annotation. Strong correlations between the predicted and experimental values for most of disorder prediction algorithms suggest that these algorithms can reliably predict disorder residues in test proteins (Table S2).

### Calculation of protein disorder content and position-wise disorder scores

To get relative estimates of structural disorder in the native sequences of each species here we first compared their disorder content with that of length conserved random sequences. Overall disorder content of proteins in each group (real vs. random of each species) was calculated as the average percentage of disordered residues (predicted by a consensus-based approach) in the proteins of that group. To reduce the effects of short flexible loops in disorder prediction [10], we considered another measure of protein disorder content, the percentage of dis_30 residues, calculated as the percentage disordered residues only in long disordered segments (more than 30 or more consecutive disordered residues).

Next, in order to check whether there is any site-specific selection for high or low disorder in naturally occurring protein sequences, we compared the position-specific disorder scores (up to first and last 150 residues) of real sequences with that of two kinds of random protein models (random models 2, 3) which were specifically generated to take into account for the position-specific bias in amino acid composition. To calculate position-specific disorder scores of real proteins, naturally occurring sequences of each species were lined up according to their start position and again according to their end position. Position-specific disorder score of real proteins at any position ‘i’ was calculated as the fraction residues predicted as disordered residues (by consensus-based approach) at that position (‘i’) to the total number of sequences in the alignment (here ‘i’ equal to 1 to 150 from both ends). Position-specific disorder scores of random sequences were calculated in the same way as the real sequences. However, for the estimation of Z-score, we divided the terminal residue conserved and column random sequences of each species into 100 randomized proteomes and estimated position-specific disorder scores for each such randomized proteome. For each position “i” of each such random model we then computed the mean and standard deviation of position-wise disorder score over the 100 randomized proteomes.

We then compared the position-wise disorder score of real protein sequences with that of random protein models at the corresponding position thorough Z-score defined as

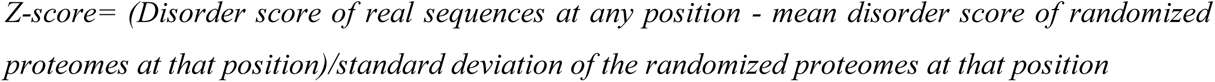

At any particular position, a positive Z-score indicates the enrichment of disordered residues in the naturally occurring sequences while a negative Z-score indicates its enrichment in random sequences. Statistical significance of the difference is accessed through *P*-values calculated via integrating over the relevant area of the related normal distribution. Here *P-*values were calculated following two-tailed distribution considering the probability that disordered scores of real sequences can be higher than that of random sequences or vice-versa.

### Calculation of GC content and generation of random models

GC content of protein-coding sequences was calculated using CodonW (http://codonw.sourceforge.net). To compare disorder content of real and random proteins in GC bins, proteins were grouped according to the GC content of their coding sequences. To encompass the maximum number of sequences while maintaining a uniform GC variation GC bins were chosen in steps of 10% GC variation whereas starting points are adjusted according to the distribution of genic GC content of each species. For instance, in eukaryotes, GC bins were chosen starting from 40% genic GC except for two low GC genomes *D. melanogaster* and *N. crassa* where GC bins were set starting from 30% genic GC. In prokaryotes, GC bins were chosen starting from 30% genic GC except for two high genomes *H. volcanii* and *D. radiodurans* where the minimum threshold for GC bins was set at 50% genic GC. GC bins defined by this way include >99% of protein-coding sequences in each species. For position-wise comparison of GC content between real and random sequences we generated random nucleotide models analogous to terminal residue conserved and column random protein models (models 2, 3). For terminal residue conserved nucleotide model, we shuffled the codons in the first and last 450 positions of coding sequences corresponding to first and last 150 positions of protein sequences. For column random model we aligned the codons of real sequences from both ends and then shuffled the codons in each position of the alignment. Z-score and *P*-values were calculated by similar approach as utilized for comparing disorder score.

### Categorization according to GO slim analysis and gene expression level

GO slim functional annotations (total 130 categories) of human protein-coding genes were retrieved from Ensembl 90 [50]. GO slim categories with less than 100 genes were removed finally 124 categories were considered. Next, we compared the position-specific disorder score of real sequences under each such category following Z-score approach as described previously with two kinds of random protein models (model 2 and 3). For terminal residue conserved model (model 2) we considered the random sequences corresponding to real sequences under each GO slim category and column random sequences (model 3) were generated separately for each GO slim category from the alignment of proteins in that category. For the expression level of human proteins at the transcript level, we considered high-throughput RNA-seq gene expression data of Uhlén et al. [64]. In this dataset FPKM (fragments per kilobase of exon model per million mapped reads) level of 20,344 genes across 32 normal human tissues has been provided. From this dataset, we retrieved tissue-averaged gene expression level of 18497 of 19961 human proteins considered in this study. Gene expression level of *E. coli* (4022 of 4064 proteins) and *S. cerevisiae*, (5122 of 5848) proteins were retrieved from Geo Gene Expression Omnibus [65] (https://www.ncbi.nlm.nih.gov/geo/query/acc.cgi?acc=GSE67218) and publication of Ghaemmaghami et al. [66] respectively. For our comparative study, in each of these three species, we compared the Z-scores of predicted disorder between the top and bottom 20% of proteins sorted according to their gene expression level. Z-scores of each group of high and low expressed proteins were calculated in reference to their corresponding terminal residue conserved random sequences and column random sequences. Column random sequences were generated for each group of highly and lowly expressed proteins separately from the alignment of proteins in that group.

### Prediction disordered binding sites with ANCHOR

To identify the residues along the proteins those may be important for protein binding we considered ANCHOR [31]. ANCHOR predicts the elements within disordered regions those play crucial roles in molecular recognition and protein-protein interactions. To predict such residues, it utilizes the same general principle (pair-wise inter-residue interaction energies) as the IUPred disorder prediction algorithm [31]. However, the basic difference between these two algorithms is that while IUPred predicts all the potential disordered regions within a protein, ANCHOR predicts the disordered regions which could act as potential protein-protein binding sites [31]. Thus the scores predicted by ANCHOR were suggested to be independent of IUPred algorithm [31]. Although these regions are usually very short in length and are associated with very high false positive detection rate, ANCHOR was shown to achieve an accuracy rate of average ∼ 67% when tested with different benchmark datasets [31]. For each position (first and last 150 positions) we compared the proportion of disordered binding residues predicted by ANCHOR between real and random protein sequences (random model 2, 3) of each species following Z-score approach as described for position-specific disorder score. The proportion of disordered binding regions in each position was calculated as the number of residues predicted as disordered binding regions at that position divided by the total number of sequences in the dataset.

### Prediction of solvent accessible surface area

Solvent accessible surface area (ASA) of each protein in our datasets was predicted using the standalone version of SPIDER3-Single algorithm [67]. SPIDER3-Single predicts the extent of solvent exposure of each residue within a protein along with several other structural information based on sequence information only *i.e.* without using homology profile. This algorithm uses similar types of neural networks as SPOT-Disorder-Single [67], however, considers different sets of input features. As tested by the authors, this algorithm is more accurate than other state-of-art ASA prediction algorithms such as ASAquick or profile-based SPIDER3 [67]. Here we calculated Z-score for predicted ASA following the steps as described for disorder prediction. For the sake of brevity, this test was done only for three species *H. sapiens*, *E. coli* and *S. Cerevisiae* using the same real and random (terminal residue conserved and column random) datasets as used for disorder prediction.

### Statistical analyses

All statistical tests were performed using R (The R Project for Statistical Computing [68]). We compared the percentages of disordered residues (not normally distributed) between real and random datasets using two samples “Wilcox.test” (commonly known as the Mann-Whitney U test) function of R. Following non-parametric distribution, all the correlation analyses were performed using Spearman’s Rank correlation test and significant levels were denoted with *P*-values.

## Authors’ contributions

TT and AP conceived the study and designed the methodologies. AP collected the data and performed all the experiments under the guidance of TT. AP and TT analyzed the data and wrote the manuscript. Both the authors have read and approved the final manuscript.

## Acknowledgments

The authors thank Mr. Michael Peeri for technical support. The authors like to thank the Israeli Concil of Higher Education and Research for granting Planning and Budgeting Committee (PBC) outstanding post-doctoral research fellowship to AP.

## Supplementary material

### Supplementary Figure legends

**Figure S1 Proteome-wise comparison of disorder content between real and random sequences predicted by consensus approach #2**

The graph shows the average disorder content of real and random proteins of each of the test species. Disordered content is calculated as the percentage of disordered residues in each protein (predicted by the consensus approach #2, see main text for details) then averaged over all the proteins in each group (real or random). Disordered content is calculated considering all predicted disordered residues (denoted as percentages of all disordered residues) and considering disordered residues only in long disordered regions (30 or more consecutive disordered residues) (denoted as % of DIS_30 residues). So, there are two panels for each species, each panel showing the proportion of disordered residues between real and random sequences calculated by these two approaches. *P*-values were calculated by Mann-Whitney U test comparing disordered content between real and random sequences of each species. Here error bars show standard error at 95% confidence interval. Significant difference (*P* < 0.05) was shown with *. Here *P* stands for the *P*-values obtained by Mann-Whitney U test.

**Figure S2 Species to species comparison of the distribution of charged residues between the test prokaryotic and eukaryotic organisms**

The graph shows the average % of charged residues in the proteomes of our test organisms. Here we compared the proportion of charged residues of each eukaryotic species (denoted as species B) with that of each prokaryotic species (denoted as species A). *P*-values were calculated by Mann-Whitney U test. Here error bars show standard error at 95% confidence interval. Significant difference (*P* < 0.05) was shown with *. Here *P* stands for the *P*-values obtained by Mann-Whitney U test.

**Figure S3 Species to species comparison of the distribution of polar residues between the test prokaryotic and eukaryotic organisms**

The graph shows the average % of polar residues in the proteomes of our test organisms. Here we compared the proportion of polar residues of each eukaryotic species (denoted as species B) with that of each prokaryotic species (denoted as species A). *P*-values were calculated by Mann-Whitney U test. Here error bars show standard error at 95% confidence interval. Significant difference (*P* < 0.05) was shown with *. Here *P* stands for the *P*-values obtained by Mann-Whitney U test.

**Figure S4 Comparison of disorder content between real and random sequences of eukaryotic species in GC bins**

In each species, naturally occurring protein sequences were grouped in GC bins according to the GC content of their coding sequences then their disorder content was compared with that of their corresponding length conserved random sequences. In each species, GC bins were defined following distribution of genic GC content in steps of 10% GC variation (see main text). Disorder content of both real and random datasets was calculated by the consensus approach #1 (see main text) considering (*i*) all the predicted disordered residues (denoted as percentages of all disordered residues), and (*ii*) considering disordered residues only in long disordered regions (denoted as % of DIS_30 residues). So, in each GC bin there are two panels, each panel showing the proportion of disordered residues between real and random sequences calculated by these two approaches. *P*-values were calculated by Mann-Whitney U test comparing disorder content between real and random sequences in each GC bin. Here error bars show standard error at 95% confidence interval. Significant difference (*P* < 0.05) was shown with *. Here *P* stands for the *P*-values obtained by Mann-Whitney U test.

**Figure S5 Comparison of disorder content between real and random sequences of prokaryotic species in GC bins**

In each species, naturally occurring protein sequences were grouped in GC bins according to the GC content of their coding sequences then their disorder content was compared with that of their corresponding length conserved random sequences. In each species, GC bins were defined following distribution of genic GC content in steps of 10% GC variation (see main text). Disorder content of both real and random datasets was calculated by consensus approach #1 (see main text) considering (*i*) all the predicted disordered residues (denoted as percentages of all disordered residues), and (*ii*) considering disordered residues only in long disordered regions (denoted as % of DIS_30 residues). So, in each GC bin there are two panels, each panel showing the proportion of disordered residues between real and random sequences calculated by these two approaches. *P*-values were calculated by Mann-Whitney U test comparing disordered content between real and random sequences in each GC bin. Here error bars show standard error at 95% confidence interval. Significant difference (*P* < 0.05) was shown with *. Here *P* stands for the *P*-value obtained by Mann-Whitney U test.

**Figure S6 Z—score profile for the position-specific disorder score of each species predicted by consensus approach #2**

This figure shows the extent of protein disorder between real and random protein models for the first 150 and last 150 positions of 12 organisms considered in this study predicted by consensus approach #2. In panels **A** and **B,** Z-scores were calculated comparing position-specific disorder scores of real and terminal residue conserved random sequences of each species and in panels **C** and **D,** Z-scores were calculated comparing position-specific disorder scores of real and column random sequences of each species. The color scale for each graph is shown on the top of each panel. Here positive Z-score indicates enrichment of protein disorder in naturally occurring sequences while negative Z-score indicates the reverse. Organisms are arranged according to their mid-point rooted species tree retrieved from NCBI taxonomic database with the help of their species taxonomic identifier.

**Figure S7 Z—score profile for position-specific disorder scores of *H. sapiens***

This figure shows the extent of protein disorder between real and random protein models for the first 150 and last 150 positions of human proteins predicted by consensus approach #1 (panels **A**, **B**, **C**, **D**) and consensus approach #2 (panels **E**, **F**, **G**, **H**). Panels **A, B, E, F:** Comparison of real and terminal residue conserved random sequences. Panels **C, D, G, H:** Comparison of real and column random sequences. In each panel, vertical blue lines stand for the average position-specific disorder scores of real proteins and grey lines that for random proteins. At each position, the statistical difference between these two measurements is accessed through Z-score (see main text) and is shown with blue line. Corresponding *P*-value at each position is shown either with a green (*P* < 0.05) or red dot (*P* >= 0.05) above the Z-score. Here positive Z-score indicates enrichment of protein disorder in naturally occurring sequences while negative Z-score indicates the reverse.

**Figure S8 Z—score profile for position-specific disorder scores of *D. melanogaster***

This figure shows the extent of protein disorder between real and random protein models for the first 150 and last 150 positions of *D. melanogaster* proteins predicted by consensus approach #1 (panels **A**, **B**, **C**, **D**) and consensus approach #2 (panels **E**, **F**, **G**, **H**). Figure representation is same as Figure S7.

**Figure S9 Z—score profile for position-specific disorder scores of *C. elegans***

This figure shows the extent of protein disorder between real and random protein models for the first 150 and last 150 positions of *C. elegans* proteins predicted by consensus approach #1 (panels **A**, **B**, **C**, **D**) and consensus approach #2 (panels **E**, **F**, **G**, **H**). Figure representation is same as Figure S7.

**Figure S10 Z—score profile for position-specific disorder scores of *S. cerevisiae***

This figure shows the extent of protein disorder between real and random protein models for the first 150 and last 150 positions of *S. cerevisiae* proteins predicted by consensus approach #1 (panels **A**, **B**, **C**, **D**) and consensus approach #2 (panels **E**, **F**, **G**, **H**). Figure representation is same as Figure S7.

**Figure S11 Z—score profile for position-specific disorder scores of *A. oryzae***

This figure shows the extent of protein disorder between real and random protein models for the first 150 and last 150 positions of *A. oryzae* proteins predicted by consensus approach #1 (panels **A**, **B**, **C**, **D**) and consensus approach #2 (panels **E**, **F**, **G**, **H**). Figure representation is same as Figure S7.

**Figure S12 Z—score profile for position-specific disorder scores of *N. crassa***

This figure shows the extent of protein disorder between real and random protein models for the first 150 and last 150 positions of *N. crassa* proteins predicted by consensus approach #1 (panels **A**, **B**, **C**, **D**) and consensus approach #2 (panels **E**, **F**, **G**, **H**). Figure representation is same as Figure S7.

**Figure S13 Z—score profile for position-specific disorder scores of *B. subtilis***

This figure shows the extent of protein disorder between real and random protein models for the first 150 and last 150 positions of *B. subtilis* proteins predicted by consensus approach #1 (panels **A**, **B**, **C**, **D**) and consensus approach #2 (panels **E**, **F**, **G**, **H**). Figure representation is same as Figure S7.

**Figure S14 Z—score profile for position-specific disorder scores of *E. coli***

This figure shows the extent of protein disorder between real and random protein models for the first 150 and last 150 positions of *E. coli* proteins predicted by consensus approach #1 (panels **A**, **B**, **C**, **D**) and consensus approach #2 (panels **E**, **F**, **G**, **H**). Figure representation is same as Figure S7.

F**igure S15 Z—score profile for position-specific disorder scores of *D. radiodurans***

This figure shows the extent of protein disorder between real and random protein models for the first 150 and last 150 positions of *D. radiodurans* proteins predicted by consensus approach #1 (panels **A**, **B**, **C**, **D**) and consensus approach #2 (panels **E**, **F**, **G**, **H**). Figure representation is same as Figure S7.

**Figure S16 Z—score profile for position-specific disorder scores of *M. mazei***

This figure shows the extent of protein disorder between real and random protein models for the first 150 and last 150 positions of *M. mazei* proteins predicted by consensus approach #1 (panels **A**, **B**, **C**, **D**) and consensus approach #2 (panels **E**, **F**, **G**, **H**). Figure representation is same as Figure S7.

**Figure S17 Z—score profile for position-specific disorder scores of *H. volcanii***

This figure shows the extent of protein disorder between real and random protein models for the first 150 and last 150 positions of *H. volcanii* proteins predicted by consensus approach #1 (panels **A**, **B**, **C**, **D**) and consensus approach #2 (panels **E**, **F**, **G**, **H**). Figure representation is same as Figure S7.

**Figure S18 Z—score profile for position-specific disorder scores of *T. gammatolerans***

This figure shows the extent of protein disorder between real and random protein models for the first 150 and last 150 positions of *T. gammatolerans* proteins predicted by consensus approach #1 (panels **A**, **B**, **C**, **D**) and consensus approach #2 (panels **E**, **F**, **G**, **H**). Figure representation is same as Figure S7.

**Figure S19 Z—score profile for position-specific disorder scores of *H. sapiens* proteins predicted by IUPred2A (version long)**

This figure shows the extent of protein disorder between real and random protein models for the first 150 and last 150 positions of *H. sapiens* proteins predicted by IUPred2A algorithm (version long). **A, B:** comparison of real and terminal residue conserved random sequences. **C, D:** Comparison of real and column random sequences. In each panel, vertical blue lines stand for the average position-specific disorder scores of real proteins and grey lines that for random proteins. At each position, the statistical difference between these two measurements is accessed through Z-score (see main text) and is shown with blue line. Corresponding *P*-value at each position is shown either with a green (*P* < 0.05) or red dot (*P* >= 0.05) above the Z-score. Here positive Z-score indicates enrichment of protein disorder in naturally occurring sequences while negative Z-score indicates the reverse.

**Figure S20 Z—score profile for position-specific disorder scores predicted by IUPred2A (version long) of *D. melanogaster***

This figure shows the extent of protein disorder between real and random protein models for the first 150 and last 150 positions of *D. melanogaster* proteins predicted by IUPred2A algorithm (version long). Figure representation is same as Figure S19.

**Figure S21 Z—score profile for position-specific disorder scores predicted by IUPred2A (version long) of *C. elegans***

This figure shows the extent of protein disorder between real and random protein models for the first 150 and last 150 positions of *C. elegans* proteins predicted by IUPred2A algorithm (version long). Figure representation is same as Figure S19.

**Figure S22 Z—score profile for position-specific disorder scores predicted by IUPred2A (version long) of *A. oryzae***

This figure shows the extent of protein disorder between real and random protein models for the first 150 and last 150 positions of *A. oryzae* proteins predicted by IUPred2A algorithm (version long). Figure representation is same as Figure S19.

**Figure S23 Z—score profile for position-specific disorder scores predicted by IUPred2A (version long) of *N. crassa***

This figure shows the extent of protein disorder between real and random protein models for the first 150 and last 150 positions of *N. crassa* proteins predicted by IUPred2A algorithm (version long). Figure representation is same as Figure S19.

**Figure S24 Z—score profile for position-specific disorder scores predicted by IUPred2A (version long) of *B. subtilis***

This figure shows the extent of protein disorder between real and random protein models for the first 150 and last 150 positions of *B. subtilis* proteins predicted by IUPred2A algorithm (version long). Figure representation is same as Figure S19.

**Figure S25 Z—score profile for position-specific disorder scores predicted by IUPred2A (version long) of *E. coli***

This figure shows the extent of protein disorder between real and random protein models for the first 150 and last 150 positions of *E. coli* proteins predicted by IUPred2A algorithm (version long). Figure representation is same as Figure S19.

**Figure S26 Z—score profile for position-specific disorder scores predicted by IUPred2A (version long) of *D. radiodurans***

This figure shows the extent of protein disorder between real and random protein models for the first 150 and last 150 positions of *D. radiodurans* proteins predicted by IUPred2A algorithm (version long). Figure representation is same as Figure S19.

**Figure S27 Z—score profile for position-specific disorder scores predicted by IUPred2A (version long) of *M. mazei***

This figure shows the extent of protein disorder between real and random protein models for the first 150 and last 150 positions of *M. mazei* proteins predicted by IUPred2A algorithm (version long). Figure representation is same as Figure S19.

**Figure S28 Z—score profile for position-specific disorder scores predicted by IUPred2A (version long) of *H. volcanii***

This figure shows the extent of protein disorder between real and random protein models for the first 150 and last 150 positions of *H. volcanii* proteins predicted by IUPred2A algorithm (version long). Figure representation is same as Figure S19.

**Figure S29 Z—score profile for position-specific disorder scores predicted by IUPred2A (version long) of *T. gammatolerans***

This figure shows the extent of protein disorder between real and random protein models for the first 150 and last 150 positions of *T. gammatolerans* proteins predicted by IUPred2A algorithm (version long). Figure representation is same as Figure S19.

**Figure S30 Z—score profile for position-specific disorder scores of *A. oryzae* after removing the first and last 50 residues**

For this test we removed the terminal regions (up to the first and last 50 amino acid positions) from the extant protein sequences of *A. oryzae* and generated random models (column random and terminal residue random models) from this terminal regions removed real protein dataset. We freshly predicted disordered residues in these sequences (both real and random) following consensus approach #1 and analyzed position-wise Z-score profile as we did for the full-length sequences (see main text). This figure shows the extent of protein disorder between real and random protein models (generated for this test) up to the first 100 and last 100 positions. **A.** and **B.**:

Comparison of real and terminal residue conserved random sequences. **C.** and **D.**: Comparison of real and column random sequences. In each panel, vertical blue lines stand for the average position-specific disorder scores of real proteins and grey lines that for random proteins. At each position, the statistical difference between these two measurements is accessed through Z-score (see main text) and is shown with blue line. Corresponding *P*-value at each position is shown either with a green (*P* < 0.05) or red dot (*P* >= 0.05) above the Z-score. Here positive Z-score indicates enrichment of protein disorder in naturally occurring sequences while negative Z-score indicates the reverse.

**Figure S31 Z—score profile for position-specific disorder scores of *C. elegans* after removing the first and last 50 residues**

For this test we removed the terminal regions (up to the first and last 50 amino acid positions) from the extant protein sequences of *C. elegans* and generated random models (column random and terminal residue random models) from this terminal regions removed real protein dataset. We freshly predicted disordered residues in these sequences (both real and random) following consensus approach #1 and analyzed position-wise Z-score profile as we did for the full-length sequences (see main text). This figure shows the extent of protein disorder between real and random protein models (generated for this test) up to the first 100 and last 100 positions. **A.** and **B.**: Comparison of real and terminal residue conserved random sequences. **C.** and **D.**: Comparison of real and column random sequences. In each panel, vertical blue lines stand for the average position-specific disorder scores of real proteins and grey lines that for random proteins. At each position, the statistical difference between these two measurements is accessed through Z-score (see main text) and is shown with blue line. Corresponding P-value at each position is shown either with a green (*P* < 0.05) or red dot (*P* >= 0.05) above the Z-score. Here positive Z-score indicates enrichment of protein disorder in naturally occurring sequences while negative Z-score indicates the reverse.

**Figure S32 Z—score profile for position-specific disorder scores of *D. melanogaster* after removing the first and last 50 residues**

For this test we removed the terminal regions (up to the first and last 50 amino acid positions) from the extant protein sequences of *D. melanogaster* and generated random models (column random and terminal residue random models) from this terminal regions removed real protein dataset. We freshly predicted disordered residues in these sequences (both real and random) following consensus approach #1 and analyzed position-wise Z-score profile as we did for the full-length sequences (see main text). This figure shows the extent of protein disorder between real and random protein models (generated for this test) up to the first 100 and last 100 positions. **A.** and **B.**: Comparison of real and terminal residue conserved random sequences. **C.** and **D.**: Comparison of real and column random sequences. In each panel, vertical blue lines stand for the average position-specific disorder scores of real proteins and grey lines that for random proteins. At each position, the statistical difference between these two measurements is accessed through Z-score (see main text) and is shown with blue line. Corresponding *P*-value at each position is shown either with a green (*P* < 0.05) or red dot (*P* >= 0.05) above the Z-score. Here positive Z-score indicates enrichment of protein disorder in naturally occurring sequences while negative Z-score indicates the reverse.

**Figure S33 Z—score profile for position-specific disorder scores of *H. sapiens* after removing the first and last 50 residues**

For this test we removed the terminal regions (up to the first and last 50 amino acid positions) from the extant protein sequences of *H. sapiens* and generated random models (column random and terminal residue random models) from this terminal regions removed real protein dataset. We freshly predicted disordered residues in these sequences (both real and random) following consensus approach #1 and analyzed position-wise Z-score profile as we did for the full-length sequences (see main text). This figure shows the extent of protein disorder between real and random protein models (generated for this test) up to the first 100 and last 100 positions. **A.** and **B.**: Comparison of real and terminal residue conserved random sequences. **C.** and **D.**: Comparison of real and column random sequences. In each panel, vertical blue lines stand for the average position-specific disorder scores of real proteins and grey lines that for random proteins. At each position, the statistical difference between these two measurements is accessed through Z-score (see main text) and is shown with blue line. Corresponding *P*-value at each position is shown either with a green (*P* < 0.05) or red dot (*P* >= 0.05) above the Z-score. Here positive Z-score indicates enrichment of protein disorder in naturally occurring sequences while negative Z-score indicates the reverse.

**Figure S34 Z—score profile for position-specific disorder scores of *N. crassa* after removing the first and last 50 residues.**

For this test we removed the terminal regions (up to the first and last 50 amino acid positions) from the extant protein sequences of *N. crassa* and generated random models (column random and terminal residue random models) from this terminal regions removed real protein dataset. We freshly predicted disordered residues in these sequences (both real and random) following consensus approach #1 and analyzed position-wise Z-score profile as we did for the full-length sequences (see main text). This figure shows the extent of protein disorder between real and random protein models (generated for this test) up to the first 100 and last 100 positions. **A.** and **B.**: Comparison of real and terminal residue conserved random sequences. **C.** and **D.**: Comparison of real and column random sequences. In each panel, vertical blue lines stand for the average position-specific disorder scores of real proteins and grey lines that for random proteins. At each position, the statistical difference between these two measurements is accessed through Z-score (see main text) and is shown with blue line. Corresponding *P*-value at each position is shown either with a green (*P* < 0.05) or red dot (*P* >= 0.05) above the Z-score. Here positive Z-score indicates enrichment of protein disorder in naturally occurring sequences while negative Z-score indicates the reverse.

**Figure S35 Z—score profile for position-specific disorder scores of *S. cerevisiae* after removing the first and last 50 residues**

For this test we removed the terminal regions (up to the first and last 50 amino acid positions) from the extant protein sequences of *S. cerevisiae* and generated random models (column random and terminal residue random models) from this terminal regions removed real protein dataset. We freshly predicted disordered residues in these sequences (both real and random) following consensus approach #1 and analyzed position-wise Z-score profile as we did for the full-length sequences (see main text). This figure shows the extent of protein disorder between real and random protein models (generated for this test) up to the first 100 and last 100 positions. **A.** and **B.**: Comparison of real and terminal residue conserved random sequences. **C.** and **D.**: Comparison of real and column random sequences. In each panel, vertical blue lines stand for the average position-specific disorder scores of real proteins and grey lines that for random proteins. At each position, the statistical difference between these two measurements is accessed through Z-score (see main text) and is shown with blue line. Corresponding *P*-value at each position is shown either with a green (*P* < 0.05) or red dot (*P* >= 0.05) above the Z-score. Here positive Z-score indicates enrichment of protein disorder in naturally occurring sequences while negative Z-score indicates the reverse.

**Figure S36 Z—score profile for GC content for the 12 species considered in this study**

This figure shows the Z-score profile calculated comparing GC content of real and randomly generated nucleotide sequences for the first 150 and last 150 positions of 12 organisms considered in this study. In panels **A** and **B,** Z-scores were calculated comparing GC content of real and terminal residue conserved random sequences of each species and in panels **C** and **D,** Z-scores were calculated comparing GC content of real and column random sequences of each species. The color scale for each graph is shown on the top of each panel. Here positive Z-score indicates enrichment of GC content in naturally occurring sequences while negative Z-score indicates the reverse. Organisms are arranged according to their mid-point rooted species tree retrieved from NCBI taxonomic database with the help of their species taxonomic identifier.

**Figure S37 Z—score profile for position-specific GC scores of *H. sapiens***

This figure shows the extent of GC content between the coding sequences of real and random models for the first 150 and last 150 amino acid positions. **A.** and **B.** Comparison of real and terminal residue conserved random sequences. **C.** and **D.** Comparison of real and column random sequences. In each panel, blue lines stand for the average GC content of real proteins coding sequences and black lines that for random DNA sequences. At each position, the statistical difference between these two measurements is accessed through Z-score (see main text) and is shown with blue line. Corresponding *P*-value at each position is shown either with a green (*P* < 0.05) or red dot (*P* >= 0.05) above the Z-score. Here positive Z-score indicates enrichment of GC content in naturally occurring sequences while negative Z-score indicates the reverse.

**Figure S38 Z—score profile for GC content of *D. melanogaster***

This figure shows the extent of GC content between real and random models for the first 150 and last 150 amino acid positions of D. *melanogaster* proteins. Figure representation is same as Figure S37.

**Figure S39 Z—score profile GC content of *C. elegans***

This figure shows the extent of GC content between real and random models for the first 150 and last 150 amino acid positions of *C. elegans* proteins. Figure representation is same as Figure S37.

**Figure S40 Z—score profile GC content of *S. cerevisiae***

This figure shows the extent of GC content between real and random models for the first 150 and last 150 amino acid positions of *S. cerevisiae* proteins. Figure representation is same as Figure S37.

**Figure S41 Z—score profile for GC content of *A. oryzae***

This figure shows the extent of GC content between real and random models for the first 150 and last 150 amino acid positions of *A. oryzae* proteins. Figure representation is same as Figure S37.

**Figure S42 Z—score profile for GC content of *N. crassa***

This figure shows the extent of GC content between real and random models for the first 150 and last150 amino acid positions of *N. crassa* proteins. Figure representation is same as Figure S37.

**Figure S43 Z—score profile for GC content of *B. subtilis***

This figure shows the extent of GC content between real and random models for the first 150 and last 150 amino acid positions of *B. subtilis* proteins. Figure representation is same as Figure S37.

**Figure S44 Z—score profile for GC content of *E. coli***

This figure shows the extent of GC content between real and random models for the first 150 and last 150 amino acid positions of *E. coli* proteins. Figure representation is same as Figure S37.

**Figure S45 Z—score profile for GC content of *D. radiodurans*.**

This figure shows the extent of GC content between real and random models for the first 150 and last 150 amino acid positions of *D. radiodurans* proteins. Figure representation is same as Figure S37.

**Figure S46 Z—score profile for GC content of *M. mazei***

This figure shows the extent of GC content between real and random models for the first 150 and last 150 amino acid positions of *M. mazei* proteins. Figure representation is same as Figure S37.

**Figure S47 Z—score profile for position-specific GC score of *H. volcanii***

This figure shows the extent of GC content between real and random models for the first 150 and last 150 amino acid positions of *H. volcanii* proteins. Figure representation is same as Figure S37.

**Figure S48 Z—score profile for GC content of *T. gammatolerans***

This figure shows the extent of GC content between real and random models for the first 150 and last 150 amino acid positions of *T. gammatolerans* proteins. Figure representation is same as Figure S37.

**Figure S49 Z—score profile for the position-specific disorder score of *A. oryzae* considering proteins without any splice junction in the coding sequences up to the first and last 100 amino acids positions**

This figure shows the extent of protein disorder in the proteins without any splice junction in the coding sequences up to the first and last 100 amino acids positions and the corresponding random sequences specifically generated from these sequences. Disordered residues in these real and random protein datasets were predicted by consensus approach #1 (see main text for details). In each panel, vertical blue lines stand for the average position-specific disorder scores of real proteins and grey lines that for random proteins. At each position, the statistical difference between these two measurements is accessed through Z-score (see main text) and is shown with blue line.). In panels **A** and **B,** Z-scores were calculated comparing position-specific disorder scores of real and terminal residue conserved random sequences of each species and in panels **C** and **D,** Z-scores were calculated comparing position-specific disorder scores of real and column random sequences of each species. Here positive Z-score indicates enrichment of protein disorder in naturally occurring sequences while negative Z-score indicates the reverse. Corresponding *P*-value at each position is shown either with a green (*P* < 0.05) or red dot (*P* >= 0.05) above the Z-score.

**Figure S50 Z—score profile for the position-specific disorder score of *C. elegans* considering proteins without any splice junction in the coding sequences up to the first and last 100 amino acids positions**

This figure shows the extent of protein disorder in the proteins without any splice junction in the coding sequences up to the first and last 100 amino acids positions and the corresponding random sequences specifically generated from these sequences. Disordered residues in these real and random protein datasets were predicted by consensus approach #1 (see main text for details). In each panel, vertical blue lines stand for the average position-specific disorder scores of real proteins and grey lines that for random proteins. At each position, the statistical difference between these two measurements is accessed through Z-score (see main text) and is shown with blue line.). In panels **A** and **B,** Z-scores were calculated comparing position-specific disorder scores of real and terminal residue conserved random sequences of each species and in panels **C** and **D,** Z-scores were calculated comparing position-specific disorder scores of real and column random sequences of each species. Here positive Z-score indicates enrichment of protein disorder in naturally occurring sequences while negative Z-score indicates the reverse. Corresponding *P*-value at each position is shown either with a green (*P* < 0.05) or red dot (*P* >= 0.05) above the Z-score.

**Figure S51 Z—score profile for the position-specific disorder score of *D. melanogaster* considering proteins without any splice junction in the coding sequences up to the first and last 100 amino acids positions** panels **A** and **B,** Z-scores were calculated comparing position-specific disorder scores of real and terminal residue conserved random sequences of each species and in panels **C** and **D,** Z-scores were calculated comparing position-specific disorder scores of real and column random sequences of each species. Here positive Z-score indicates enrichment of protein disorder in naturally occurring sequences while negative Z-score indicates the reverse. Corresponding *P*-value at each position is shown either with a green (*P* < 0.05) or red dot (*P* >= 0.05) above the Z-score.

**Figure S52 Z—score profile for the position-specific disorder score of *H. sapiens* considering proteins without any splice junction in the coding sequences up to the first and last 100 amino acids positions**

This figure shows the extent of protein disorder in the proteins without any splice junction in the coding sequences up to the first and last 100 amino acids positions and the corresponding random sequences specifically generated from these sequences. Disordered residues in these real and random protein datasets were predicted by consensus approach #1 (see main text for details). In each panel, vertical blue lines stand for the average position-specific disorder scores of real proteins and grey lines that for random proteins. At each position, the statistical difference between these two measurements is accessed through Z-score (see main text) and is shown with blue line.). In panels **A** and **B,** Z-scores were calculated comparing position-specific disorder scores of real and terminal residue conserved random sequences of each species and in panels **C** and **D,** Z-scores were calculated comparing position-specific disorder scores of real and column random sequences of each species. Here positive Z-score indicates enrichment of protein disorder in naturally occurring sequences while negative Z-score indicates the reverse. Corresponding *P*-value at each position is shown either with a green (*P* < 0.05) or red dot (*P* >= 0.05) above the Z-score.

**Figure S53 Z—score profile for the position-specific disorder score of *N. crassa* considering proteins without any splice junction in the coding sequences up to the first and last 100 amino acids positions** panels **A** and **B,** Z-scores were calculated comparing position-specific disorder scores of real and terminal residue conserved random sequences of each species and in panels **C** and **D,** Z-scores were calculated comparing position-specific disorder scores of real and column random sequences of each species. Here positive Z-score indicates enrichment of protein disorder in naturally occurring sequences while negative Z-score indicates the reverse. Corresponding *P*-value at each position is shown either with a green (*P* < 0.05) or red dot (*P* >= 0.05) above the Z-score.

**Figure S54 Z—score profile for the position-specific disorder score of *S. cerevisiae* considering proteins without any splice junction in the coding sequences up to the first and last 100 amino acids positions**

This figure shows the extent of protein disorder in the proteins without any splice junction in the coding sequences up to the first and last 100 amino acids positions and the corresponding random sequences specifically generated from these sequences. Disordered residues in these real and random protein datasets were predicted by consensus approach #1 (see main text for details). In each panel, vertical blue lines stand for the average position-specific disorder scores of real proteins and grey lines that for random proteins. At each position, the statistical difference between these two measurements is accessed through Z-score (see main text) and is shown with blue line.). In panels **A** and **B,** Z-scores were calculated comparing position-specific disorder scores of real and terminal residue conserved random sequences of each species and in panels **C** and **D,** Z-scores were calculated comparing position-specific disorder scores of real and column random sequences of each species. Here positive Z-score indicates enrichment of protein disorder in naturally occurring sequences while negative Z-score indicates the reverse. Corresponding *P*-value at each position is shown either with a green (*P* < 0.05) or red dot (*P* >= 0.05) above the Z-score.

**Figure S55 Z—score profile for the predicted solvent accessibility for the 12 species considered in this study**

This figure shows the Z-score profile for the predicted solvent accessibility of real and randomly generated protein sequences for the first 150 and last 150 positions of 12 organisms considered in this study. In panels A and B, Z-scores were calculated comparing position-specific disorder scores of real and terminal residue conserved random sequences of each species and in panels C and D, Zscores were calculated comparing position-specific disorder scores of real and column random sequences of each species. The color scale for each graph is shown on the top of each panel. Here positive Z-score indicates enrichment of predicted solvent accessibility in naturally occurring sequences while negative Z-score indicates the reverse. Organisms are arranged according to theirmid-point rooted species tree retrieved from NCBI taxonomic database with the help of their species taxonomic identifier.

**Figure S56 Z—score profile for predicted solvent accessibility of *H. sapiens***

This figure shows the extent of predicted solvent accessibility between the real and random proteins of *H. sapiens* for the first 150 and last 150 positions. **A, B.** Comparison of real and terminal residue conserved random sequences. **C, D.** Comparison of real and column random sequences. In each panel, blue line stand for the average predicted solvent accessibility scores of real proteins and black line that for random proteins. At each position, the statistical difference between these two measurements is accessed through Z-score (see main text) and is shown with blue line. Corresponding *P*-value at each position is shown either with a green (*P* < 0.05) or red dot (*P* >= 0.05) above the Z-score. Here positive Z-score indicates enrichment of predicted solvent accessibility in naturally occurring sequences while negative Z-score indicates the reverse.

**Figure S57 Z—score profile for predicted solvent accessibility of *D. melanogaster***

This figure shows the extent of predicted solvent accessibility between real and random proteins of *D. melanogaster* proteins for the first 150 and last 150 positions. Figure representation is same as Figure S56.

**Figure S58 Z—score profile predicted solvent accessibility of *C. elegans***

This figure shows the extent of predicted solvent accessibility between real and random proteins of *C. elegans* proteins for the first 150 and last 150 positions. Figure representation is same as Figure S56.

**Figure S59 Z—score profile solvent accessibility of *S. cerevisiae***

This figure shows the extent of predicted solvent accessibility between real and random proteins of *S. cerevisiae* proteins for the first 150 and last 150 positions. Figure representation is same as Figure S56.

**Figure S60 Z—score profile for predicted solvent accessibility of *A. oryzae***

This figure shows the extent of predicted solvent accessibility between real and random proteins of *A. oryzae* proteins for the first 150 and last 150 positions. Figure representation is same as Figure S56.

**Figure S61 Z—score profile for position-specific predicted solvent accessibility of *N. crassa***

This figure shows the extent of predicted solvent accessibility between real and random proteins of *N. crassa* proteins for the first 150 and last 150 positions. Figure representation is same as Figure S56.

**Figure S62 Z—score profile for position-specific predicted solvent accessibility of *B. subtilis***

This figure shows the extent of predicted solvent accessibility between real and random proteins of *B. subtilis* proteins for the first 150 and last 150 positions. Figure representation is same as Figure S56.

**Figure S63 Z—score profile for position-specific predicted solvent accessibility of *E. coli***

This figure shows the extent of predicted solvent accessibility between real and random proteins of *E. coli* proteins for the first 150 and last 150 positions. Figure representation is same as Figure S56.

**Figure S64 Z—score profile for position-specific predicted solvent accessibility of *D. radiodurans*.**

This figure shows the extent of predicted solvent accessibility between real and random proteins of *D. radiodurans* proteins for the first 150 and last 150 positions. Figure representation is same as Figure S56.

**Figure S65 Z—score profile for position-specific predicted solvent accessibility of *M. mazei***

This figure shows the extent of predicted solvent accessibility between real and random proteins of *M. mazei* proteins for the first 150 and last 150 positions. Figure representation is same as Figure S56.

**Figure S66 Z—score profile for position-specific predicted solvent accessibility of *H. volcanii***

This figure shows the extent of predicted solvent accessibility between real and random proteins of *H. volcanii* proteins for the first 150 and last 150 positions. Figure representation is same as Figure S56.

**Figure S67 Z—score profile for position-specific predicted solvent accessibility of *T. gammatolerans***

This figure shows the extent of predicted solvent accessibility between real and random proteins of *T. gammatolerans* proteins for the first 150 and last 150 positions. Figure representation is same as Figure S56.

**Figure S68 Mean Z-score profile for the last 150 positions of *H. sapiens* proteins under GO slim biological process ontologies in reference to terminal residue conserved random**

This heat map represents position-wise Z-score of predicted disordered residues for the last 150 positions near the C-terminal region of human proteins under each GO slim biological process ontology. For each such GO-slim category, Z-scores for predicted disordered residues were calculated considering proteins (more than 200 residues in length) under that category in reference to their corresponding terminal residue conserved random sequences. Here rows represent the positions along the protein sequence. Z-scores were coded in color scheme (color legend). Only the ontologies with more than 100 proteins were shown.

**Figure S69 Mean Z-score profile for the first 150 positions of *H. sapiens* proteins under GO slim biological process ontologies in reference to column random model**

This heat map represents position-wise Z-score of predicted disordered residues for the first 150 positions near the N-terminal region of human proteins under each GO slim biological process ontology. In each functional category, Z-scores for predicted disordered residues were calculated considering proteins (more than 200 residues in length) under that category in reference to their corresponding column random sequences. Here rows represent the positions along the protein sequence and columns represents Z-score coded in color scheme (color legend). Only the ontologies with more than 100 proteins were shown.

**Figure S70 Mean Z-score profile for the last 150 positions of *H. sapiens* proteins under GO slim biological process ontologies in reference to column random model**

This heat map represents position-wise Z-score of predicted disordered residues for the last 150 positions near the C-terminal region of *H. sapiens* proteins under each GO slim biological process ontology. In each functional category, Z-scores for predicted disordered residues were calculated considering proteins (more than 200 residues in length) under that category in reference to their corresponding column random sequences. Here rows represent the positions along the protein sequence and columns represents Z-score coded in color scheme (color legend). Only the ontologies with more than 100 proteins were shown.

**Figure S71 Mean Z-score profile for the first 150 positions of *H. sapiens* proteins under GO slim molecular function ontologies in reference to terminal residue conserved random model**

This heat map represents position-wise Z-score of predicted disordered residues for the first 150 positions near the N-terminal region of *H. sapiens* proteins under each GO slim molecular function ontology. In each functional category, Z-scores for predicted disordered residues were calculated considering proteins (more than 200 residues in length) under that category in reference to their corresponding terminal residue conserved random sequences. Here rows represent the positions along the protein sequence and columns represents Z-score coded in color scheme (color legend). Only the ontologies with more than 100 proteins were shown.

**Figure S72 Mean Z-score profile for the last 150 positions of *H. sapiens* proteins under GO slim molecular function ontologies in reference to terminal residue conserved random model**

This heat map represents position-wise Z-score of predicted disordered residues for the last 150 positions near the C-terminal region of *H. sapiens* proteins under each GO slim molecular function ontology. In each functional category, Z-scores for predicted disordered residues were calculated considering proteins (more than 200 residues in length) under that category in reference their corresponding terminal residue conserved random sequences. Here rows represent the positions along the protein sequence and columns represents Z-score coded in color scheme (color legend). Only the ontologies with more than 100 proteins were shown.

**Figure S73 Mean Z-score profile for the first 150 positions of *H. sapiens* proteins under GO slim molecular function ontologies in reference to column random model**

This heat map represents position-wise Z-score of predicted disordered residues for the first 150 positions near the N-terminal region of *H. sapiens* proteins under each GO slim molecular function ontology. In each functional category, Z-scores for predicted disordered residues were calculated considering proteins (more than 200 residues in length) under that category in reference to their corresponding column random sequences. Here rows represent the positions along the protein sequence and columns represents Z-score coded in color scheme (color legend). Only the ontologies with more than 100 proteins were shown.

**Figure S74 Mean Z-score profile for the last 150 positions of *H. sapiens* proteins under GO slim molecular function ontologies in reference to column random model**

This heat map represents position-wise Z-score of predicted disordered residues for the last 150 positions near the C-terminal region of *H. sapiens* proteins under each GO slim molecular function ontology. In each functional category, Z-scores for predicted disordered residues were calculated considering proteins (more than 200 residues in length) under that category in reference to their corresponding column random sequences. Here rows represent the positions along the protein sequence and columns represents Z-score coded in color scheme (color legend). Only the ontologies with more than 100 proteins were shown.

**Figure S75 Mean Z-score profile for the first 150 positions of *H. sapiens* proteins under their GO slim cellular component ontologies in reference to terminal residue conserved random model**

This heat map represents position-wise Z-score of predicted disordered residues for the first 150 positions near the N-terminal region of *H. sapiens* proteins under each GO slim cellular component ontology. In each functional category, Z-scores for predicted disordered residues were calculated considering proteins (more than 200 residues in length) under that category in reference to their corresponding terminal residue random sequences. Here rows represent the positions along the protein sequence and columns represents Z-score coded in color scheme (color legend). Only the ontologies with more than 100 proteins were shown.

**Figure S76 Mean Z-score profile for the last 150 positions of *H. sapiens* proteins under GO slim cellular component ontologies in reference to terminal residue conserved random model**

This heat map represents position-wise Z-score of predicted disordered residues for the last 150 positions near the C-terminal region of *H. sapiens* proteins under each GO slim cellular component ontology. In each functional category, Z-scores for predicted disordered residues were calculated considering proteins (more than 200 residues in length) under that category in reference to their corresponding terminal residue conserved random sequences. Here rows represent the positions along the protein sequence and columns represents Z-score coded in color scheme (color legend). Only the ontologies with more than 100 proteins were shown.

**Figure S77 Mean Z-score profile for the first 150 positions of *H. sapiens* proteins under GO slim cellular component ontologies in reference to column random model**

This heat map represents position-wise Z-score of predicted disordered residues for the first 150 positions near the N-terminal region of *H. sapiens* proteins under each GO slim cellular component ontology. In each functional category, Z-scores for predicted disordered residues were calculated considering proteins (more than 200 residues in length) under that category in reference to their corresponding column random sequences. Here rows represent the positions along the protein sequence and columns represents Z-score coded in color scheme (color legend). Only the ontologies with more than 100 proteins were shown.

**Figure S78 Mean Z-score profile for the last 150 positions of *H. sapiens* proteins under GO slim cellular component ontologies in reference to column random model**

This heat map represents position-wise Z-score of predicted disordered residues for the last 150 positions near the C-terminal region of *H. sapiens* proteins under each GO slim cellular component ontology. In each functional category, Z-scores for predicted disordered residues were calculated considering proteins (more than 200 residues in length) under that category in reference to their corresponding column random sequences. Here rows represent the positions along the protein sequence and columns represents Z-score coded in color scheme (color legend). Only the ontologies with more than 100 proteins were shown.

**Figure S79 Comparison of Z-score profile between highly and lowly expressed genes-terminal residue conserved random model**

This figure shows the position-wise (first and last 150 positions) Z-scores for predicted disordered residues of proteins encoded by highly and lowly expressed genes of three organisms *H. sapiens, E. coli,* and *S. cerevisiae*. For each group of proteins, at each position, we first calculated Z-scores for predicted disordered residues with respect to their terminal residues conserved random sequences then compared Z-scores between highly and lowly expressed groups. Here blue line stands for Z-scores of highly expressed proteins and red line stands for Z-scores of lowly expressed proteins. To access statistical significance of the difference we compared Z-scores between highly and lowly expressed genes considering sliding window of 21 residues starting from the first positions towards the end position with an interval of 1 position. Z-scores were compared window-wise and *P*-values were calculated for each such window. Here we plotted the mean Z-score (and associated *P*-values) of highly and lowly expressed group in each window positioned at the center of the window. Statistical significance was accessed through Mann-Whitney U test following non-parametric distribution. *P* –values were shown in color codes at the top of Z-scores.

**Figure S80 Comparison of Z-score profile between highly and lowly expressed genes-column random model**

This figure shows the position-wise (first and last 150 positions) Z-scores for predicted disordered residues of proteins encoded by highly and lowly expressed genes of three organisms *H. sapiens*, *E. coli,* and *S. cerevisiae*. For each group of proteins, at each position, we first calculated Z-scores for predicted residues with respect to their column random sequences then compared Z-scores between highly and lowly expressed groups. Here blue line stands for Z-scores of highly expressed proteins and red line stands for Z-scores of lowly expressed proteins. To access statistical significance of the difference we compared Z-scores between highly and lowly expressed genes considering sliding window of 21 residues starting from the first positions towards the end position with an interval of 1 position. Z-scores were compared window-wise and *P*-values were calculated for each such window. Here we plotted the mean Z-score (and associated *P*-values) of highly and lowly expressed group in each window positioned at the center of the window. Statistical significance was accessed through Mann-Whitney U test following non-parametric distribution. *P* –values were shown in color codes at the top of Z-scores.

### Supplementary tables legends

**Table S1 List of genomes used in the analysis**

**Table S2 Table of correlation between the results of different disorder prediction algorithms and experimental annotation**

