## Supplemental figures and tables for "Exploring Potential Signals of Selection for Disordered Residues in Naturally Occurring Prokaryotic and Eukaryotic Proteins"

**Running title:** *Panda A and Tuller T/ Evidences Of Selection In Disordered Proteins*

**Figure S1**

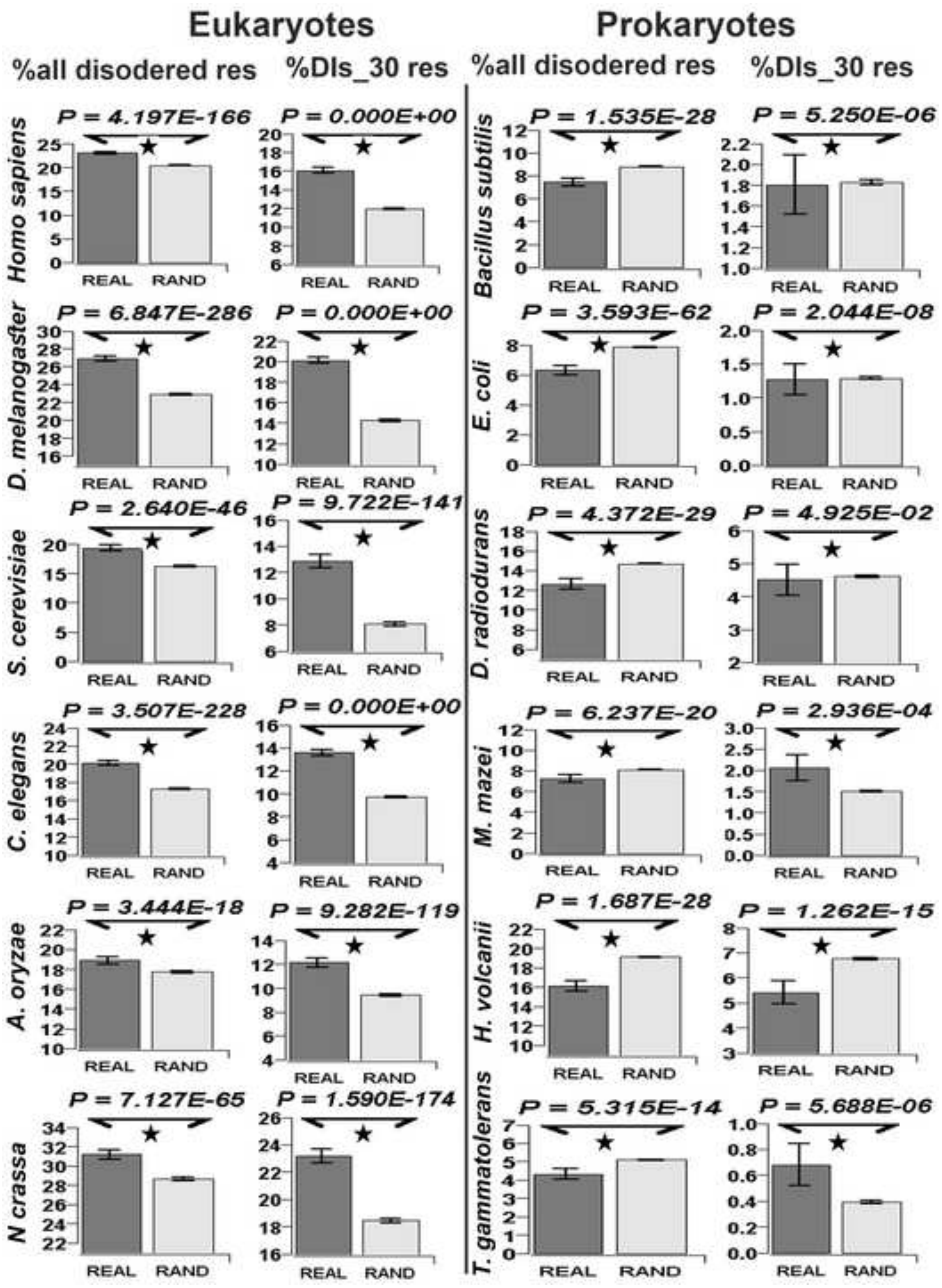

Eukaryotes
Prokaryotes
%all disodered res
%DIs_30 res
%all disodered res
%DIs_30 res
Homo sapiens
P = 4.197E-166
P = 0.000E+00
D. melanogaster
P = 6.847E-286
P = 0.000E+00
S. cerevisiae
P = 2.640E-46
P = 9.722E-141
C. elegans
P = 3.507E-228
P = 0.000E+00
A. oryzae
P = 3.444E-18
P = 9.282E-119
N crassa
P = 7.127E-65
P = 1.590E-174
Bacillus subtilis
P = 1.535E-28
P = 5.250E-06
E. coli
P = 3.593E-62
P = 2.044E-08
D. radiodurans
P = 4.372E-29
P = 4.925E-02
M. mazei
P = 6.237E-20
P = 2.936E-04
H. volcanii
P = 1.687E-28
P = 1.262E-15
T. gammatolerans
P = 5.315E-14
P = 5.688E-06
REAL
RAND

**Figure S2**

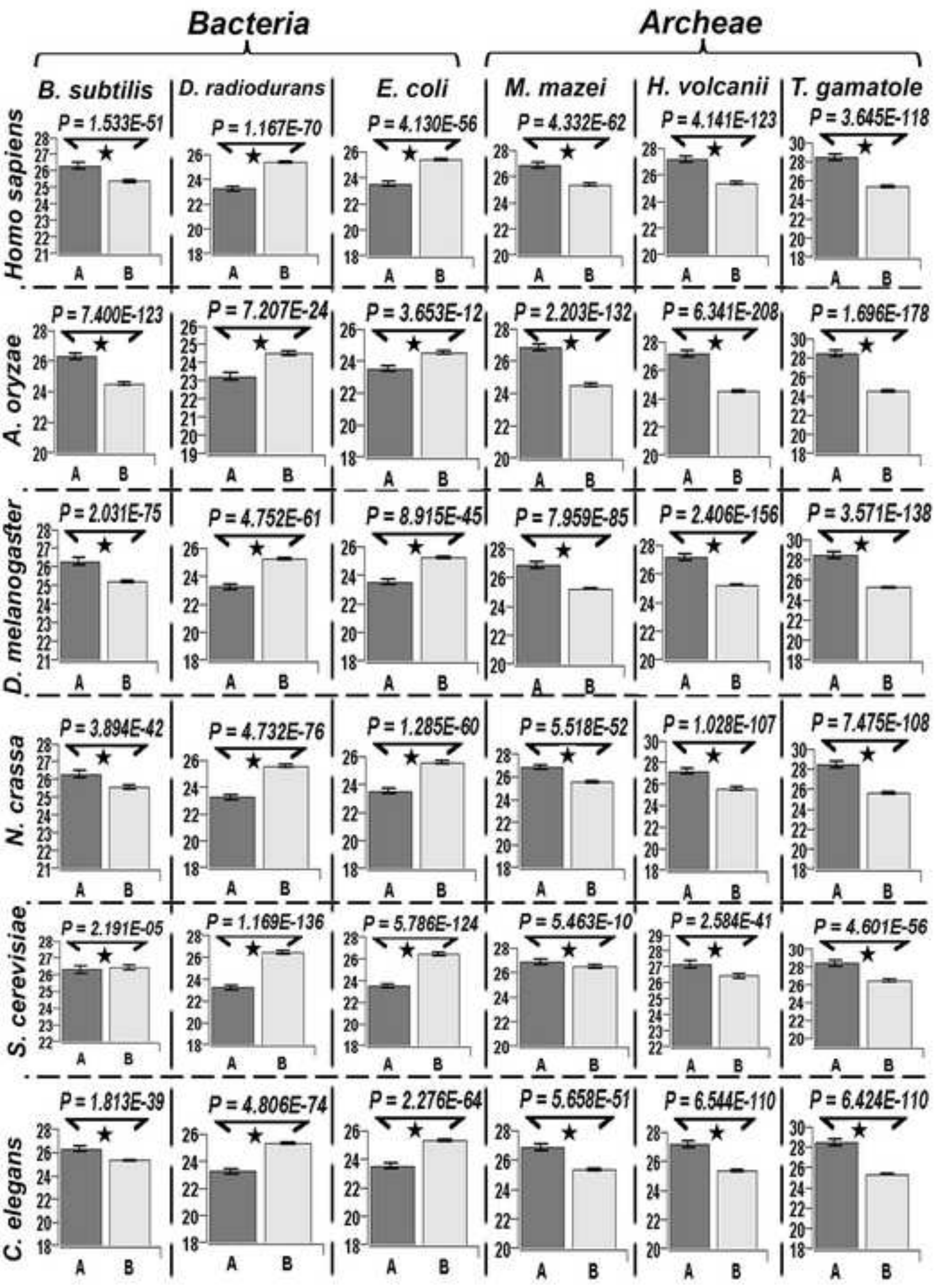

Bacteria
Archeae
B. subtilis
D. radiodurans
E. coli
M. mazei
H. volcanii
T. gamatole
Homo sapiens
A. oryzae
D. melanogaster
N. crassa
S. cerevisiae
C. elegans
P = 1.533E-51
P = 1.167E-70
P = 4.130E-56
P = 4.332E-62
P = 4.141E-123
P = 3.645E-118
P = 7.400E-123
P = 7.207E-24
P = 3.653E-12
P = 2.203E-132
P = 6.341E-208
P = 1.696E-178
P = 2.031E-75
P = 4.752E-61
P = 8.915E-45
P = 7.959E-85
P = 2.406E-156
P = 3.571E-138
P = 3.894E-42
P = 4.732E-76
P = 1.285E-60
P = 5.518E-52
P = 1.028E-107
P = 7.475E-108
P = 2.191E-05
P = 1.169E-136
P = 5.786E-124
P = 5.463E-10
P = 2.584E-41
P = 4.601E-56
P = 1.813E-39
P = 4.806E-74
P = 2.276E-64
P = 5.658E-51
P = 6.544E-110
P = 6.424E-110
A
B

**Figure S3**

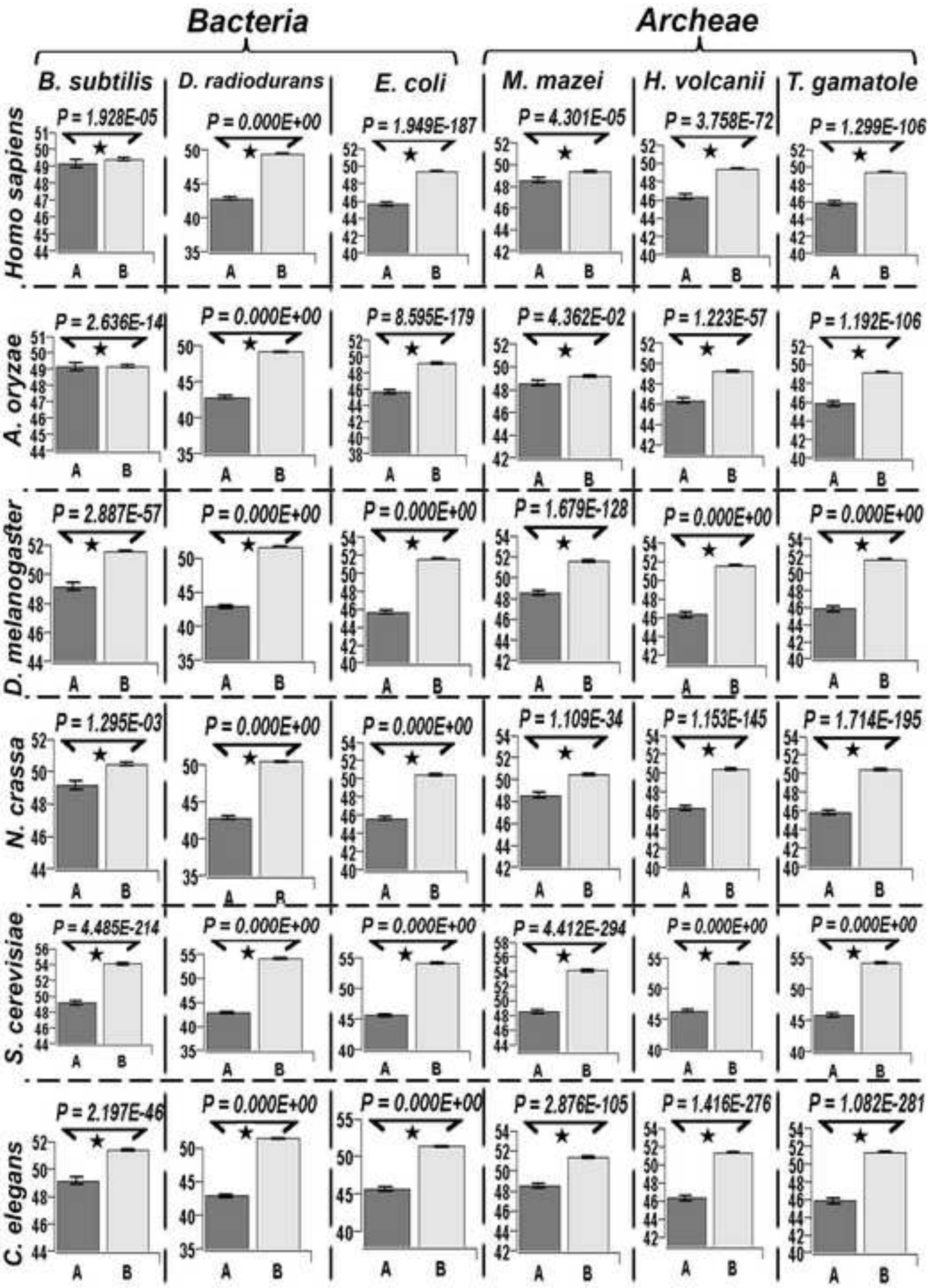
Bacteria
Archeae
B. subtilis
D. radiodurans
E. coli
M. mazei
H. volcanii
T. gamatole
Homo sapiens
A. oryzae
D. melanogaster
N. crassa
S. cerevisiae
C. elegans
P = 1.928E-05
P = 0.000E+00
P = 1.949E-187
P = 4.301E-05
P = 3.758E-72
P = 1.299E-106
P = 2.636E-14
P = 0.000E+00
P = 8.595E-179
P = 4.362E-02
P = 1.223E-57
P = 1.192E-106
P = 2.887E-57
P = 0.000E+00
P = 0.000E+00
P = 1.679E-128
P = 0.000E+00
P = 0.000E+00
P = 1.295E-03
P = 0.000E+00
P = 0.000E+00
P = 1.109E-34
P = 1.153E-145
P = 1.714E-195
P = 4.485E-214
P = 0.000E+00
P = 0.000E+00
P = 4.412E-294
P = 0.000E+00
P = 0.000E+00
P = 2.197E-46
P = 0.000E+00
P = 0.000E+00
P = 2.876E-105
P = 1.416E-276
P = 1.082E-281
A
B

Figure S4

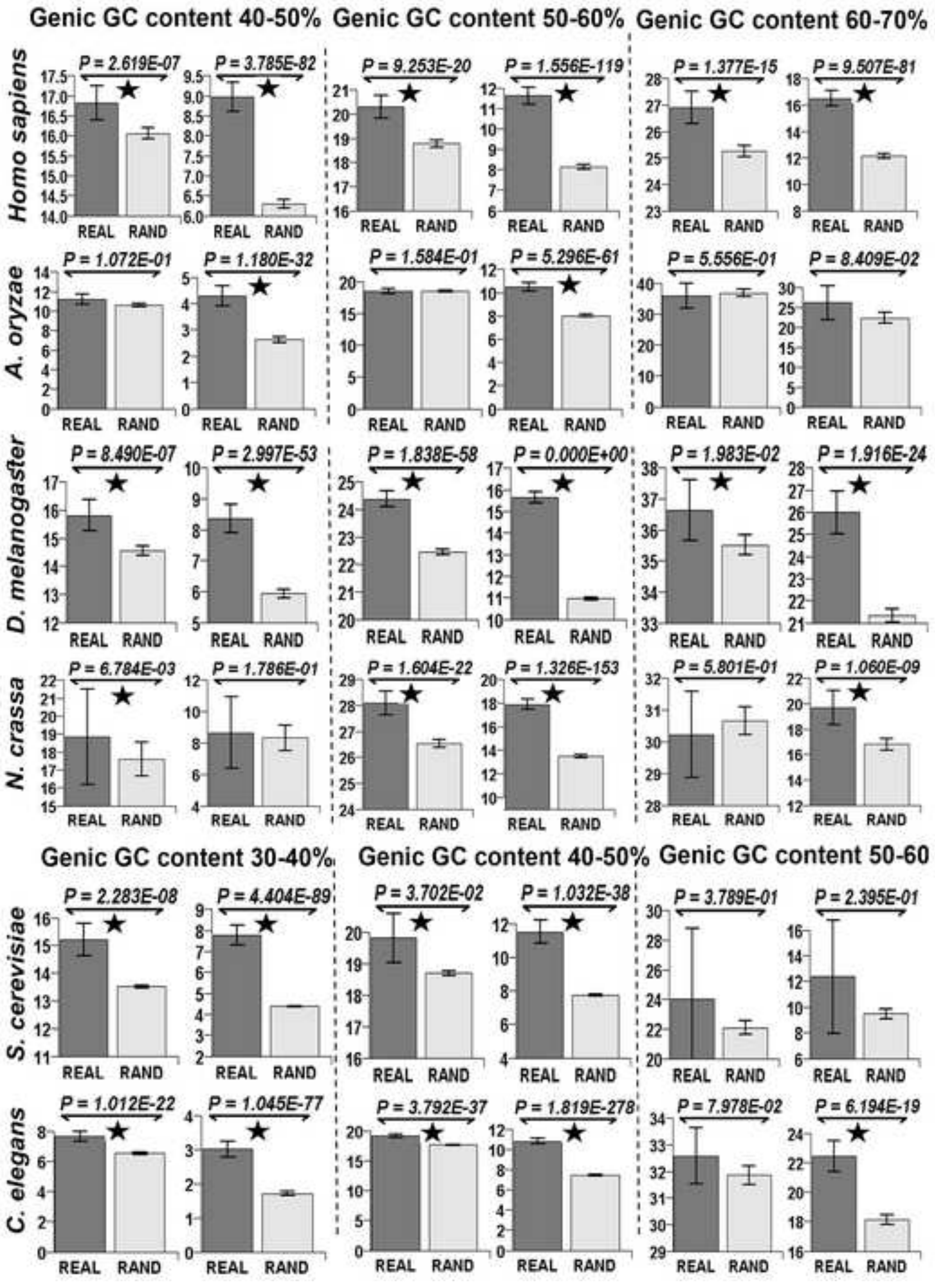

Genic GC content 40-50%
Genic GC content 50-60%
Genic GC content 60-70%
Homo sapiens
P = 2.619E-07
P = 3.785E-82
P = 9.253E-20
P = 1.556E-119
P = 1.377E-15
P = 9.507E-81
A. oryzae
P = 1.072E-01
P = 1.180E-32
P = 1.584E-01
P = 5.296E-61
P = 5.556E-01
P = 8.409E-02
D. melanogaster
P = 8.490E-07
P = 2.997E-53
P = 1.838E-58
P = 0.000E+00
P = 1.983E-02
P = 1.916E-24
N. crassa
P = 6.784E-03
P = 1.786E-01
P = 1.604E-22
P = 1.326E-153
P = 5.801E-01
P = 1.060E-09
Genic GC content 30-40%
Genic GC content 40-50%
Genic GC content 50-60
S. cerevisiae
P = 2.283E-08
P = 4.404E-89
P = 3.702E-02
P = 1.032E-38
P = 3.789E-01
P = 2.395E-01
C. elegans
P = 1.012E-22
P = 1.045E-77
P = 3.792E-37
P = 1.819E-278
P = 7.978E-02
P = 6.194E-19
REAL
RAND

Figure S5

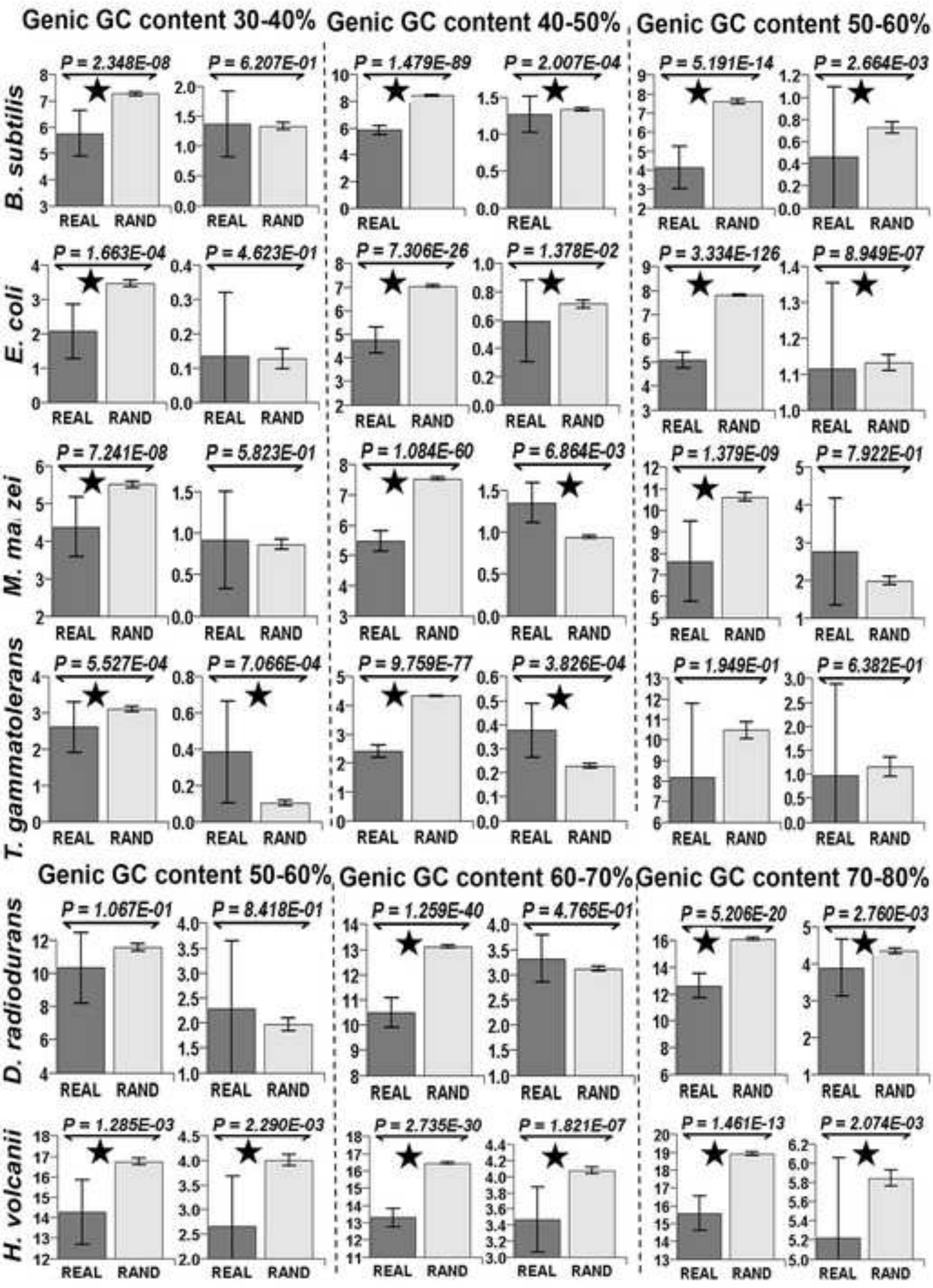

Genic GC content 30-40%
Genic GC content 40-50%
Genic GC content 50-60%
B. subtilis
P = 2.348E-08
P = 6.207E-01
P = 1.479E-89
P = 2.007E-04
P = 5.191E-14
P = 2.664E-03
E. coli
P = 1.663E-04
P = 4.623E-01
P = 7.306E-26
P = 1.378E-02
P = 3.334E-126
P = 8.949E-07
M. mazei
P = 7.241E-08
P = 5.823E-01
P = 1.084E-60
P = 6.864E-03
P = 1.379E-09
P = 7.922E-01
T. gammatolerans
P = 5.527E-04
P = 7.066E-04
P = 9.759E-77
P = 3.826E-04
P = 1.949E-01
P = 6.382E-01
Genic GC content 50-60%
Genic GC content 60-70%
Genic GC content 70-80%
D. radiodurans
P = 1.067E-01
P = 8.418E-01
P = 1.259E-40
P = 4.765E-01
P = 5.206E-20
P = 2.760E-03
H. volcanii
P = 1.285E-03
P = 2.290E-03
P = 2.735E-30
P = 1.821E-07
P = 1.461E-13
P = 2.074E-03
REAL
RAND

**Figure S6**

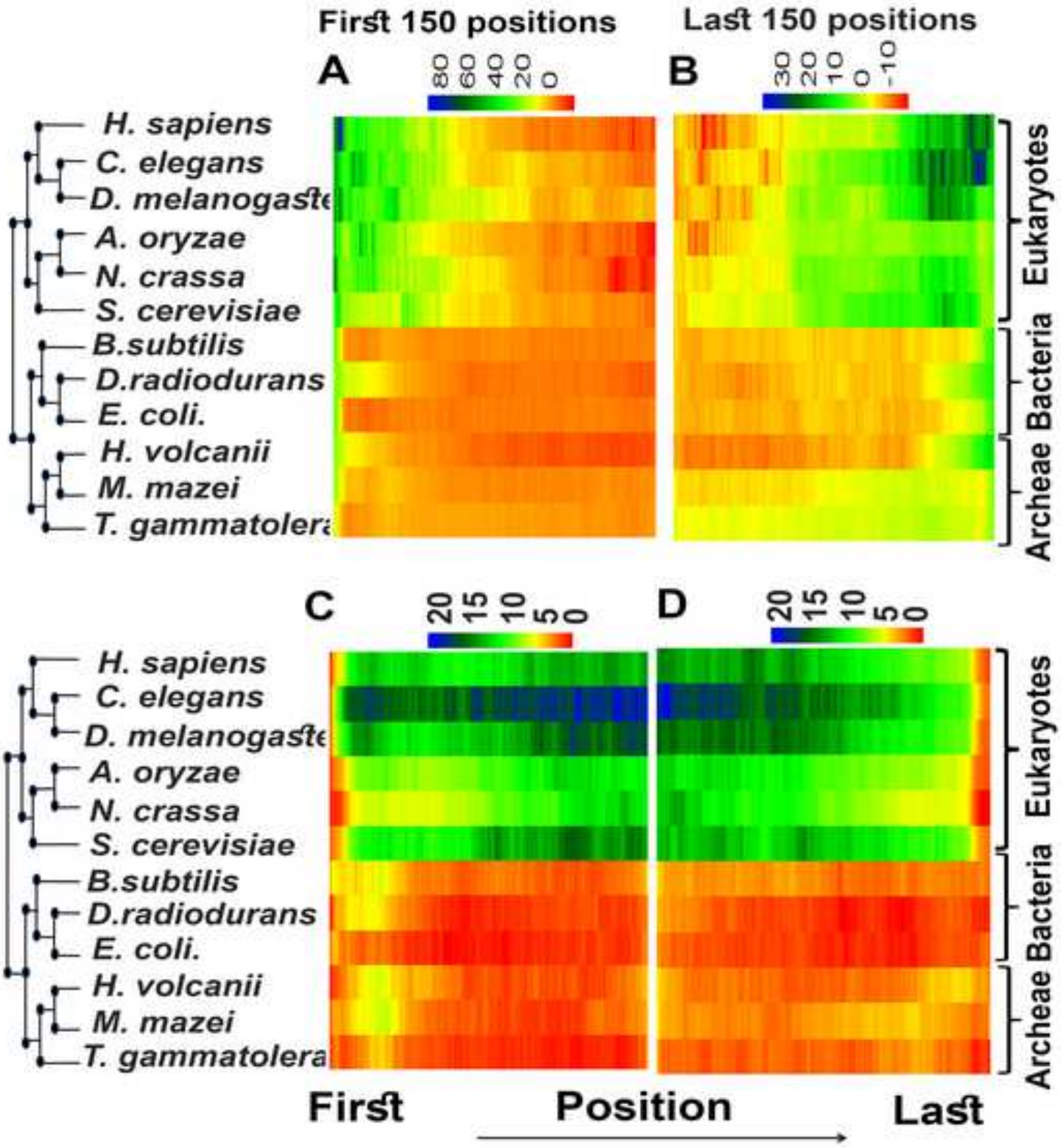

First 150 positions
Last 150 positions
A
80 60 40 20 0
B
30 20 10 0 -10
H. sapiens
C. elegans
D. melanogaste
A. oryzae
N. crassa
S. cerevisiae
B.subtilis
D.radiodurans
E. coli.
H. volcanii
M. mazei
T. gammatolera
Eukaryotes
Bacteria
Archeae
C
20 15 10 5 0
D
20 15 10 5 0
First
Position
Last

Figure S7

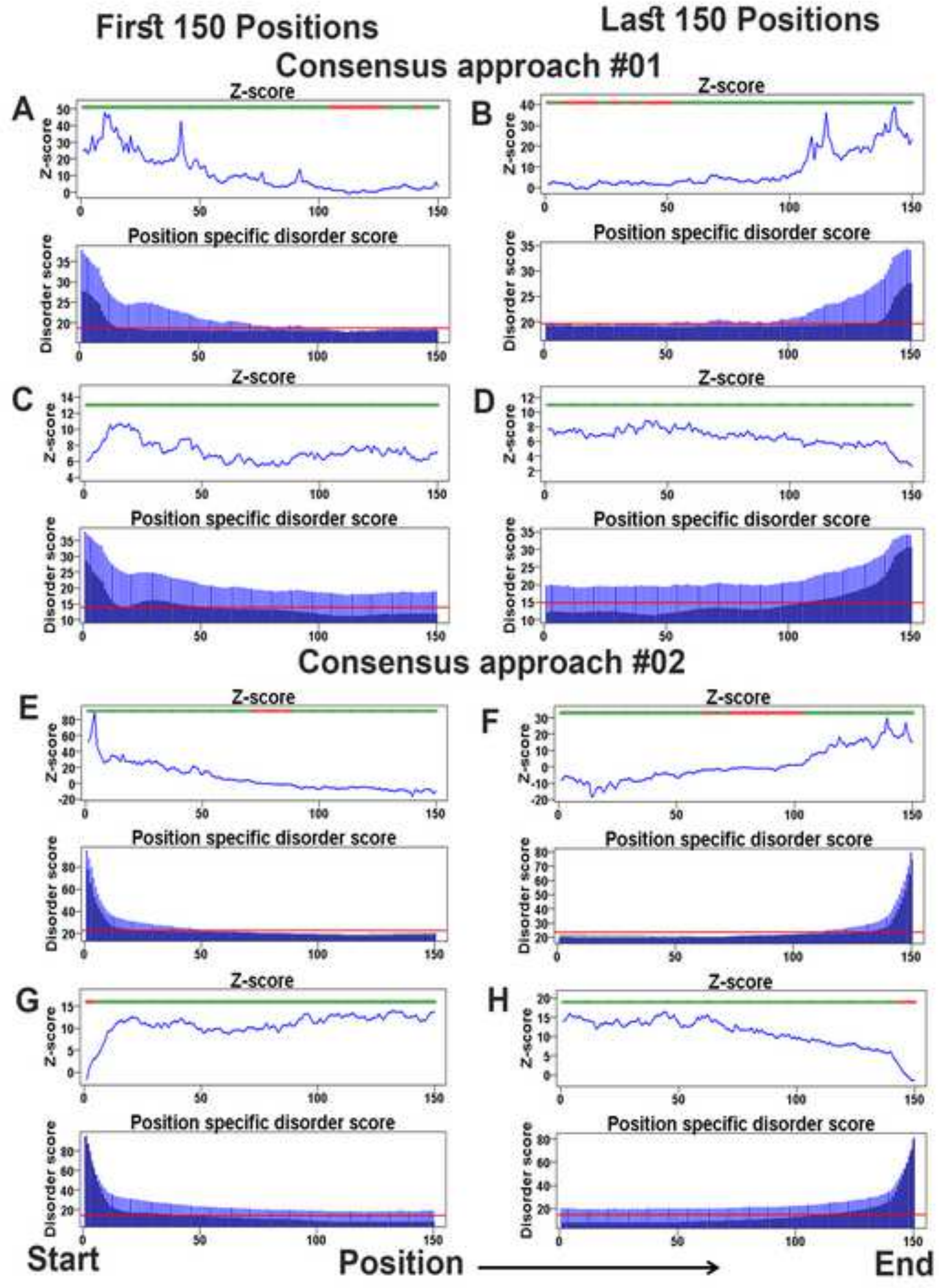

First 150 Positions
Last 150 Positions
Consensus approach #01
A
B
C
D
Z-score
Position specific disorder score
Disorder score
Consensus approach #02
E
F
G
H
Start
Position
End

Figure S8

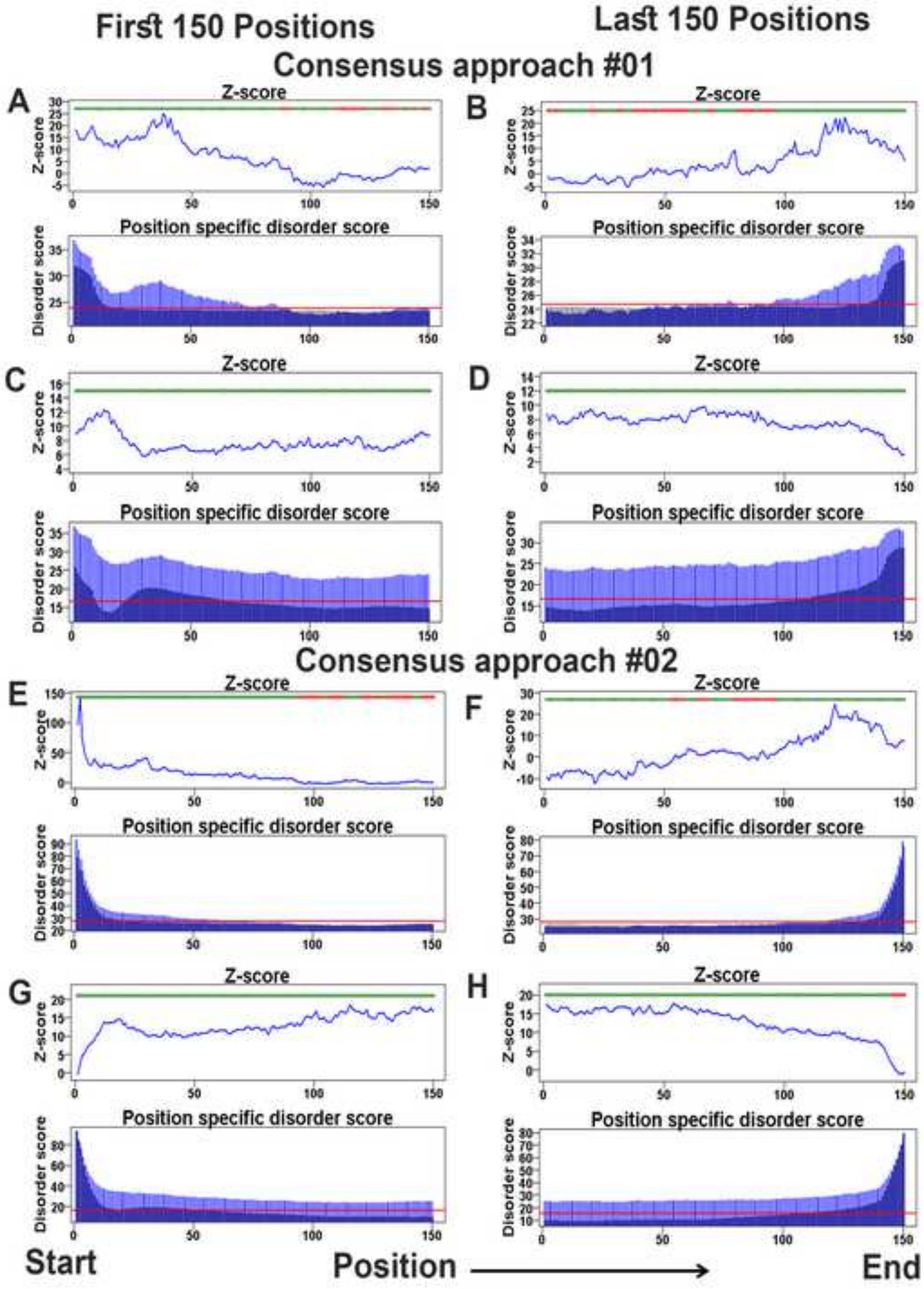

First 150 Positions
Last 150 Positions
Consensus approach #01
A
B
C
D
Z-score
Position specific disorder score
Disorder score
Consensus approach #02
E
F
G
H
Start
Position
End

**Figure S9**

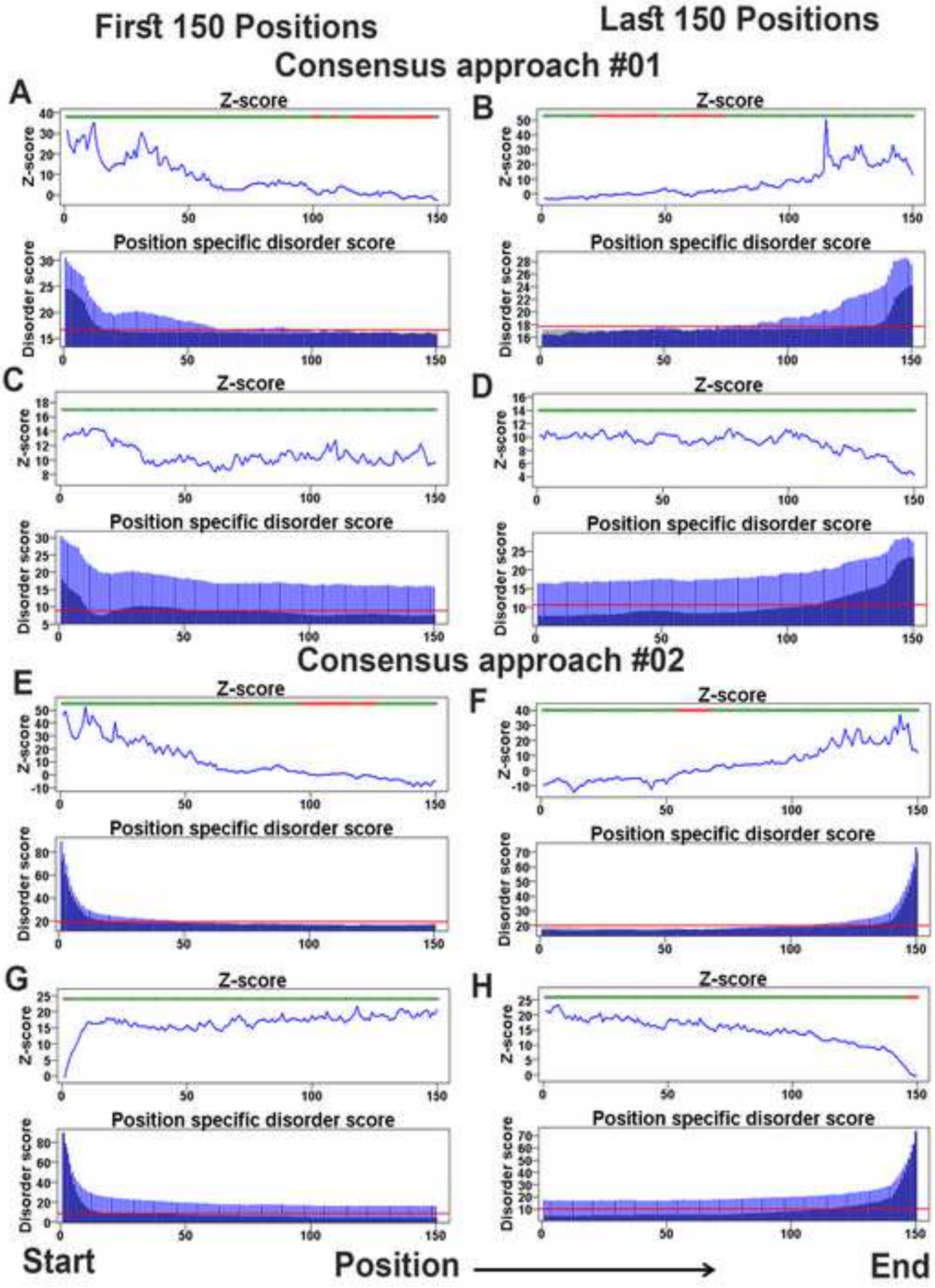

First 150 Positions
Last 150 Positions
Consensus approach #01
A
B
C
D
Z-score
Position specific disorder score
Disorder score
Consensus approach #02
E
F
G
H
Start
Position
End

Figure S10

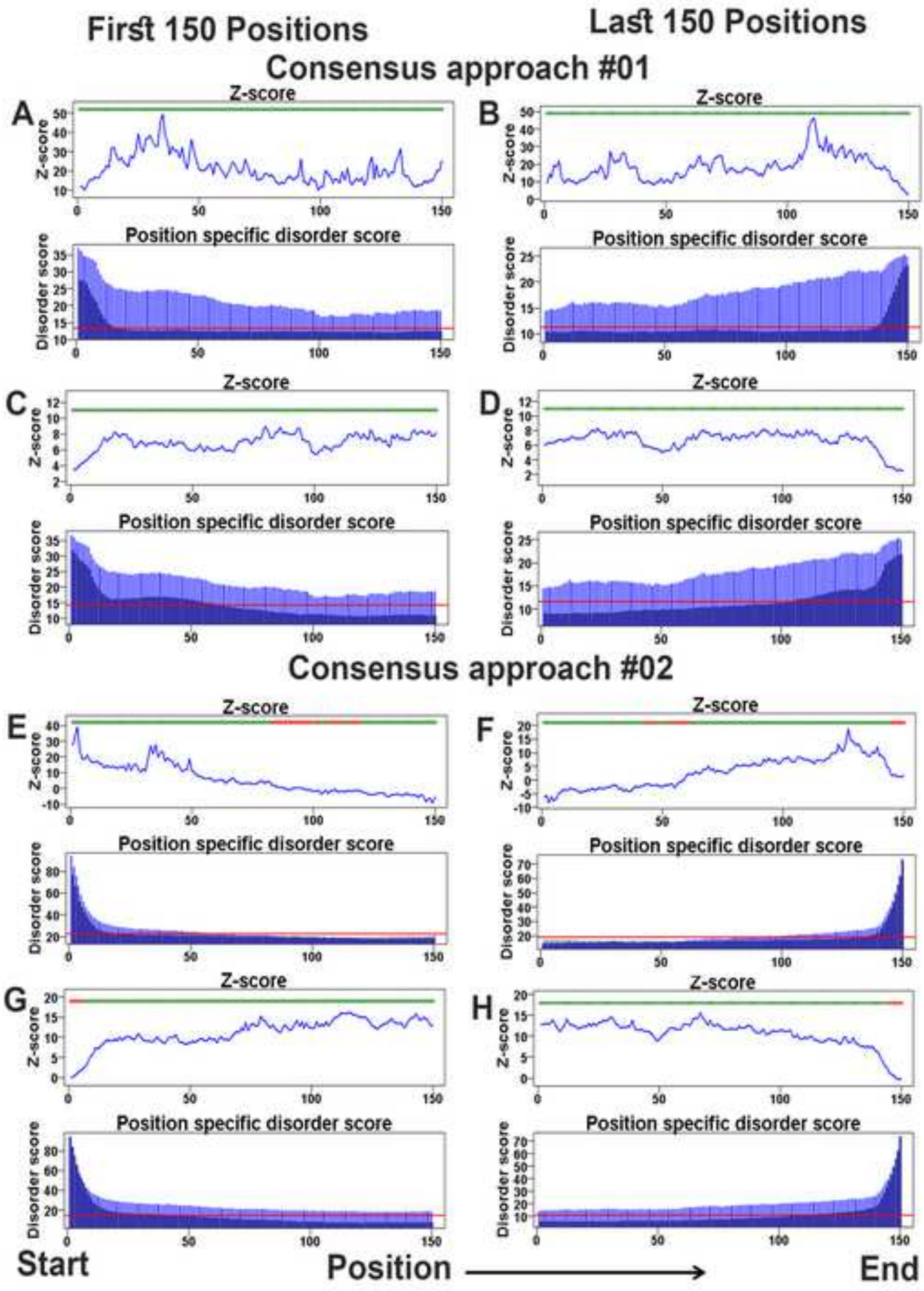
First 150 Positions
Last 150 Positions
Consensus approach #01
Consensus approach #02
A
B
C
D
E
F
G
H
Z-score
Position specific disorder score
Disorder score
Start
Position
End

Figure S11

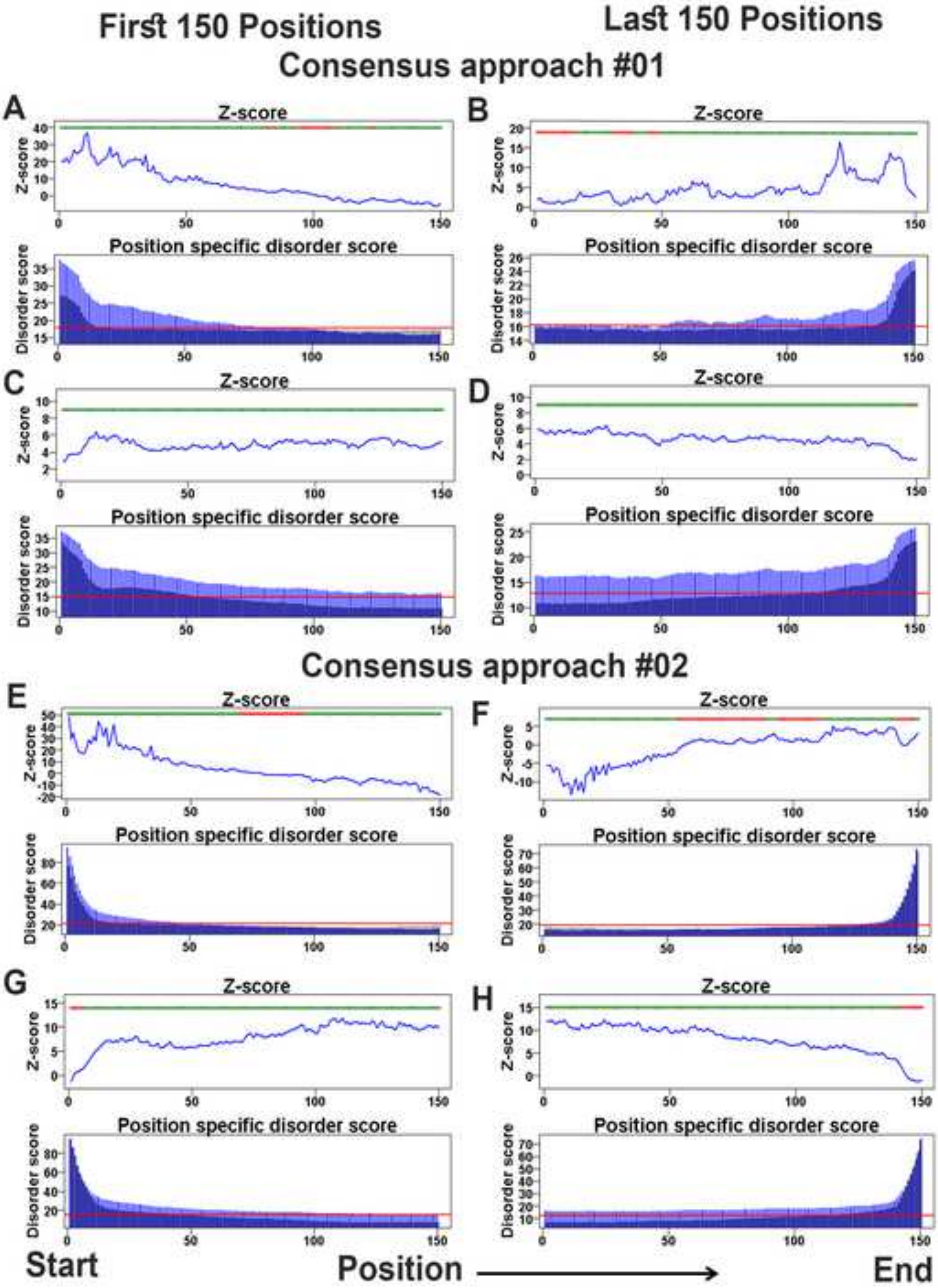
First 150 Positions
Last 150 Positions
Consensus approach #01
A
B
C
D
Z-score
Position specific disorder score
Disorder score
Consensus approach #02
E
F
G
H
Start
Position
End

Figure S12

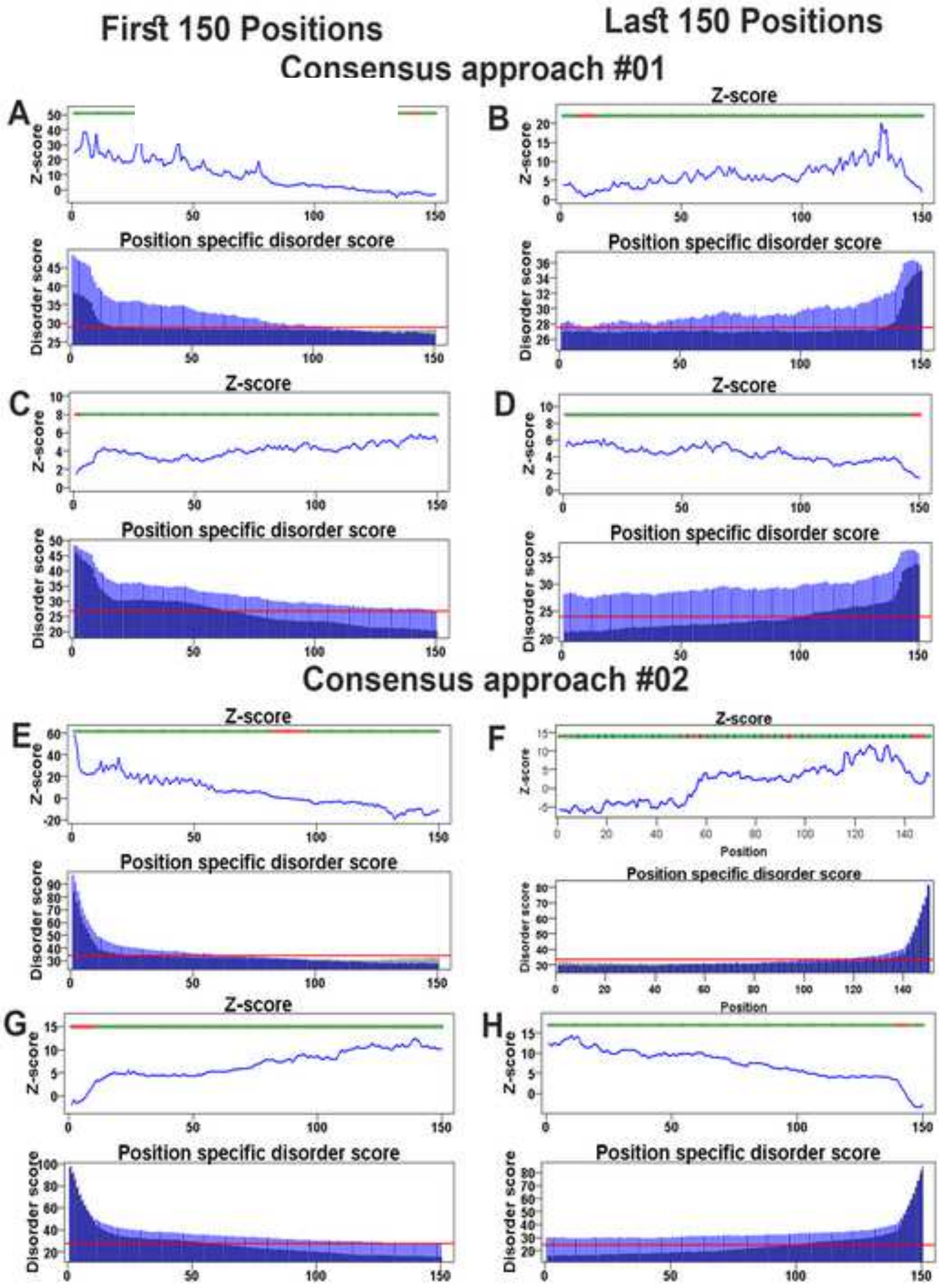

First 150 Positions
Last 150 Positions
Consensus approach #01
Consensus approach #02
A
B
C
D
E
F
G
H
Z-score
Position specific disorder score
Disorder score
Position

Figure S13

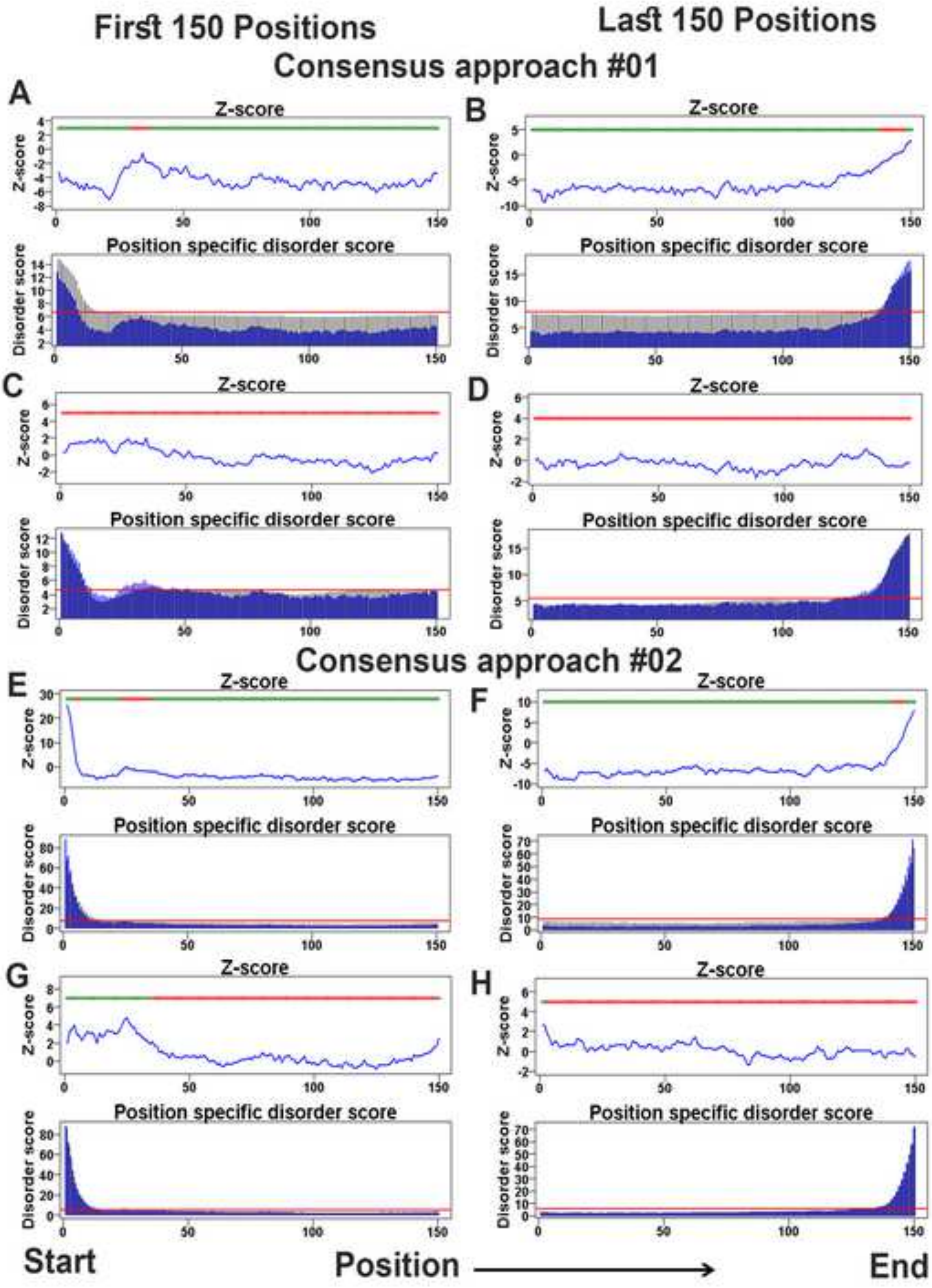

First 150 Positions
Last 150 Positions
Consensus approach #01
A
B
C
D
Consensus approach #02
E
F
G
H
Z-score
Position specific disorder score
Disorder score
Start
Position
End

Figure S14

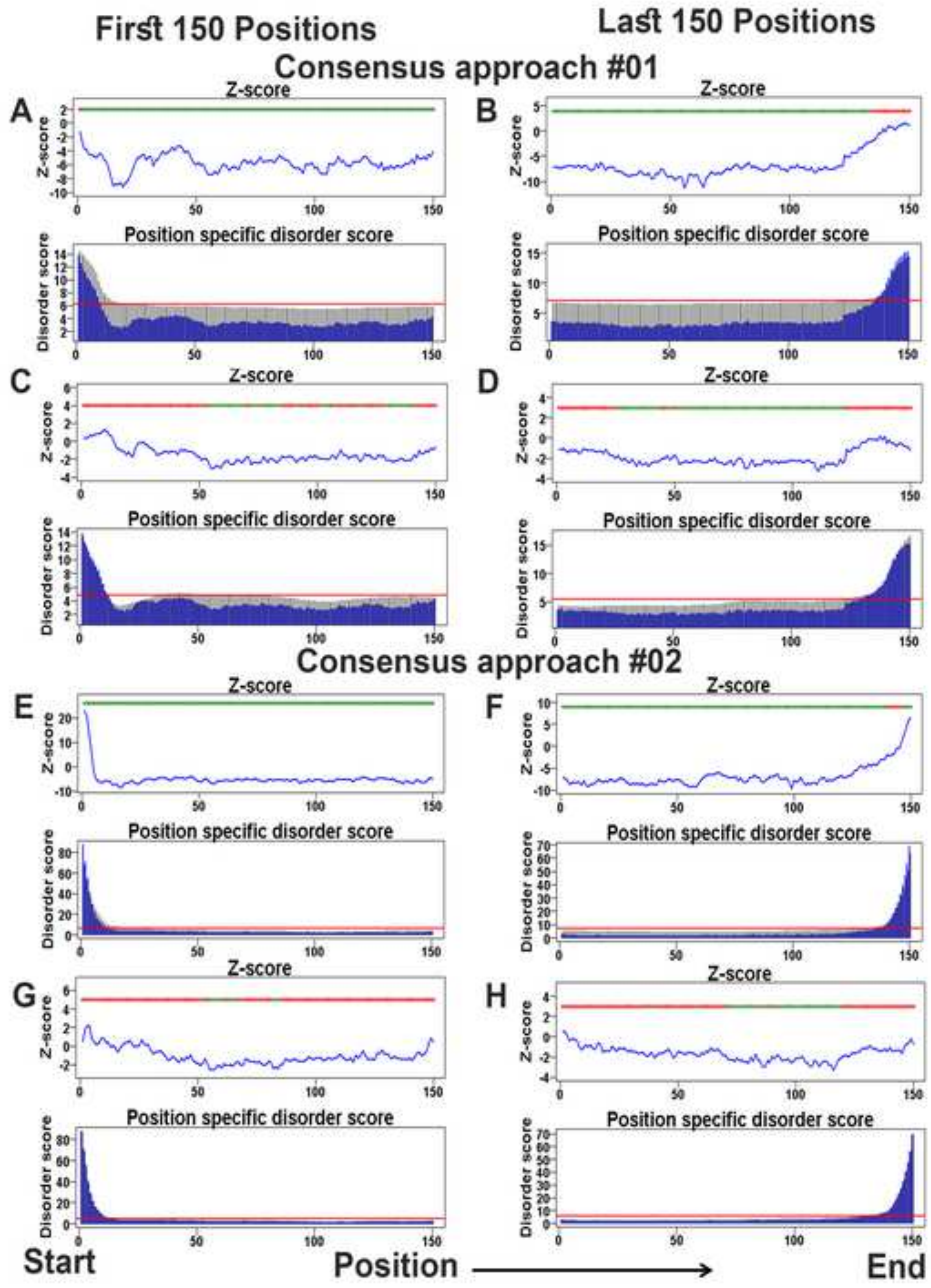

First 150 Positions
Last 150 Positions
Consensus approach #01
A
B
C
D
Z-score
Position specific disorder score
Disorder score
Consensus approach #02
E
F
G
H
Start
Position
End

Figure S15

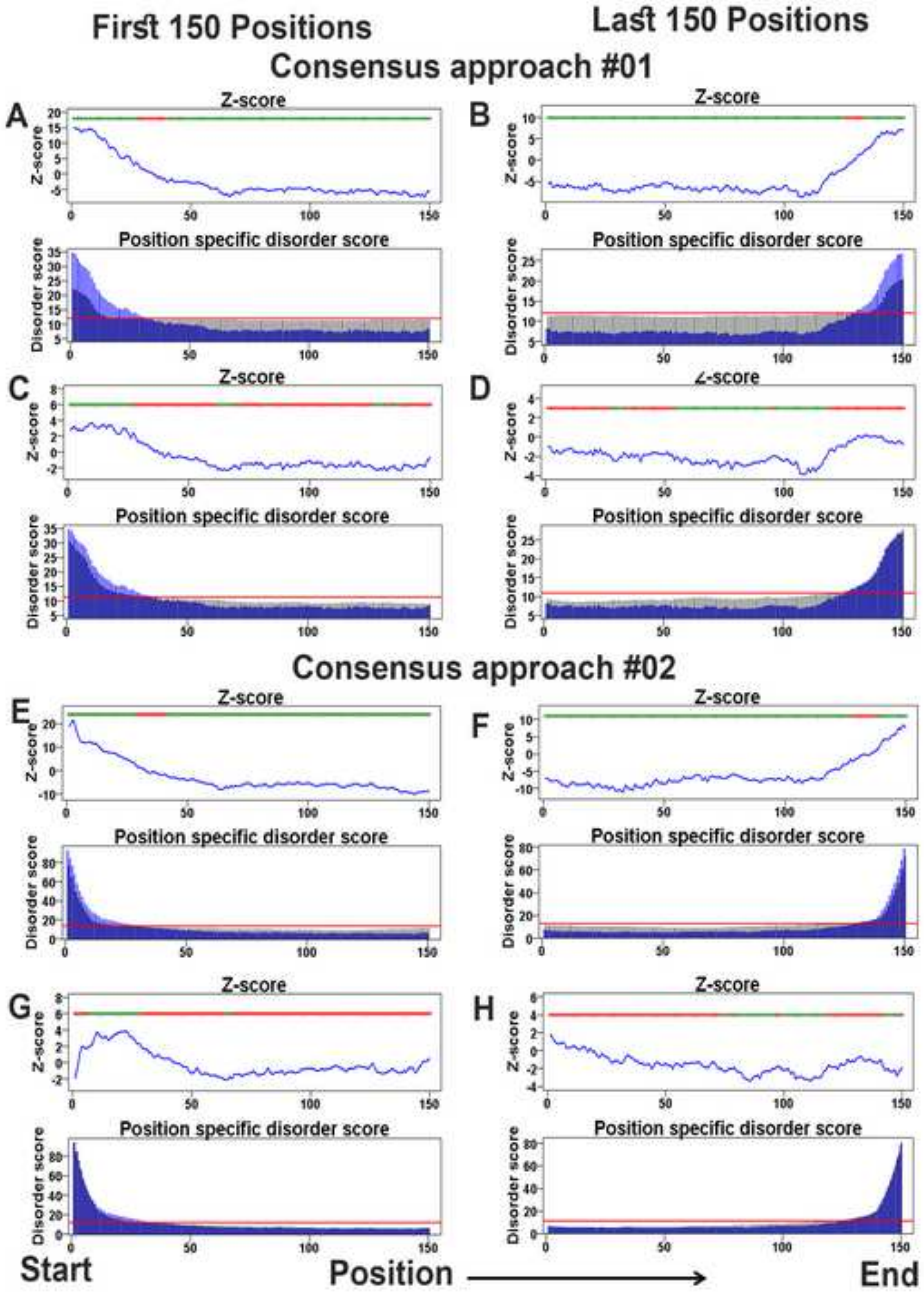

First 150 Positions
Last 150 Positions
Consensus approach #01
A
B
C
D
Z-score
Position specific disorder score
Disorder score
Consensus approach #02
E
F
G
H
Start
Position
End

**Figure S16**

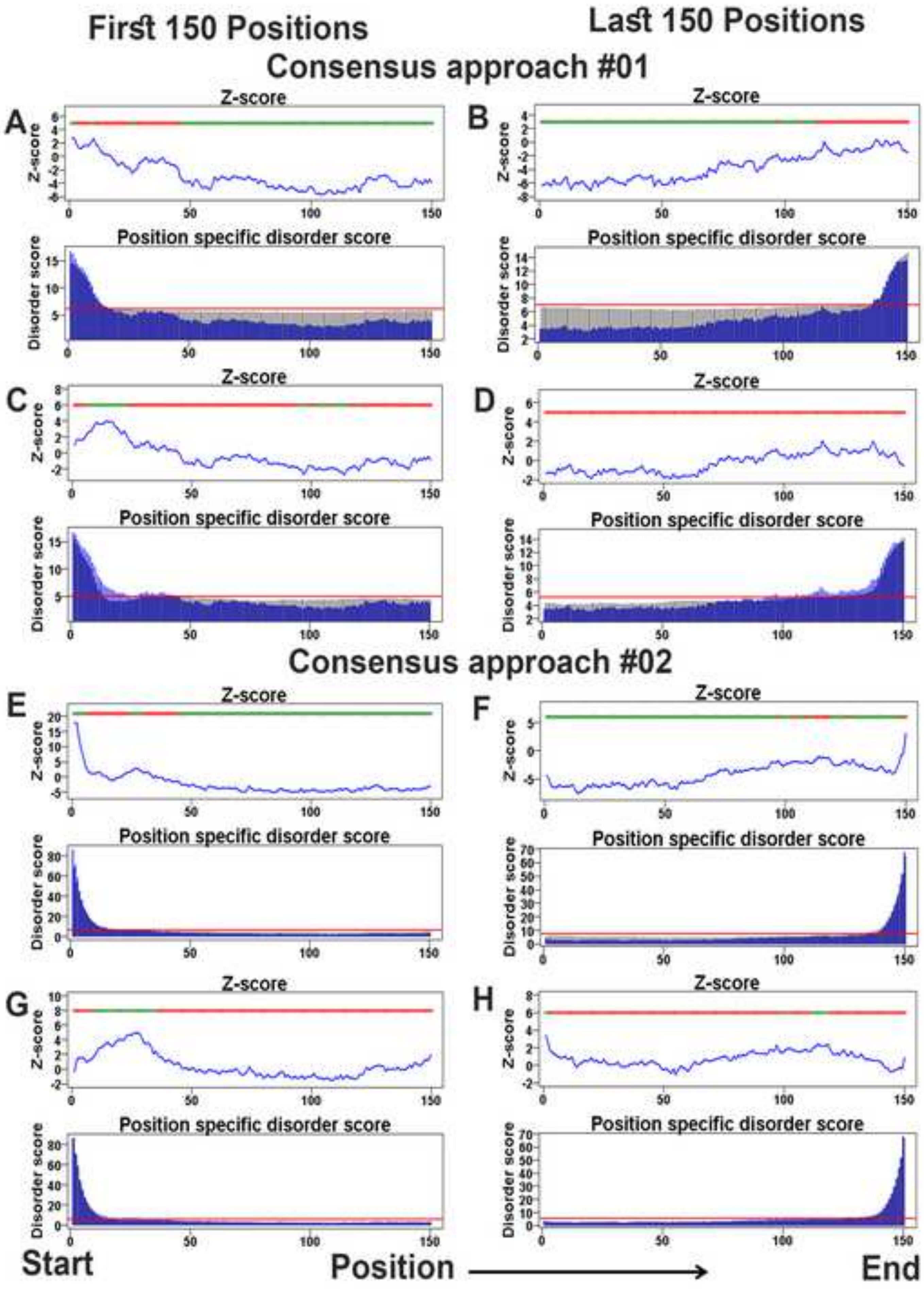

First 150 Positions
Last 150 Positions
Consensus approach #01
Consensus approach #02
A
B
C
D
E
F
G
H
Z-score
Position specific disorder score
Disorder score
Start
Position
End

Figure S17

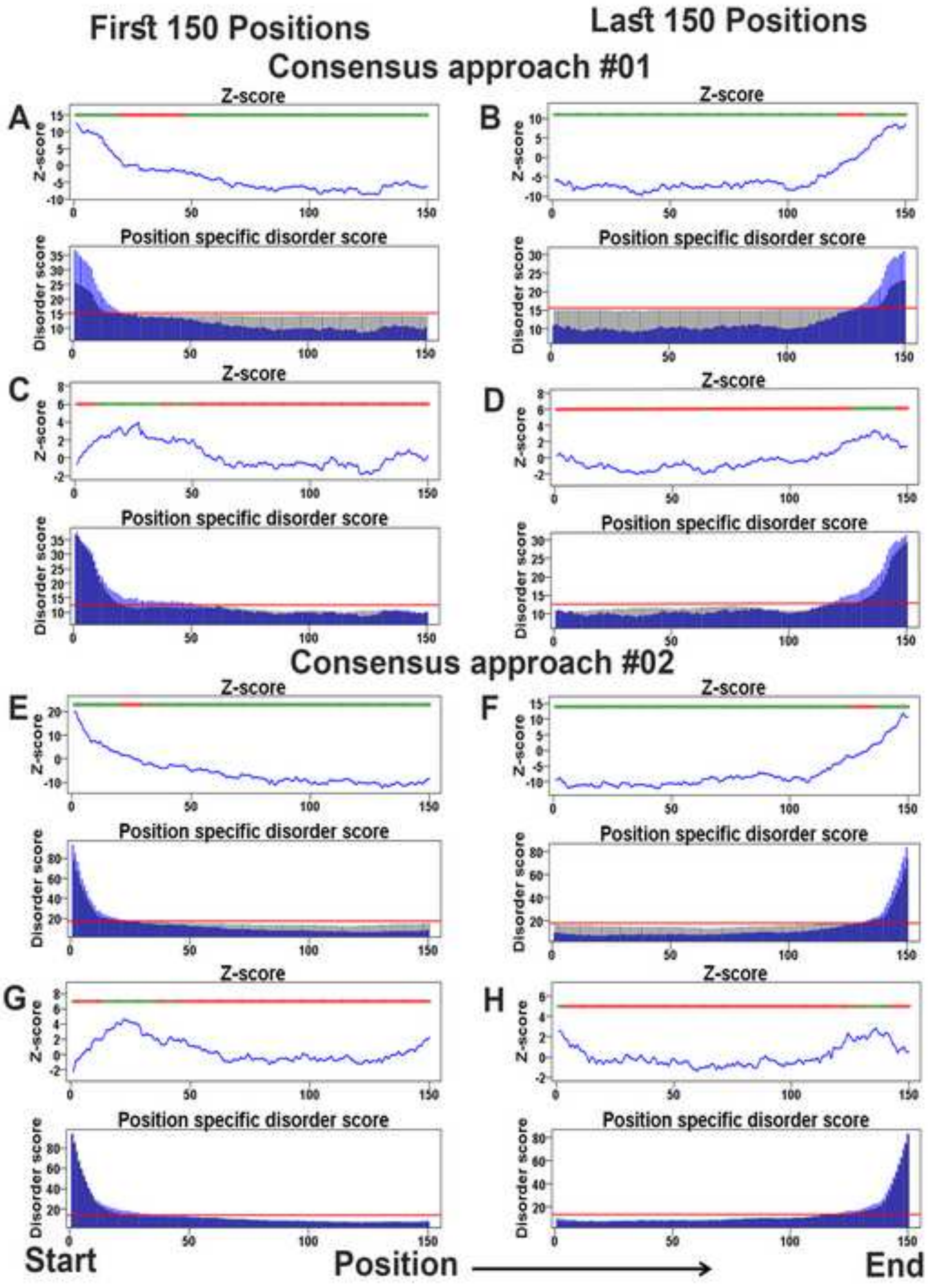

First 150 Positions
Last 150 Positions
Consensus approach #01
A
B
C
D
Consensus approach #02
E
F
G
H
Z-score
Position specific disorder score
Disorder score
Start
Position
End

Figure S18

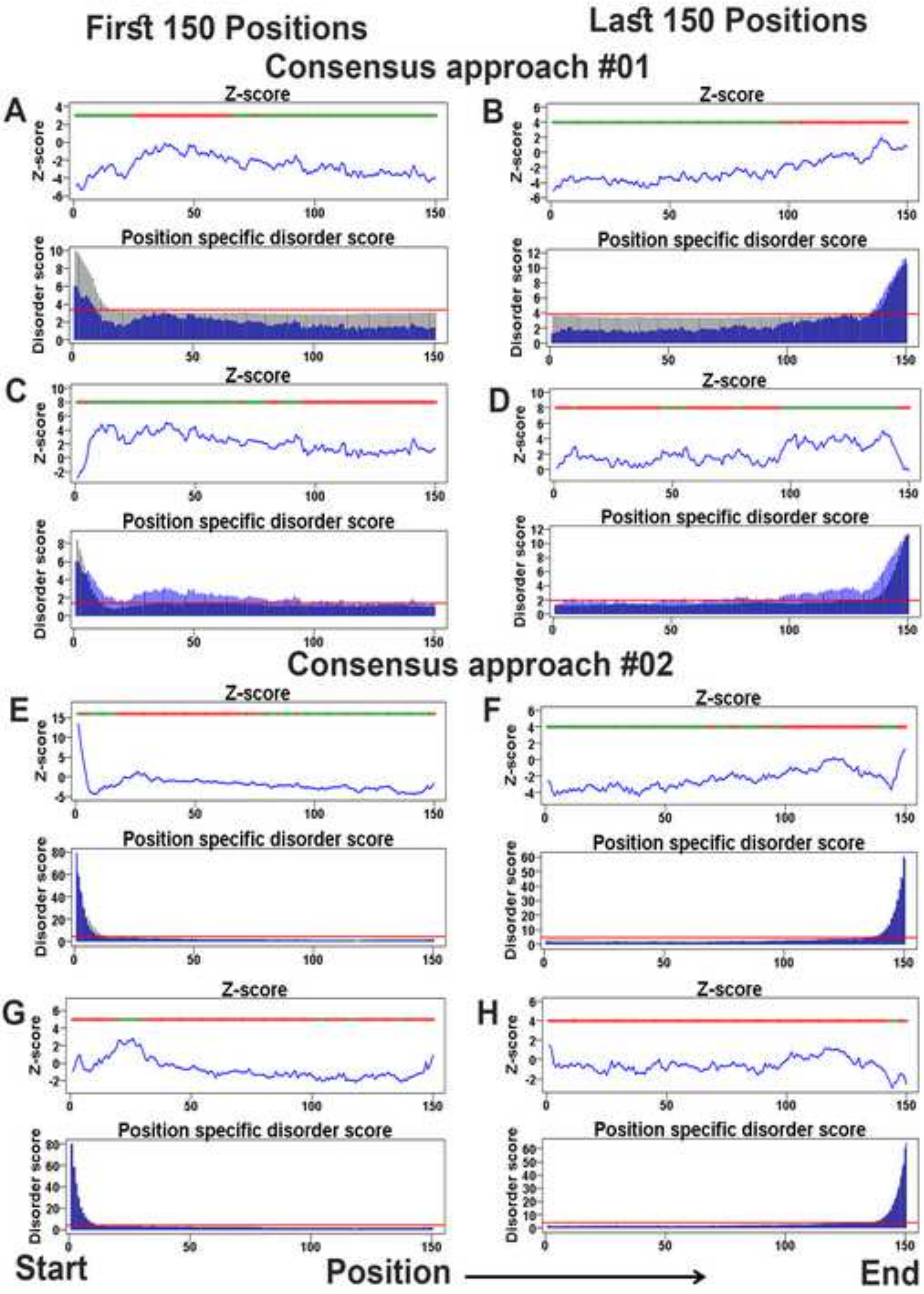

First 150 Positions
Last 150 Positions
Consensus approach #01
A
B
C
D
Z-score
Position specific disorder score
Disorder score
Consensus approach #02
E
F
G
H
Start
Position
End

**Figure S19**

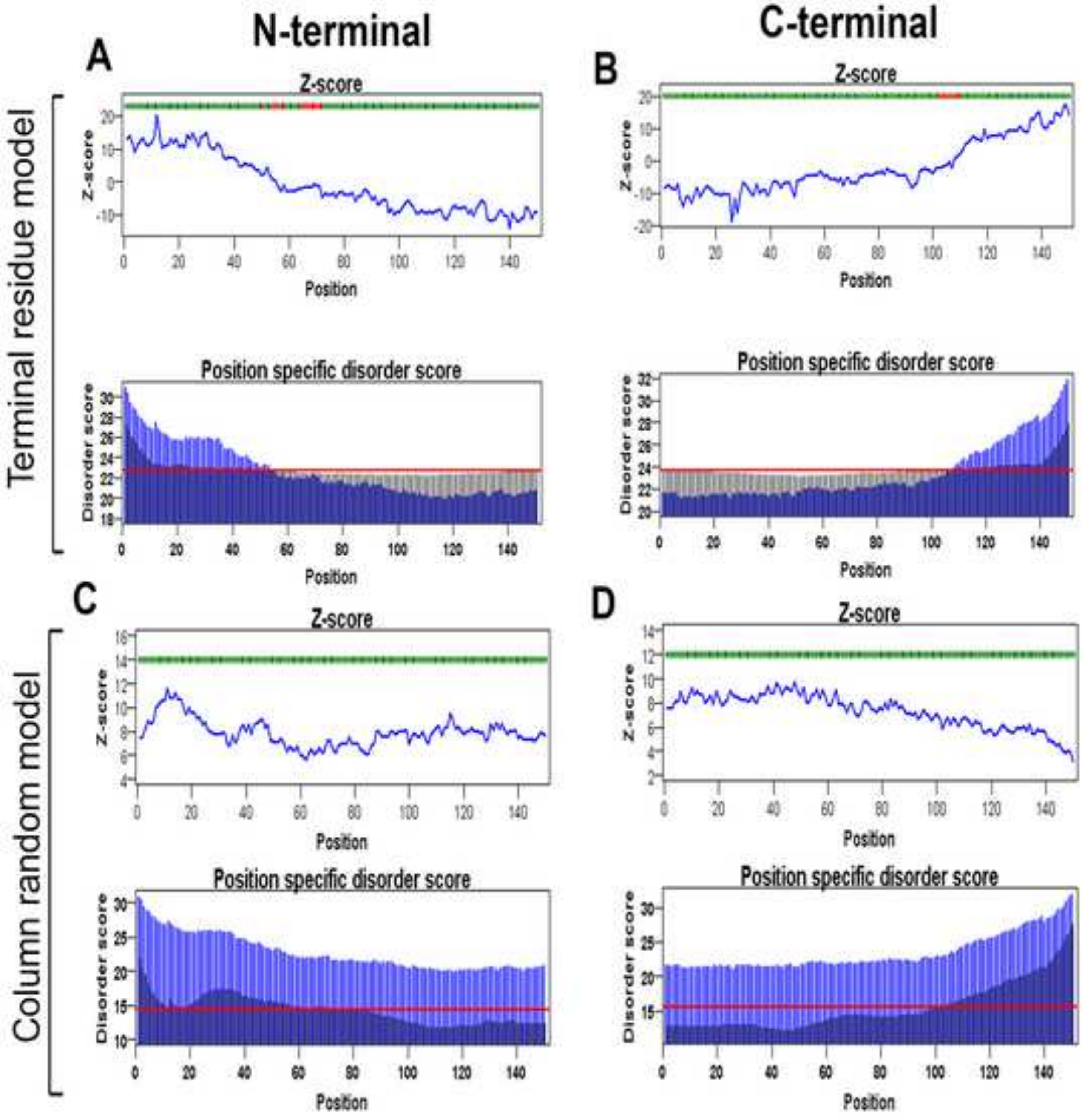

N-terminal
C-terminal
A
B
C
D
Terminal residue model
Column random model
Z-score
Position specific disorder score
Z-score
Disorder score
Position

**Figure S20**

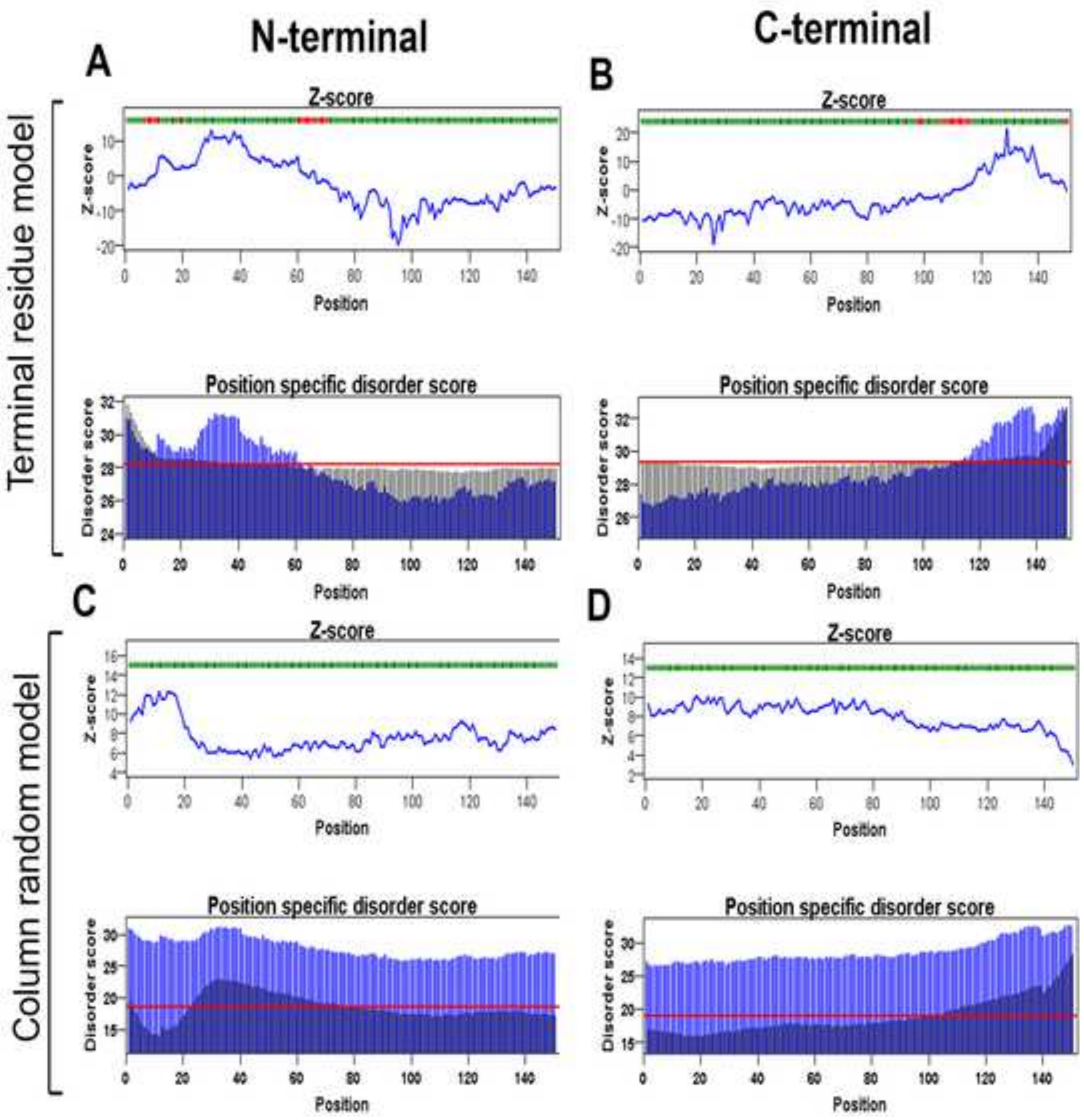
N-terminal
C-terminal
A
B
C
D
Terminal residue model
Column random model
Z-score
Position specific disorder score
Position
Disorder score

**Figure S21**

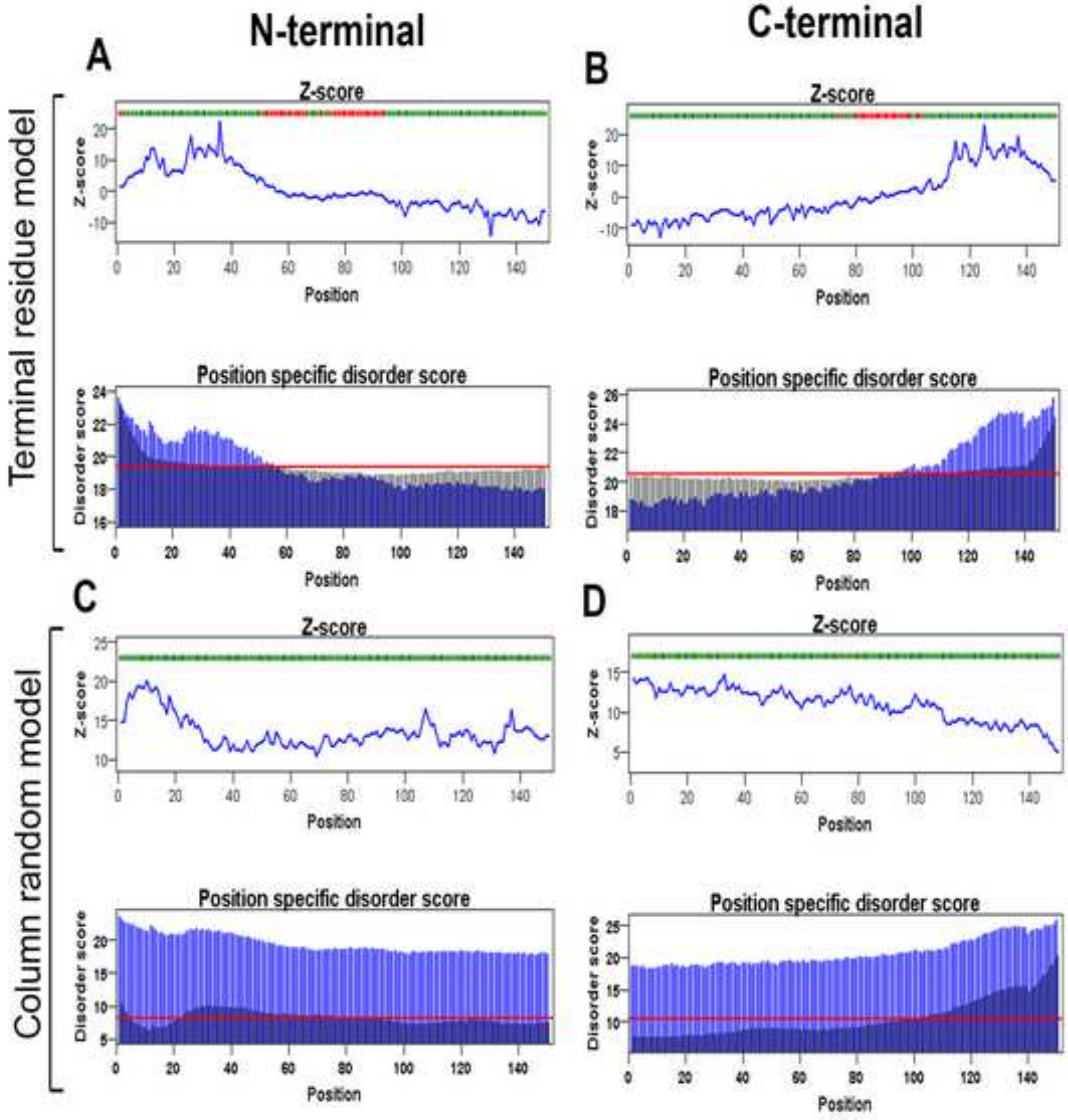
N-terminal
C-terminal
A
B
C
D
Terminal residue model
Column random model
Z-score
Position specific disorder score
Z-score
Disorder score
Position

**Figure S22**

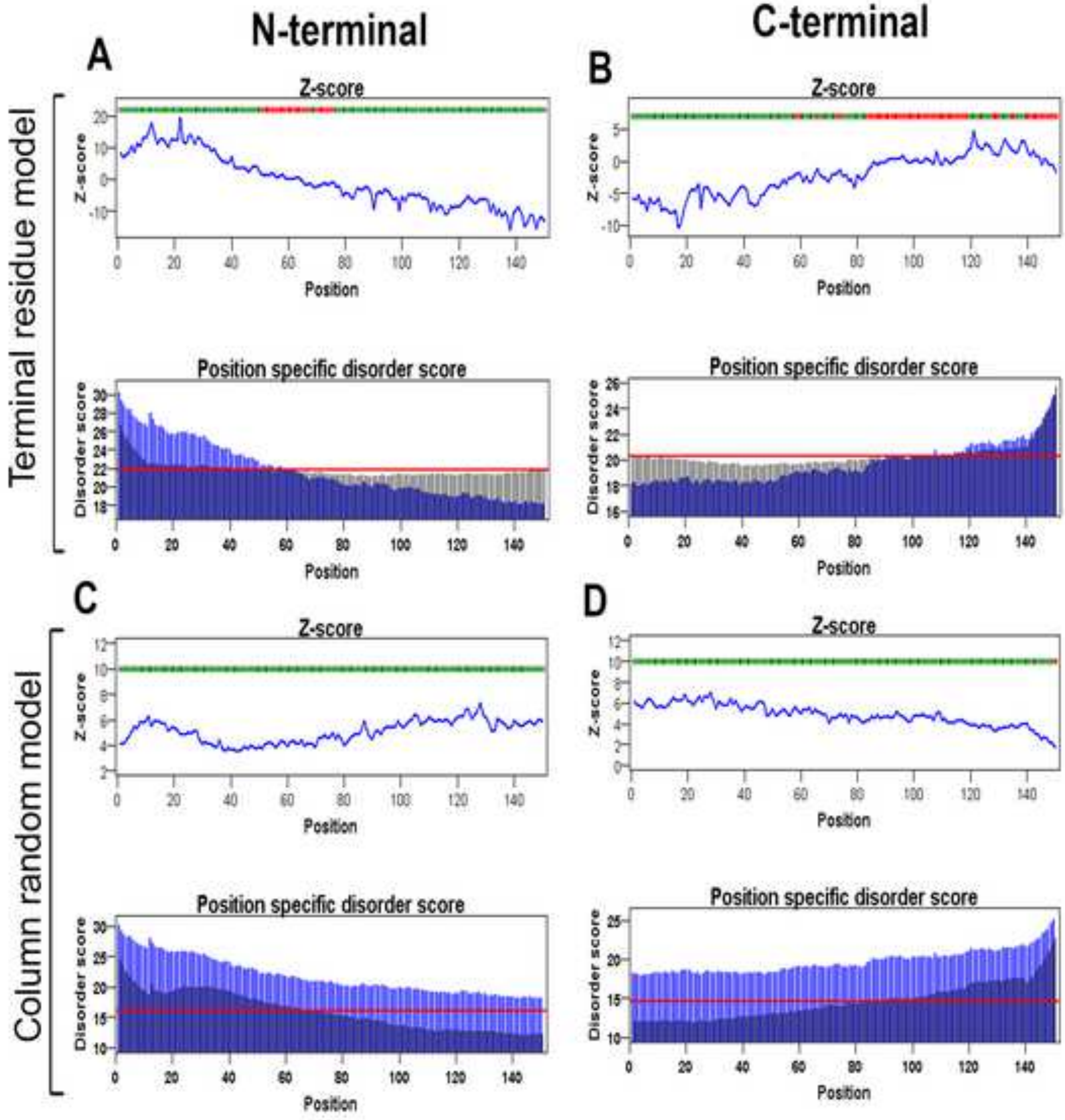

N-terminal
C-terminal
A
B
C
D
Terminal residue model
Column random model
Z-score
Position
Position specific disorder score
Disorder score

**Figure S23**

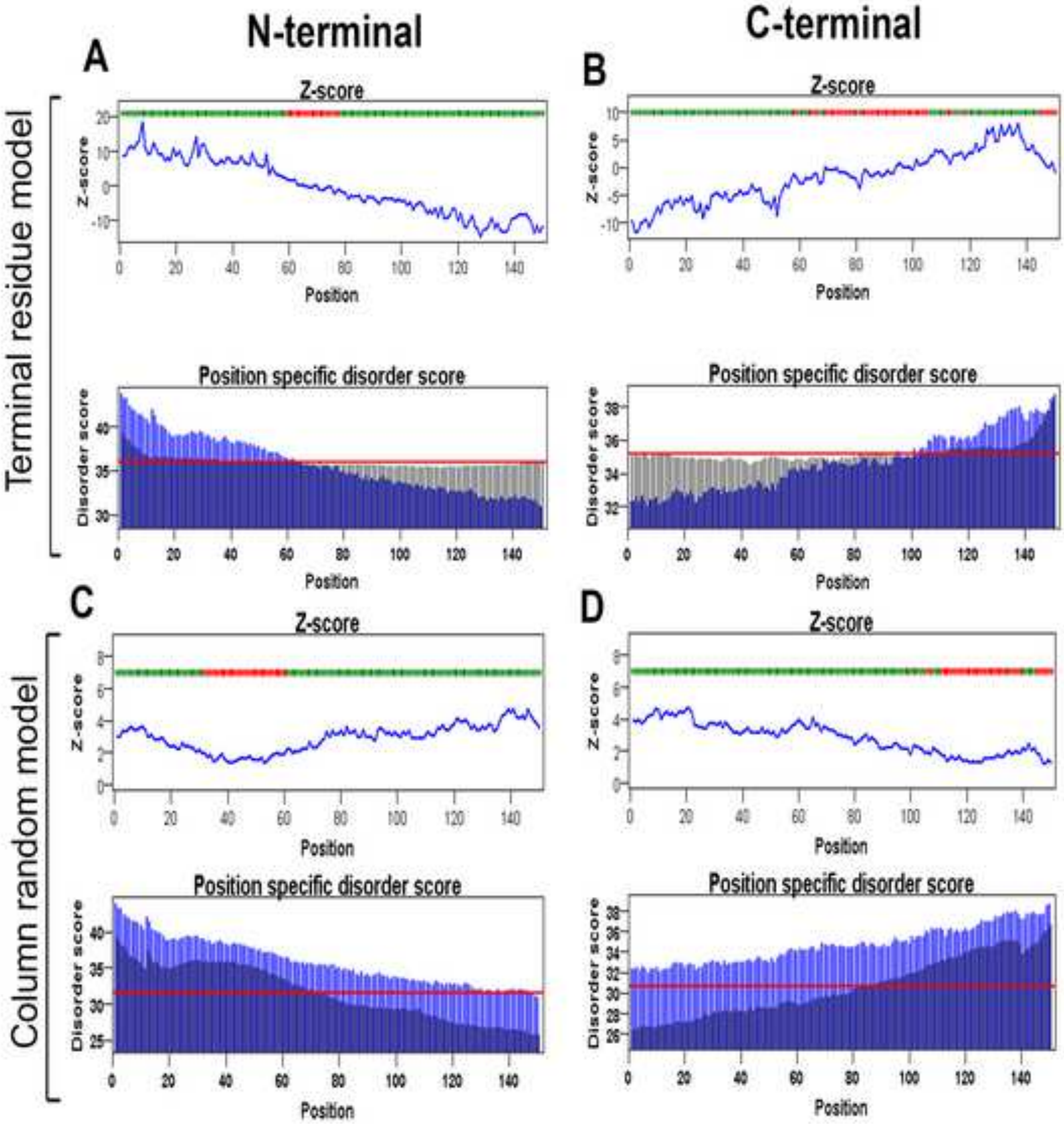

N-terminal
C-terminal
A
B
C
D
Terminal residue model
Column random model
Z-score
Position specific disorder score
Position
Z-score
Disorder score

**Figure S24**

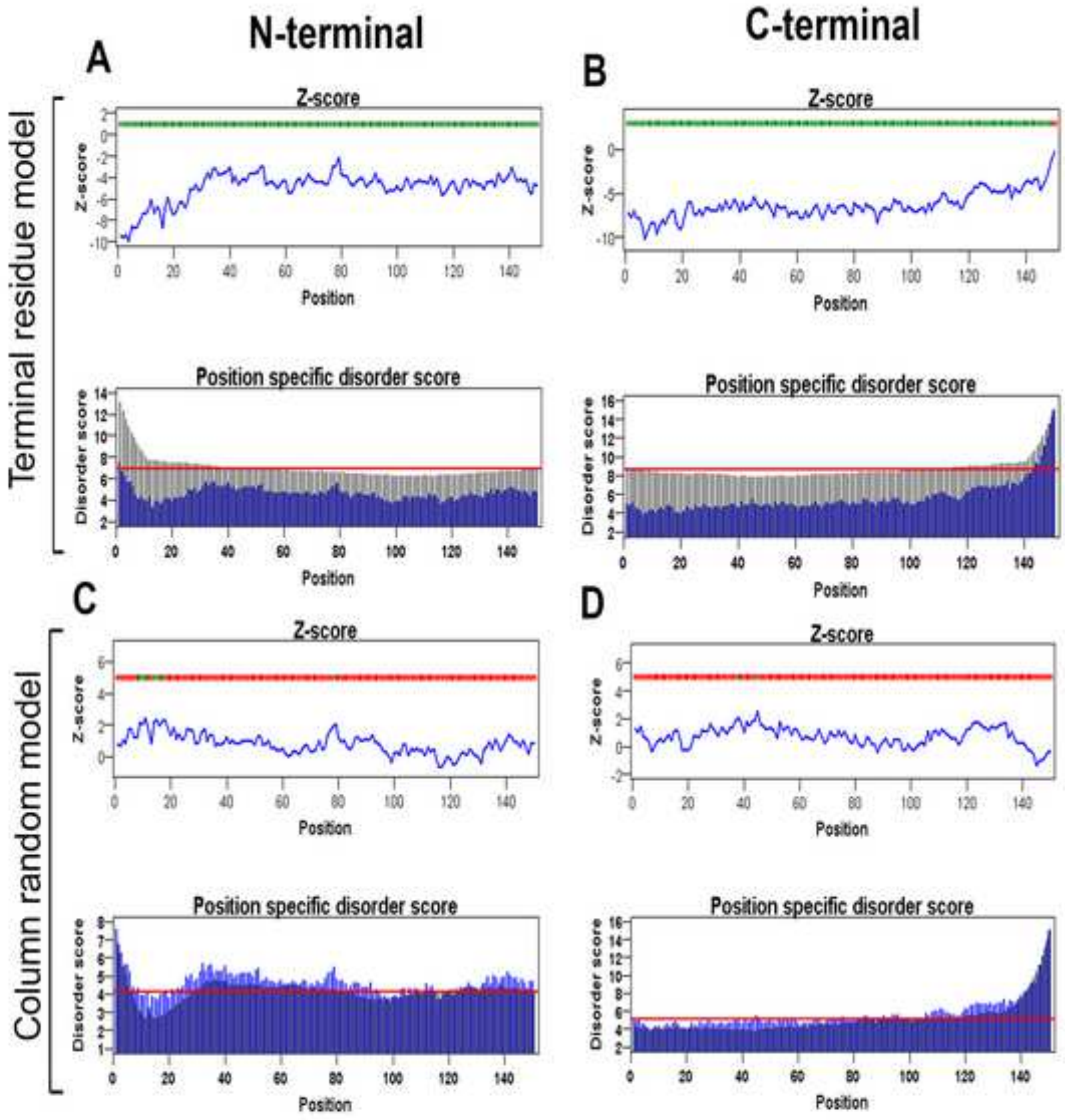
N-terminal
C-terminal
A
B
C
D
Terminal residue model
Column random model
Z-score
Position specific disorder score
Z-score
Disorder score
Position

**Figure S25**

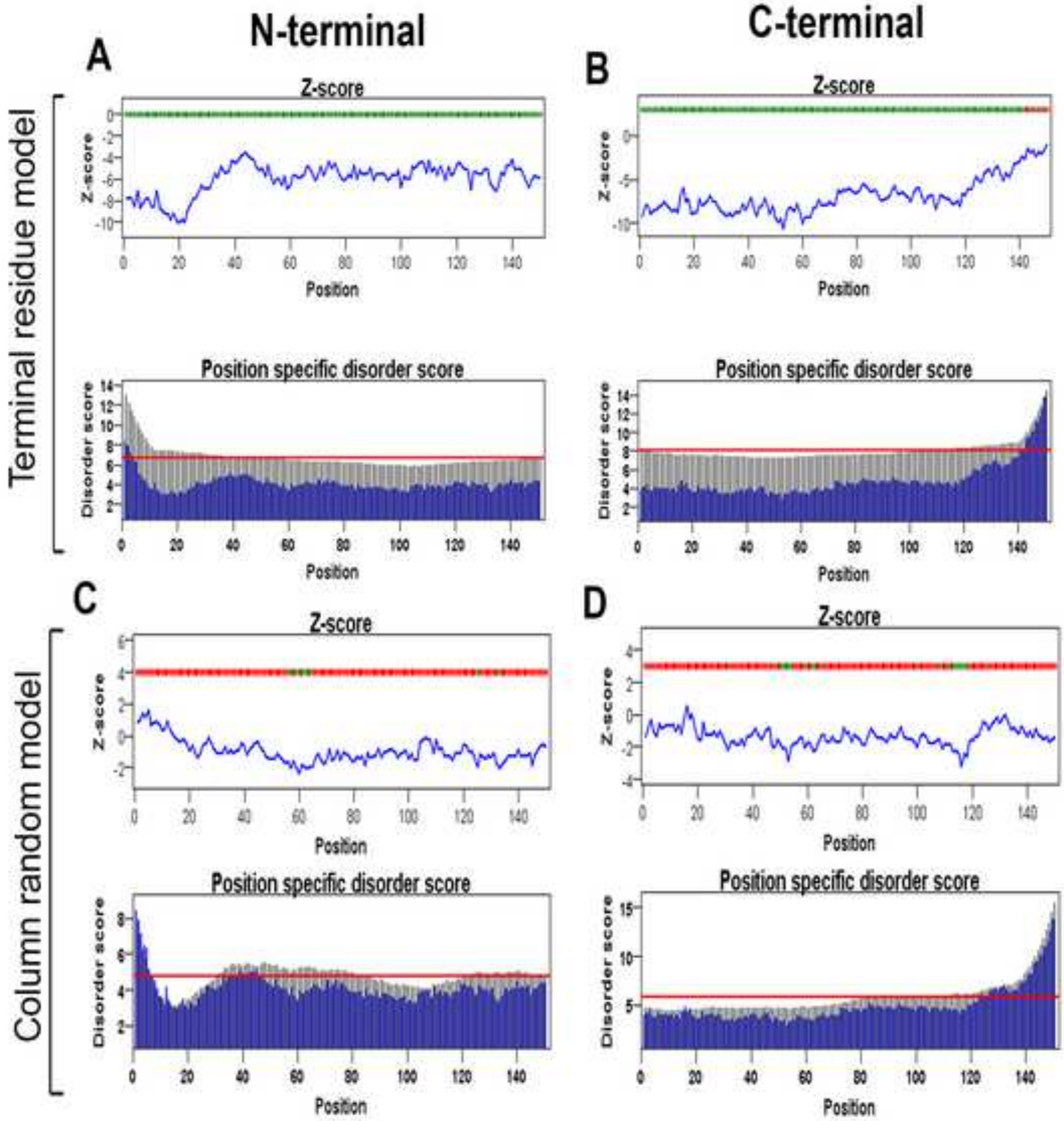

N-terminal
C-terminal
A
B
C
D
Terminal residue model
Column random model
Z-score
Position specific disorder score
Position
Z-score
Disorder score

**Figure S26**

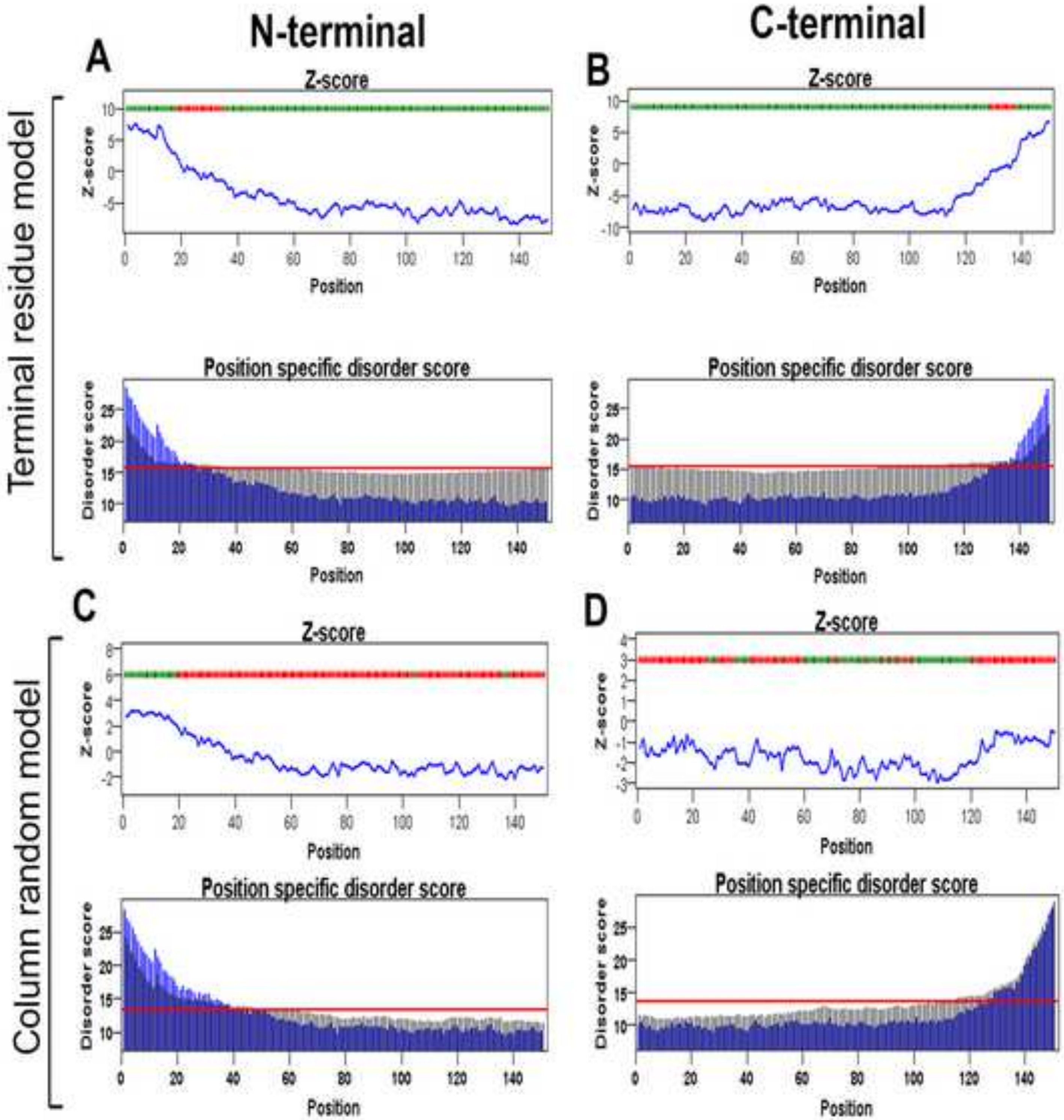
N-terminal
C-terminal
Terminal residue model
Column random model
A
B
C
D
Z-score
Position
Position specific disorder score
Disorder score

**Figure S27**

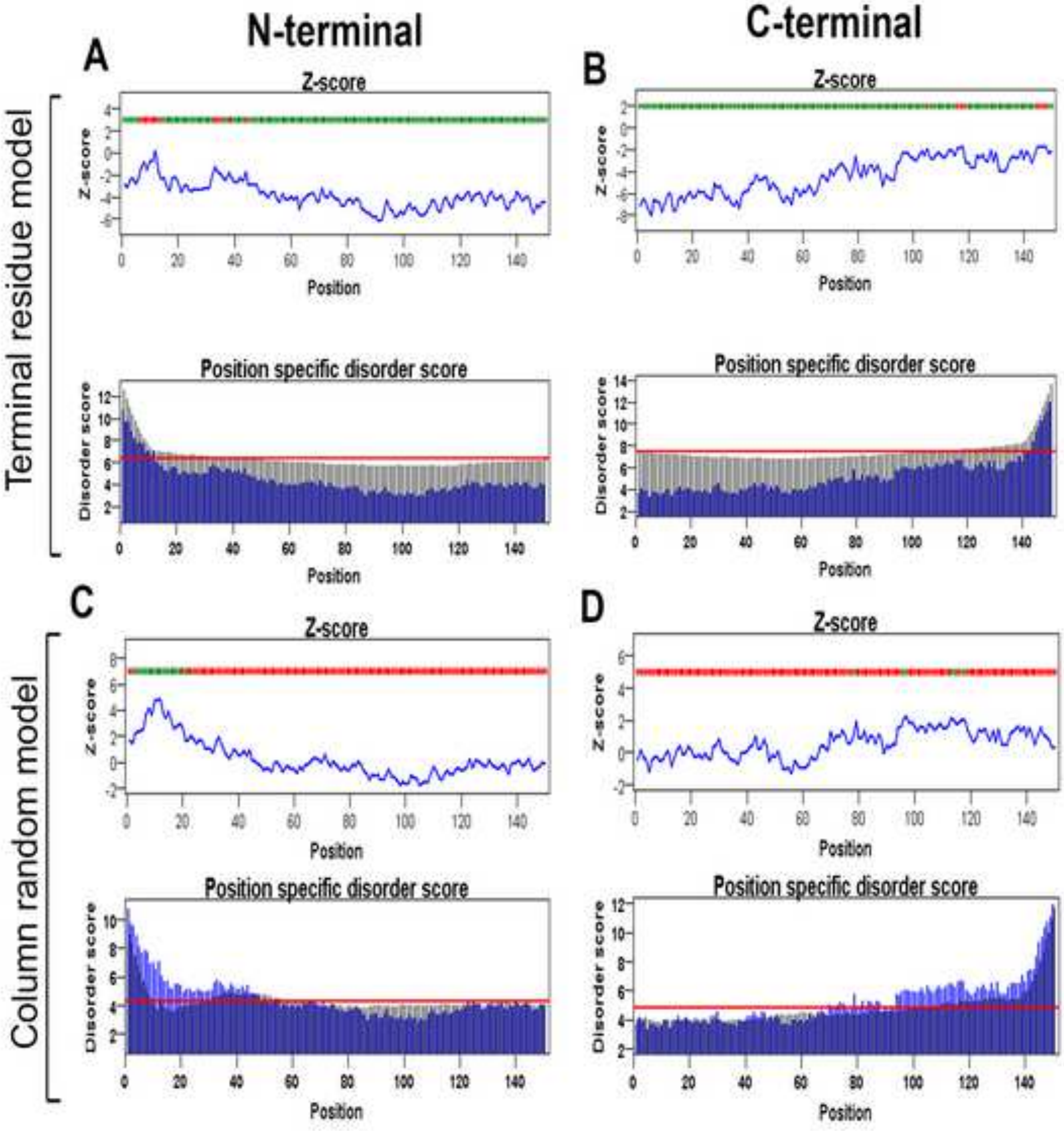

N-terminal
C-terminal
Terminal residue model
Column random model
A
B
C
D
Z-score
Position
Position specific disorder score
Disorder score

**Figure S28**

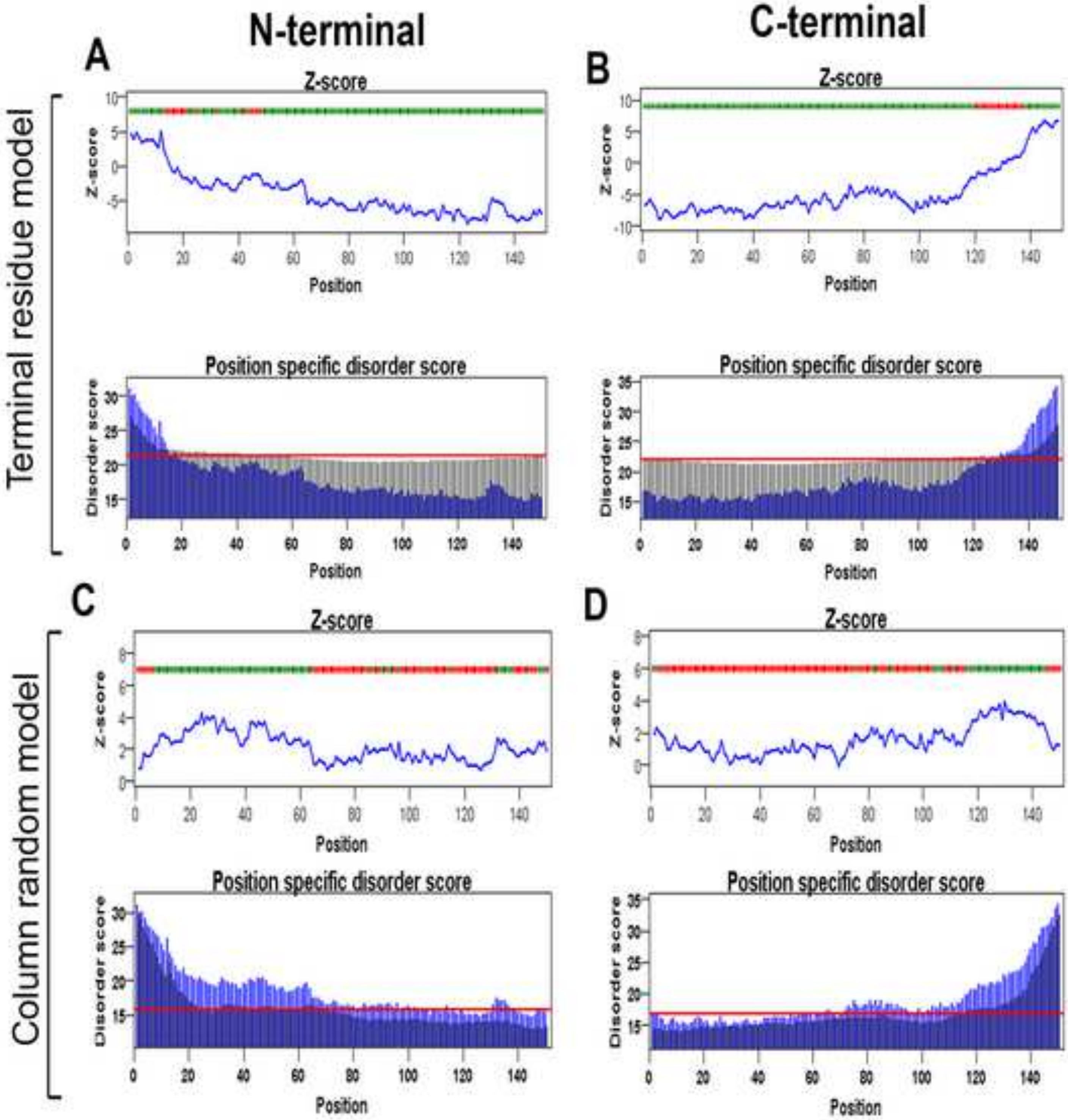
N-terminal
C-terminal
A
B
C
D
Terminal residue model
Column random model
Z-score
Position specific disorder score
Z-score
Disorder score
Position

**Figure S29**

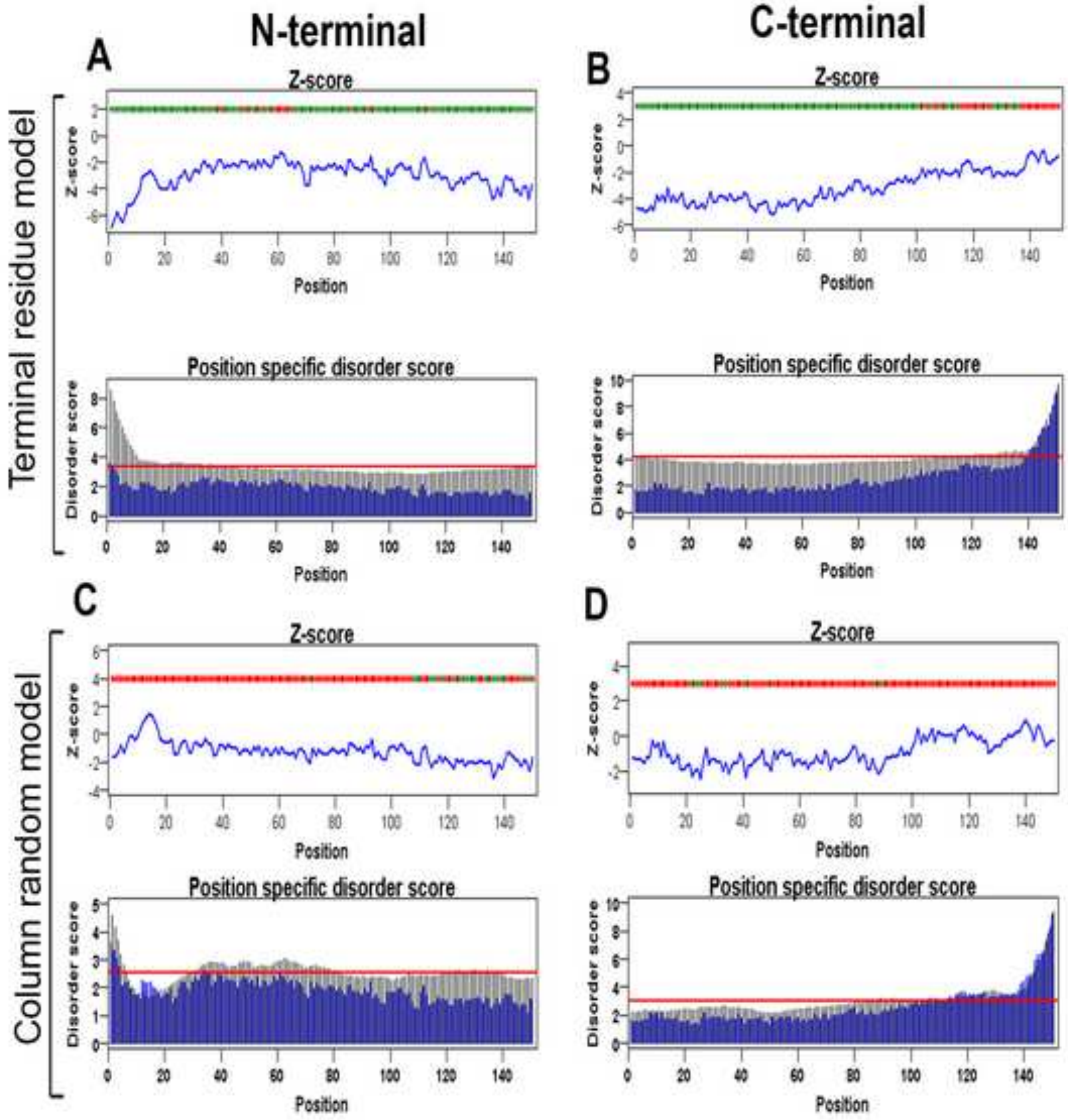

N-terminal
C-terminal
A
B
C
D
Terminal residue model
Column random model
Z-score
Z-score
Position
Disorder score
Position specific disorder score

**Figure S30**

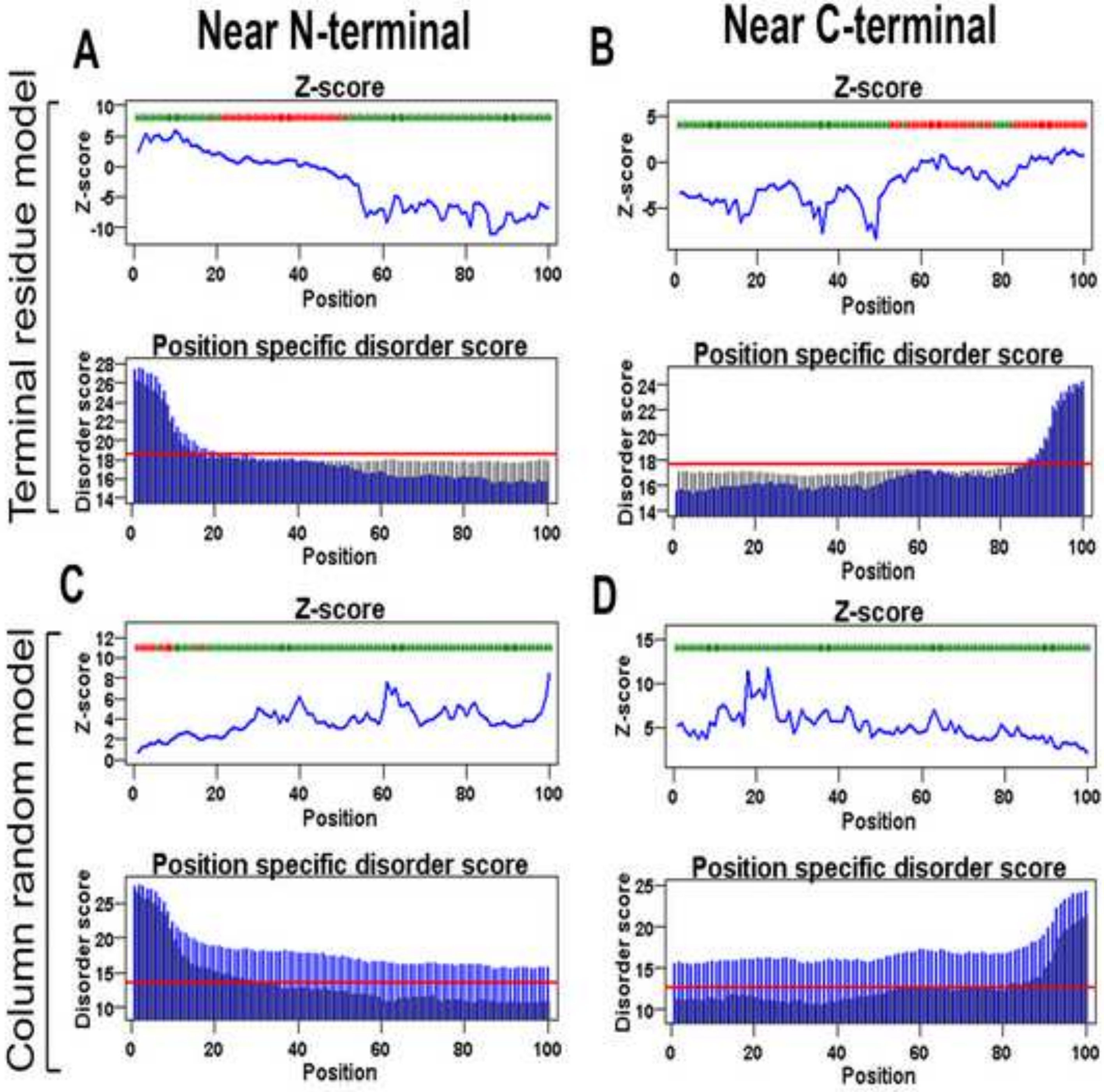

Near N-terminal
Near C-terminal
A
B
C
D
Terminal residue model
Column random model
Z-score
Position specific disorder score
Z-score
Disorder score
Position

**Figure S31**

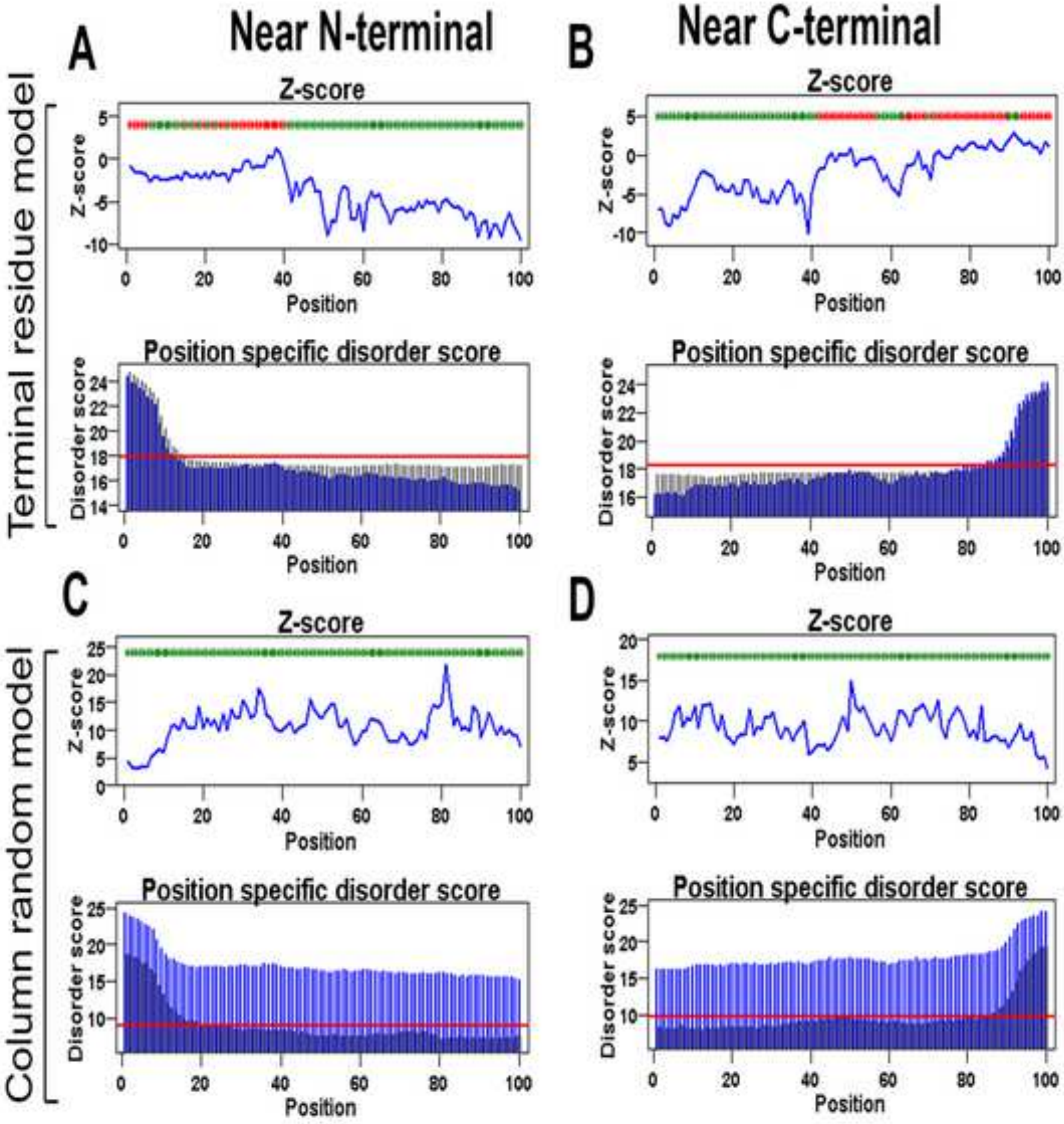

Near N-terminal
Near C-terminal
Terminal residue model
Column random model
A
B
C
D
Z-score
Position specific disorder score
Z-score
Disorder score
Position

**Figure S32**

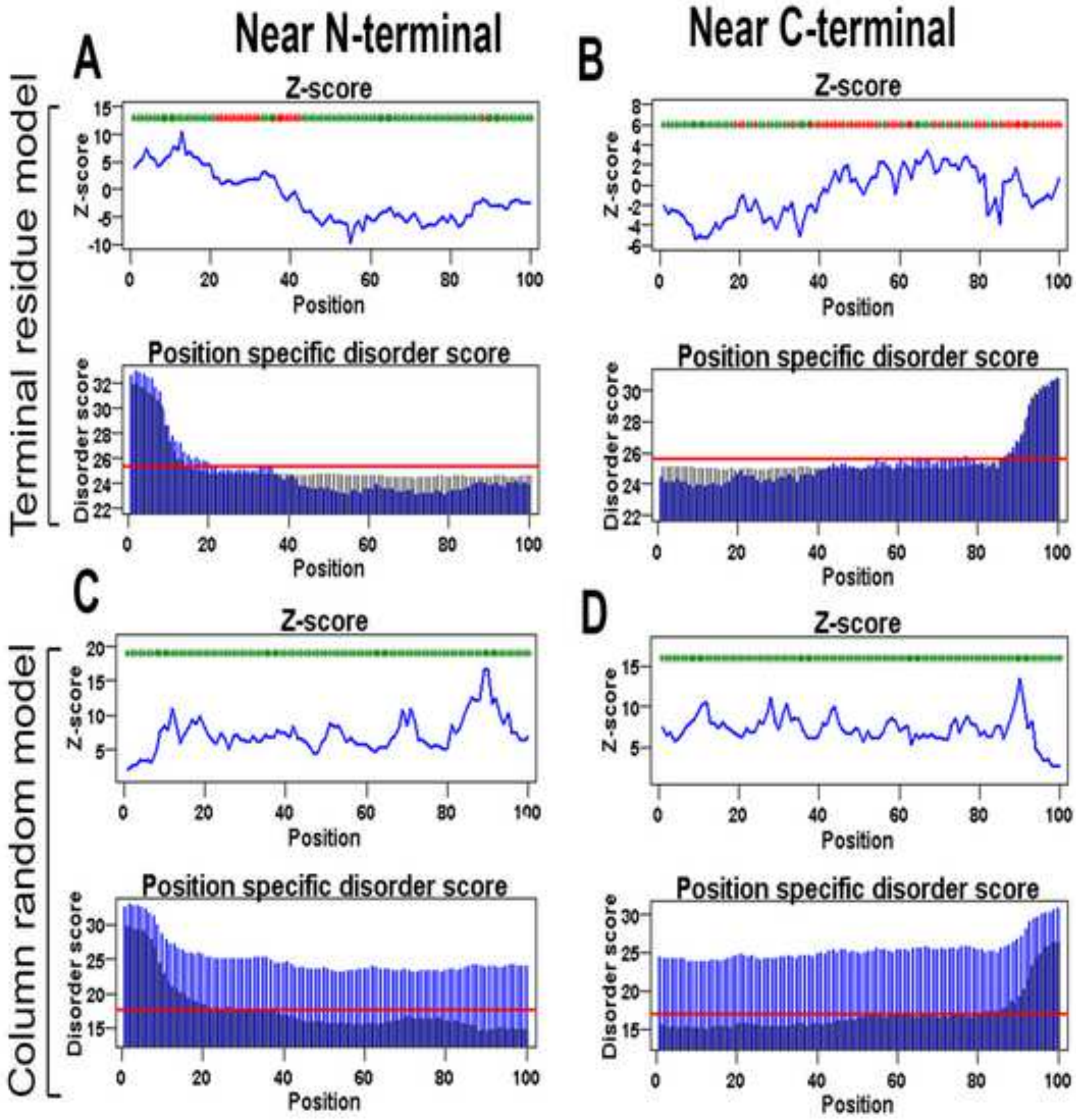

Near N-terminal
Near C-terminal
A
B
C
D
Terminal residue model
Column random model
Z-score
Position specific disorder score
Position
Disorder score

**Figure S33**

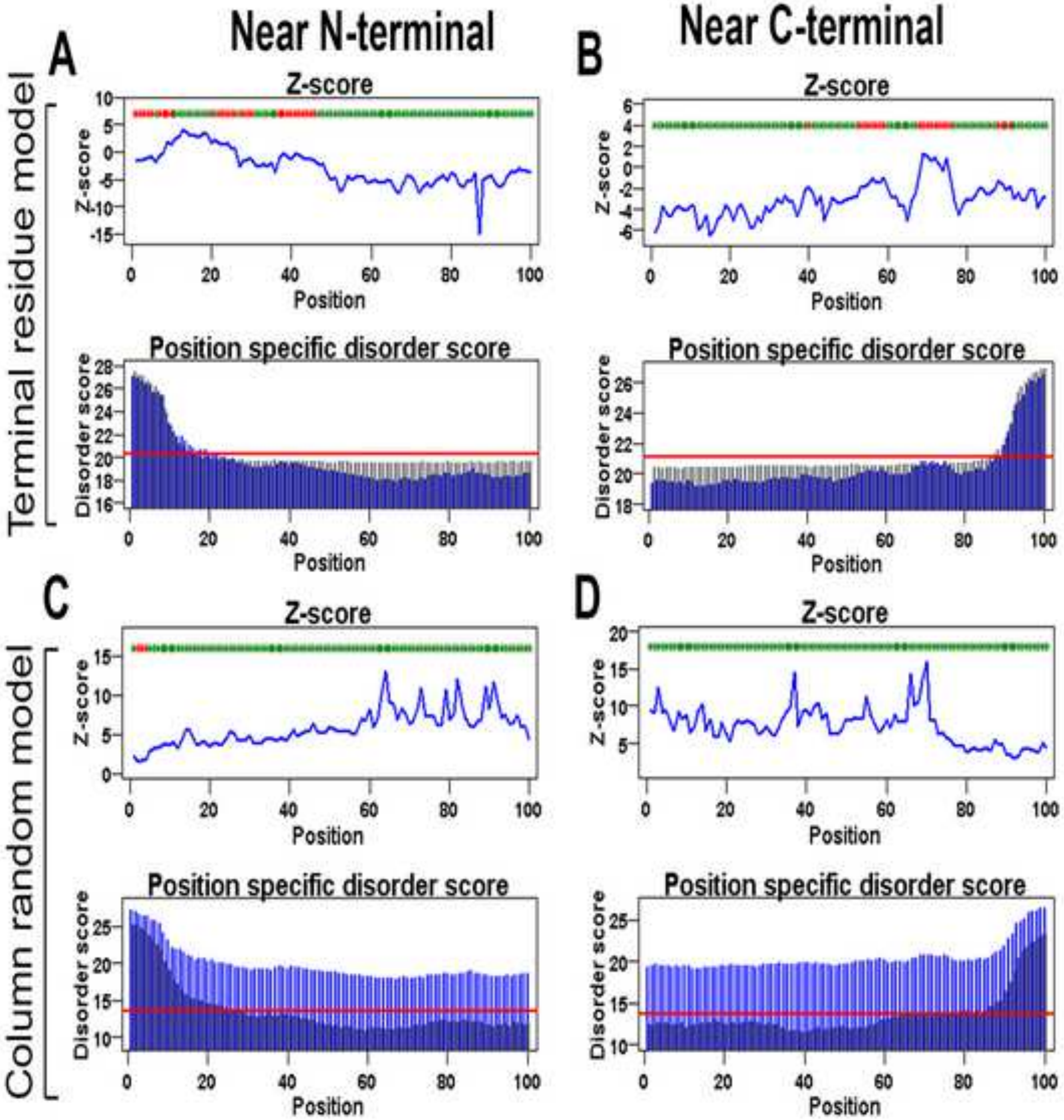

Near N-terminal
Near C-terminal
A
B
C
D
Terminal residue model
Column random model
Z-score
Position
Position specific disorder score
Disorder score

**Figure S34**

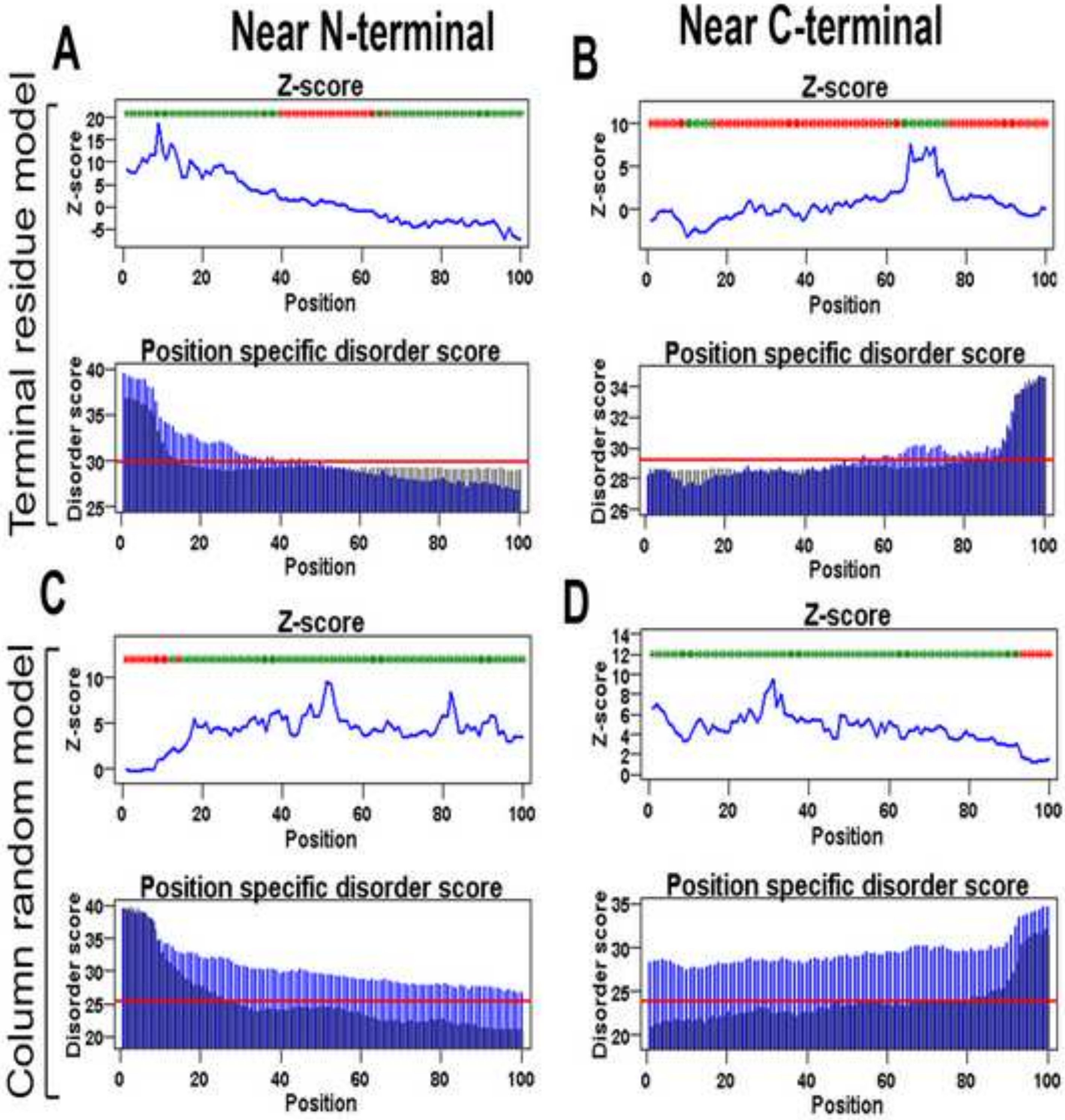

Terminal residue model
Column random model
A
Near N-terminal
B
Near C-terminal
Z-score
Position
Position specific disorder score
Disorder score
C
D

**Figure S35**

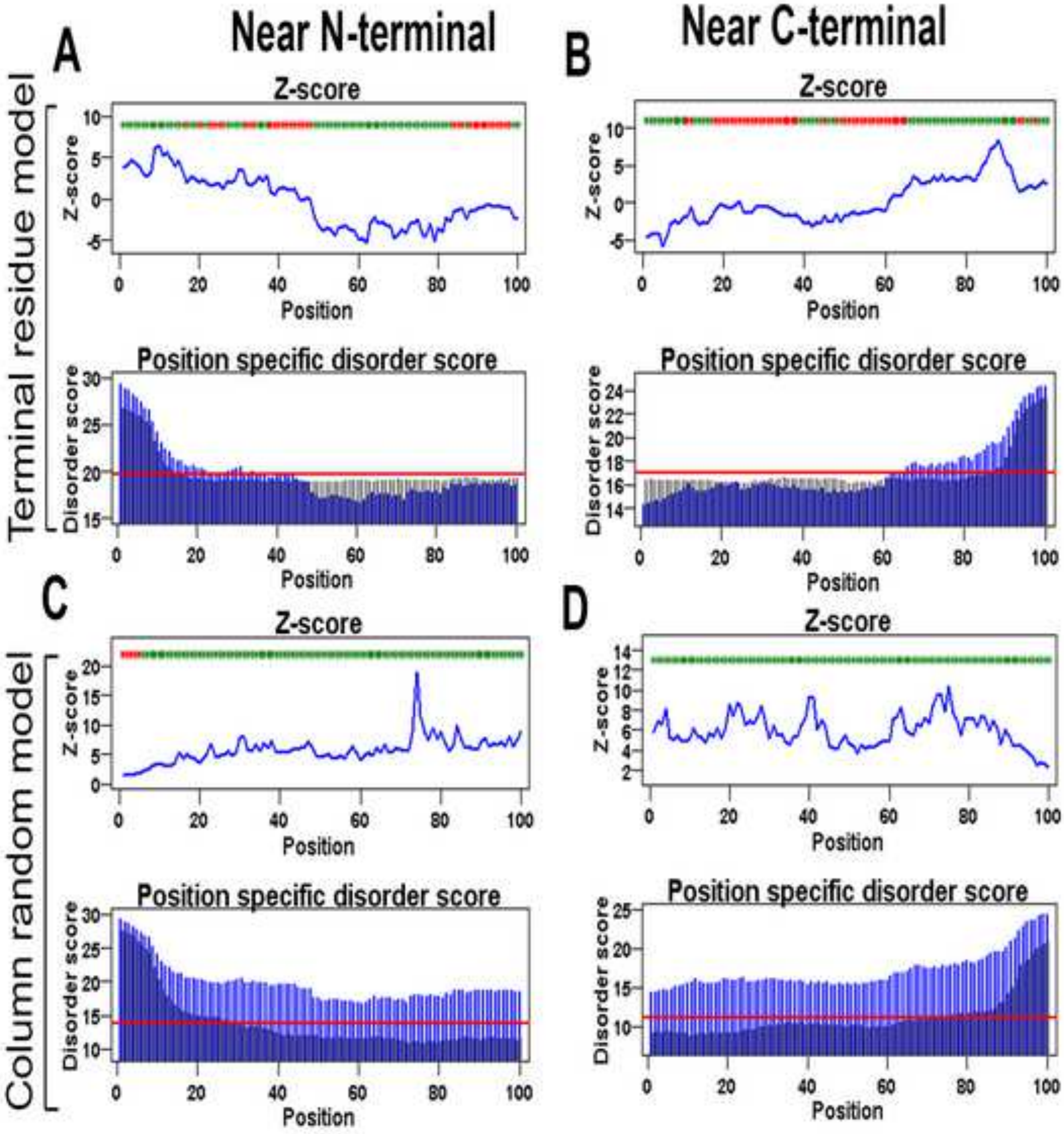

A
Near N-terminal
B
Near C-terminal
Terminal residue model
Z-score
Z-score
Position
Position specific disorder score
Disorder score
Position
C
D
Column random model

**Figure S36**

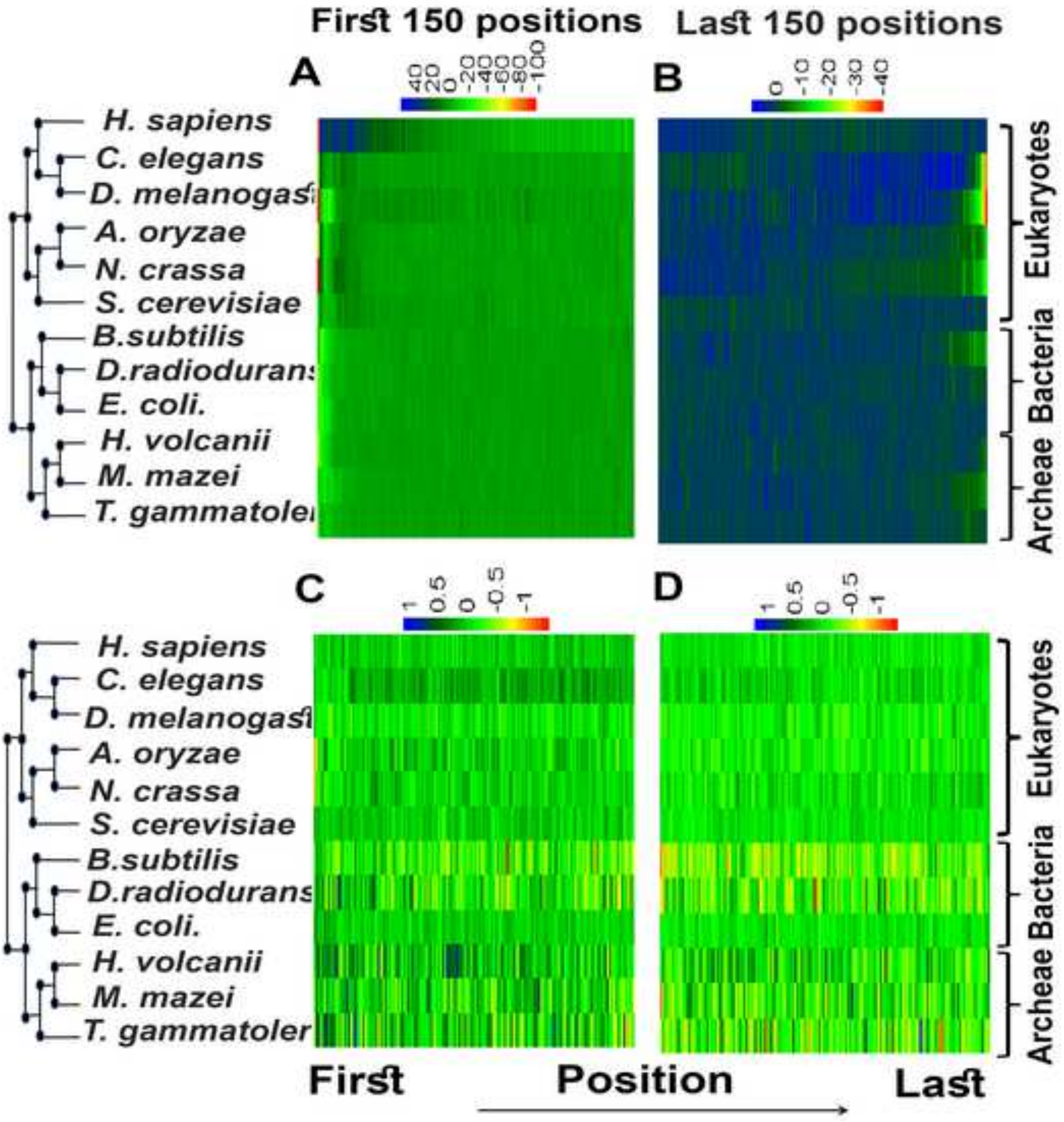

First 150 positions
Last 150 positions
A
40
20
0
-20
-40
-60
-80
-100
B
0
-10
-20
-30
-40
C
1
0.5
0
-0.5
-1
D
1
0.5
0
-0.5
-1
H. sapiens
C. elegans
D. melanogast
A. oryzae
N. crassa
S. cerevisiae
B.subtilis
D.radiodurans
E. coli.
H. volcanii
M. mazei
T. gammatoler
Eukaryotes
Bacteria
Archeae
First
Position
Last

**Figure S37**

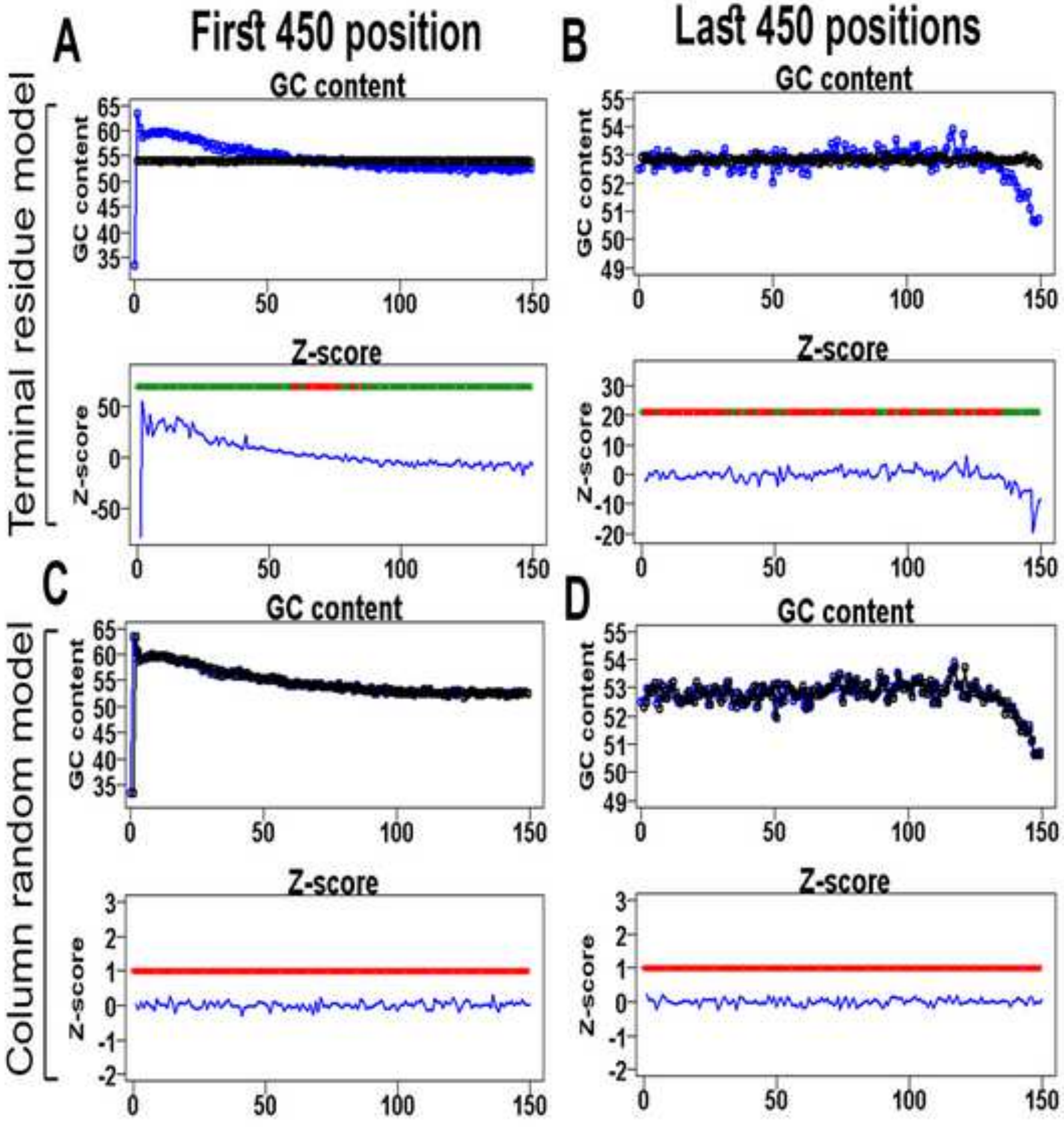
A
First 450 position
B
Last 450 positions
Terminal residue model
GC content
GC content
Z-score
Z-score
C
D
Column random model
GC content
GC content
Z-score
Z-score

**Figure S38**

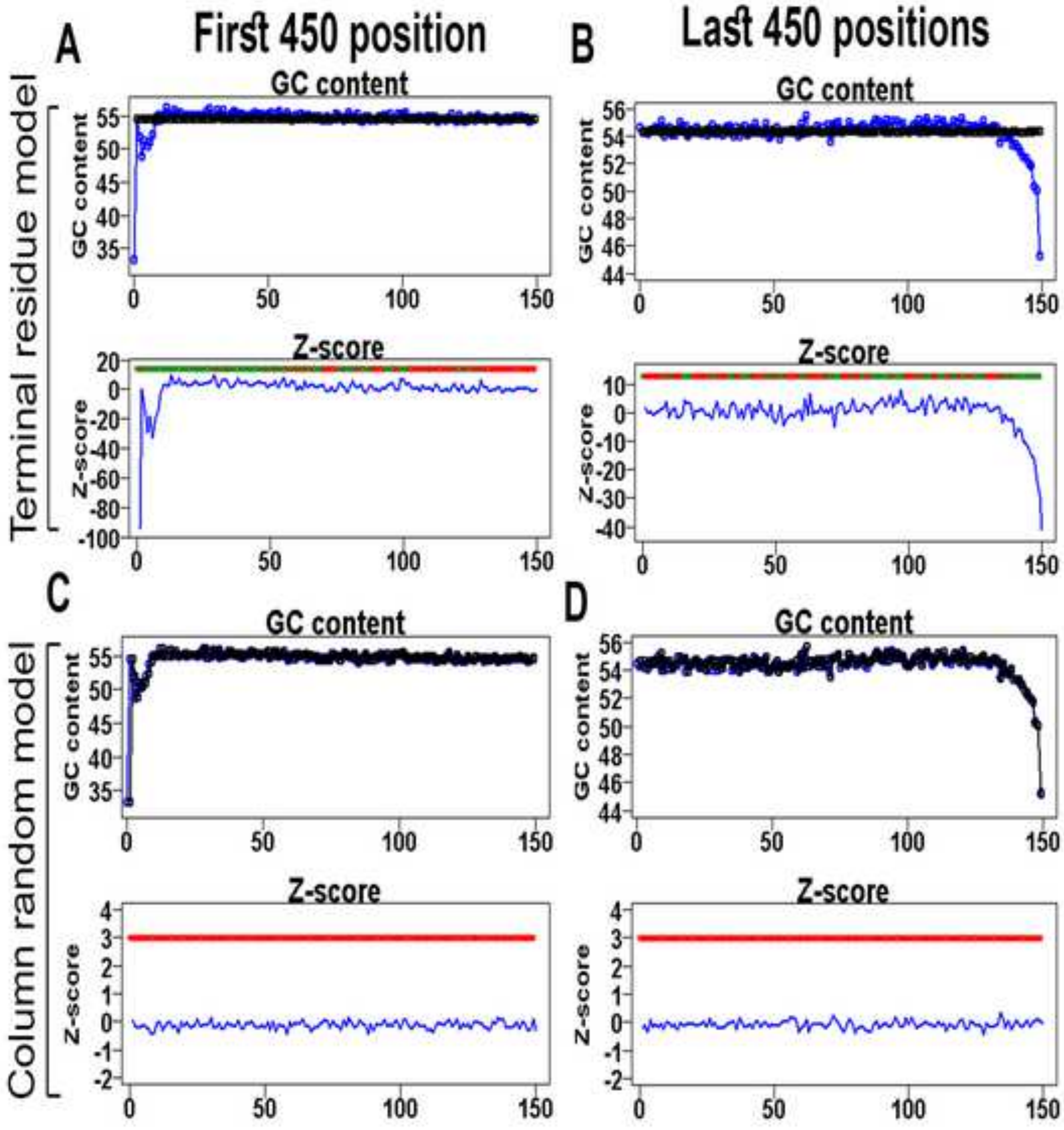
A
First 450 position
B
Last 450 positions
Terminal residue model
Column random model
C
D
GC content
Z-score

**Figure S39**

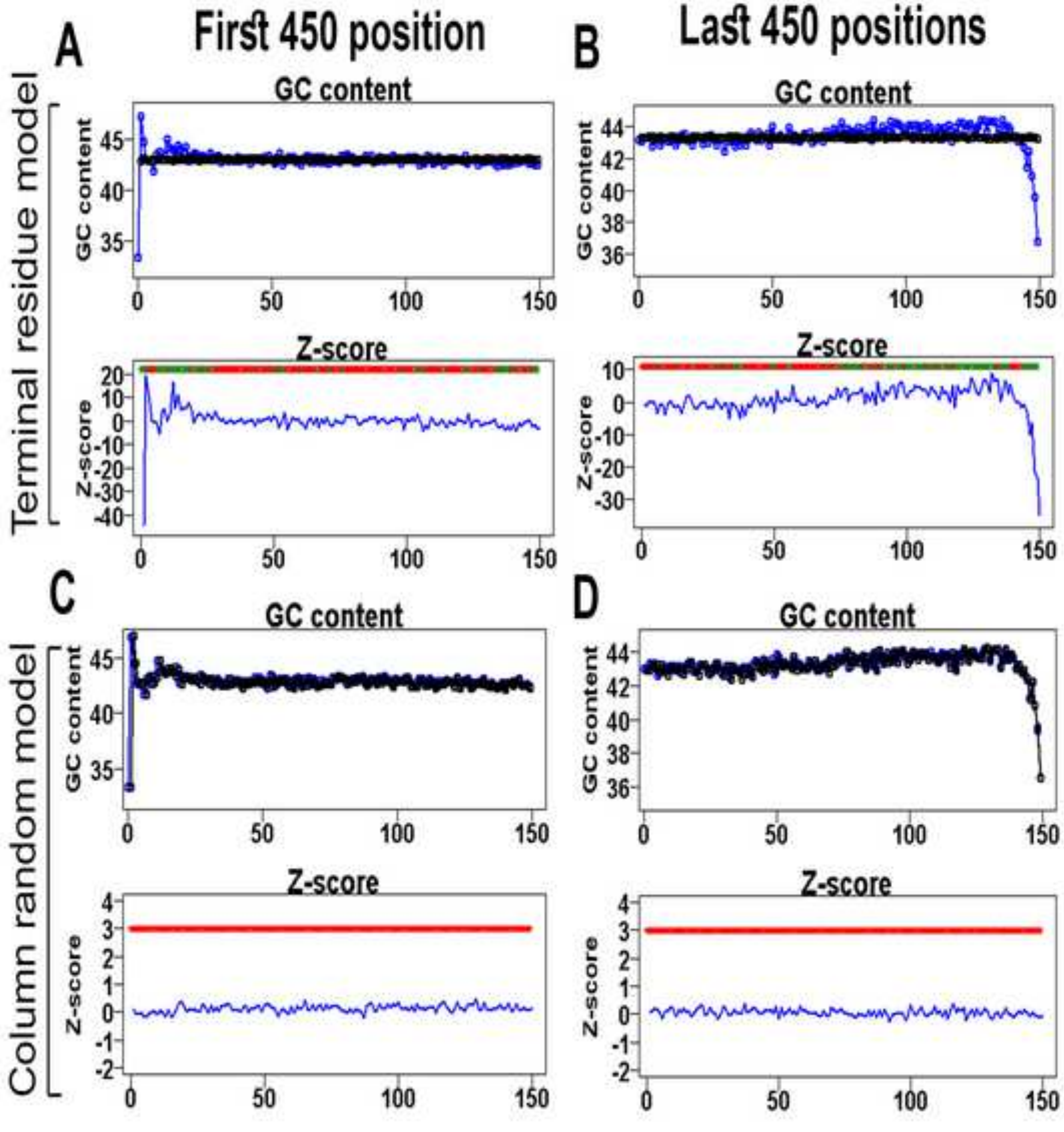

A
First 450 position
B
Last 450 positions
Terminal residue model
GC content
GC content
Z-score
Z-score
C
D
Column random model
GC content
GC content
Z-score
Z-score

Figure S40

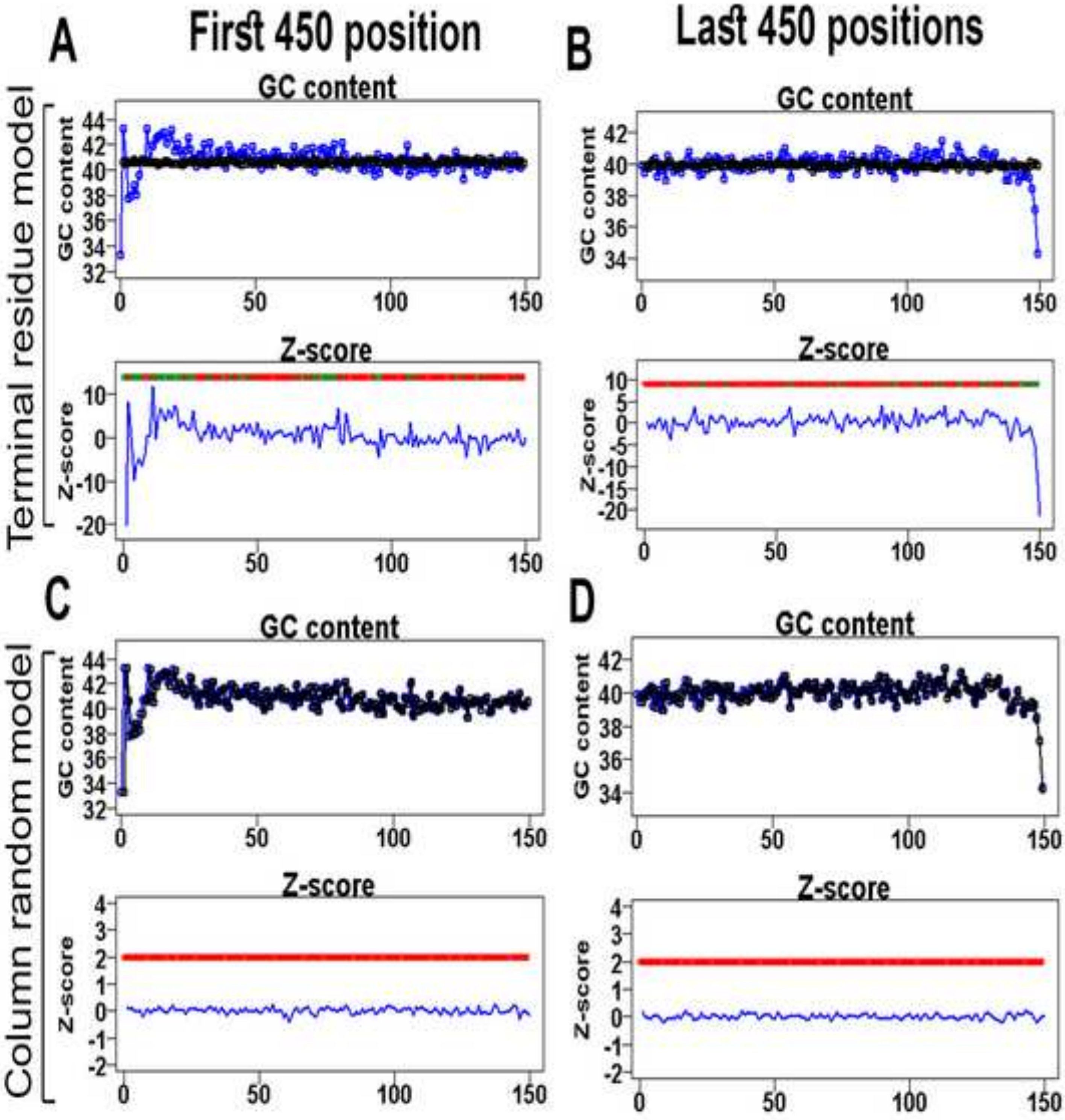

A
First 450 position
B
Last 450 positions
Terminal residue model
GC content
GC content
Z-score
C
D
Column random model

**Figure S41**

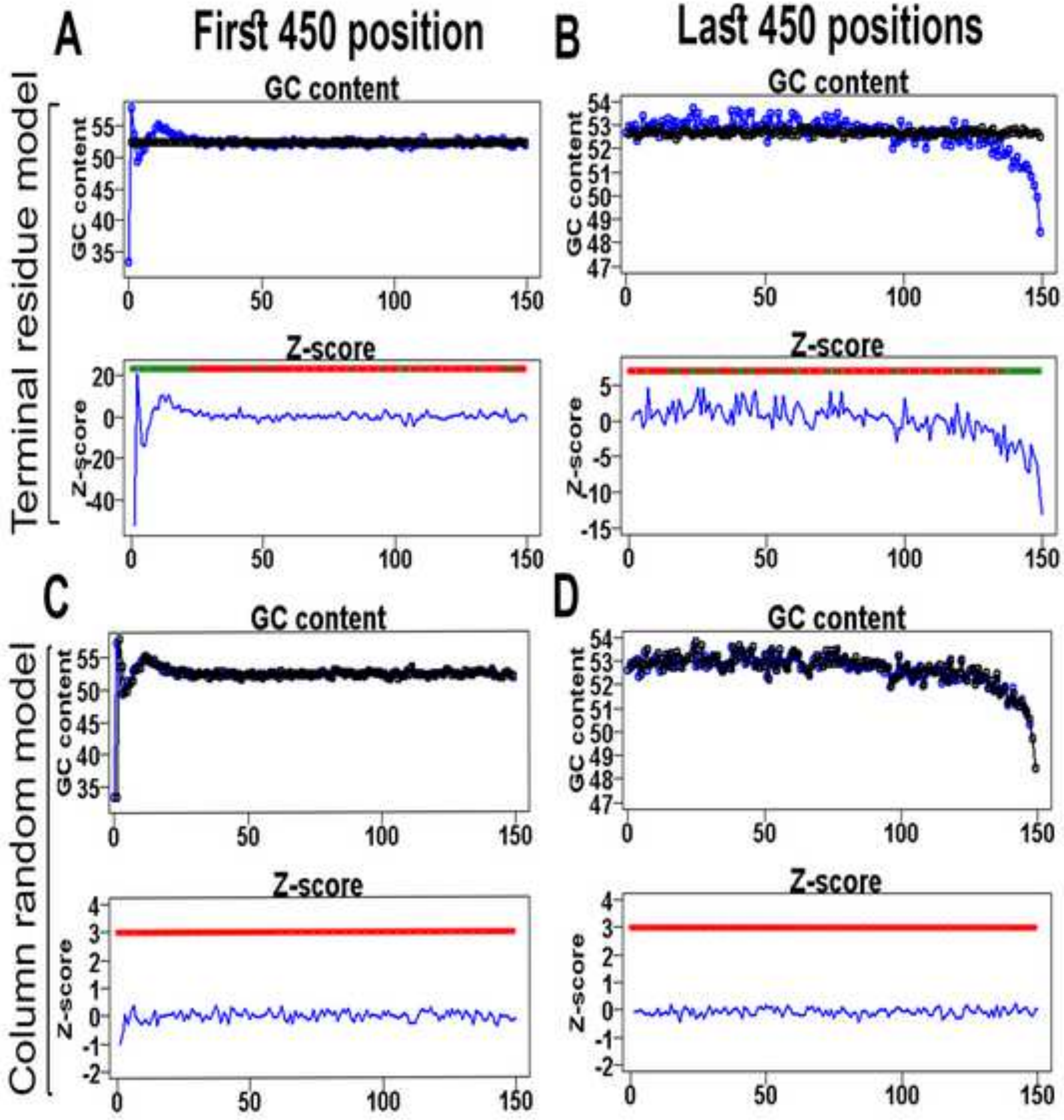

A
First 450 position
B
Last 450 positions
Terminal residue model
GC content
GC content
Z-score
Z-score
C
D
Column random model
GC content
GC content
Z-score
Z-score

Figure S42

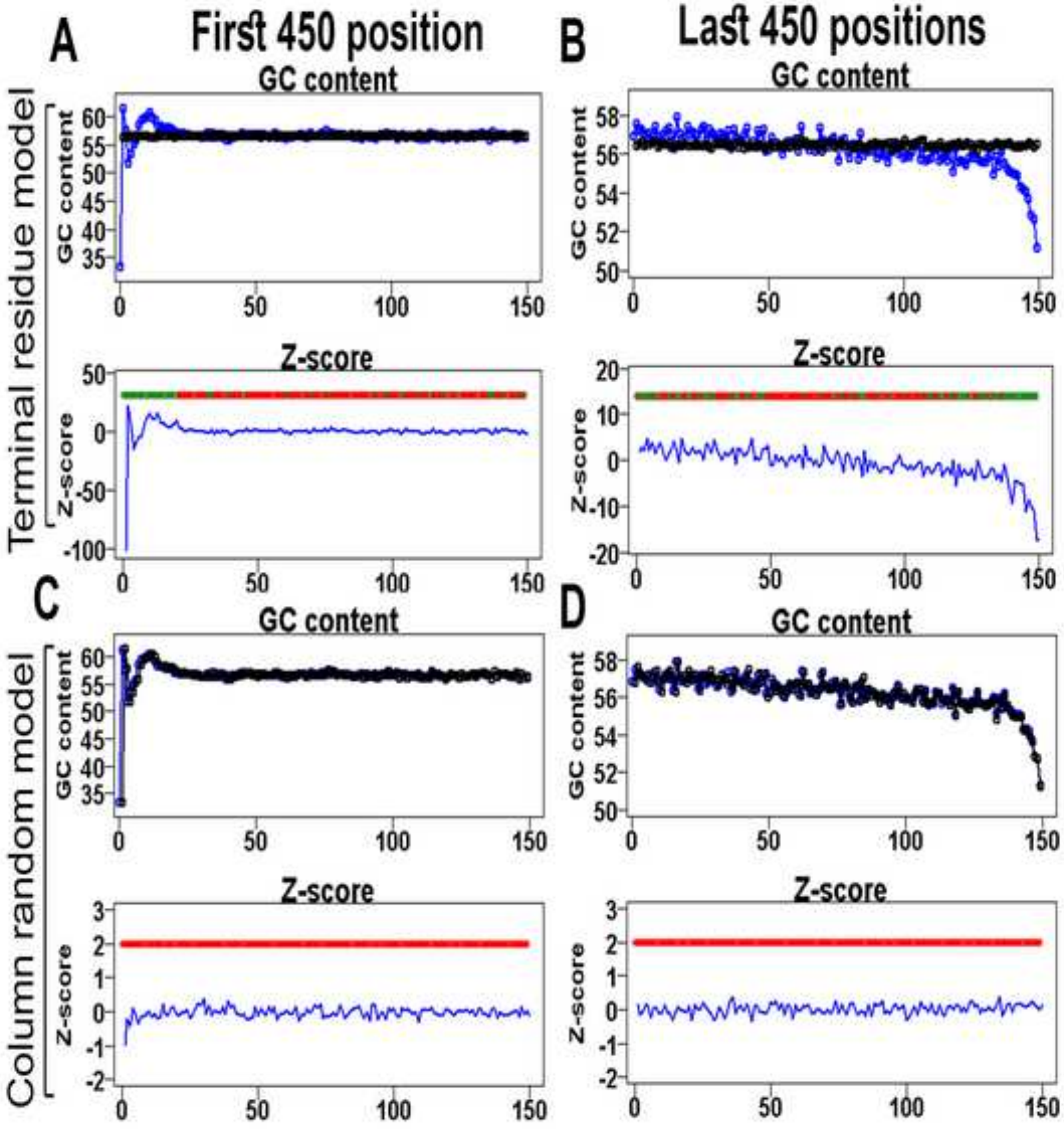
A
First 450 position
B
Last 450 positions
Terminal residue model
GC content
GC content
Z-score
Z-score
C
D
Column random model
GC content
GC content
Z-score
Z-score
0
50
100
150

**Figure S43**

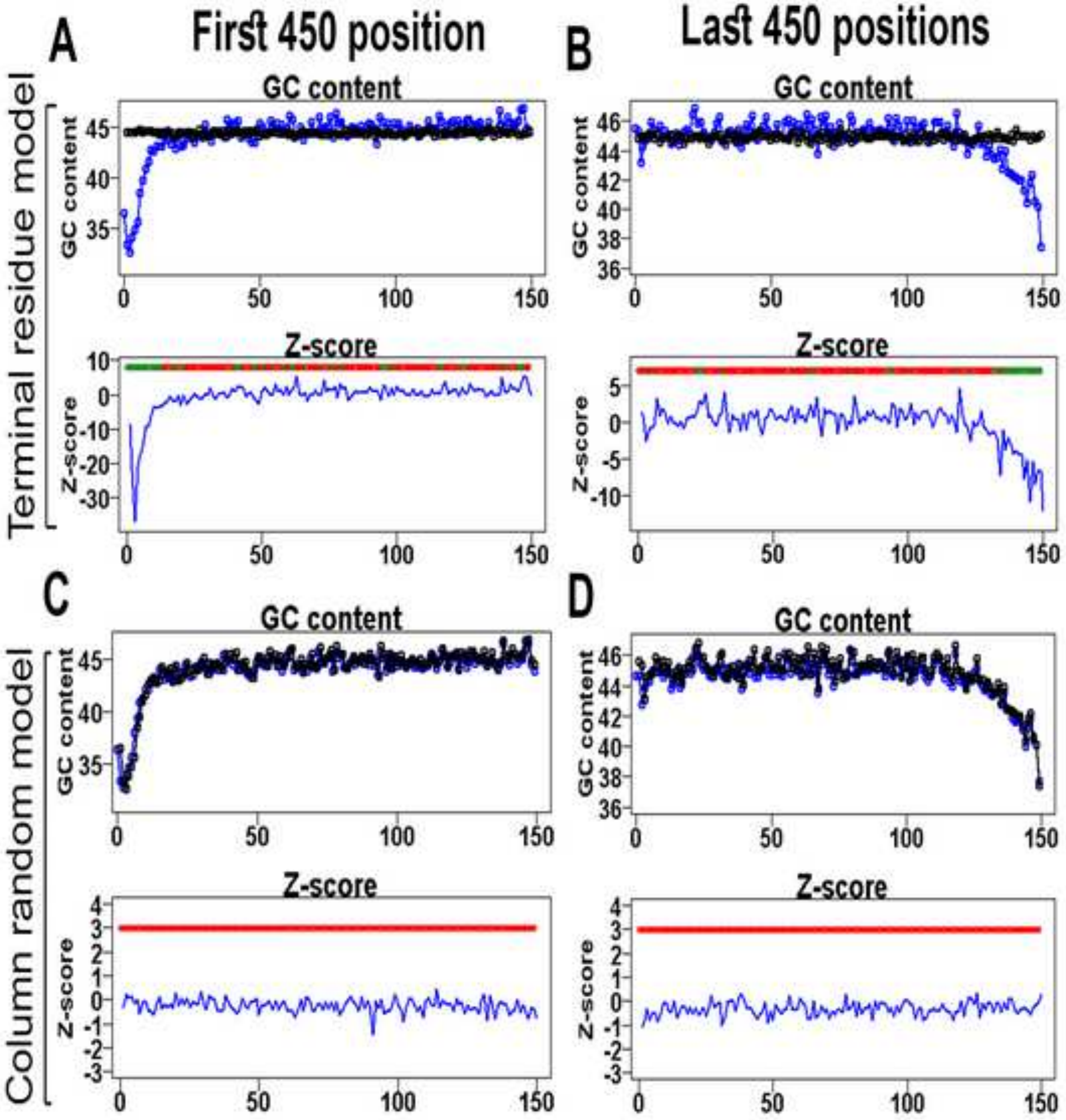

A
First 450 position
B
Last 450 positions
Terminal residue model
GC content
Z-score
C
D
Column random model

**Figure S44**

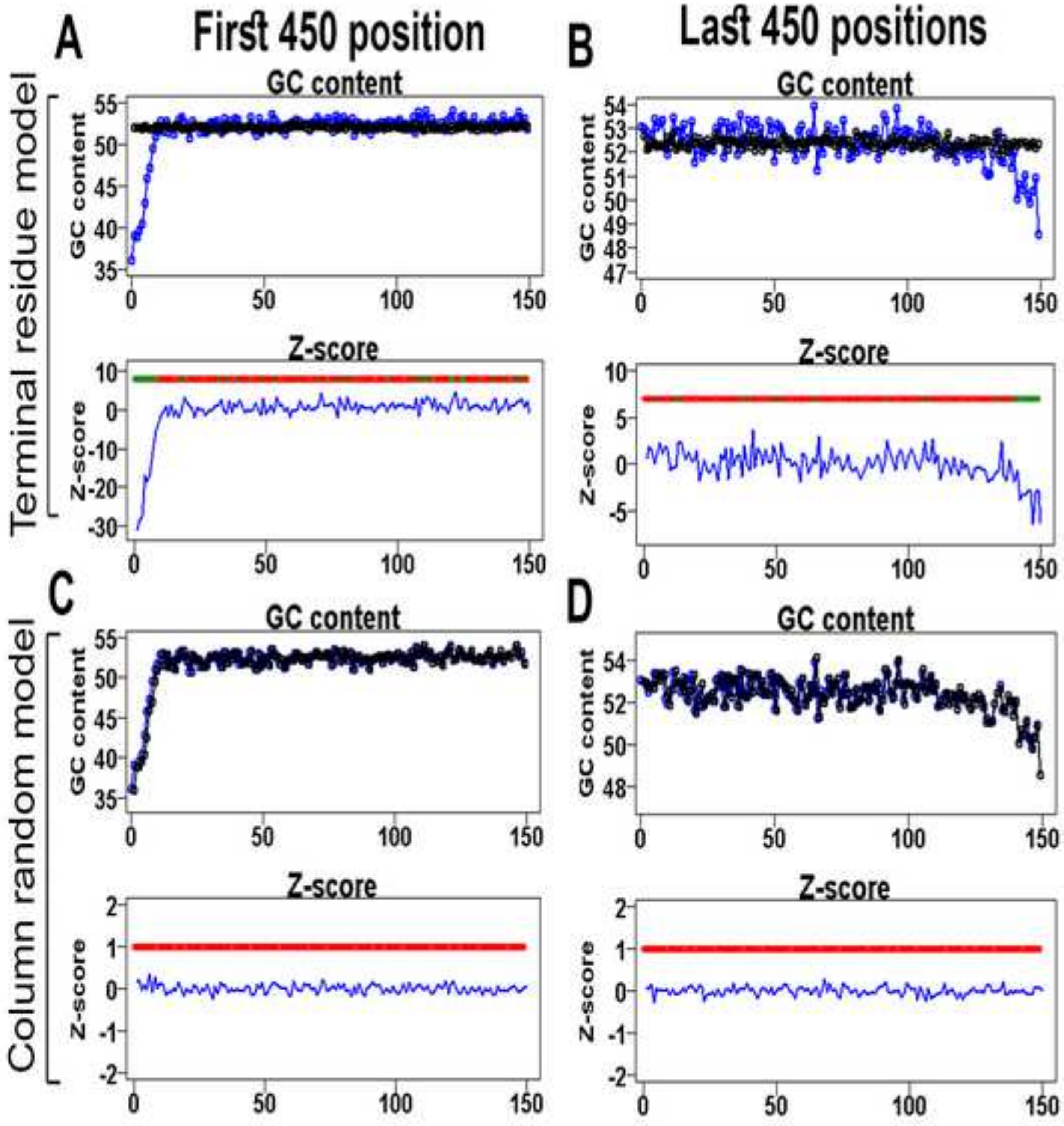

A
First 450 position
B
Last 450 positions
Terminal residue model
GC content
GC content
Z-score
Z-score
C
D
Column random model
GC content
GC content
Z-score
Z-score

Figure S45

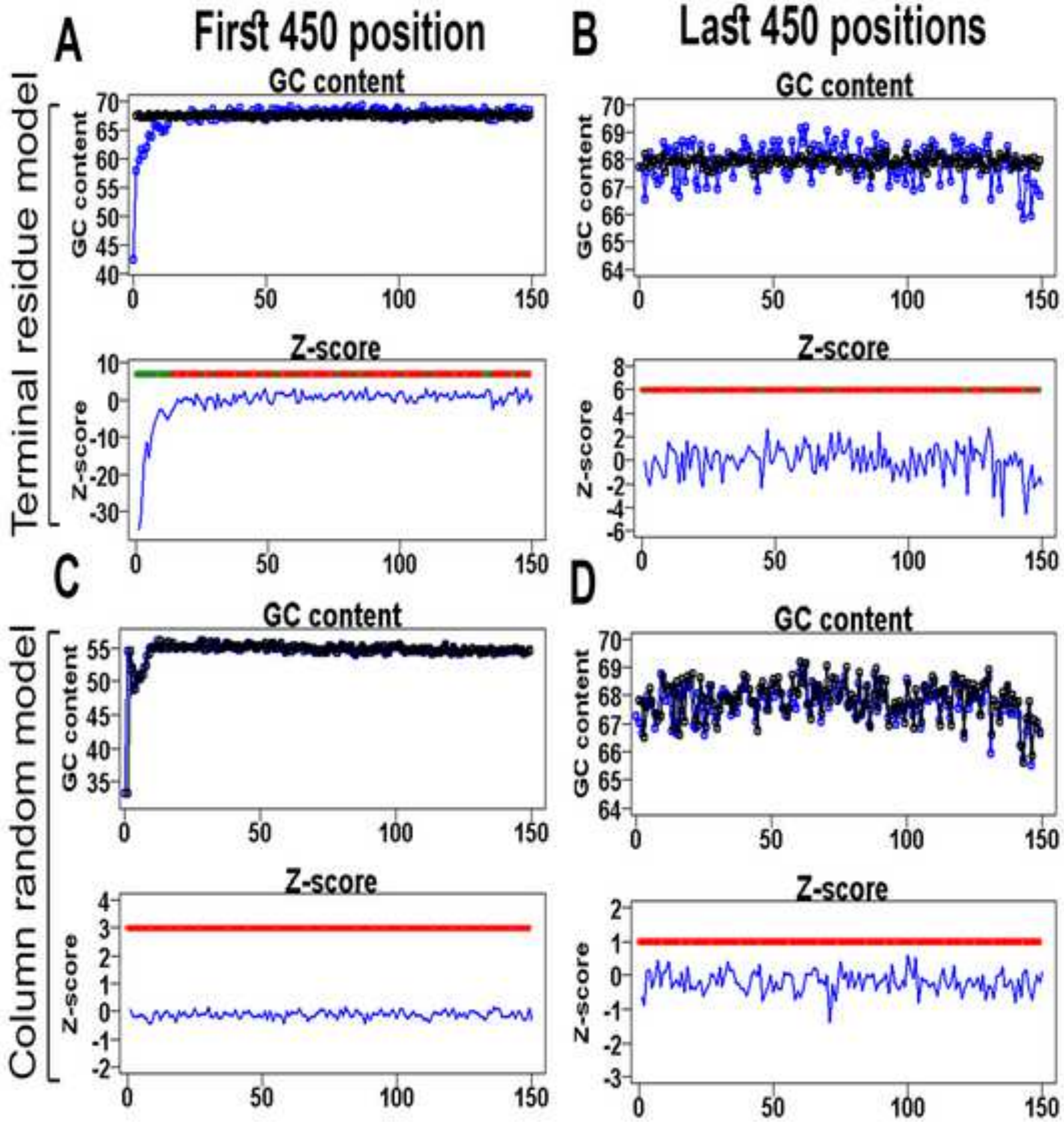

A
First 450 position
B
Last 450 positions
Terminal residue model
GC content
GC content
Z-score
Z-score
C
D
Column random model
GC content
GC content
Z-score
Z-score

Figure S46

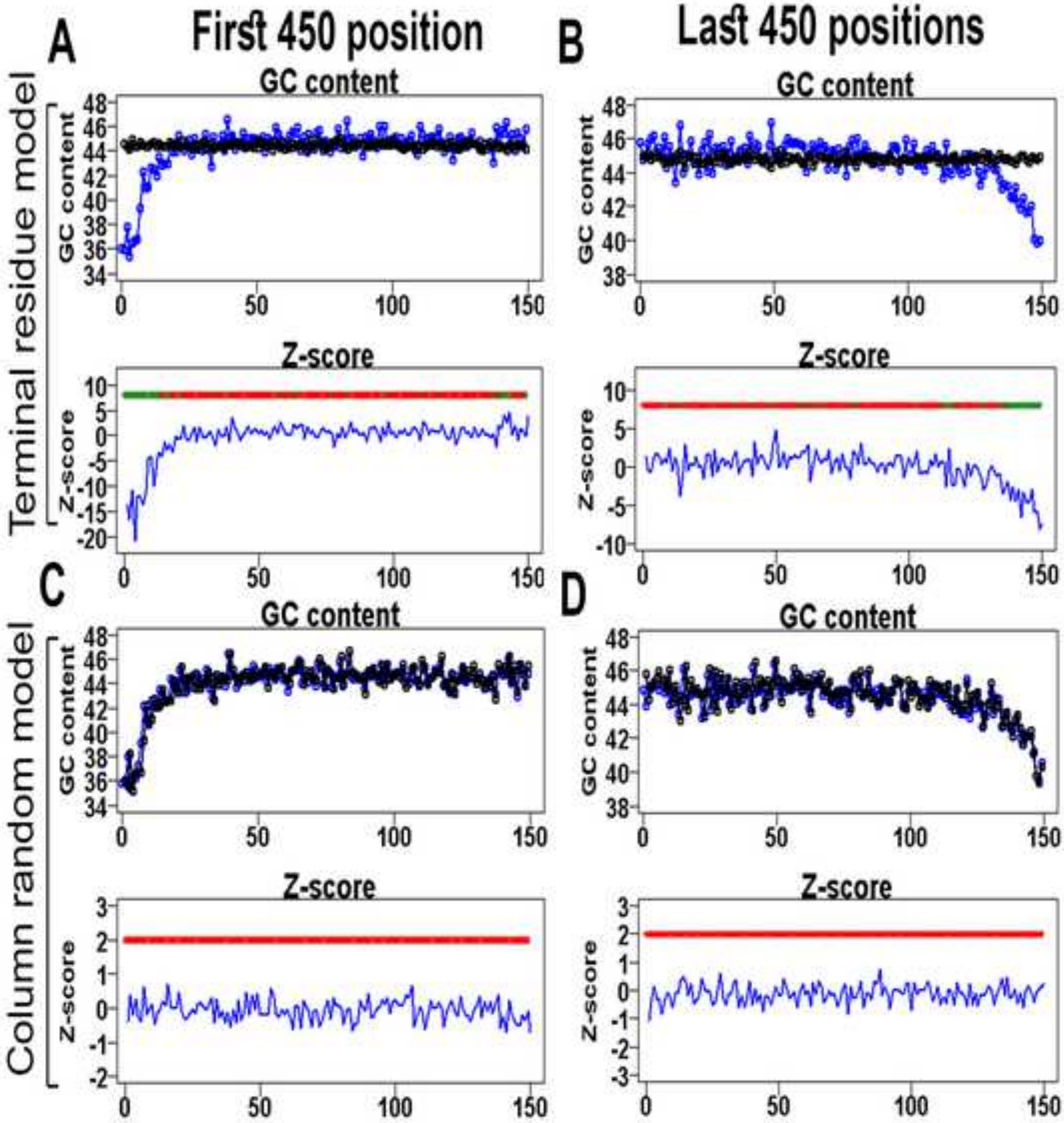
A
First 450 position
B
Last 450 positions
C
D
Terminal residue model
Column random model
GC content
Z-score
GC content
Z-score
GC content
Z-score
GC content
Z-score
GC content
Z-score

Figure S47

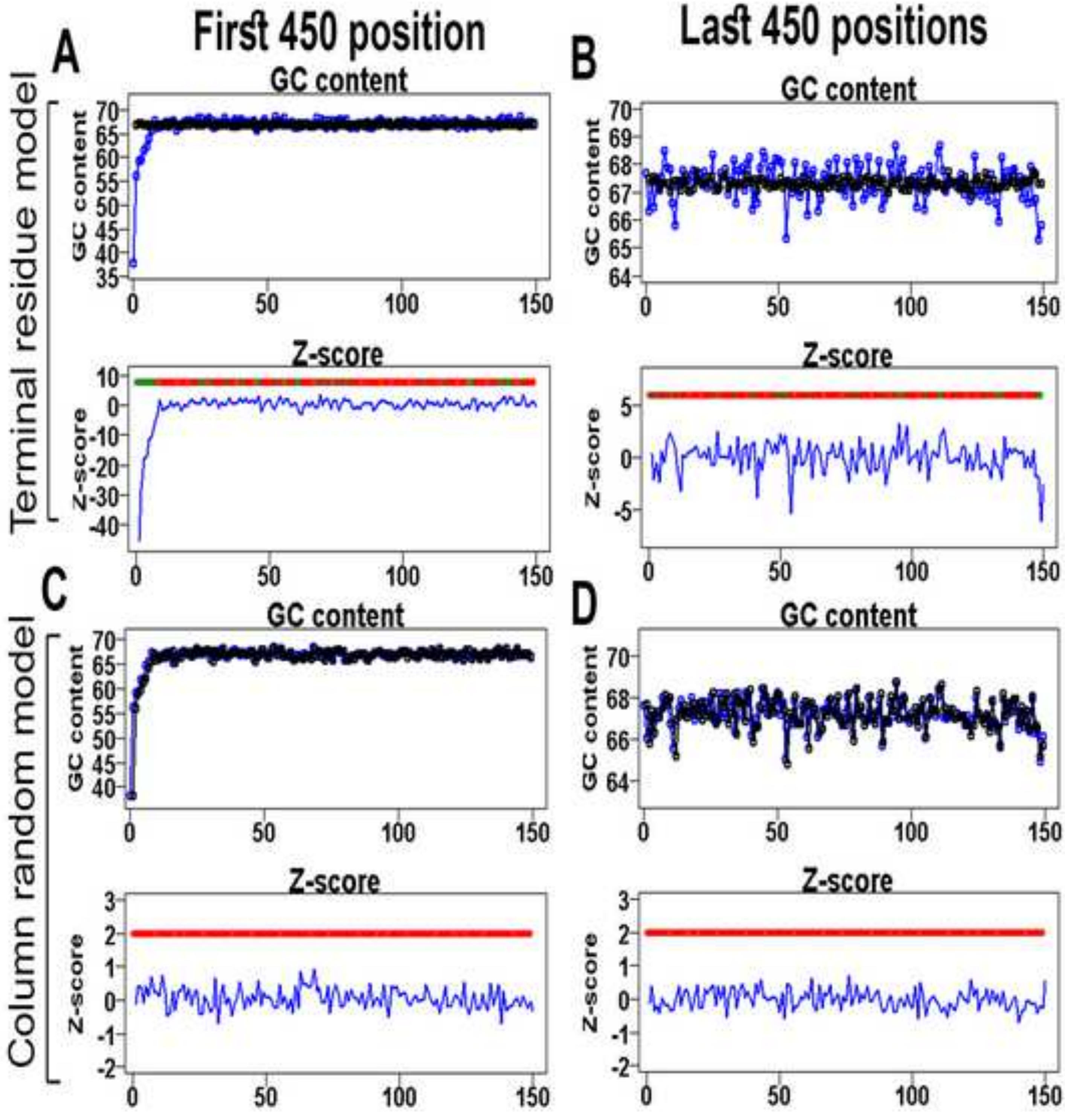

Terminal residue model
Column random model
A
First 450 position
B
Last 450 positions
C
D
GC content
Z-score

Figure S48

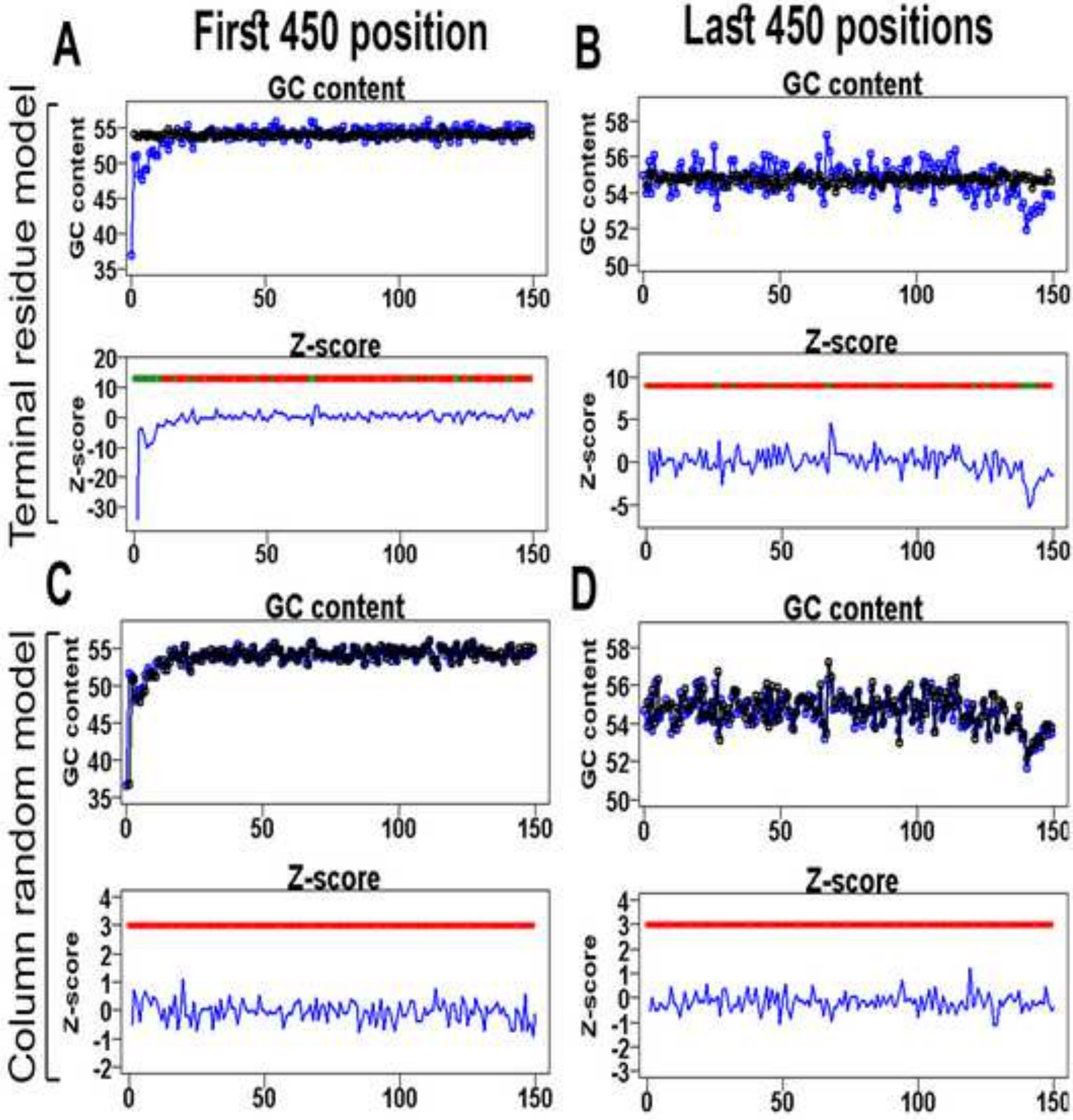
A
First 450 position
B
Last 450 positions
GC content
Z-score
GC content
Z-score
Terminal residue model
C
D
Column random model
GC content
Z-score

**Figure S49**

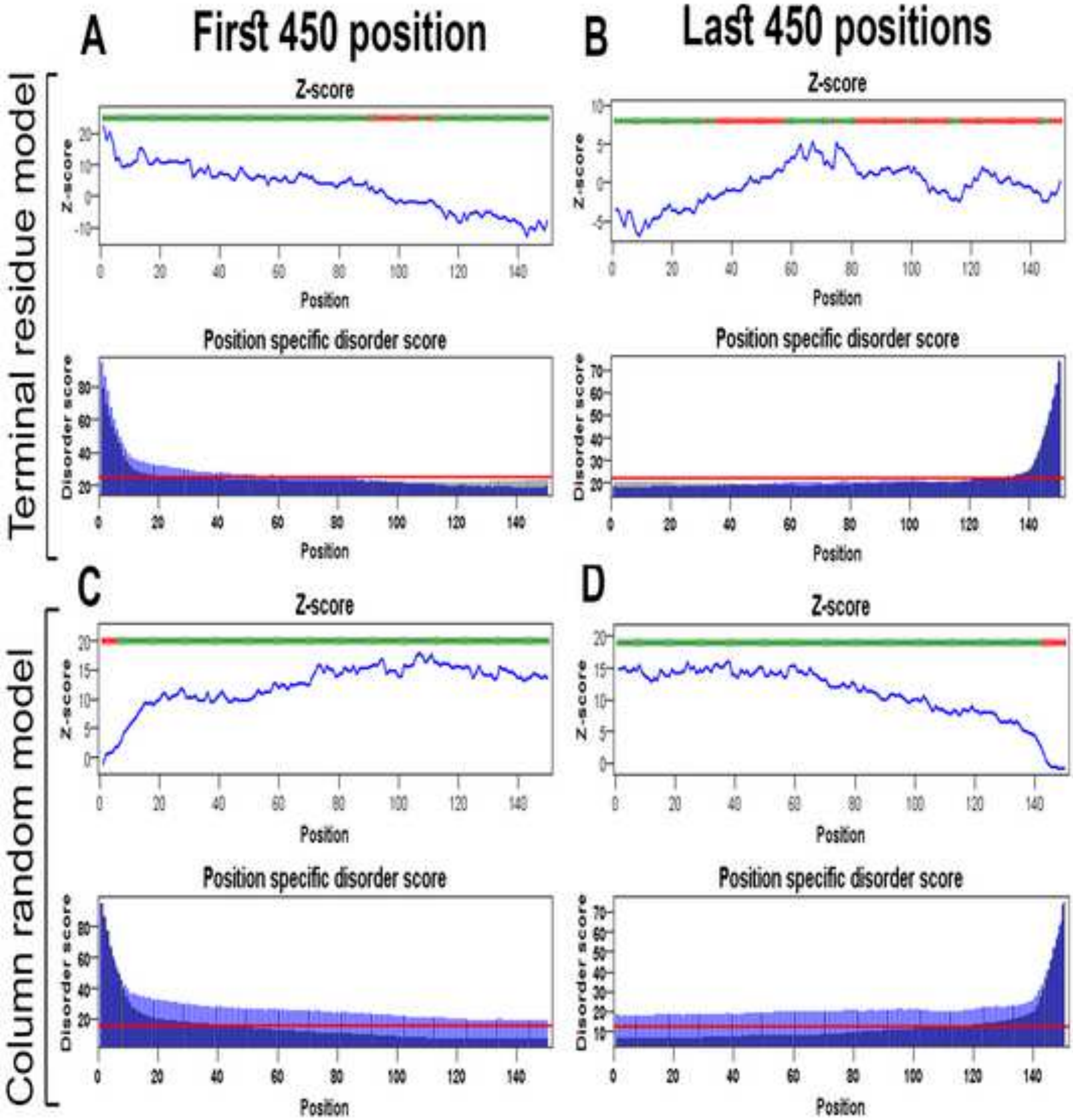

A
First 450 position
B
Last 450 positions
Terminal residue model
Column random model
C
D
Z-score
Position specific disorder score
Z-score
Disorder score
Position

Figure S50

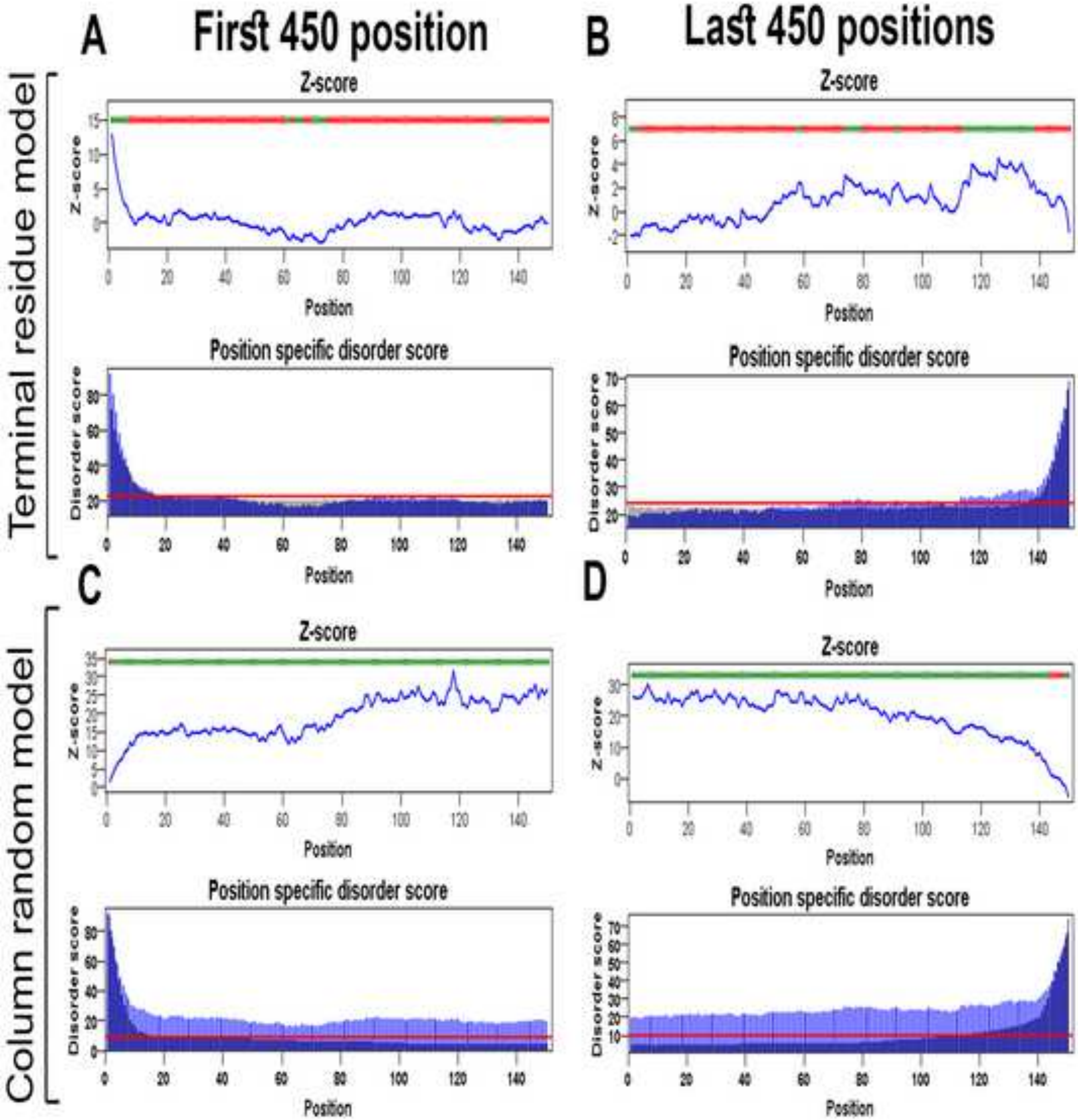
A
First 450 position
B
Last 450 positions
Terminal residue model
Column random model
C
D
Z-score
Position
Position specific disorder score
Disorder score

**Figure S51**

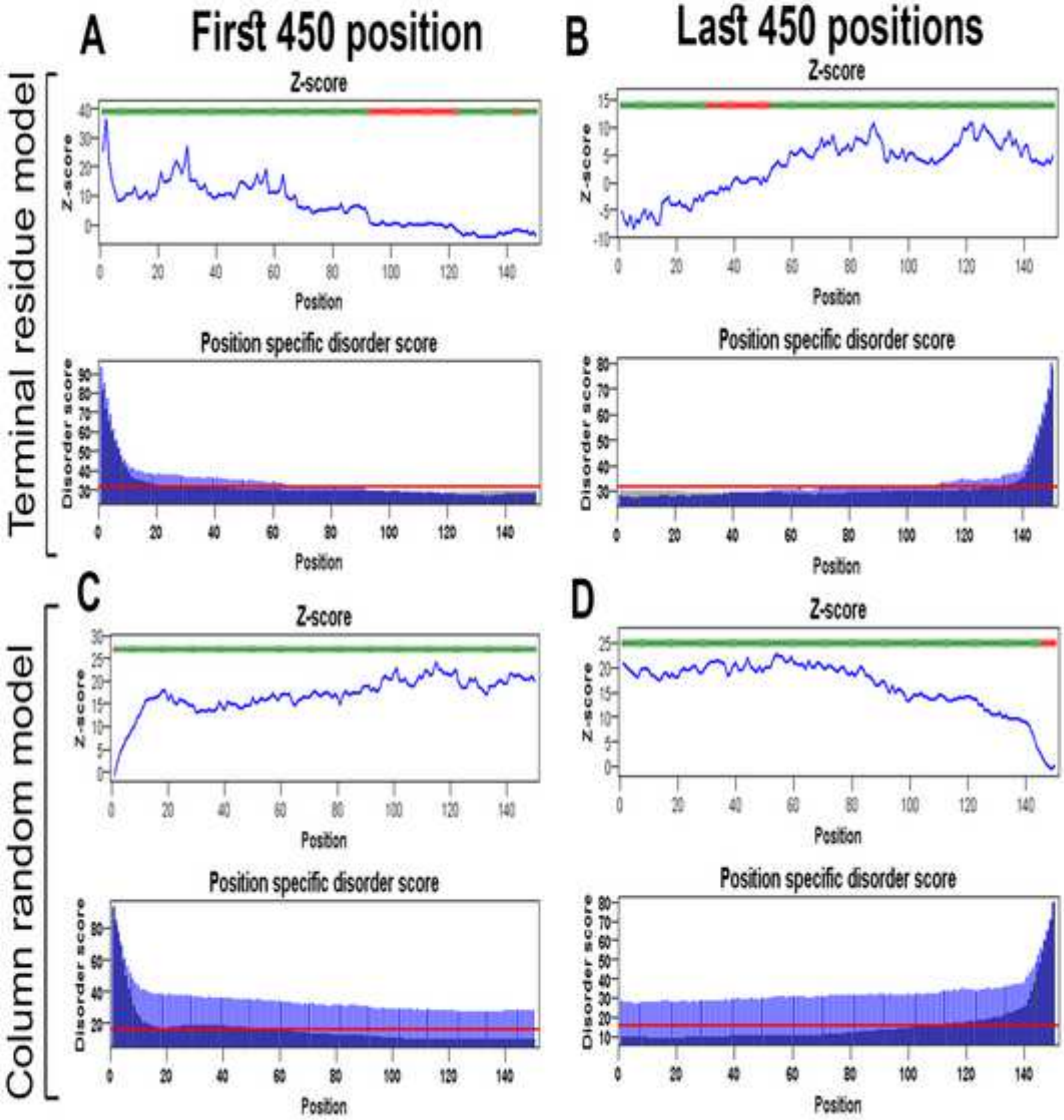
A
First 450 position
B
Last 450 positions
C
D
Terminal residue model
Column random model
Z-score
Position specific disorder score
Position
Disorder score

**Figure S52**

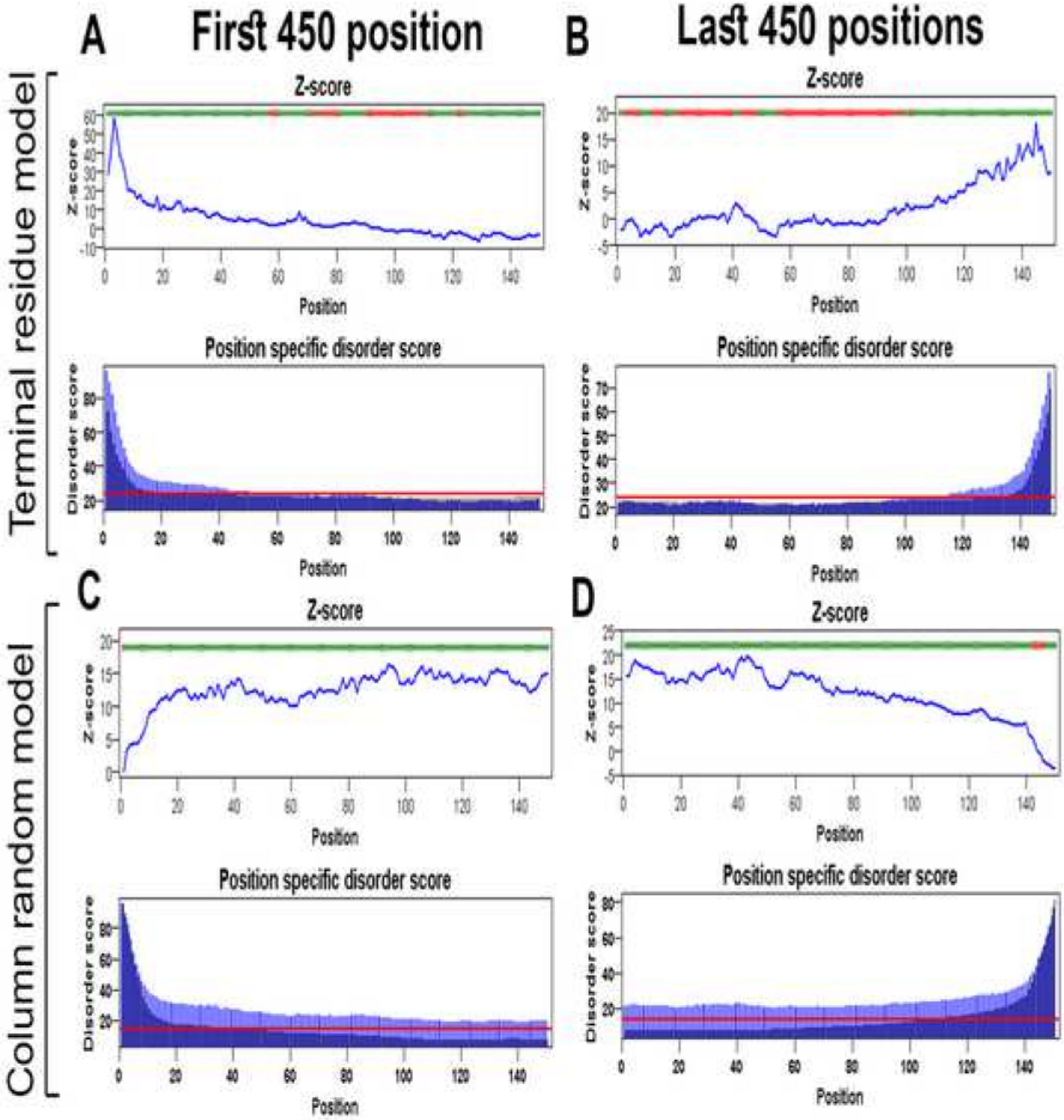
A
First 450 position
B
Last 450 positions
Terminal residue model
Column random model
Z-score
Position specific disorder score
Disorder score
Position
C
D

Figure S53

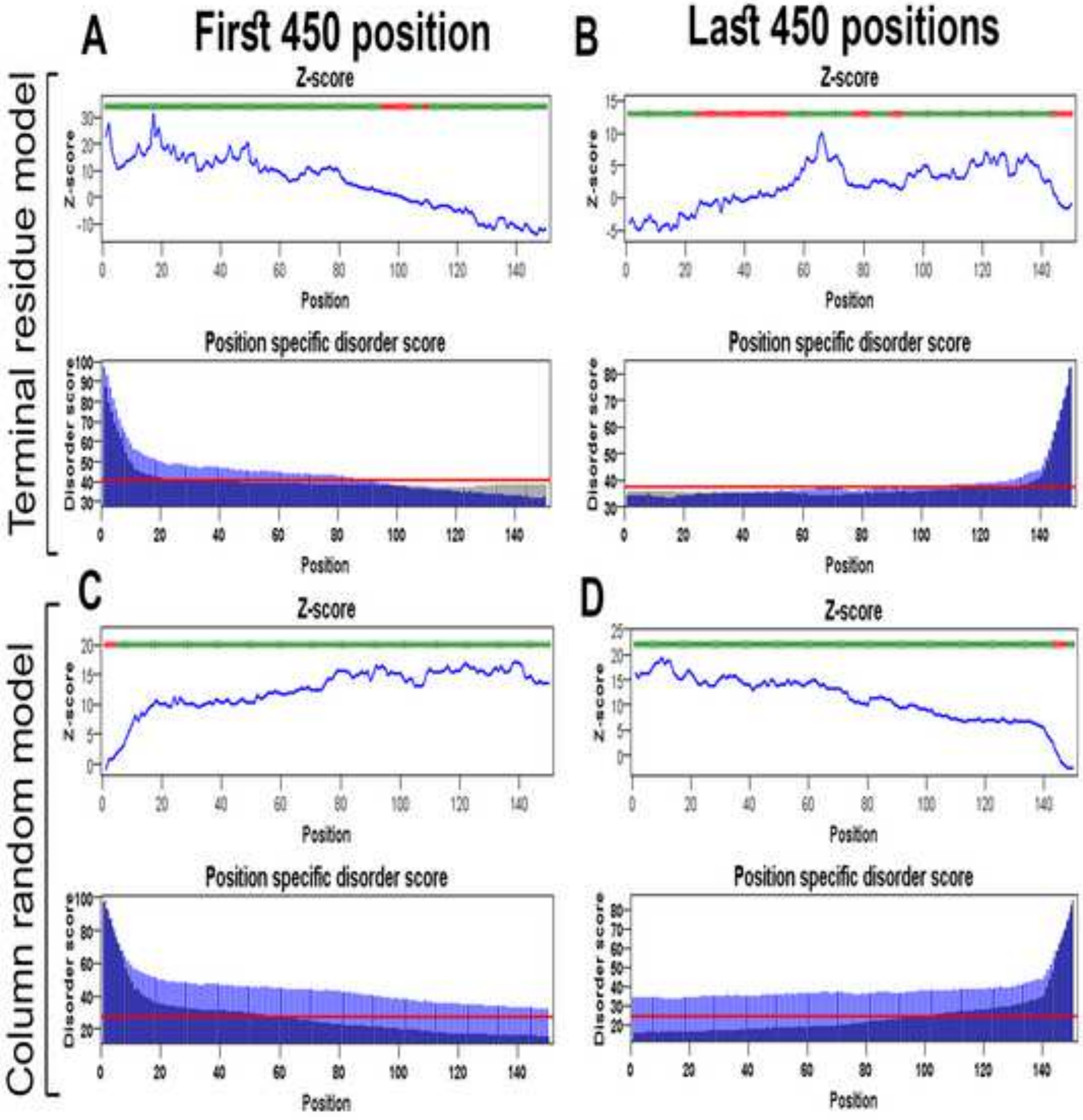
A
First 450 position
B
Last 450 positions
Terminal residue model
Column random model
C
D
Z-score
Position
Position specific disorder score
Disorder score

**Figure S54**

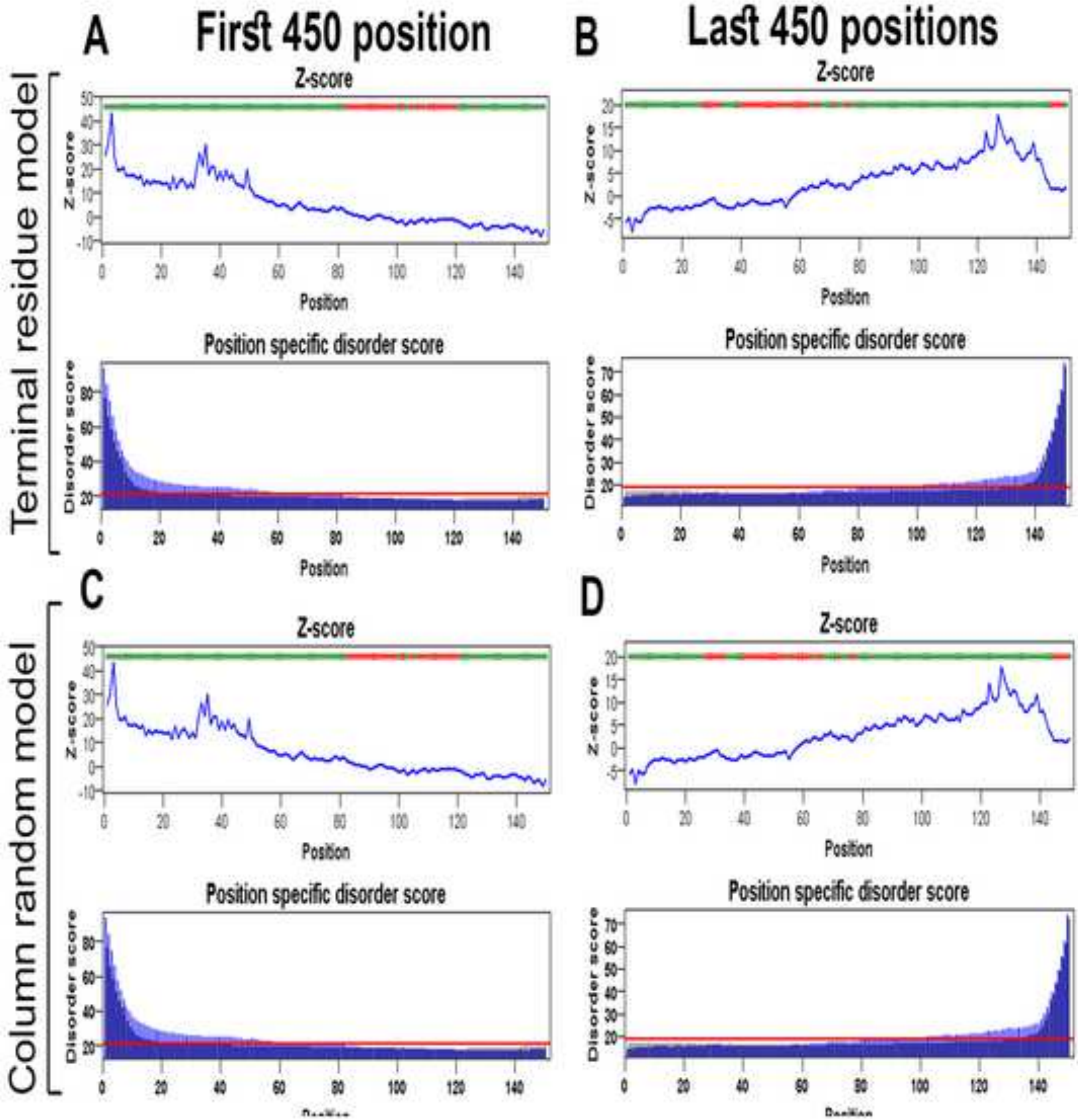

A
First 450 position
B
Last 450 positions
Terminal residue model
Column random model
C
D
Z-score
Position
Position specific disorder score
Disorder score

Figure S55

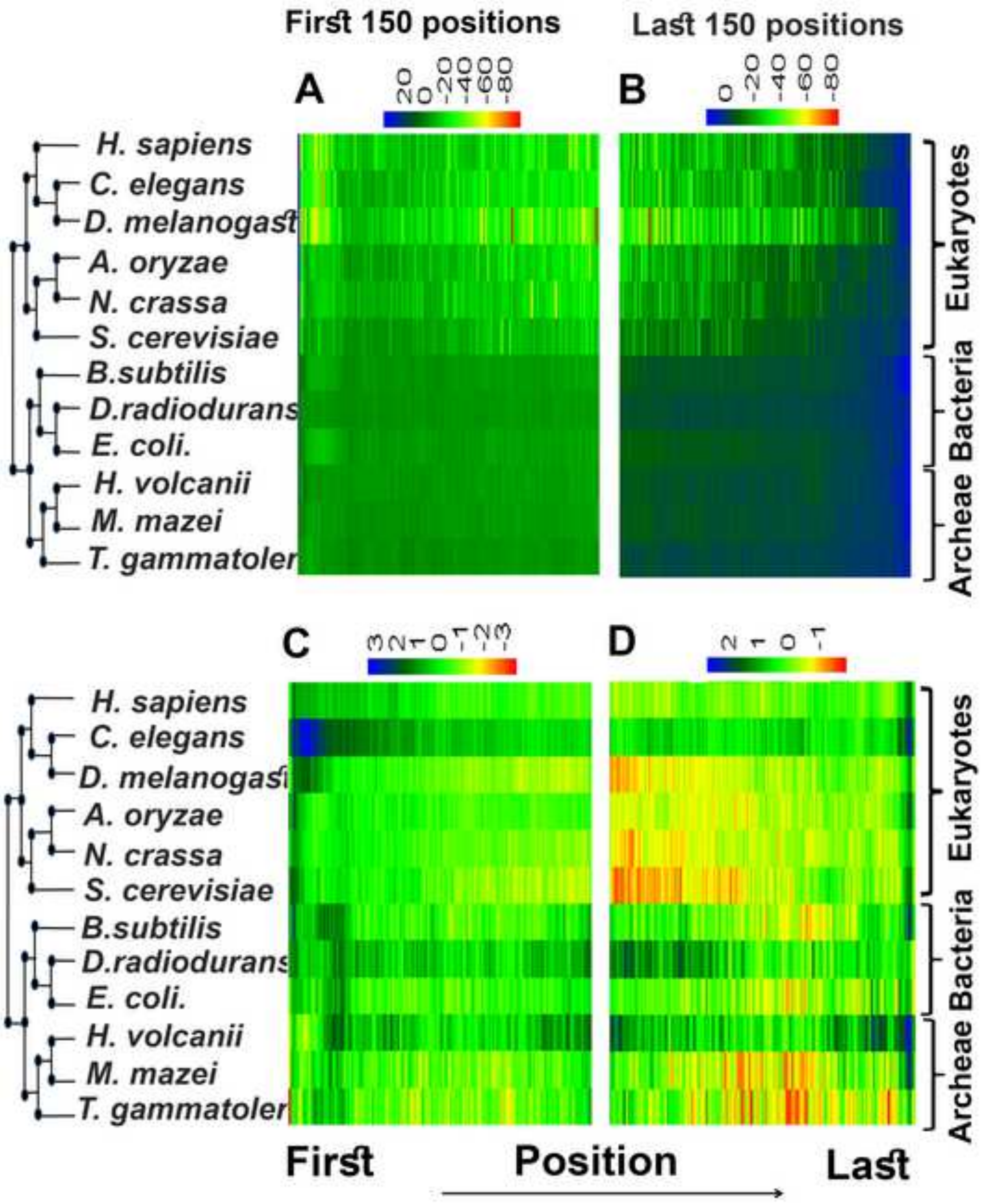

First 150 positions
Last 150 positions
A
20 0 -20 -40 -60 -80
B
0 -20 -40 -60 -80
H. sapiens
C. elegans
D. melanogast
A. oryzae
N. crassa
S. cerevisiae
B.subtilis
D.radiodurans
E. coli.
H. volcanii
M. mazei
T. gammatoler
Eukaryotes
Bacteria
Archeae
C
3 2 1 0 -1 -2 -3
D
2 1 0 -1
First
Position
Last

Figure S56

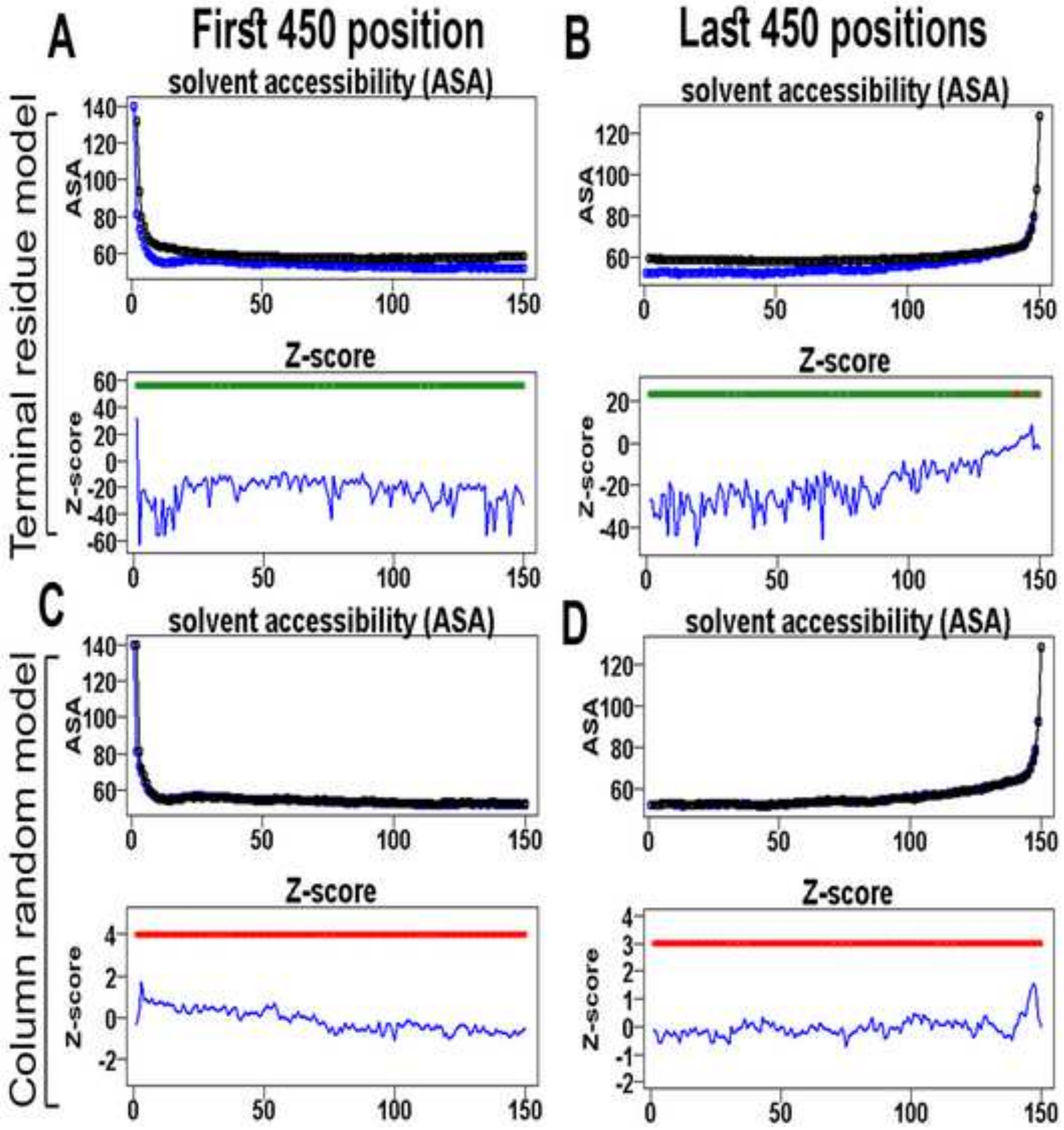

A
First 450 position
B
Last 450 positions
Terminal residue model
Column random model
solvent accessibility (ASA)
ASA
Z-score
C
D

**Figure S57**

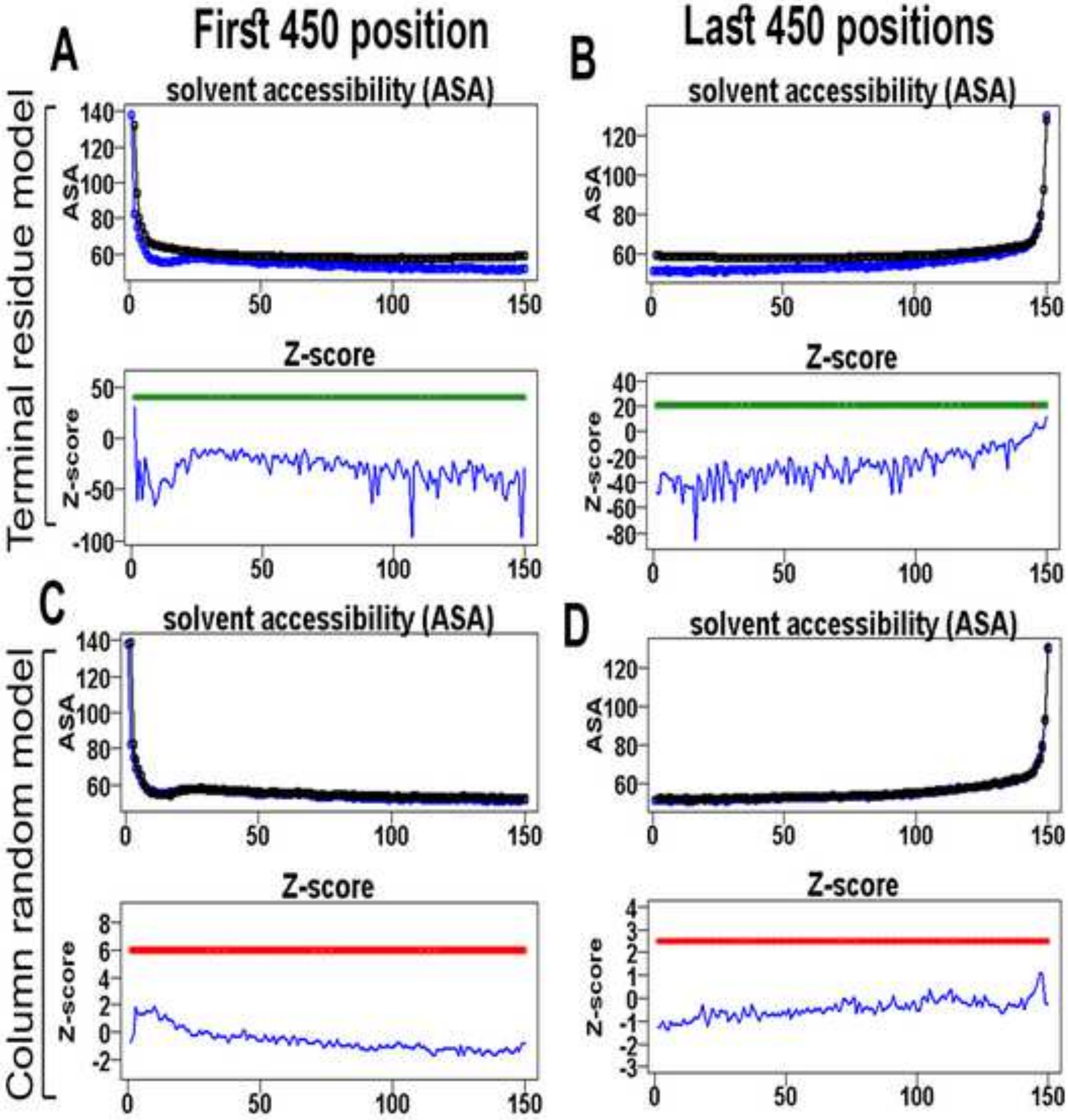

First 450 position
Last 450 positions
A
B
C
D
Terminal residue model
Column random model
solvent accessibility (ASA)
Z-score
ASA
Z-score
140
120
100
80
60
50
0
-50
-100
40
20
-20
-40
-60
-80
8
6
4
2
-2
4
3
1
-1
-3
100
150

**Figure S58**

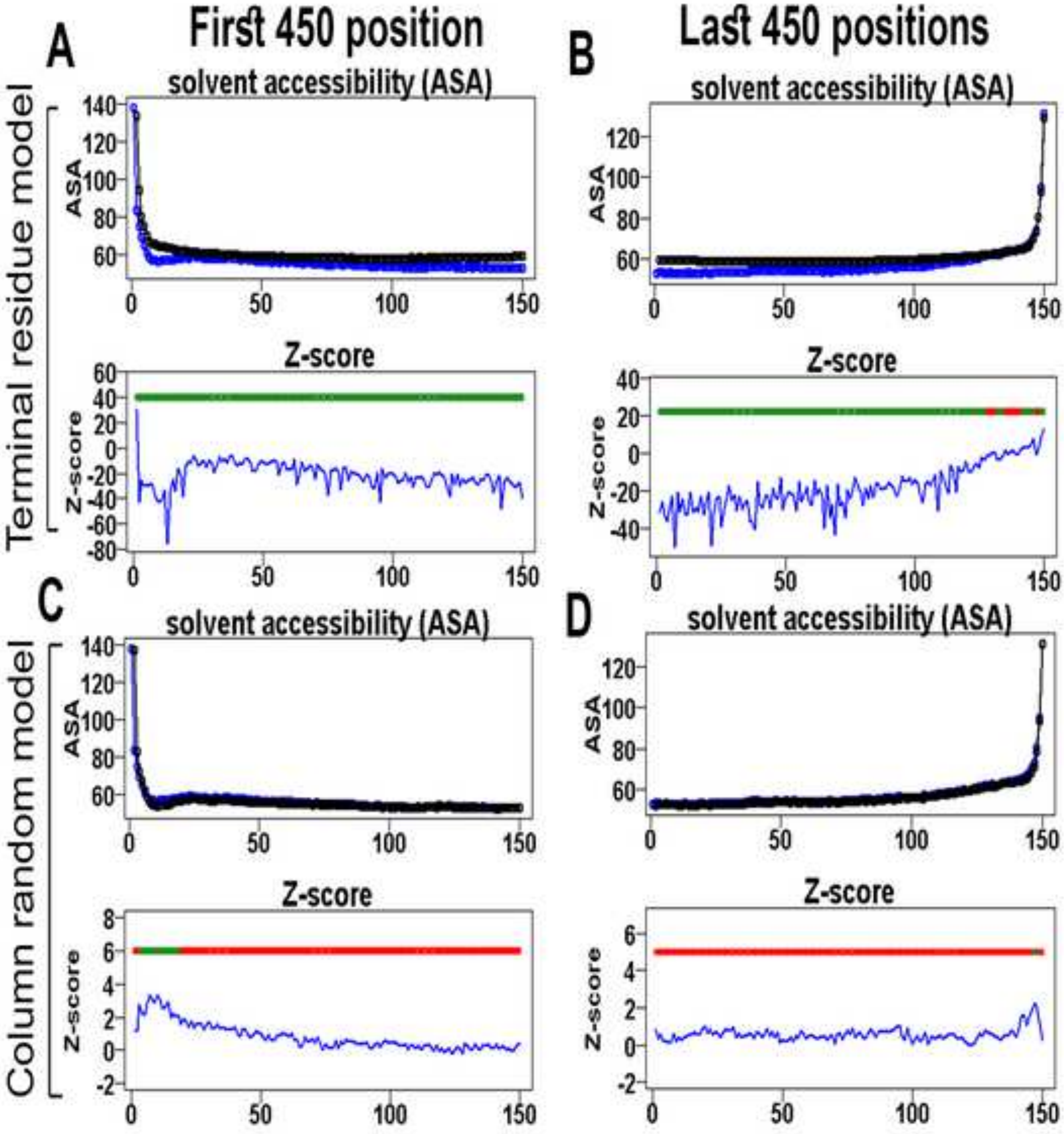
A
First 450 position
B
Last 450 positions
Terminal residue model
solvent accessibility (ASA)
ASA
Z-score
C
D
Column random model

**Figure S59**

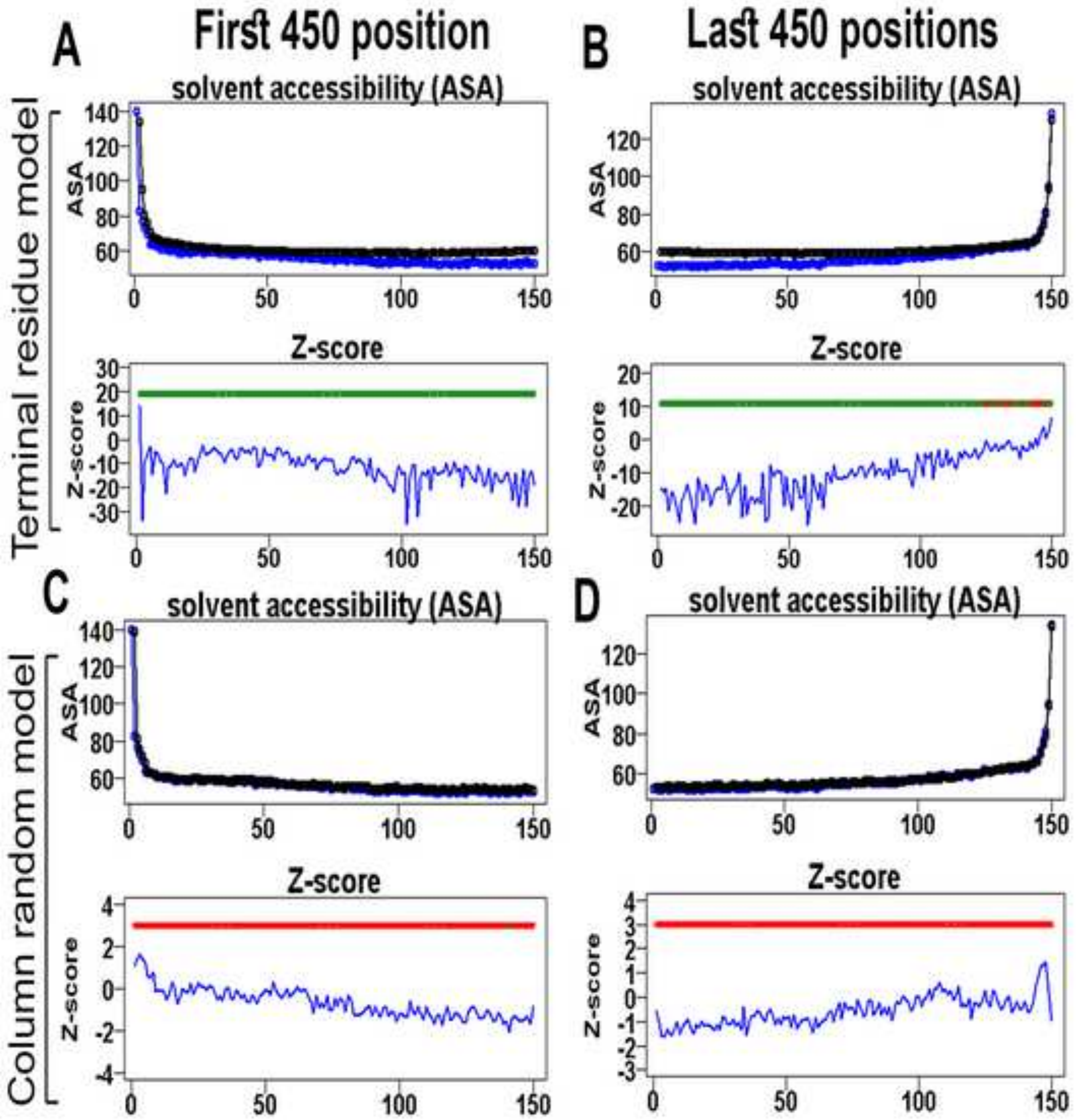
A
First 450 position
B
Last 450 positions
Terminal residue model
Column random model
solvent accessibility (ASA)
ASA
Z-score
C
D

**Figure S60**

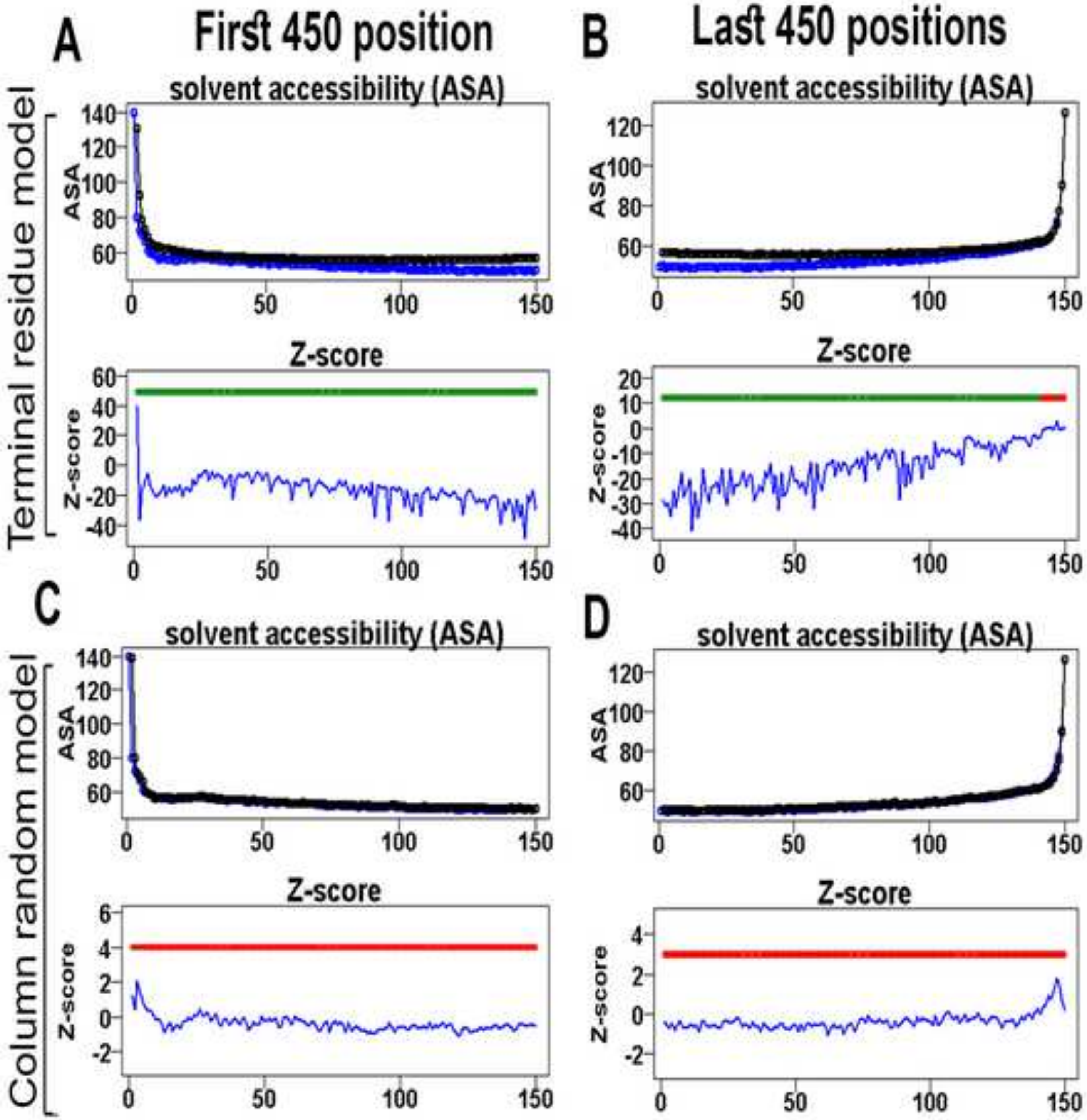

A
First 450 position
B
Last 450 positions
Terminal residue model
solvent accessibility (ASA)
ASA
Z-score
C
D
Column random model

**Figure S61**

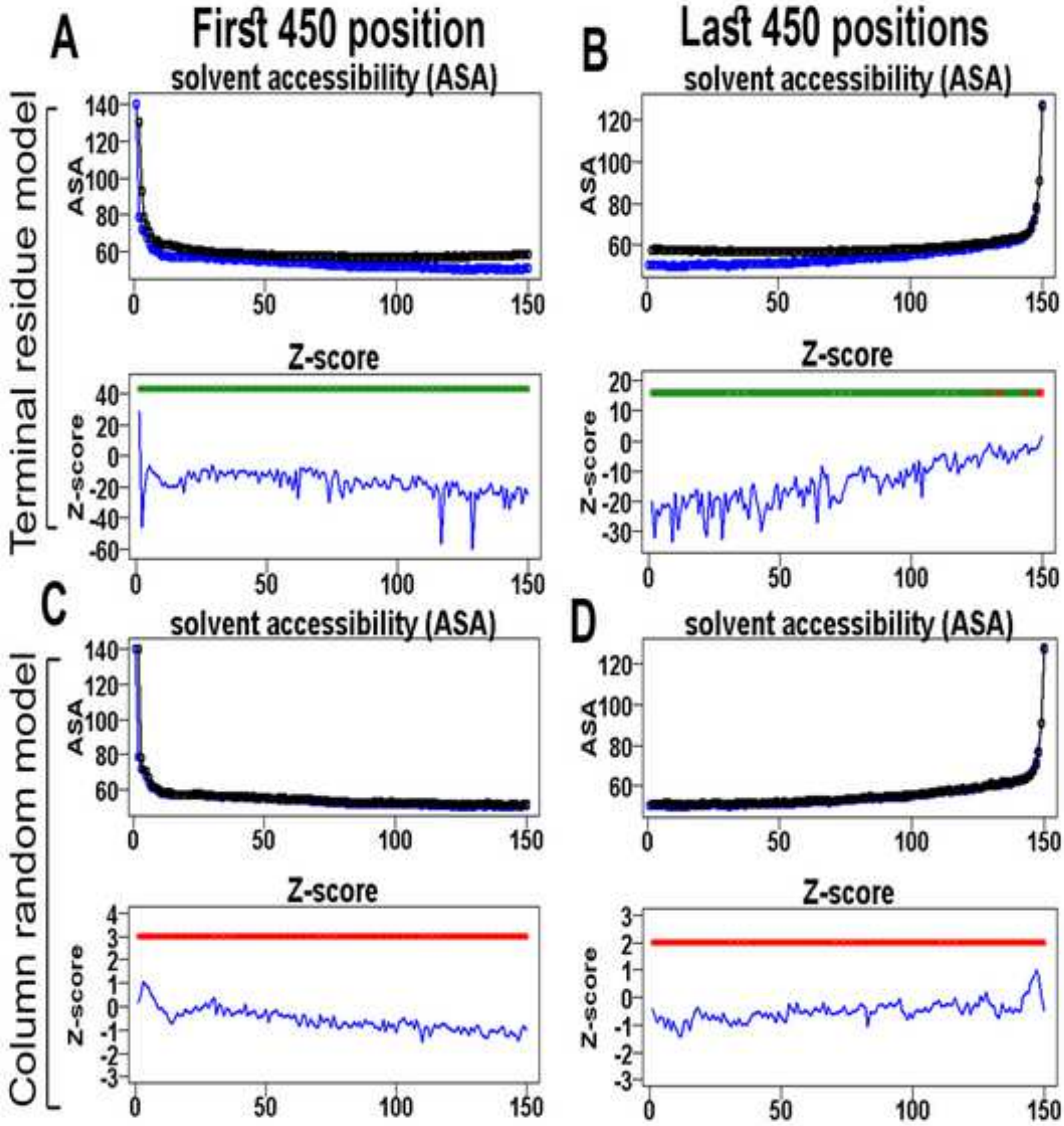
A
First 450 position
B
Last 450 positions
Terminal residue model
solvent accessibility (ASA)
ASA
140
120
100
80
60
0
50
100
150
Z-score
Z-score
40
20
0
-20
-40
-60
10
-10
-30
C
D
Column random model
4
3
2
1
-1
-2
-3

Figure S62

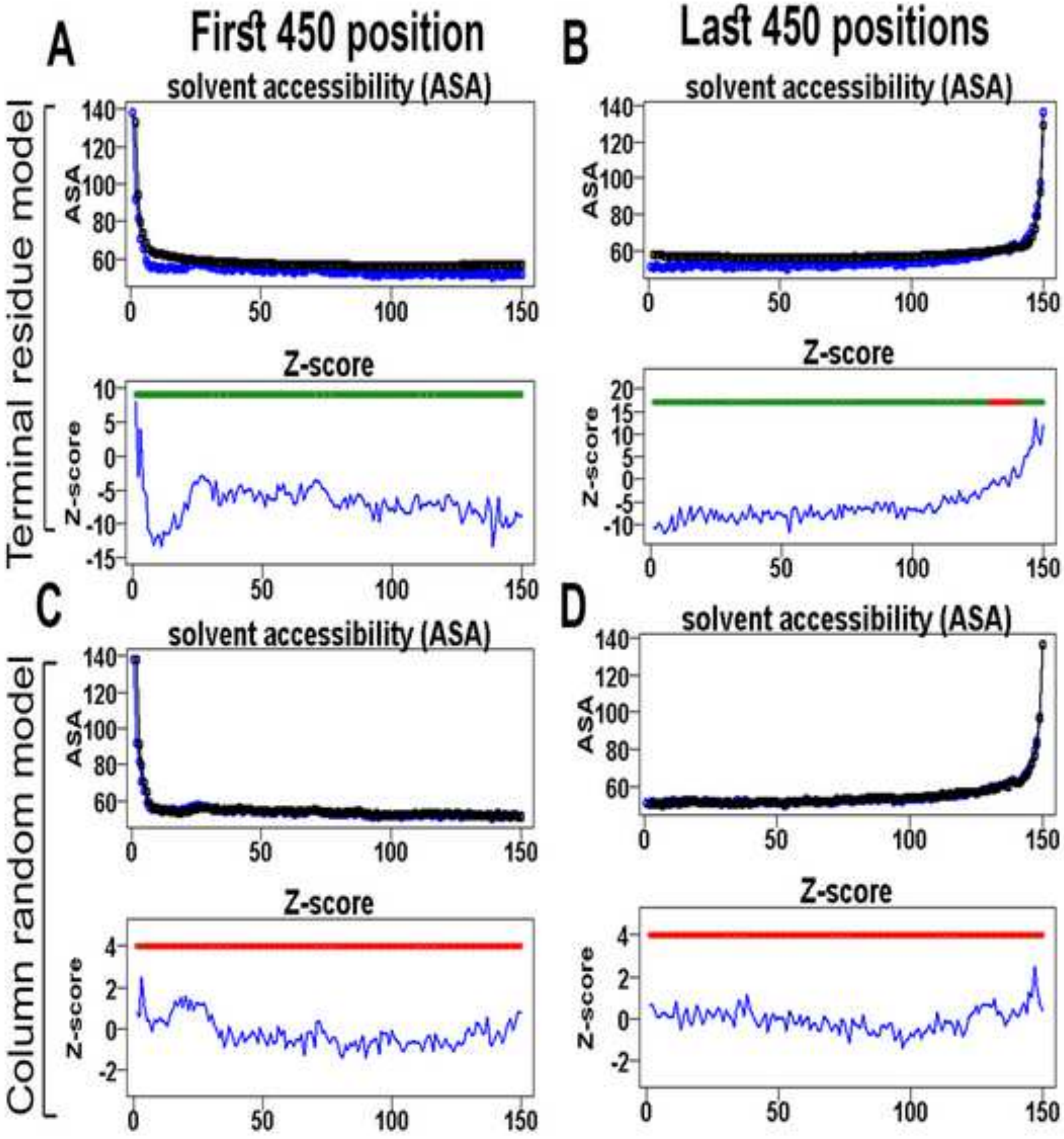

A
First 450 position
B
Last 450 positions
solvent accessibility (ASA)
Z-score
C
D
Terminal residue model
Column random model
ASA
Z-score

**Figure S63**

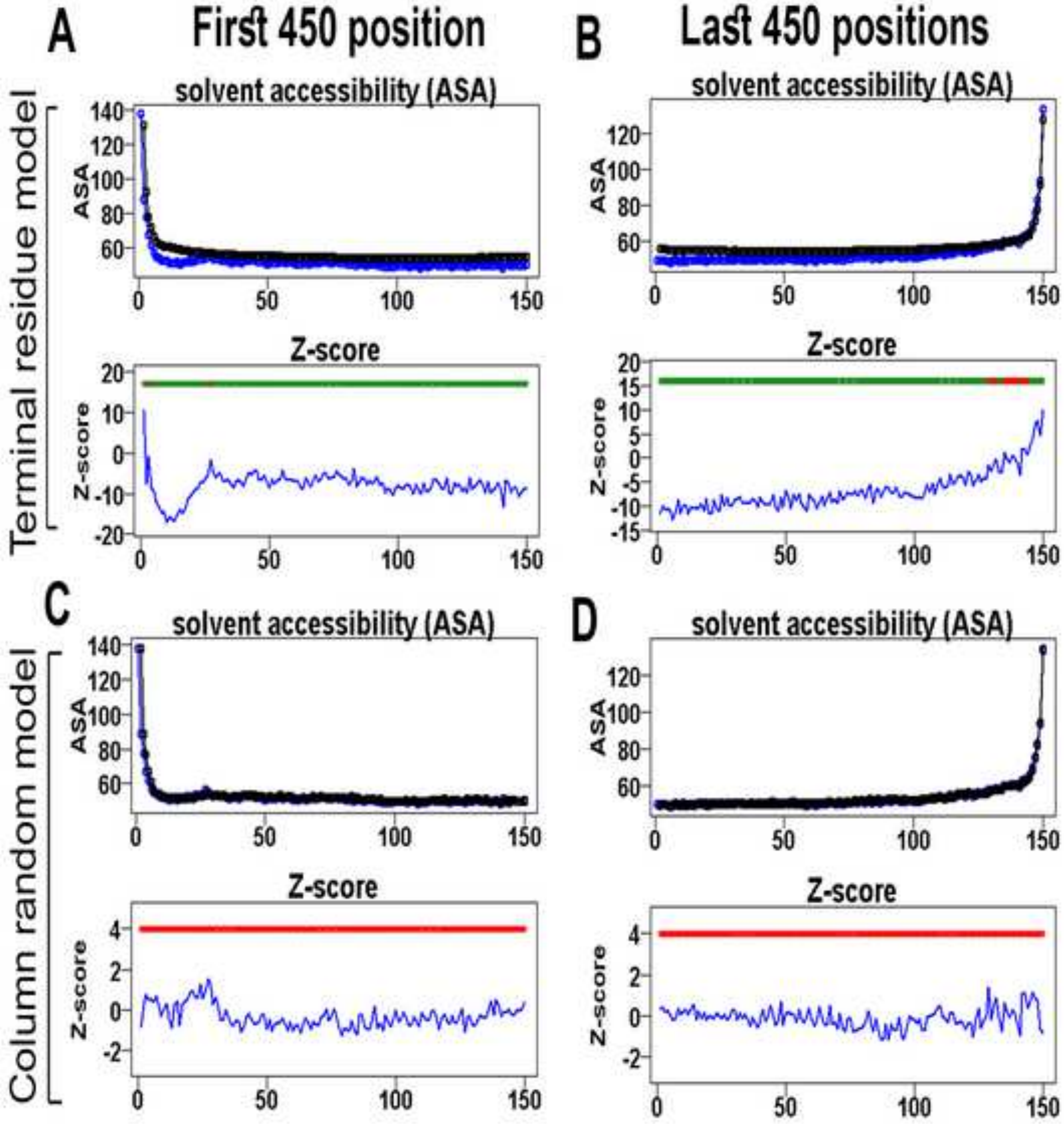
A
First 450 position
B
Last 450 positions
Terminal residue model
solvent accessibility (ASA)
ASA
Z-score
C
D
Column random model

**Figure S64**

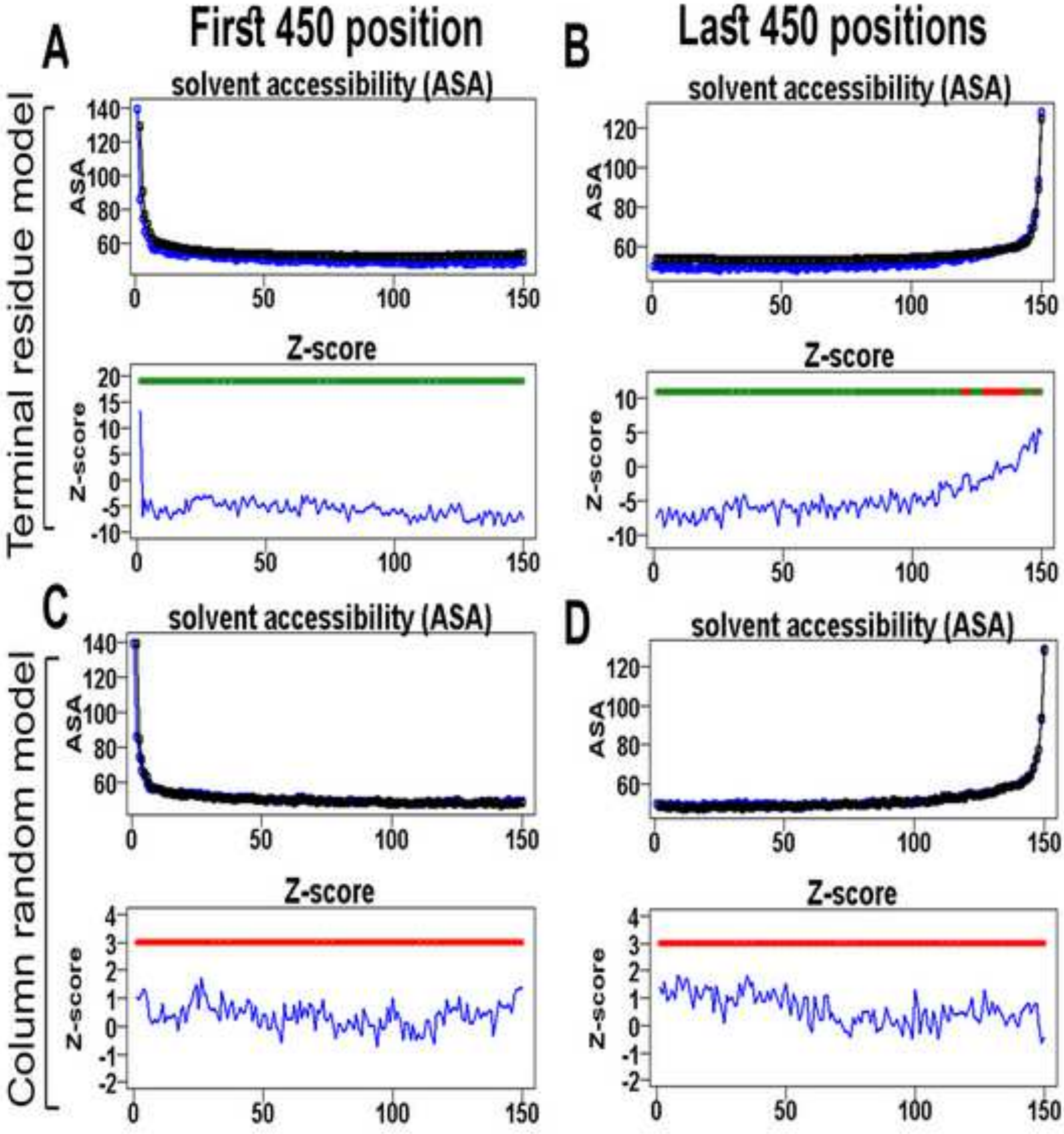
A
First 450 position
B
Last 450 positions
Terminal residue model
solvent accessibility (ASA)
ASA
140
120
100
80
60
0
50
100
150
Z-score
Z-score
20
15
10
5
0
-5
-10
C
D
Column random model
4
3
2
1
0
-1
-2

Figure S65

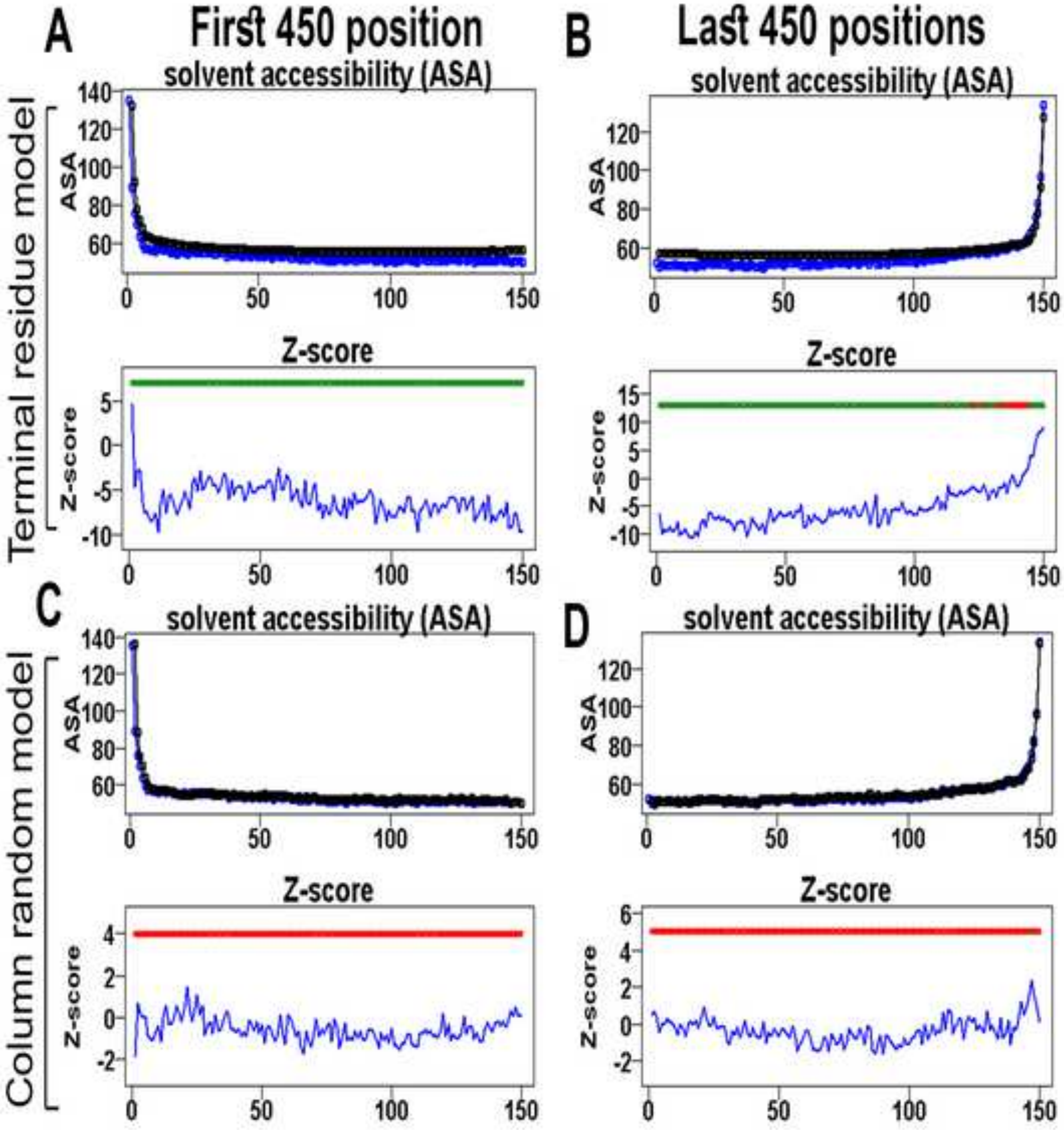

A
First 450 position
B
Last 450 positions
Terminal residue model
Column random model
solvent accessibility (ASA)
ASA
Z-score
Z-score
C
D
solvent accessibility (ASA)
Z-score

Figure S66

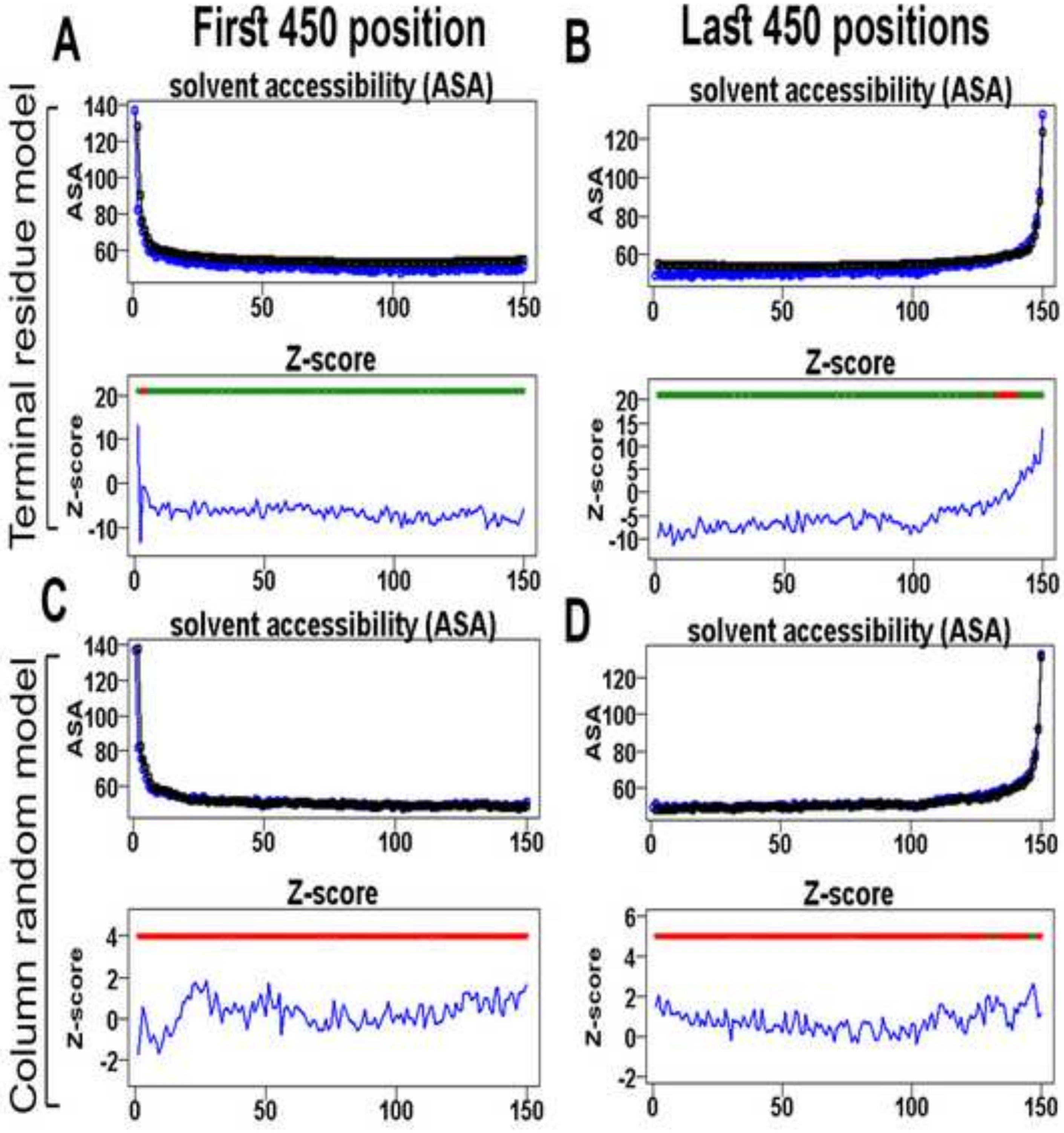

A
First 450 position
B
Last 450 positions
Terminal residue model
Column random model
solvent accessibility (ASA)
ASA
Z-score
C
D

Figure S67

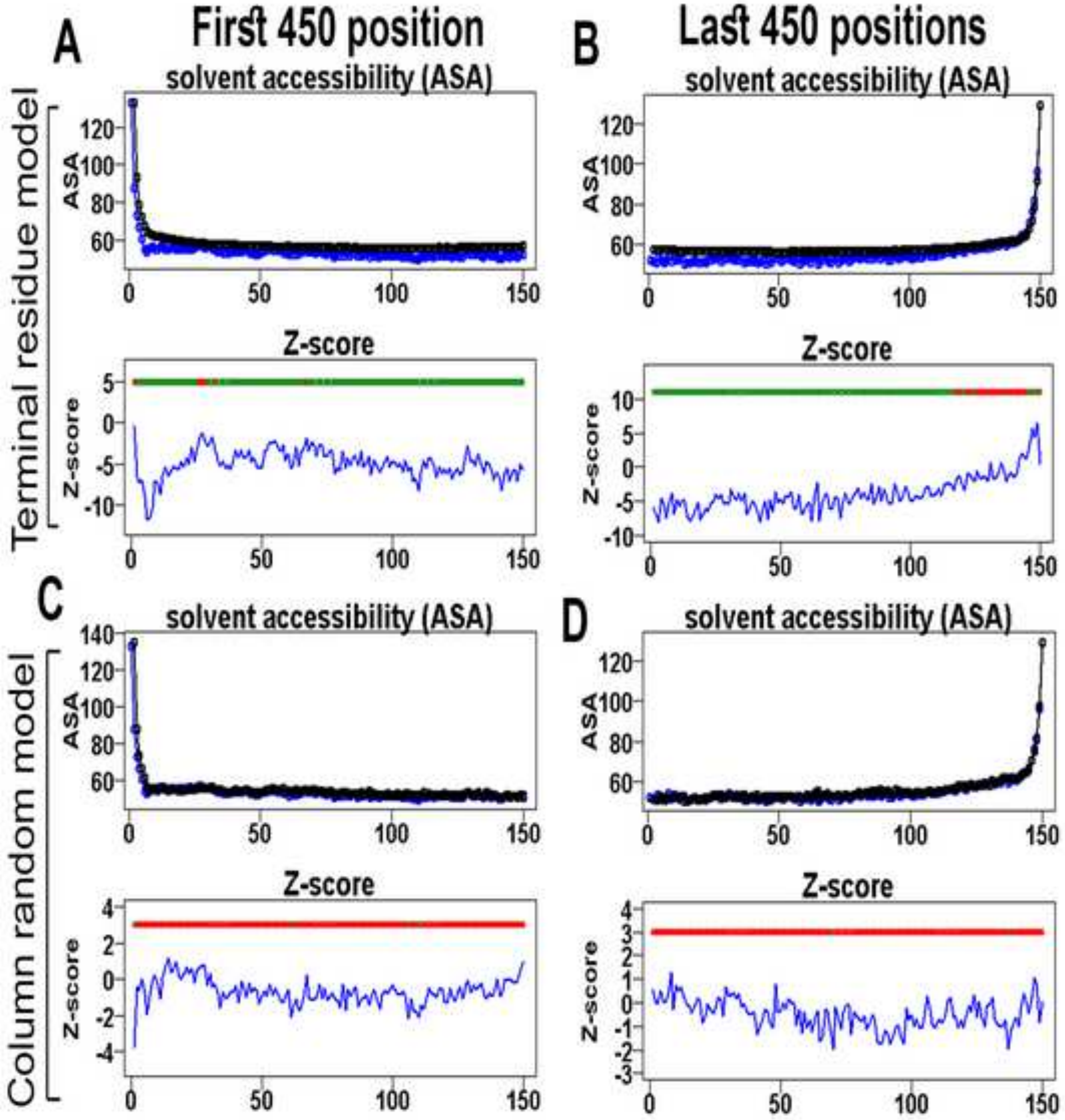
A
First 450 position
B
Last 450 positions
Terminal residue model
Column random model
solvent accessibility (ASA)
ASA
Z-score
C
D
0
50
100
150

**Figure S68**

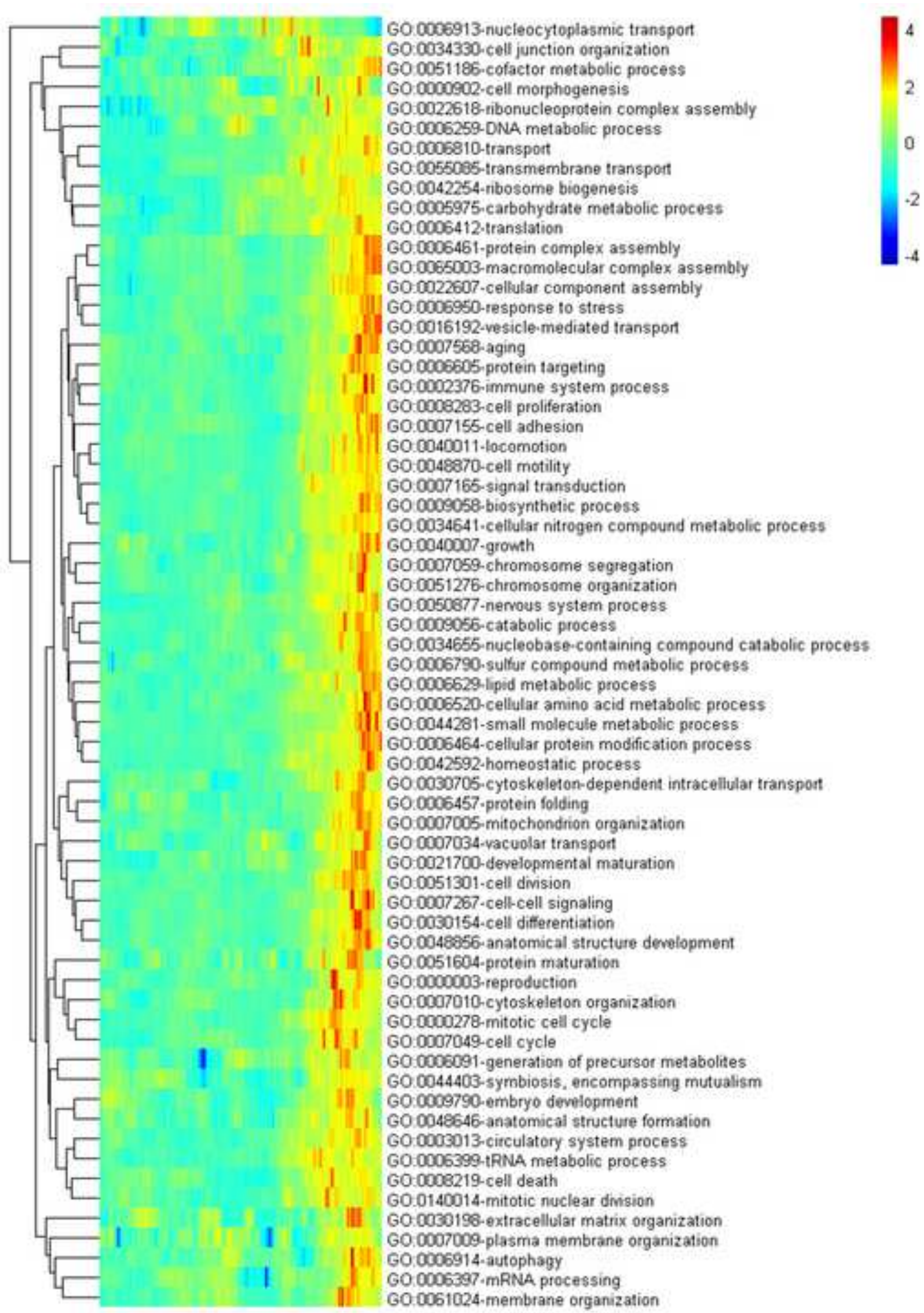

GO:0006913-nucleocytoplasmic transport
GO:0034330-cell junction organization
GO:0051186-cofactor metabolic process
GO:0000902-cell morphogenesis
GO:0022618-ribonucleoprotein complex assembly
GO:0006259-DNA metabolic process
GO:0006810-transport
GO:0055085-transmembrane transport
GO:0042254-ribosome biogenesis
GO:0005975-carbohydrate metabolic process
GO:0006412-translation
GO:0006461-protein complex assembly
GO:0065003-macromolecular complex assembly
GO:0022607-cellular component assembly
GO:0006950-response to stress
GO:0016192-vesicle-mediated transport
GO:0007568-aging
GO:0006605-protein targeting
GO:0002376-immune system process
GO:0008283-cell proliferation
GO:0007155-cell adhesion
GO:0040011-locomotion
GO:0048870-cell motility
GO:0007165-signal transduction
GO:0009058-biosynthetic process
GO:0034641-cellular nitrogen compound metabolic process
GO:0040007-growth
GO:0007059-chromosome segregation
GO:0051276-chromosome organization
GO:0050877-nervous system process
GO:0009056-catabolic process
GO:0034655-nucleobase-containing compound catabolic process
GO:0006790-sulfur compound metabolic process
GO:0006629-lipid metabolic process
GO:0006520-cellular amino acid metabolic process
GO:0044281-small molecule metabolic process
GO:0006464-cellular protein modification process
GO:0042592-homeostatic process
GO:0030705-cytoskeleton-dependent intracellular transport
GO:0006457-protein folding
GO:0007005-mitochondrion organization
GO:0007034-vacuolar transport
GO:0021700-developmental maturation
GO:0051301-cell division
GO:0007267-cell-cell signaling
GO:0030154-cell differentiation
GO:0048856-anatomical structure development
GO:0051604-protein maturation
GO:0000003-reproduction
GO:0007010-cytoskeleton organization
GO:0000278-mitotic cell cycle
GO:0007049-cell cycle
GO:0006091-generation of precursor metabolites
GO:0044403-symbiosis, encompassing mutualism
GO:0009790-embryo development
GO:0048646-anatomical structure formation
GO:0003013-circulatory system process
GO:0006399-tRNA metabolic process
GO:0008219-cell death
GO:0140014-mitotic nuclear division
GO:0030198-extracellular matrix organization
GO:0007009-plasma membrane organization
GO:0006914-autophagy
GO:0006397-mRNA processing
GO:0061024-membrane organization
4
2
0
-2
-4

**Figure S69**

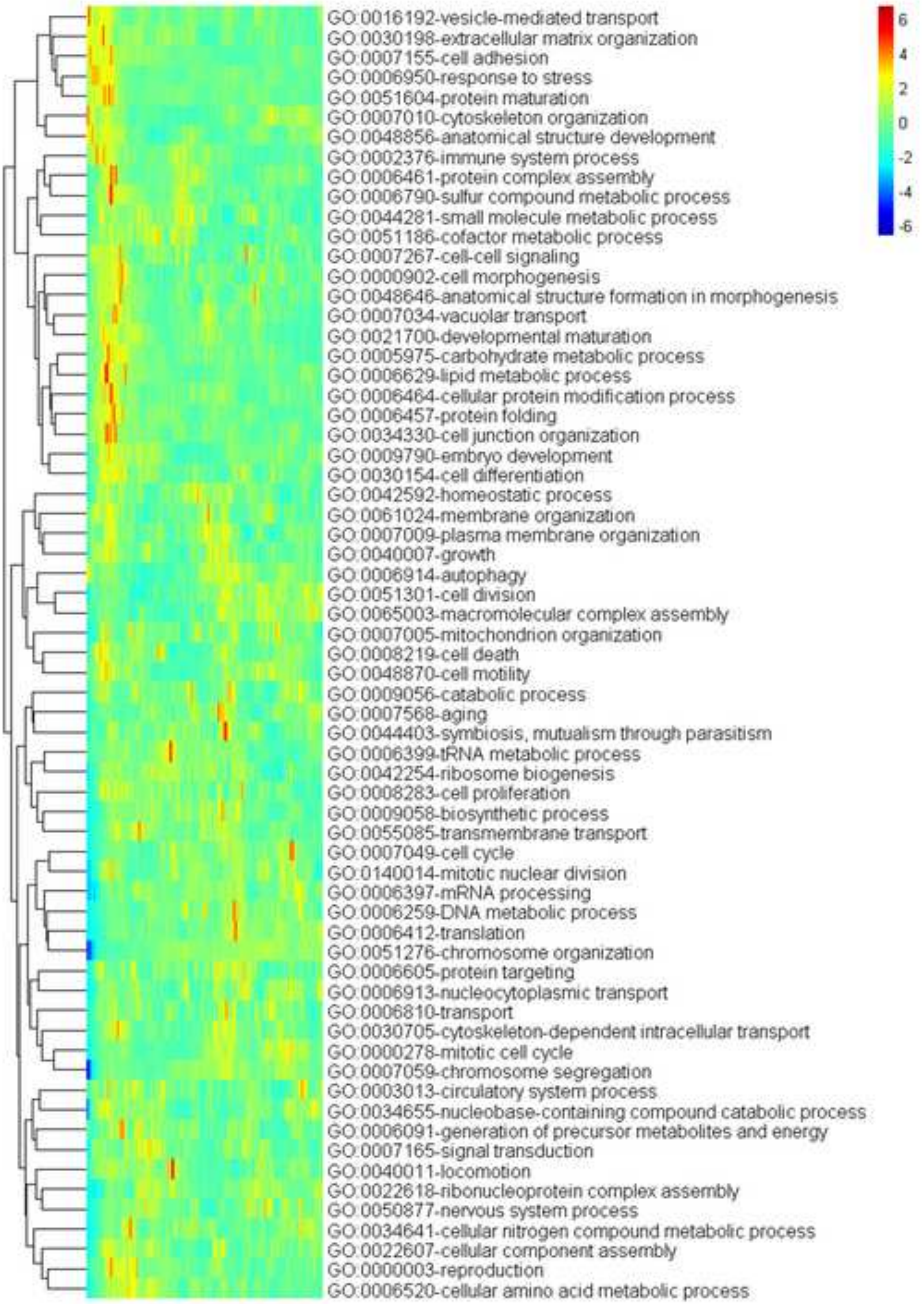
GO:0016192-vesicle-mediated transport
GO:0030198-extracellular matrix organization
GO:0007155-cell adhesion
GO:0006950-response to stress
GO:0051604-protein maturation
GO:0007010-cytoskeleton organization
GO:0048856-anatomical structure development
GO:0002376-immune system process
GO:0006461-protein complex assembly
GO:0006790-sulfur compound metabolic process
GO:0044281-small molecule metabolic process
GO:0051186-cofactor metabolic process
GO:0007267-cell-cell signaling
GO:0000902-cell morphogenesis
GO:0048646-anatomical structure formation in morphogenesis
GO:0007034-vacuolar transport
GO:0021700-developmental maturation
GO:0005975-carbohydrate metabolic process
GO:0006629-lipid metabolic process
GO:0006464-cellular protein modification process
GO:0006457-protein folding
GO:0034330-cell junction organization
GO:0009790-embryo development
GO:0030154-cell differentiation
GO:0042592-homeostatic process
GO:0061024-membrane organization
GO:0007009-plasma membrane organization
GO:0040007-growth
GO:0006914-autophagy
GO:0051301-cell division
GO:0065003-macromolecular complex assembly
GO:0007005-mitochondrion organization
GO:0008219-cell death
GO:0048870-cell motility
GO:0009056-catabolic process
GO:0007568-aging
GO:0044403-symbiosis, mutualism through parasitism
GO:0006399-tRNA metabolic process
GO:0042254-ribosome biogenesis
GO:0008283-cell proliferation
GO:0009058-biosynthetic process
GO:0055085-transmembrane transport
GO:0007049-cell cycle
GO:0140014-mitotic nuclear division
GO:0006397-mRNA processing
GO:0006259-DNA metabolic process
GO:0006412-translation
GO:0051276-chromosome organization
GO:0006605-protein targeting
GO:0006913-nucleocytoplasmic transport
GO:0006810-transport
GO:0030705-cytoskeleton-dependent intracellular transport
GO:0000278-mitotic cell cycle
GO:0007059-chromosome segregation
GO:0003013-circulatory system process
GO:0034655-nucleobase-containing compound catabolic process
GO:0006091-generation of precursor metabolites and energy
GO:0007165-signal transduction
GO:0040011-locomotion
GO:0022618-ribonucleoprotein complex assembly
GO:0050877-nervous system process
GO:0034641-cellular nitrogen compound metabolic process
GO:0022607-cellular component assembly
GO:0000003-reproduction
GO:0006520-cellular amino acid metabolic process
6
4
2
0
-2
-4
-6

**Figure S70**

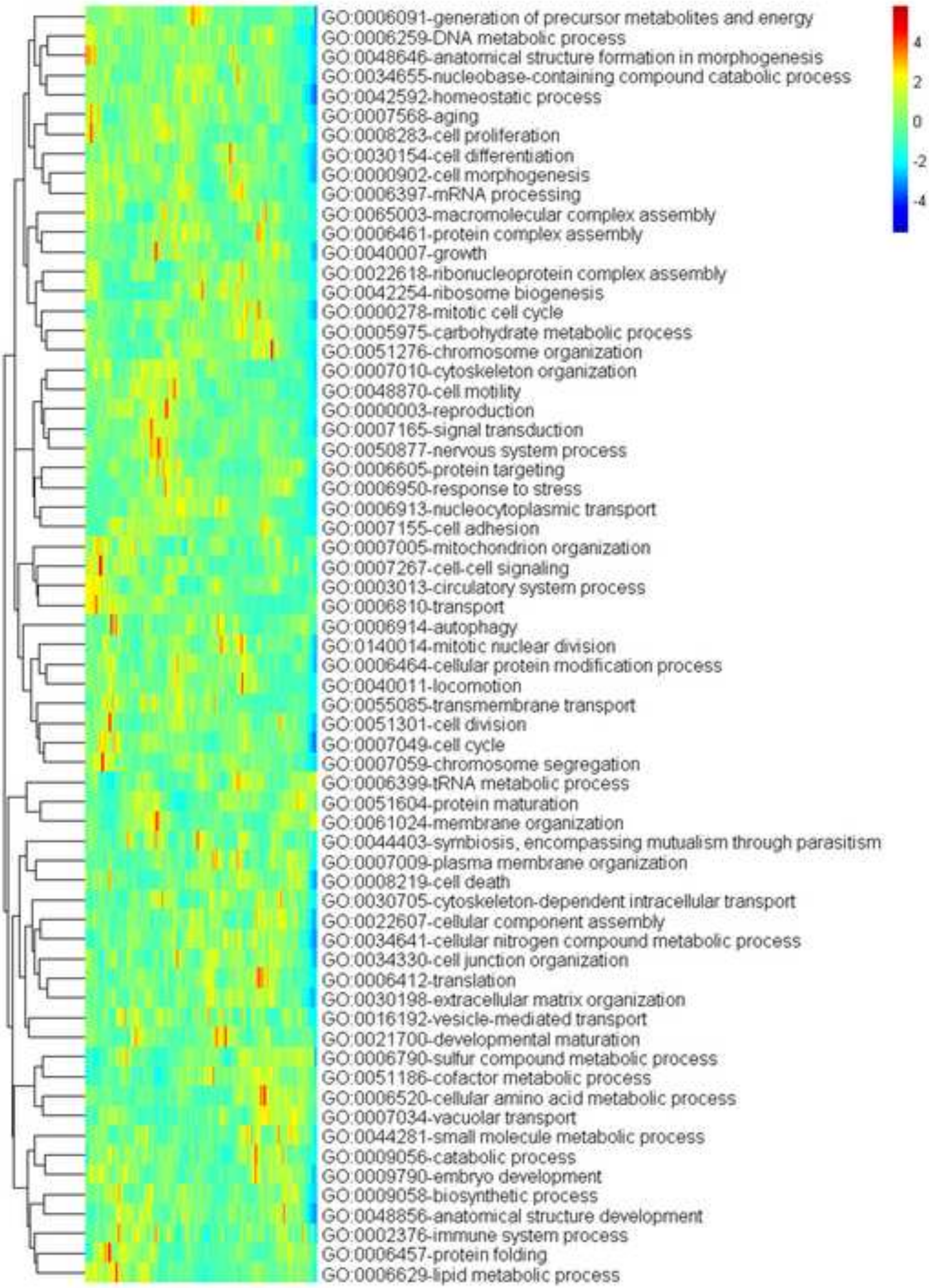
GO:0006091-generation of precursor metabolites and energy
GO:0006259-DNA metabolic process
GO:0048646-anatomical structure formation in morphogenesis
GO:0034655-nucleobase-containing compound catabolic process
GO:0042592-homeostatic process
GO:0007568-aging
GO:0008283-cell proliferation
GO:0030154-cell differentiation
GO:0000902-cell morphogenesis
GO:0006397-mRNA processing
GO:0065003-macromolecular complex assembly
GO:0006461-protein complex assembly
GO:0040007-growth
GO:0022618-ribonucleoprotein complex assembly
GO:0042254-ribosome biogenesis
GO:0000278-mitotic cell cycle
GO:0005975-carbohydrate metabolic process
GO:0051276-chromosome organization
GO:0007010-cytoskeleton organization
GO:0048870-cell motility
GO:0000003-reproduction
GO:0007165-signal transduction
GO:0050877-nervous system process
GO:0006605-protein targeting
GO:0006950-response to stress
GO:0006913-nucleocytoplasmic transport
GO:0007155-cell adhesion
GO:0007005-mitochondrion organization
GO:0007267-cell-cell signaling
GO:0003013-circulatory system process
GO:0006810-transport
GO:0006914-autophagy
GO:0140014-mitotic nuclear division
GO:0006464-cellular protein modification process
GO:0040011-locomotion
GO:0055085-transmembrane transport
GO:0051301-cell division
GO:0007049-cell cycle
GO:0007059-chromosome segregation
GO:0006399-tRNA metabolic process
GO:0051604-protein maturation
GO:0061024-membrane organization
GO:0044403-symbiosis, encompassing mutualism through parasitism
GO:0007009-plasma membrane organization
GO:0008219-cell death
GO:0030705-cytoskeleton-dependent intracellular transport
GO:0022607-cellular component assembly
GO:0034641-cellular nitrogen compound metabolic process
GO:0034330-cell junction organization
GO:0006412-translation
GO:0030198-extracellular matrix organization
GO:0016192-vesicle-mediated transport
GO:0021700-developmental maturation
GO:0006790-sulfur compound metabolic process
GO:0051186-cofactor metabolic process
GO:0006520-cellular amino acid metabolic process
GO:0007034-vacuolar transport
GO:0044281-small molecule metabolic process
GO:0009056-catabolic process
GO:0009790-embryo development
GO:0009058-biosynthetic process
GO:0048856-anatomical structure development
GO:0002376-immune system process
GO:0006457-protein folding
GO:0006629-lipid metabolic process
4
2
0
-2
-4

**Figure S71**

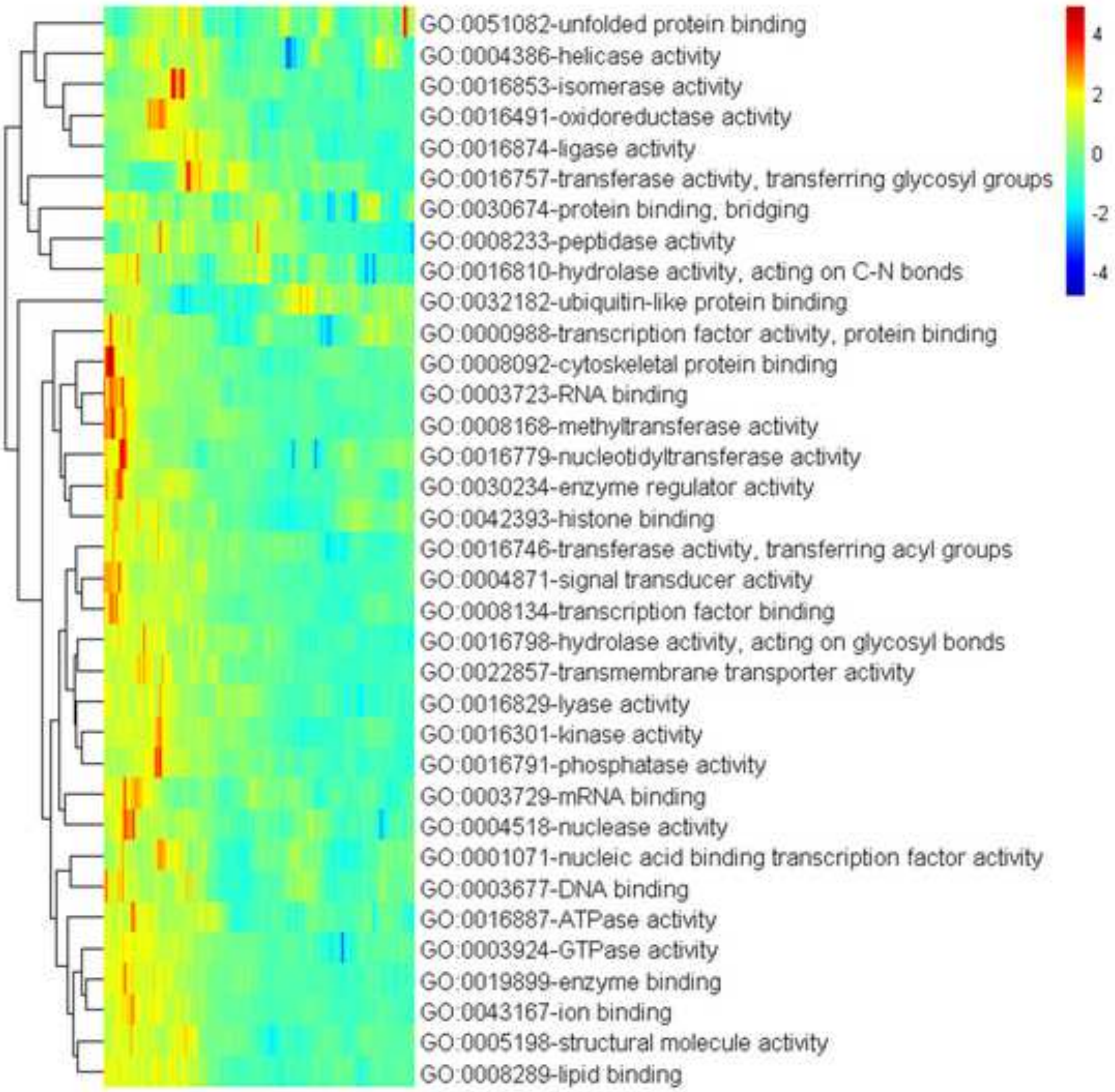
GO:0051082-unfolded protein binding
GO:0004386-helicase activity
GO:0016853-isomerase activity
GO:0016491-oxidoreductase activity
GO:0016874-ligase activity
GO:0016757-transferase activity, transferring glycosyl groups
GO:0030674-protein binding, bridging
GO:0008233-peptidase activity
GO:0016810-hydrolase activity, acting on C-N bonds
GO:0032182-ubiquitin-like protein binding
GO:0000988-transcription factor activity, protein binding
GO:0008092-cytoskeletal protein binding
GO:0003723-RNA binding
GO:0008168-methyltransferase activity
GO:0016779-nucleotidyltransferase activity
GO:0030234-enzyme regulator activity
GO:0042393-histone binding
GO:0016746-transferase activity, transferring acyl groups
GO:0004871-signal transducer activity
GO:0008134-transcription factor binding
GO:0016798-hydrolase activity, acting on glycosyl bonds
GO:0022857-transmembrane transporter activity
GO:0016829-lyase activity
GO:0016301-kinase activity
GO:0016791-phosphatase activity
GO:0003729-mRNA binding
GO:0004518-nuclease activity
GO:0001071-nucleic acid binding transcription factor activity
GO:0003677-DNA binding
GO:0016887-ATPase activity
GO:0003924-GTPase activity
GO:0019899-enzyme binding
GO:0043167-ion binding
GO:0005198-structural molecule activity
GO:0008289-lipid binding
4
2
0
-2
-4

**Figure S72**

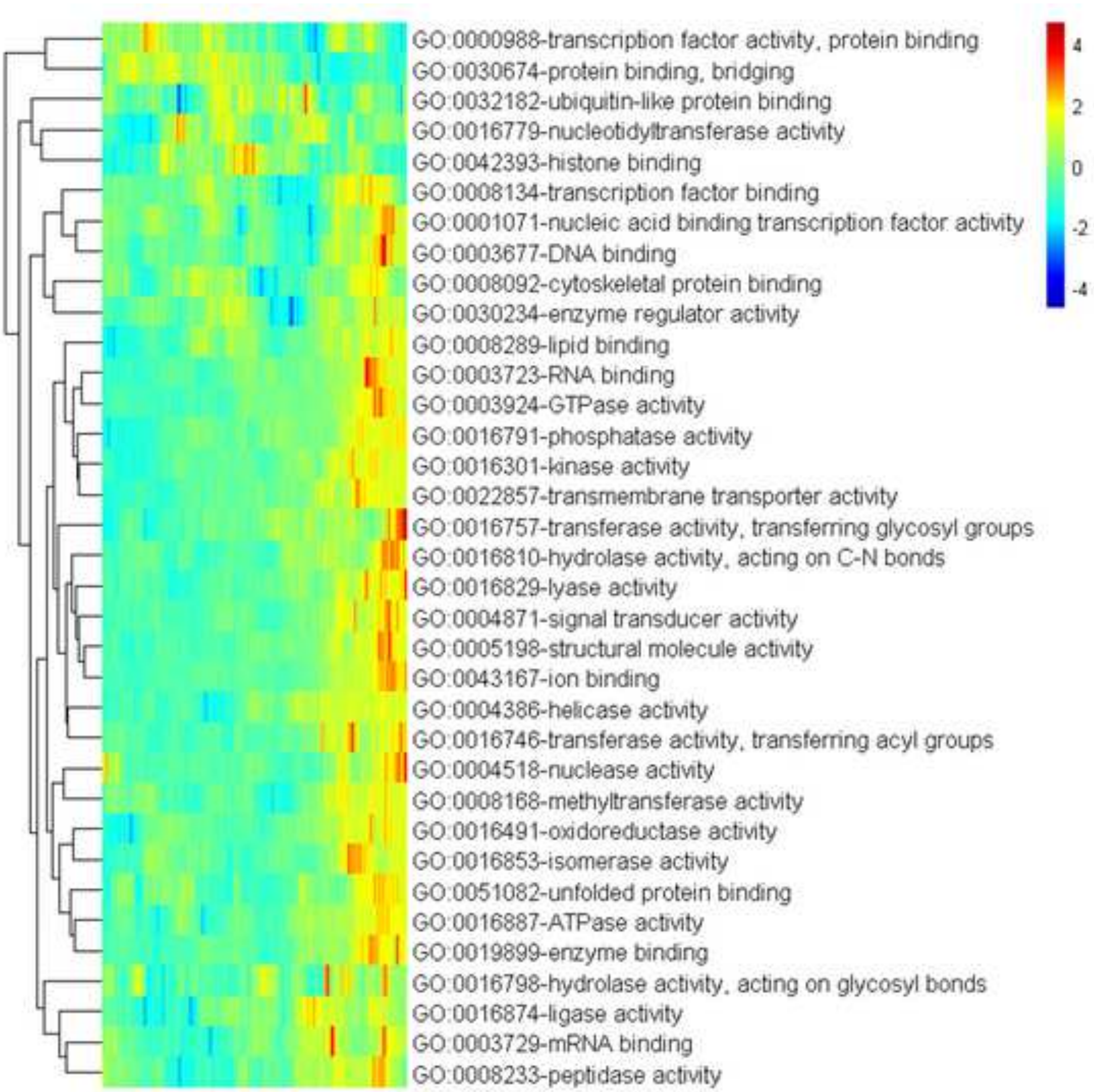
GO:0000988-transcription factor activity, protein binding
GO:0030674-protein binding, bridging
GO:0032182-ubiquitin-like protein binding
GO:0016779-nucleotidyltransferase activity
GO:0042393-histone binding
GO:0008134-transcription factor binding
GO:0001071-nucleic acid binding transcription factor activity
GO:0003677-DNA binding
GO:0008092-cytoskeletal protein binding
GO:0030234-enzyme regulator activity
GO:0008289-lipid binding
GO:0003723-RNA binding
GO:0003924-GTPase activity
GO:0016791-phosphatase activity
GO:0016301-kinase activity
GO:0022857-transmembrane transporter activity
GO:0016757-transferase activity, transferring glycosyl groups
GO:0016810-hydrolase activity, acting on C-N bonds
GO:0016829-lyase activity
GO:0004871-signal transducer activity
GO:0005198-structural molecule activity
GO:0043167-ion binding
GO:0004386-helicase activity
GO:0016746-transferase activity, transferring acyl groups
GO:0004518-nuclease activity
GO:0008168-methyltransferase activity
GO:0016491-oxidoreductase activity
GO:0016853-isomerase activity
GO:0051082-unfolded protein binding
GO:0016887-ATPase activity
GO:0019899-enzyme binding
GO:0016798-hydrolase activity, acting on glycosyl bonds
GO:0016874-ligase activity
GO:0003729-mRNA binding
GO:0008233-peptidase activity
4
2
0
-2
-4

Figure S73

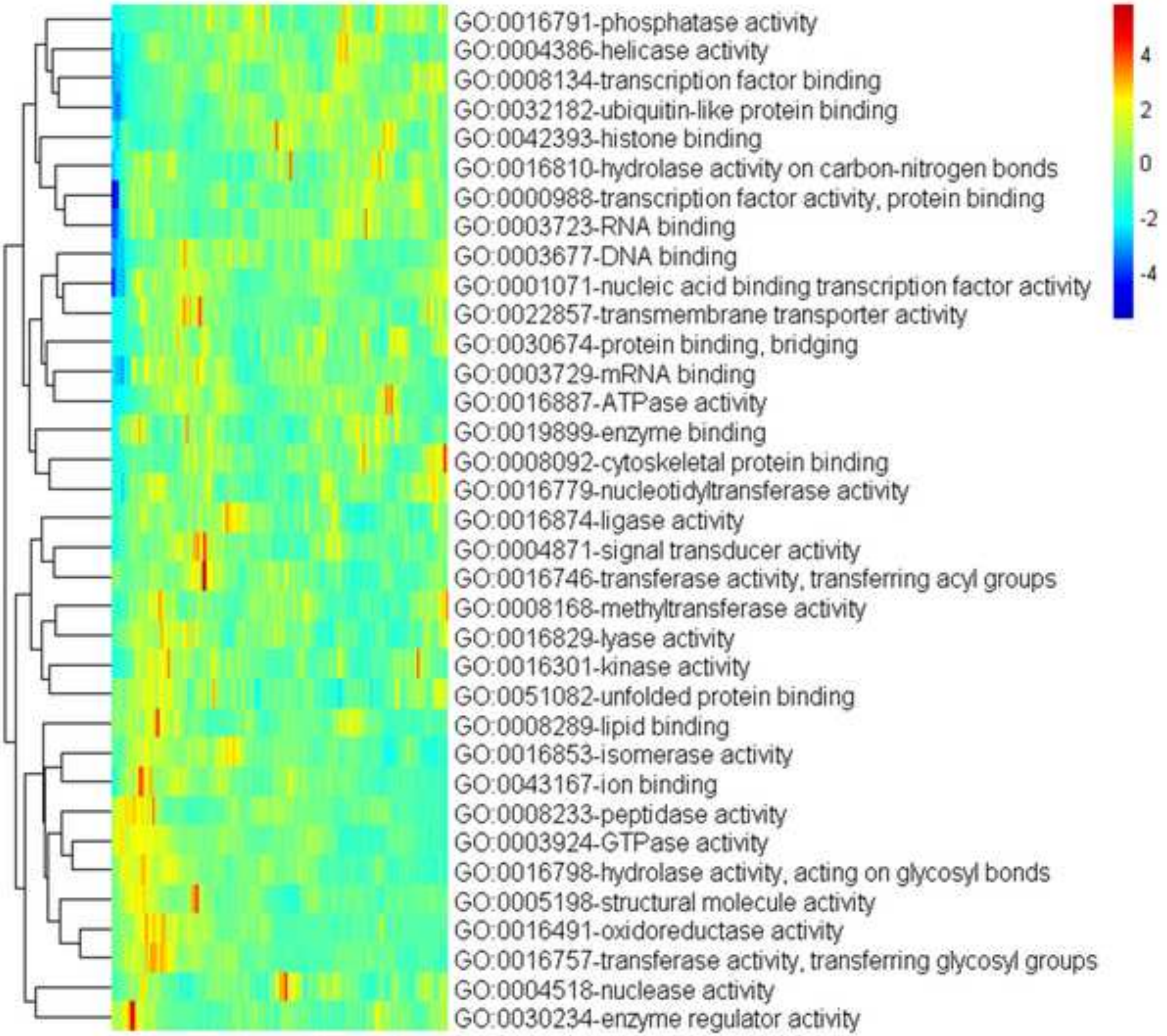

GO:0016791-phosphatase activity
GO:0004386-helicase activity
GO:0008134-transcription factor binding
GO:0032182-ubiquitin-like protein binding
GO:0042393-histone binding
GO:0016810-hydrolase activity on carbon-nitrogen bonds
GO:0000988-transcription factor activity, protein binding
GO:0003723-RNA binding
GO:0003677-DNA binding
GO:0001071-nucleic acid binding transcription factor activity
GO:0022857-transmembrane transporter activity
GO:0030674-protein binding, bridging
GO:0003729-mRNA binding
GO:0016887-ATPase activity
GO:0019899-enzyme binding
GO:0008092-cytoskeletal protein binding
GO:0016779-nucleotidyltransferase activity
GO:0016874-ligase activity
GO:0004871-signal transducer activity
GO:0016746-transferase activity, transferring acyl groups
GO:0008168-methyltransferase activity
GO:0016829-lyase activity
GO:0016301-kinase activity
GO:0051082-unfolded protein binding
GO:0008289-lipid binding
GO:0016853-isomerase activity
GO:0043167-ion binding
GO:0008233-peptidase activity
GO:0003924-GTPase activity
GO:0016798-hydrolase activity, acting on glycosyl bonds
GO:0005198-structural molecule activity
GO:0016491-oxidoreductase activity
GO:0016757-transferase activity, transferring glycosyl groups
GO:0004518-nuclease activity
GO:0030234-enzyme regulator activity
4
2
0
-2
-4

**Figure S74**

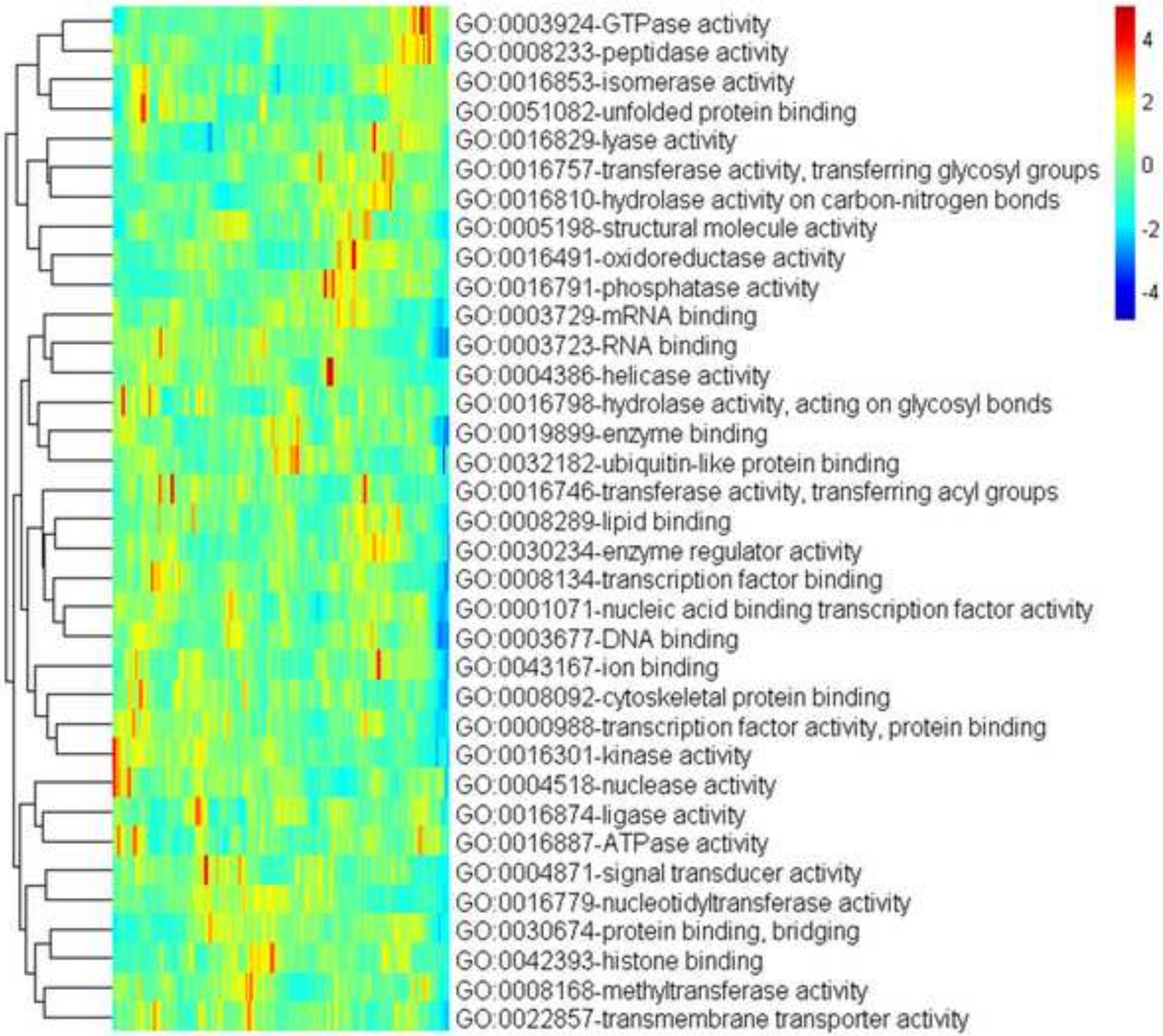
GO:0003924-GTPase activity
GO:0008233-peptidase activity
GO:0016853-isomerase activity
GO:0051082-unfolded protein binding
GO:0016829-lyase activity
GO:0016757-transferase activity, transferring glycosyl groups
GO:0016810-hydrolase activity on carbon-nitrogen bonds
GO:0005198-structural molecule activity
GO:0016491-oxidoreductase activity
GO:0016791-phosphatase activity
GO:0003729-mRNA binding
GO:0003723-RNA binding
GO:0004386-helicase activity
GO:0016798-hydrolase activity, acting on glycosyl bonds
GO:0019899-enzyme binding
GO:0032182-ubiquitin-like protein binding
GO:0016746-transferase activity, transferring acyl groups
GO:0008289-lipid binding
GO:0030234-enzyme regulator activity
GO:0008134-transcription factor binding
GO:0001071-nucleic acid binding transcription factor activity
GO:0003677-DNA binding
GO:0043167-ion binding
GO:0008092-cytoskeletal protein binding
GO:0000988-transcription factor activity, protein binding
GO:0016301-kinase activity
GO:0004518-nuclease activity
GO:0016874-ligase activity
GO:0016887-ATPase activity
GO:0004871-signal transducer activity
GO:0016779-nucleotidyltransferase activity
GO:0030674-protein binding, bridging
GO:0042393-histone binding
GO:0008168-methyltransferase activity
GO:0022857-transmembrane transporter activity
4
2
0
-2
-4

**Figure S75**

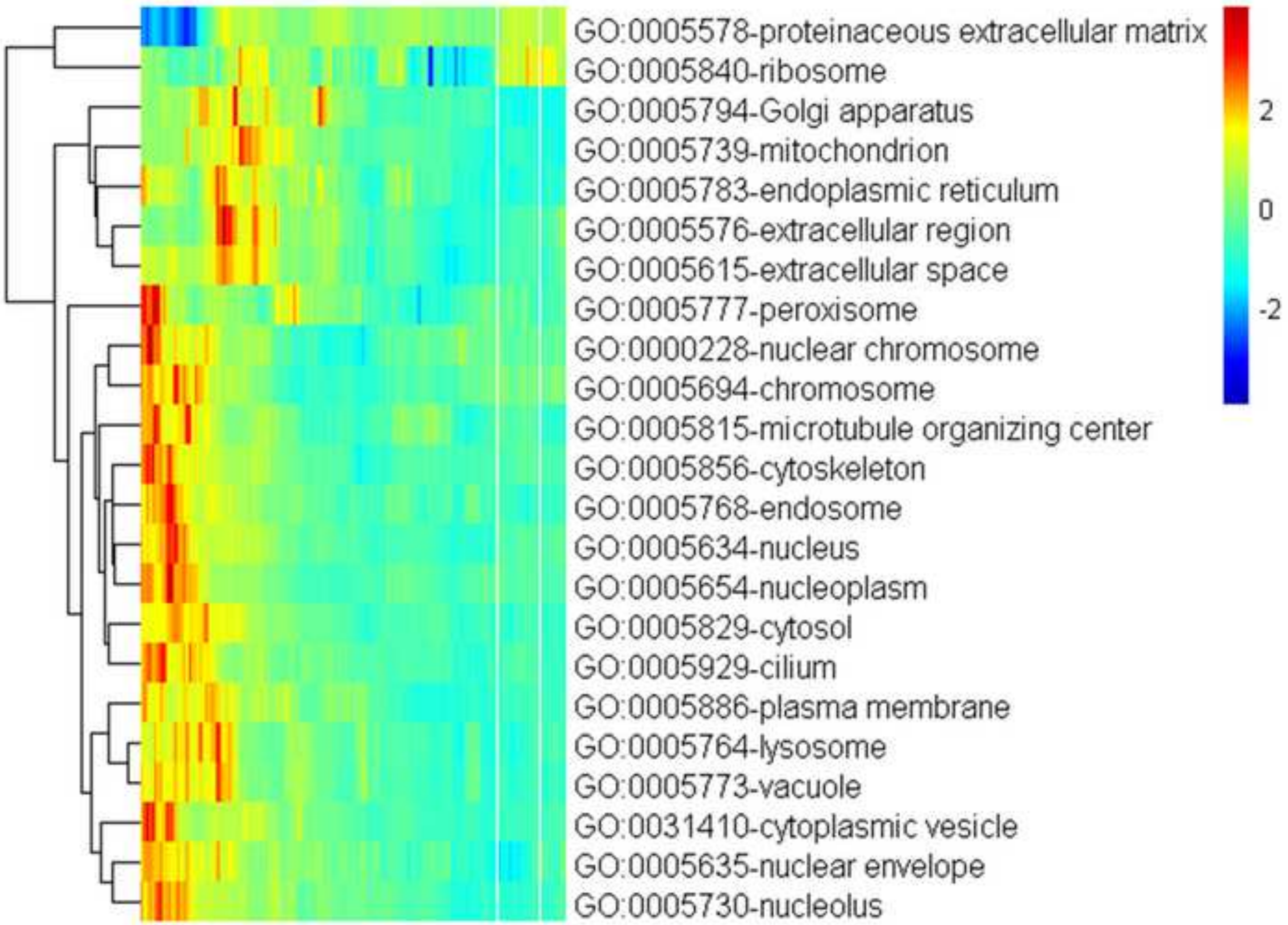
GO:0005578-proteinaceous extracellular matrix
GO:0005840-ribosome
GO:0005794-Golgi apparatus
GO:0005739-mitochondrion
GO:0005783-endoplasmic reticulum
GO:0005576-extracellular region
GO:0005615-extracellular space
GO:0005777-peroxisome
GO:0000228-nuclear chromosome
GO:0005694-chromosome
GO:0005815-microtubule organizing center
GO:0005856-cytoskeleton
GO:0005768-endosome
GO:0005634-nucleus
GO:0005654-nucleoplasm
GO:0005829-cytosol
GO:0005929-cilium
GO:0005886-plasma membrane
GO:0005764-lysosome
GO:0005773-vacuole
GO:0031410-cytoplasmic vesicle
GO:0005635-nuclear envelope
GO:0005730-nucleolus
2
0
-2

**Figure S76**

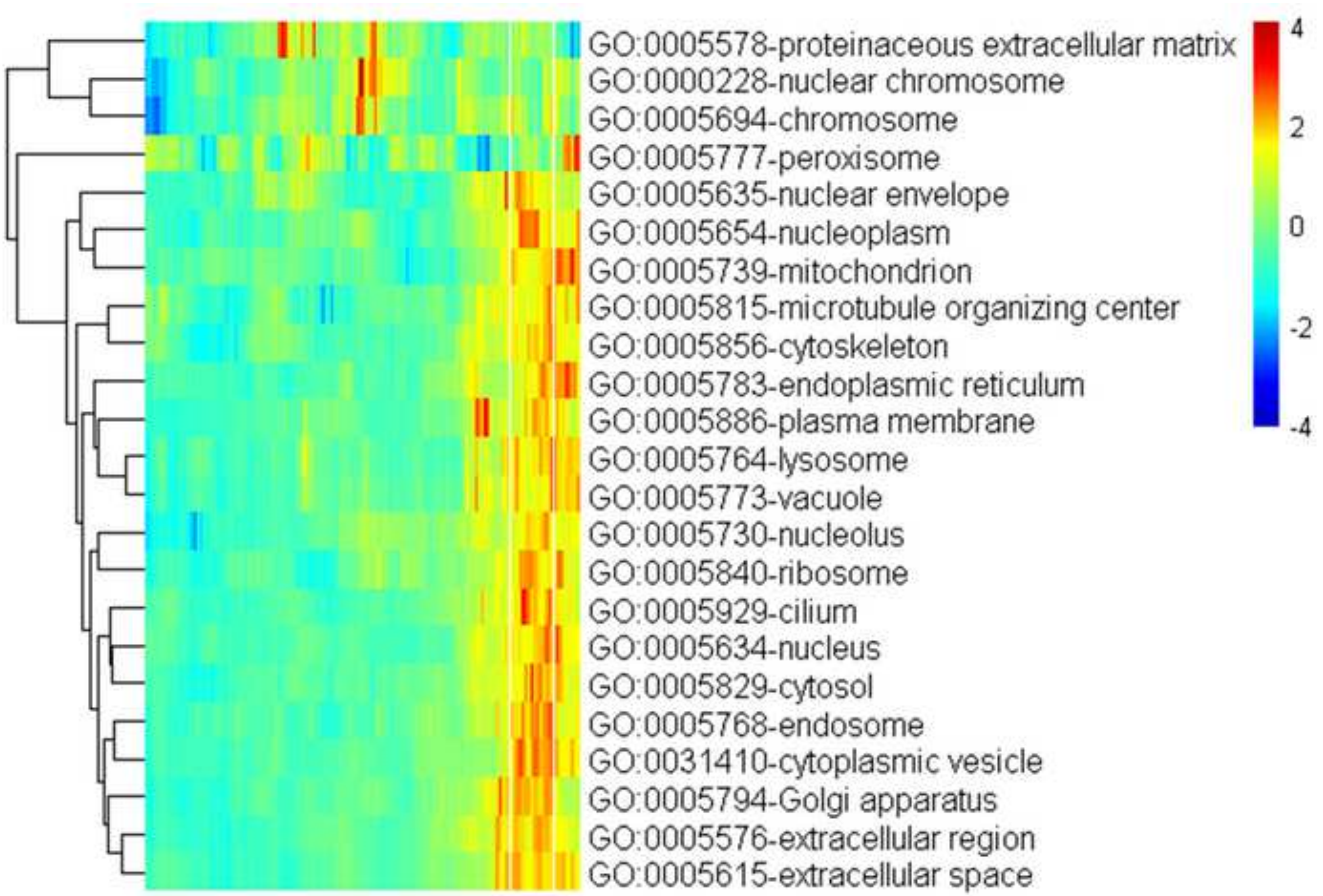
GO:0005578-proteinaceous extracellular matrix
GO:0000228-nuclear chromosome
GO:0005694-chromosome
GO:0005777-peroxisome
GO:0005635-nuclear envelope
GO:0005654-nucleoplasm
GO:0005739-mitochondrion
GO:0005815-microtubule organizing center
GO:0005856-cytoskeleton
GO:0005783-endoplasmic reticulum
GO:0005886-plasma membrane
GO:0005764-lysosome
GO:0005773-vacuole
GO:0005730-nucleolus
GO:0005840-ribosome
GO:0005929-cilium
GO:0005634-nucleus
GO:0005829-cytosol
GO:0005768-endosome
GO:0031410-cytoplasmic vesicle
GO:0005794-Golgi apparatus
GO:0005576-extracellular region
GO:0005615-extracellular space
4
2
0
-2
-4

**Figure S77**

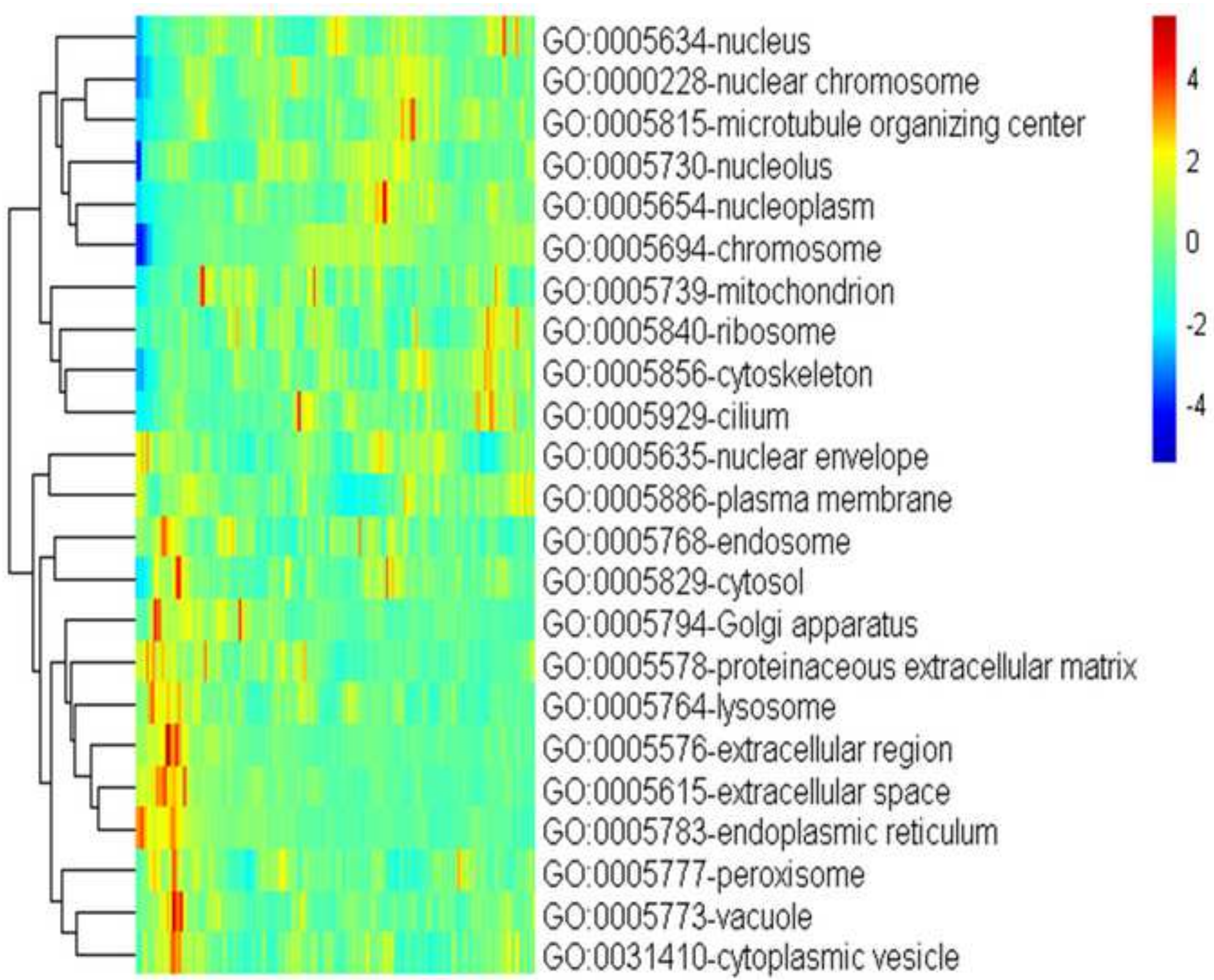
GO:0005634-nucleus
GO:0000228-nuclear chromosome
GO:0005815-microtubule organizing center
GO:0005730-nucleolus
GO:0005654-nucleoplasm
GO:0005694-chromosome
GO:0005739-mitochondrion
GO:0005840-ribosome
GO:0005856-cytoskeleton
GO:0005929-cilium
GO:0005635-nuclear envelope
GO:0005886-plasma membrane
GO:0005768-endosome
GO:0005829-cytosol
GO:0005794-Golgi apparatus
GO:0005578-proteinaceous extracellular matrix
GO:0005764-lysosome
GO:0005576-extracellular region
GO:0005615-extracellular space
GO:0005783-endoplasmic reticulum
GO:0005777-peroxisome
GO:0005773-vacuole
GO:0031410-cytoplasmic vesicle
4
2
0
-2
-4

**Figure S78**

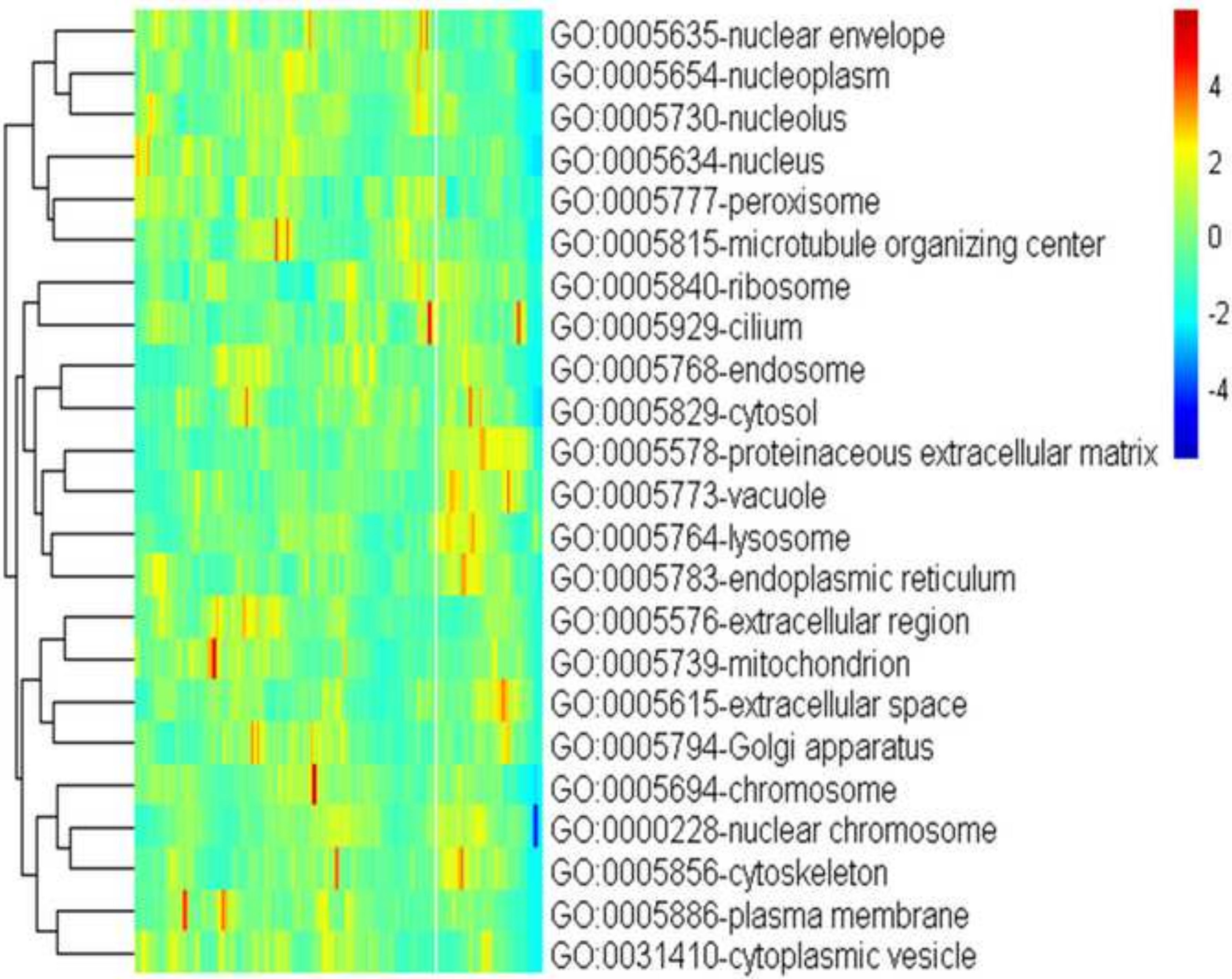
GO:0005635-nuclear envelope
GO:0005654-nucleoplasm
GO:0005730-nucleolus
GO:0005634-nucleus
GO:0005777-peroxisome
GO:0005815-microtubule organizing center
GO:0005840-ribosome
GO:0005929-cilium
GO:0005768-endosome
GO:0005829-cytosol
GO:0005578-proteinaceous extracellular matrix
GO:0005773-vacuole
GO:0005764-lysosome
GO:0005783-endoplasmic reticulum
GO:0005576-extracellular region
GO:0005739-mitochondrion
GO:0005615-extracellular space
GO:0005794-Golgi apparatus
GO:0005694-chromosome
GO:0000228-nuclear chromosome
GO:0005856-cytoskeleton
GO:0005886-plasma membrane
GO:0031410-cytoplasmic vesicle
4
2
0
-2
-4

**Figure S79**

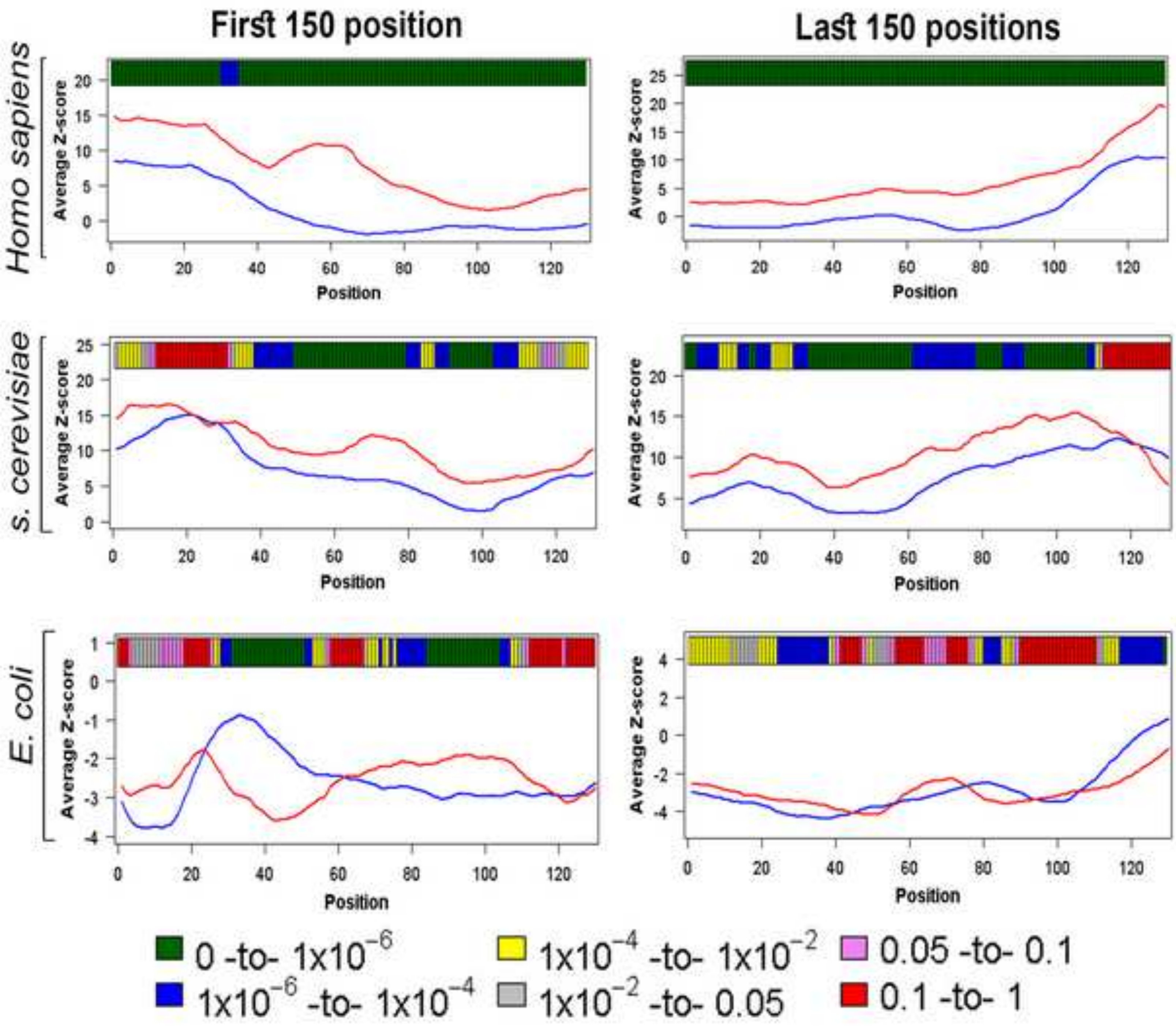

First 150 position
Last 150 positions
Homo sapiens
S. cerevisiae
E. coli
Average Z-score
Position
0 -to- 1x10$^{-6}$
1x10$^{-6}$ -to- 1x10$^{-4}$
1x10$^{-4}$ -to- 1x10$^{-2}$
1x10$^{-2}$ -to- 0.05
0.05 -to- 0.1
0.1 -to- 1

**Figure S80**

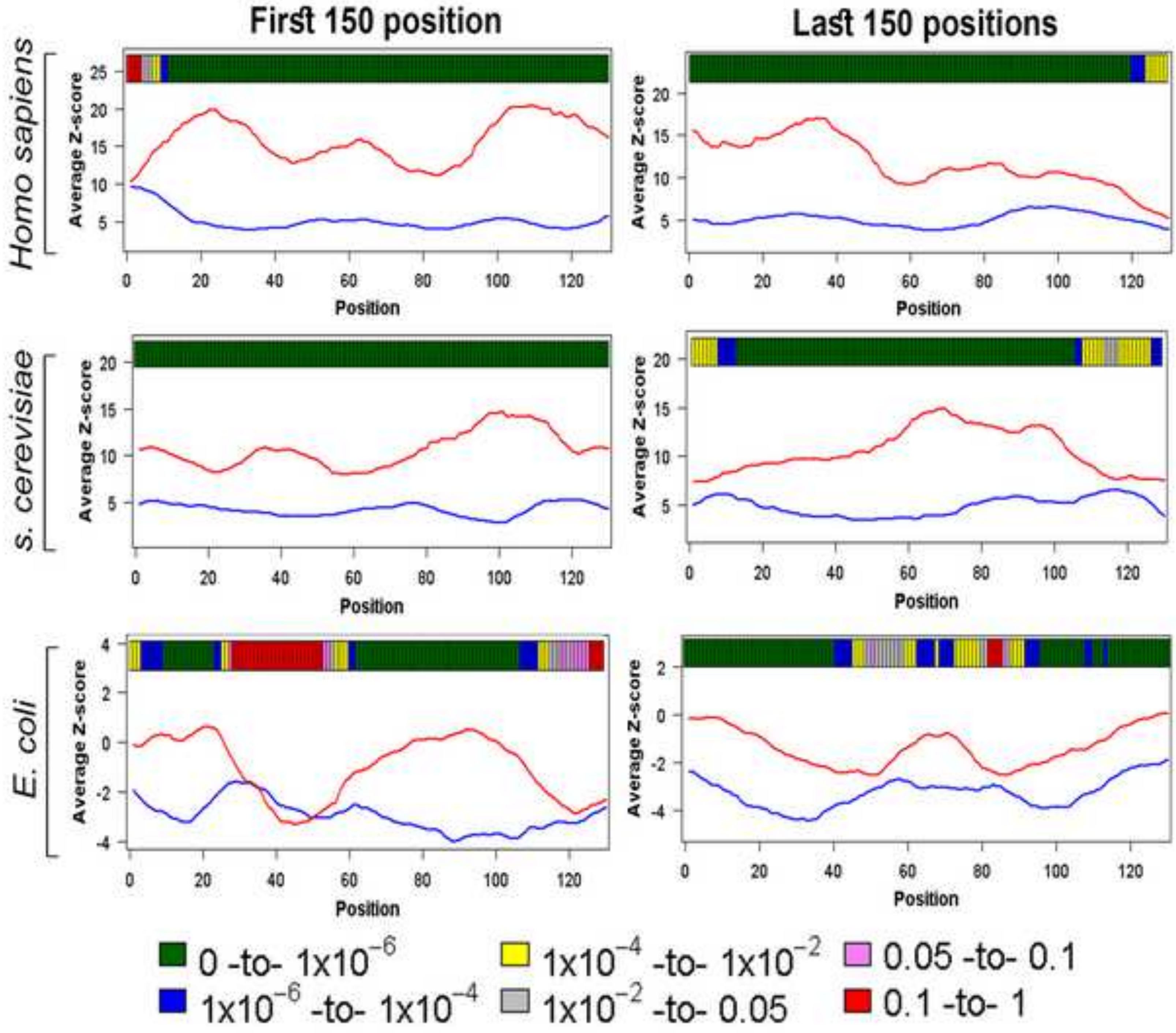

First 150 position
Last 150 positions
Homo sapiens
s. cerevisiae
E. coli
Average Z-score
Position
0 -to- 1x10^-6
1x10^-6 -to- 1x10^-4
1x10^-4 -to- 1x10^-2
1x10^-2 -to- 0.05
0.05 -to- 0.1
0.1 -to- 1

**Table S1**

**Table S1 List of genomes used in the analysis**

| Lineage | Species name (RefSeq accession number) | Genome type | Strain/ Genome assembly | Number of real proteins | No of proteins > 200 residues in length After filtering | Number of length conserved random sequences generated | No of terminal residue conserved random sequences | No column random sequences generated |
|---|---|---|---|---|---|---|---|---|
| Mammal | *Homo sapiens* (GCA_000001405.27) | Reference | GRCh38.p12 | 19,961 | 16,384 | 163,840 | 327,680 | 200,000 |
| Insect | *Drosophila melanogaster* (GCF_000001215.4) | Reference | 7227 | 30,482 | 24,799 | 247,990 | 495,980 | 200,000 |
| Worm | *Caenorhabditis elegans* (GCF_000002985.6) | Reference | Bristol N2 | 28,146 | 21,187 | 211,870 | 423,740 | 200,000 |
| Fungi | *Saccharomyces cerevisiae* | Reference | S288C | 5848 | 4772 | 47,720 | 95,440 | 200,000 |
| Fungi | *Aspergillus oryzae* (GCF_000184455.2) | Representative | RIB40 | 12,074 | 9830 | 98,300 | 196,660 | 200,000 |

| | | | | | | | | |
|---|---|---|---|---|---|---|---|---|
| Fungi | *Neurospora crassa* (GCF_000182925.2) | Representative | OR74A | 10,812 | 8899 | 88990 | 177980 | 200,000 |
| Bacteria | *Bacillus subtilis* (GCF_000009045.1) | Reference | str. 168 | 4174 | 2588 | 25,880 | 51,760 | 200,000 |
| Bacteria | *Escherichia coli* (GCF_000005845.2) | Reference | K-12 substr. MG1655 | 4064 | 2838 | 28,380 | 56,760 | 200,000 |
| Bacteria | *Deinococcus radiodurans* (GCF_000008565.1) | Reference | R1 | 3167 | 2080 | 20,800 | 41,600 | 200,000 |
| Archaea | *Methanosarcina mazei* (GCF_000970205.1) | Representative | S-6 | 3338 | 2063 | 20,630 | 41,260 | 200,000 |
| Archaea | *Haloferax volcanii* (GCF_000025685.1) | Representative | DS2 | 3827 | 2410 | 24,100 | 48,200 | 200,000 |
| Archaea | *Thermococcus gammatolerans* (GCF_000022365.1) | Reference | EJ3 | 2117 | 1346 | 13,460 | 26,920 | 200,000 |

**Table S2 Table of correlation between the results of different disorder prediction algorithms and experimental annotation**

| Algorithm | For the first 150 positions | For the last 150 positions |
|---|---|---|
| IUPred | $\rho = 0.918$; $P < 1 \times 10^{-6}$ | $\rho = 0.768$; $P < 1 \times 10^{-6}$ |
| VSl2B | $\rho = 0.952$; $P < 1 \times 10^{-6}$ | $\rho = 0.829$; $P < 1 \times 10^{-6}$ |
| DisEMBL | $\rho = 0.828$; $P < 1 \times 10^{-6}$ | $\rho = 0.51$; $P < 1 \times 10^{-6}$ |
| MoreRONN | $\rho = 0.935$; $P < 1 \times 10^{-6}$ | $\rho = 0.653$; $P < 1 \times 10^{-6}$ |
| Espritz (D) | $\rho = 0.855$; $P < 1 \times 10^{-6}$ | $\rho = 0.87$; $P < 1 \times 10^{-6}$ |
| Espritz (N) | $\rho = 0.45$; $P < 1 \times 10^{-6}$ | $\rho = 0.066$; $P = 4.19 \times 10^{-2}$ |
| Espritz (X) | $\rho = 0.923$; $P < 1 \times 10^{-6}$ | $\rho = 0.805$; $P < 1 \times 10^{-4}$ |
| IUPred 2A (long) | $\rho = 0.916$; $P < 1 \times 10^{-6}$ | $\rho = 0.741$; $P < 1 \times 10^{-4}$ |
| IUPred 2A (short) | $\rho = 0.799$; $P < 1 \times 10^{-6}$ | $\rho = 0.639$; $P < 1 \times 10^{-4}$ |
| Spot disorder Single | $\rho = 0.935$; $P < 1 \times 10^{-6}$ | $\rho = 0.623$; $P < 1 \times 10^{-4}$ |

*Note:* This table represents the correlation between the proportions of disordered residues predicted by different disorder prediction algorithms with that calculated based on experimental order/disorder annotation. For this test, we calculated proportion of disordered residues up to the first and last 150 positions (separately) of DisProt protein dataset (*i*) once following order or disorder annotation as suggested in this database based on experimental methods and (*ii*) then based on the prediction by each of algorithms used for our main analysis. To calculate the proportion of disordered residues position-wise, we aligned the proteins once from start to end and then from end to start. Proportion of disordered residues at any position is calculated as the number of disorder residues (either annotated by DisProt or predicted by algorithm) at that position to the number of proteins in the dataset. Currently, there are 803 sequences in the DisProt database. Here we considered proteins more than 150 residues in length (a total of 604 proteins). Following non-parametric distribution, here we performed Spearman's Rank correlation analysis and denoted correlation coefficients with ρ and significance level with *P*-values.
